# Building Neurovascular tissue from autologous blood for modeling brain activity

**DOI:** 10.1101/2024.10.16.617820

**Authors:** Rhythm Arora, Alka Bhardwaj, Naresh K Panda, Sanhita Sinharay, Jaimanti Bakshi, Ramandeep Singh Virk, Sanjay K Munjal, N. Banumathy, Gyanaranjan Nayak, Sourabh K Patro, Anuradha Sharma, Reena Das, Tulika Gupta, Sanjay Kumar Bhadada, Arnab Pal, Nabhajit Mallik, Rimesh Pal, Madhumita Premkumar, Ritin Mohindra, Ravi Dixit, Meenakshi Pal, Sajid Rashid, Maryada Sharma

## Abstract

There are no faithful individualized stem cell-based bioengineered neuro-vascularized models that can recapitulate the physiological hemodynamic phenomenon of neuro-vascular coupling (NVC)-the principal behind BOLD (blood oxygen level-dependent) signal in functional neuroimaging, thereby dissuading the research in exploring the brain activity-based investigative studies in neurological/neurosensory diseases. This encouraged us to establish a preclinical optoacoustic (Hb/dHb hemoglobin/deoxyhemoglobin) imaging-competent *in vitro* neuro-vascularized model by employing a novel cellular reprograming PITTRep (Plasma Induced Transcriptomics/ epi-Transcriptomics Reprograming) approach. The current reprograming approach is based on coaxing autologous blood components to ecto-mesodermal lineage intermediates that can subsequently self-pattern into neurovascular tissue by harnessing the hemorheological properties of RBCs. The nature of blood flow is non-Newtonian and is a function of RBC concentration /haematocrit when they flow through the regions of low shear rates as seen in cerebral microcirculation. The current reprograming approach is a modification of our previous cellular reprograming approach that employed a Newtonian plasma fluid. The autologous blood-derived neurovascular tissue is free from exogenous genetic modification, external growth factors, and induced pluripotent stem cell (iPSC) derivation. This model uniquely integrates functional vasculature and neurogenesis.

The current reprogramming approach resulted (in part) serendipitously while testing a potential (yet completely unexplored) hypothesis of haemodynamic reprograming by leveraging the fluid mechanic feature of blood erythrocytes as seen in thrombus formation during cerebral ischemic stroke, that is characterized by physiologically intriguing yet clinically meaningful neurological recovery (neuroplasticity) during an early time window. The current study attempted to induce “a post stroke-like model” of adult neurogenesis with functional synaptogenesis by instructing autologous blood components into thrombus formation through incorporation of erythrocytes in varying concentrations. We tried to instruct adult neurogenesis and neuroplasticity (a relatively non-resilient phenomenon under *in vitro* conditions) by co-induction of a neuro-vascular niche (NVN). These NVNs are marked by dendrites, synapses, astrogliosis, microglia activation, and growth factor signaling, thus phenocopying molecular and cellular aspects of post-stroke recovery window.

The induction of neuro-vascularized niches and functional neuro-vascular coupling (NVC) was characterized by confocal microscopy, scanning electron microscopy, proteomic profiling, and Hb/dHb spectra based optoacoustic imaging. The blood thrombus formation was checked by rotational thromboelastometry (ROTEM), and switching of adult-to-embryonic hemoglobin was confirmed by routine hemoglobin typing. We also attempted to establish patient-specific neuro-vascularized niches from autologous blood of sensorineural hearing loss (SNHL) patients. The individualized neovascularised tissues are intended to be employed for investigating deregulated synaptic plasticity/ long term potentiation underlying poor auditory comprehension outcomes in school going kids suffering from SNHL that greatly compromises their academic performance and socio-behavioural-cognitive development. The attendant multiomics of patient-specific NVNs may have potential implications in developing stem-cell based therapies for neurosensory and cerebrovascular diseases.

## Introduction

The complexity of human brain cytoarchitecture is tough to mimic in any in-vitro model system. Although there has been an enormous growth in the field of stem cell biology especially the area concerning brain organoids in past decade but still the structural integrity and functional connectivity of the human brain hasn’t been fully recapitulated in any of the brain organoid models, making it a challenge for the researchers to model and understand complex neurological and sensory neural disorders. The brain is a highly vascularized organ, the need for this can be understood by the extensive functional neuronal signalling and regulation it has to perform which requires constant oxygen supply. From the developmental perspective, the multi-step process of neurogenesis involving migration and differentiation of neuroblast cells, which culminates with the integration of mature neurons into the neural circuit is guided and coordinated by the cues coming from the developing vasculature (Bjornsson et al., 2015). Due to absence of active blood flow the current brain and cortical organoids or assembloids have insufficient maturation, lack of long-term potentiation (LTP) or synaptic plasticity hence these fall short of fully replicating the functionality of brain tissue. This limitation hinders their use in studies involving brain activity, particularly sensory-evoked neural responses. Therefore, it is envisioned that the generation of in vitro vascularized brain tissue or organoid could represent a potential framework to investigate the altered neurosensory and neuromotor molecular pathways reflecting the brain activity.

Its been more than a decade today since the first whole brain organoid was created using hPSC’s in Lancaster’s lab (Lancaster et al., 2013), ever since then there has been a remarkable progress in this area but there is a huge scope for improvement when it comes to addressing the limitations of these organoids. Due to the absence of functional vasculature and supporting cells such as microglia and astrocytes, these organoids do not fully replicate the cytoarchitecture and functionality of in vivo brain tissue and lack maturation. Over time, they tend to develop a necrotic core, primarily caused by cellular stress from inadequate oxygen and nutrient perfusion. The findings by Bhaduri et al., (2020) indicate the increased activation of glycolysis, ER stress pathways and distorted sub-specification in the cells in the brain organoids (Bhaduri et al., 2020). This problem has been addressed by employing several strategies such the use of spinning bioreactors (Lancaster and Knoblich, 2014) or culturing the cortical organoid slices on air-liquid interface (Qian et al., 2020; Giandomenico et al., 2021), to ensure improved oxygen supply using small molecule cocktail like CEPT (chroman 1, emricasan, polyamines, trans-ISRIB) to improve cell survival and promote cytoprotection (Ryu et al., 2023), transplanting the organoid to improve organoid maturity, complexity and functioning (Jgamadze et al.,2023; Cao et al., 2023; Wang et al., 2023).

The brain is a complex structure comprising of the cerebrum, cerebellum and the brain stem which are home to different types of neurons and non-neuronal cells. The connections of neurons (the synapses and tracts) between these areas are responsible for its sophisticated functioning. Although unguided protocols generate diverse cells types and brain regions but they lack functional vasculature, possess significant variation and lack reproducibility. On the other hand the guided protocols create region specific structures with great reproducibility but with limited cell types (Chiaradia et al., 2023) and absence of vascular niches, since they don’t resemble the morphology and functioning of the brain tissue therefore they can’t be robustly used for modelling various neurosensory, neurodevelopmental and neurodegenerative disorders. Assembloid approach of generating organoids provides a partial solution to this problem. Region-specific organoids can be fused together to model these interconnections like ventral and dorsal forebrain organoids are co-cultured to model interneuron migration (Bagley et al., 2017; Samarasinghe et al., 2021), cortical-thalamus organoids to model thalamus dysfunction related to psychiatric disorders (Angulo Salavarria et al., 2023), cortical striatum assembloids for modeling neural circuit dysfunction in the Autism spectrum disorders (ASD) and schizophrenia (Miura et al., 2022 and 2020). Using this approach, several attempts have been made to form vascularised cortical organoids by co culturing neuroepithelium cells with endothelial cells which has resulted in the formation of vascular-like network (Sun et al., 2022 and 2023; Nwokoye et al., 2024) but these protocols lack active flow.

Bioengineering has made important contributions to resolving this challenge by simulating the neurovascular unit and blood-brain barrier (BBB) with microfluidic platforms. Early models comprised endothelial cells (ECs), astrocytes, and pericytes, but no neurones, making them unsuitable for thorough research of brain activity. Later models included neurones, glial cells, and ECs, albeit they frequently used cells from different species, reducing their relevance to human biology. Recent research has focused on humanised models that use iPSC-derived neurones, astrocytes, and ECs. These models demonstrate functional connectivity between neurones and the BBB, as determined by procedures such as trans-endothelial electrical resistance (TEER) and dextran permeability experiments. Despite these advancements, challenges remain, such as variability in vascular network formation, the non-physiological nature of the materials used (e.g., collagen-I), and reproducibility issues across different models (Caffrey et al., 2021). The next generation technology involves building brain on a chip (BoC) model devices, microphysiological systems offer precise control over the physical and biochemical conditions of brain tissues, which enables the study of brain function and disease at an unprecedented level. However, several challenges remain before BoC models can reach their full potential. One of the main hurdles is the need to co-culture multiple cell types, including microglia and immune cells, which is difficult with current systems. There is also a need for developing universal culture media and methods for scaling up these systems to match organ size, which is critical for drug testing and disease investigative studies. The use of foreign biosynthetic materials for structural support and fabrication makes the system less natural, hinders reproducibility, decreases cost effectiveness hence making it a challenge for its direct use in the field of personalised medicine and as the preferred adopted preclinical methodology (Rodrigues et al., 2024; Amirifar et al., 2022). Moreover the functional inputs and output from any BoC would require physiological validation from animal and clinical studies reducing its direct translational impact (Amirifar et al., 2022)

Neuronal activity is the key feature to determine the functional aspect of the developed brain organoids (Mulder et al., 2023) and neurovascular unit. Calcium imaging and patch clamp (Miura et al.,2020; Eura et al. 2020; Xiang et al.,2019; Andersen et al., 2018;Birey et al., 2017; Lancaster et al., 2013) are most frequently used assays for measuring the neuronal activity in brain organoids. Excitatory and inhibitory neurons in cortical organoids are functional, but unsurprisingly less mature, compared to adult neurons (Zourray et al., 2022; Foliaki et al.,2021; Birey et al., 2017). Because cortical organoids model early to mid-embryonic development, electrophysiological characterization often reveals highly variable functionality and maturation in randomly selected cells (Qian et al., 2019). Despite their variability, the basic electrophysiological properties of neurons, including resting membrane potential, resistance and the amplitude and frequency of spontaneous postsynaptic currents, exhibit statistically significant maturation trends as organoids age (Porciúncula et al., 2021; Qian et al., 2016). Synaptogenesis is also seen in the developing organoids when examined using immunostaining or electron microscopy (Yakoub and Sadek, 2019; Birey et al., 2017; Quadrato et al., 2017; Sloan et al., 2017). Neural circuit formation and integration reflects about the complex activity developing in the brain organoids which is very essential for recapitulating any disease physiology. In a recent study by Osaki et al., 2024 they have been successful in recording the complex activity and short term plasticity in cerebral organoids (Osaki et al., 2024). But due to insufficient maturation and lack of long term potentiation they can’t completely replicate the functionality of brain tissue indicating scope for the attaining improvement.

Brain activity is correlated with blood flow and oxygenation driving corresponding blood flow increase over the locally activated brain-region, a phenomena known as neuro-vascular coupling. NVC is a delicate process of regulating spatio-temporal local cerebral blood flow which involves a coordinated response of neurons, glial cells and vascular cells (Caffrey et al., 2021). The lack of active blood flow in the existing brain organoid protocols constraints them from mimicking the natural physiology of the brain tissue. Microglia are the brain’s resident immune cells and are crucial for brain development (Zourray et al., 2022). They help with synaptic maturation, transmission, and pruning, shaping the brain’s cortical networks thus their presence is vital for recapitulating the microenvironment in the developing organoids (Fagerlund et al., 2021; Zhang et al., 2023). Astrocytes and oligodendrocytes play crucial roles in maintaining the health and function of neurons hence they must co-exist for attaining maturity in neuronal function in the brain organoid models (Acharya et al., 2024). The aspect of using pluripotent stem cells and inducing reprogramming by transgene overexpression or morphogenetic factors presence in special culture media makes these methods cost ineffective. Recent studies have also reported the prevalence of acquired cancer-related mutations in human pluripotent stem cell lines and their differentiated derivatives which affect the results derived in basic research and disease modeling (Lezmi et al.,2024). The more concerning aspect is the potential presence of such cancer-driving mutations in cells designated for clinical application(Lezmi et al., 2024).

Sensory neural hearing loss (SNHL) results from the loss of function or impairment of the sensory hair cells in cochlea or an auditory neuropathy of the cochlear nerve or the brain’s central processing centers (Tanna et al., 2024, Liu et al., 2020). Congenital SNHL affects approximately one to two out of every thousand live births, establishing it as among the most prevalent congenital conditions (van Beeck Calkoen et al, 2019, Smith et al., 2005). A cochlear implant (CI) is currently the only FDA-approved biomedical device that can restore hearing for the majority of individuals with severe-to-profound sensorineural hearing loss (SNHL). The burden of poor and highly variable speech perception outcomes are reported to be a big hindrance to the success of CI (Holden et al., 2013). There are various intrinsic and extrinsic factors that have been identified which contribute to these divergent outcomes which include gender, IQ, socio-economic status, age of onset of hearing-loss, or associated disabilities (cognitive or developmental) (intrinsic factors) (Geers et al., 2007); age at implantation, educational and rehabilitative communication strategy, hearing aid use, course of CI experience (extrinsic factors) (Zeng, 2004; Tobey et al., 2004). However, the mechanistic basis of how these factors manifest these variable outcomes has not been discovered. The most acceptable hypothesis in this regard is the contribution from top-down neurocognitive factors driving neuroplasticity in cortical regions, as other biological and audiological factors could not fully attribute to the reported divergent outcomes in CI patients (Beckers et al., 2023).

## Results

This figure illustrates the differentiation of mesodermal lineage progenitors into vessel-like structures using the PITTRep approach.

In Panel IA, the process of mesodermal progenitor induction from whole blood cells is showcased. The PITTRep method activates specific transcriptomic pathways that guide these cells toward a mesodermal fate. This activation leads to the emergence of mesodermal and endothelial cell precursors, which are key contributors to the formation of early vascular structures. The image demonstrates the early stages of lineage commitment, where blood-derived cells begin to express markers characteristic of the mesodermal lineage.

Panel IB highlights the formation of “red vessel-like” structures under varying experimental conditions. These structures are surrounded by primitive mesodermal progenitors, suggesting that early vascular organization is occurring within these cultures. The vessel-like formations appear to be supported by the surrounding progenitors, which are integral to the stabilization and maturation of these nascent vascular networks. The red colour indicates the presence of RBCs in developing vasculature, while the surrounding mesodermal cells provide a supportive environment for further differentiation.

In Panel IC, the differentiation of meso-ectodermal lineages into neuro-vascular progenitors is depicted under specific conditions, denoted as C1 and C3. These conditions have been optimized to promote the formation of both mesodermal and ectodermal progenitors, which together contribute to the generation of neurovascular tissues. The image illustrates that under condition C1(160) and C3(160), differentiation into mesodermal-endothelial progenitors is particularly robust, indicating that specific environmental cues can direct the formation of endothelial cells that are critical for vascular development.

Panel ID provides further validation of these developing structures through confocal microscopy, which allows for high-resolution imaging of the structural organization and marker expression within the cultures. In this panel, VE-cadherin (stained in red) is prominently expressed, marking the formation of vessel-like networks that resemble early blood vessels. VE-cadherin is a key protein in endothelial cells, facilitating cell-to-cell adhesion and signaling within developing vascular tissues. The presence of VE-cadherin confirms the formation of endothelial structures that are integral to the developing vasculature.

Additionally, c-Kit (also in red) highlights the presence of common neuro-melanin progenitors alongside hematopoietic stem cells, indicating a shared developmental pathway between these progenitor types in the PITTRep model. c-Kit is a stem cell marker that plays a crucial role in the regulation of cell proliferation and differentiation, particularly in neuro-melanin and hematopoietic lineages. The co-expression of c-Kit in these progenitors suggests that the PITTRep approach successfully induces a mixed population of progenitor cells capable of contributing to both vascular and neural tissues.

Moreover, alpha-SMA (smooth muscle actin) and CD31 (PECAM-1) staining provide additional confirmation of vascular development. Alpha-SMA marks the presence of vascular smooth muscle cells, which are essential for the stabilization and contraction of blood vessels. CD31 is a well-known marker of endothelial cells, confirming the identity of the cells lining the developing vessel-like structures. Together, these markers indicate that the PITTRep method successfully directs the differentiation of blood-derived cells into a complex, multi-lineage system that includes both endothelial and smooth muscle progenitors, facilitating the formation of functional vascular networks.

This figure provides analysis of the proteomic and hemoglobin profiling of neurovascular niches that were developed using the PITTRep approach. The figure details the transition from adult to embryonic hemoglobin types, reflecting significant changes in cellular and molecular phenotypes associated with vascular mesodermal lineage differentiation and blood vessel development.

In Panel IIA, the focus is on the induction of embryonic and fetal hemoglobin within the neurovascular niches, highlighting a switch from adult to embryonic hemoglobin types. The data presented here demonstrate that the PITTRep approach stimulates a reprogramming of hemoglobin expression, leading to the activation of genes that are typically associated with embryonic and fetal development. This shift is indicative of a developmental recapitulation in the system, where progenitor cells within the neurovascular niche adopt characteristics of earlier developmental stages, including the expression of hemoglobin types that are typically seen during fetal development.

Further validation of this switch in hemoglobin expression is provided in Panel IIE, which shows the results of hemoglobin typing performed using High-Performance Liquid Chromatography (HPLC). The HPLC analysis reveals distinct peaks corresponding to fetal hemoglobin (Bartz), with the red arrow in the panel pointing to the specific peaks that indicate a significant presence of fetal hemoglobin in the culture. This is a critical observation, as the presence of fetal hemoglobin suggests that the neurovascular niches are undergoing a process of reprogramming that mimics early developmental stages, potentially enhancing the plasticity and regenerative capacity of the progenitor cells within the niches.

In Panel IIB, the proteomic analysis focuses on erythrocyte homeostasis and the regulation of ion channels in erythrocytes, which are essential for maintaining the balance of ions and water within red blood cells. The bar graph presented in this panel quantifies the expression levels of key proteins involved in erythrocyte function, suggesting that the neurovascular niches are not only supporting the formation of vascular structures but are also fostering the development of blood cells that exhibit functional properties associated with erythrocyte biology.

A more detailed bioinformatic analysis of the proteomic data is presented in Panel IIB1, where the ShinyGo v0.80 platform is used to explore the molecular pathways that are enriched in the neurovascular niches. This analysis highlights several key pathways involved in erythrocyte development, myeloid progenitor differentiation, and hematopoiesis. The activation of these pathways suggests that the PITTRep model is capable of supporting hematopoietic differentiation, further validating the system as a robust model for studying blood and vascular development. The proteomic data align with the morphological and functional changes observed in the niches, demonstrating that the PITTRep approach effectively induces the differentiation of mesodermal progenitors into vascular and hematopoietic lineages.

In Panel IIC, the proteomic data specifically focusing on vasculature development is illustrated. This panel demonstrates the formation of vascular structures within the neurovascular niches, corroborating the earlier findings shown in Figure I. Using the Appyter, R language-based platform, the analysis reveals significant processes related to blood vessel sprouting and hematopoiesis, with a highly significant p-value < 0.01, indicating the robustness of the vascular development in the PITTRep model. The data highlight key proteins involved in the formation of blood vessels, including those that regulate endothelial cell migration, proliferation, and organization into tubular structures, all of which are essential for the formation of functional vasculature.

**Figure I:**
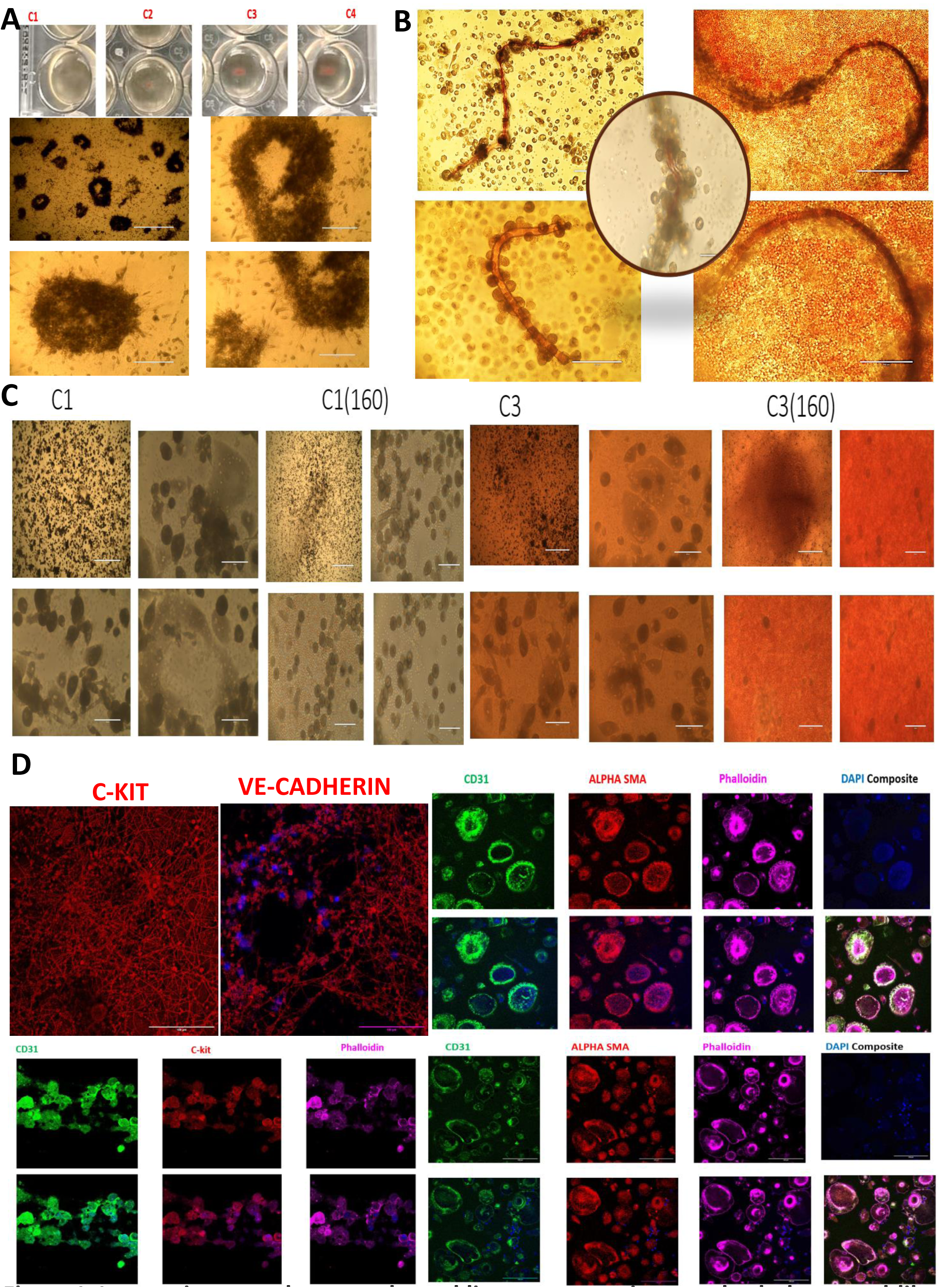
Instructing Vascular Mesodermal Lineage Progenitors and Tubular Vessel-Like Structures by Employing a Novel PITTRep (Plasma Induced Transcriptomics and epi-Transcriptomic Reprogramming) Approach.

Further validation of these findings is provided in Panel IID, where the vascular patterns observed in the neurovascular niches are visualized.

The induction of embryonic and fetal hemoglobin, coupled with the development of functional vasculature, suggests that the PITTRep approach effectively recapitulates key aspects of early developmental processes, enhancing the regenerative potential of the neurovascular niches.

### Panel IIIA: Developmental Phases of Neurogenesis in Culture

This panel highlights the various stages of neurovascular progenitor development in a culture plate, showing a time-based progression:

- **Upper Phase**: In the initial stages, **neuroepithelial cells** begin to form, which are among the first progenitors of the nervous system. As the culture matures, these neuroepithelial cells differentiate into **radial glia progenitors**. Radial glial cells are essential in brain development, acting as both scaffolding for neuron migration and progenitors for neurons and glia.

**Radial Glia Progenitors** give rise to **intermediate, basal, and apical progenitors** as time progresses. These progenitor types represent different layers and roles in neurogenesis, with the intermediate progenitors leading to various neuronal subtypes, while the basal and apical progenitors are involved in forming the supportive structure of the developing neuronal tissue.

- **Lower Phase**: The lower part of the panel shows further neuronal differentiation. Here, **neurons** begin to mature and organize into structured **neuronal tissue**. This process results in the development of a functional neuronal network, integrating into the neurovascular structure.

### Panel IIIB: Visual Development of Neurovascular Niches

In this panel, a photograph of the culture plate is shown at **day 8** of development. The image captures the formation of neurovascular niches within the plate. These niches are regions where neurogenesis and angiogenesis are taking place simultaneously, mimicking natural tissue environments. By day 8, the progenitor cells have differentiated significantly, leading to visible neurovascular organization, which is a hallmark of successful neurogenesis in culture.

### Panel IIIC: Brightfield Imaging of Meso-Ectodermal Progenitors

This brightfield image provides a clear view of the **meso-ectodermal progenitors** as they support the development of neurovascular tissue. These progenitors originate from the induced blood cells and differentiate unguided into both mesodermal (vascular) and ectodermal (neuronal) lineages, a process critical for the proper formation of neurovascular structures.

### Panel IIID: Validation through Confocal Imaging from condition C1 to C4

To validate the results, confocal imaging is employed using specific markers:

- **KI-67 (Red)**: This marker identifies **proliferative cells** within the culture, highlighting areas of active cell division. The presence of proliferative cells is essential for neurogenesis, as it indicates the continuous formation of progenitors.
- **Beta 3 Tubulin (Green)**: A marker for **early neuronal progenitors**, indicating the early stages of neuron formation. Beta 3 Tubulin is typically expressed in immature neurons, and its presence here shows the successful differentiation of neural progenitors from the blood-derived cells.

### Panel IIIE: CD34 and TCF21 Expression

This confocal image reveals the expression of:

- **CD34 (Green)**: A marker for early neuronal progenitors, and hematopoietic and endothelial progenitors. Its expression here reflects the transition of progenitor cells into a neural lineage.
- **TCF21 (Red)**: A marker showing the activation of the **Wnt signaling pathway**, which plays a crucial role in regulating cell fate, proliferation, and differentiation. The activation of Wnt signaling is indicative of the initiation of neurogenesis and tissue patterning.

### Panel IIIF: FGF2 Signaling

This panel focuses on the expression of **Fibroblast Growth Factor 2 (FGF2)**:

- **FGF2 (Green)**: The presence of basic FGF2 is essential for the development of neurovascular tissue. It plays a key role in the proliferation and differentiation of neural progenitors.
- **Phospho FGF2 (Red)**: The phosphorylated form of FGF2 represents its active state, signaling the pathway’s involvement in neurogenesis. The combination of FGF2 and its phosphorylated form shows the importance of **FGF signaling** in maintaining the development of both neural and vascular components in the culture.

### Panel IIIG: Ectodermal Progenitor Marker

This panel displays the expression of **early ectodermal progenitor markers**, which confirm the commitment of a subset of the progenitor cells to an ectodermal lineage. This is a critical step in neurogenesis, as the ectoderm gives rise to the nervous system during development.

### Panel IIIH: Expression of Stem Cell Transcription Factor OCT4

In this final panel, the expression of the transcription factor **OCT4 (Green)** is visualized. OCT4 is a pluripotency marker and is essential for maintaining stem cell properties. Its presence in the culture indicates the retention of stem cell-like characteristics in some cells, which is necessary for the continued generation of progenitors during the neurogenesis process.

This figure presents the detailed characterization of neurogenesis using various imaging and staining techniques. Following the observations made in Figure III, where brightfield imaging and confocal validation confirmed the early stages of neuronal development, Figure IV offers further validation through multiple methodologies, providing a deeper insight into the structural and molecular aspects of neuronal differentiation and vascular development.

**Figure II:**
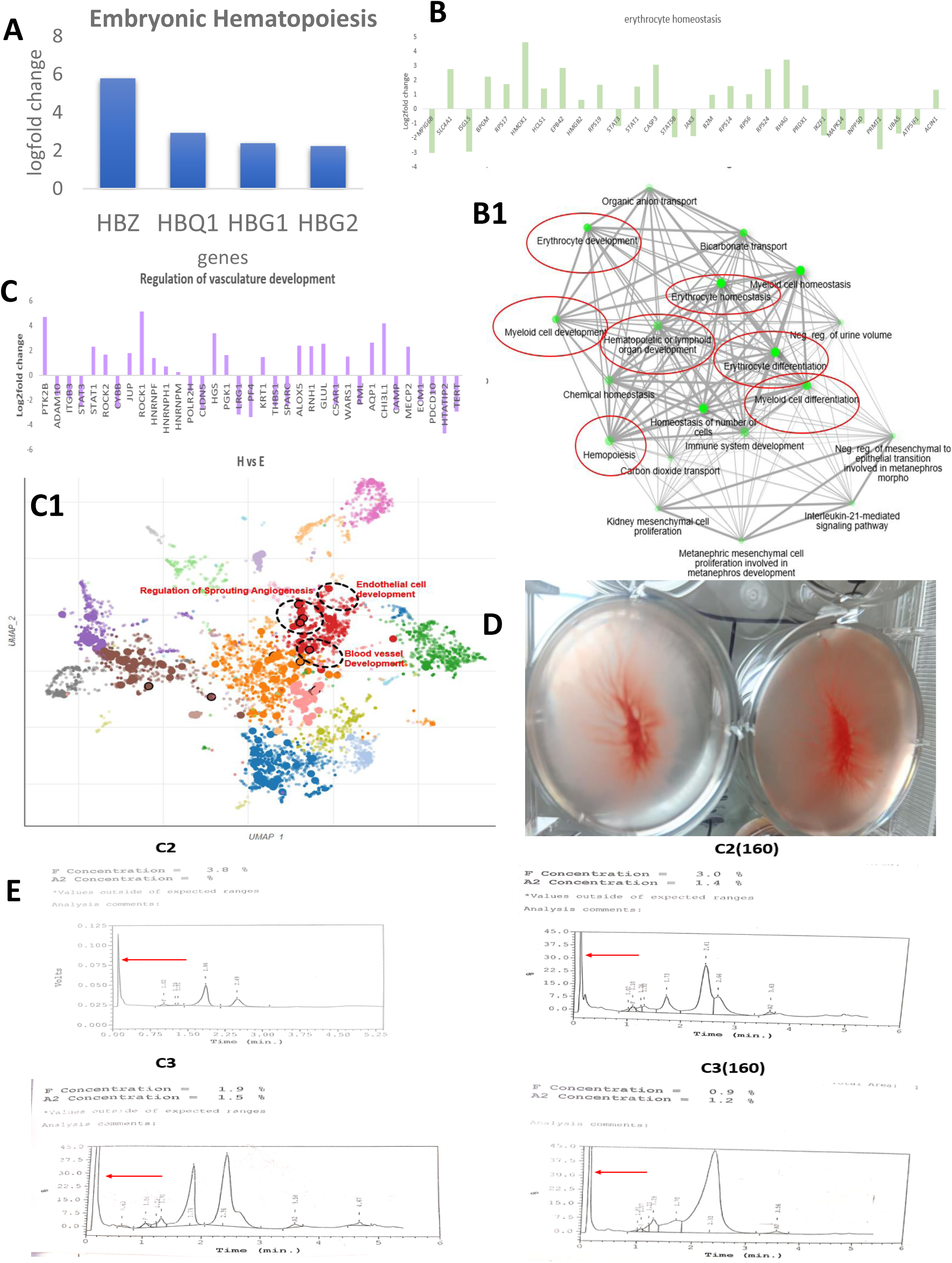
Proteomics Profiling and Hemoglobin Typing of Neurovascular Niches Indicating the Induction of Vascular Mesodermal Lineage Progenitors and the Switch from Adult to Embryonic Hemoglobin Types.

**Figure III:**
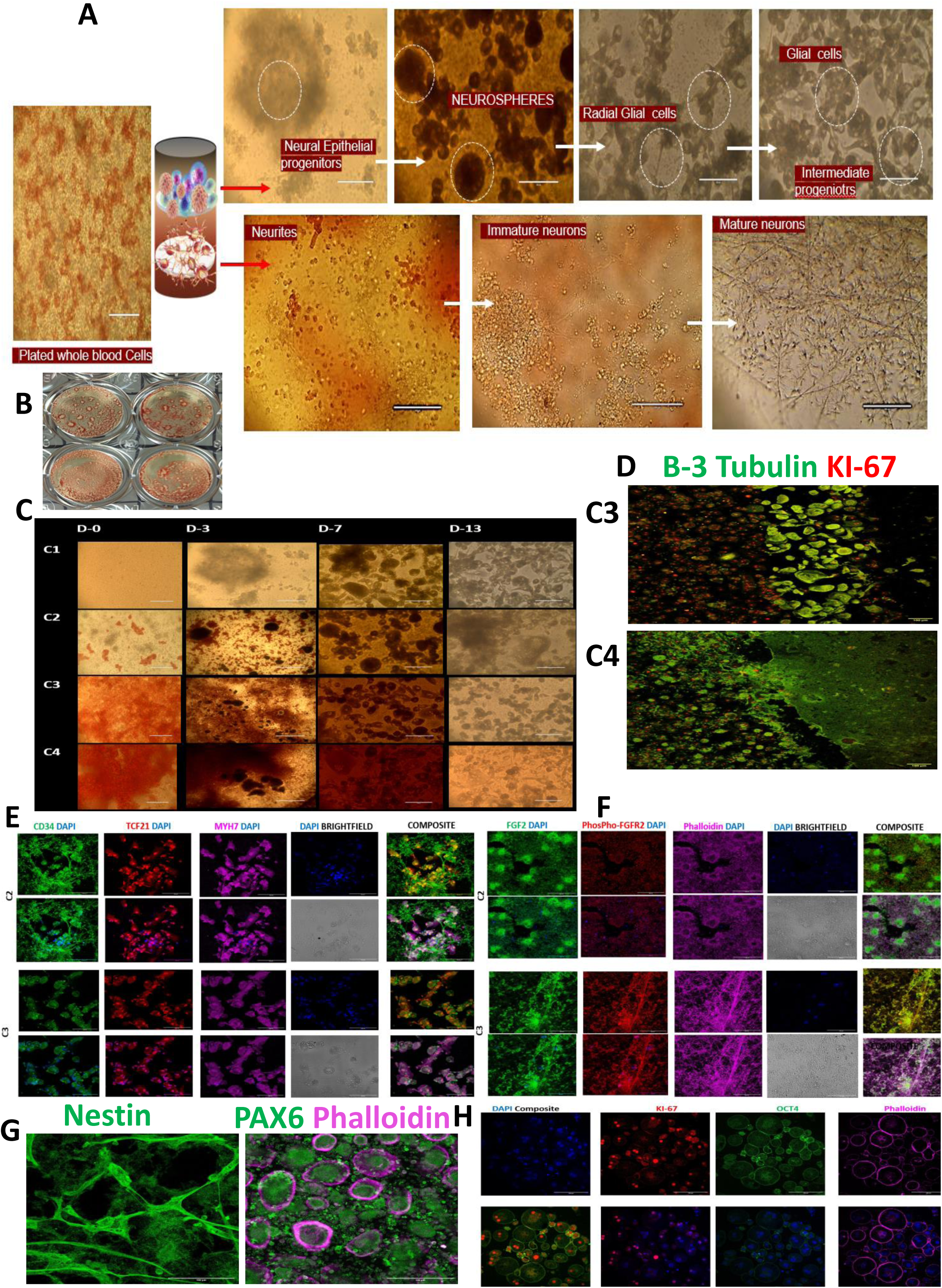
Instructing Neurogenesis from Autologous Blood by Employing the PITTRep Approach. Provides the data for neurogenesis, and induction of both mesodermal and ectodermal lineages, along with neural crest stem cell development. This process ultimately leads to the formation of functional neurovascular tissue.

**Figure IV:**
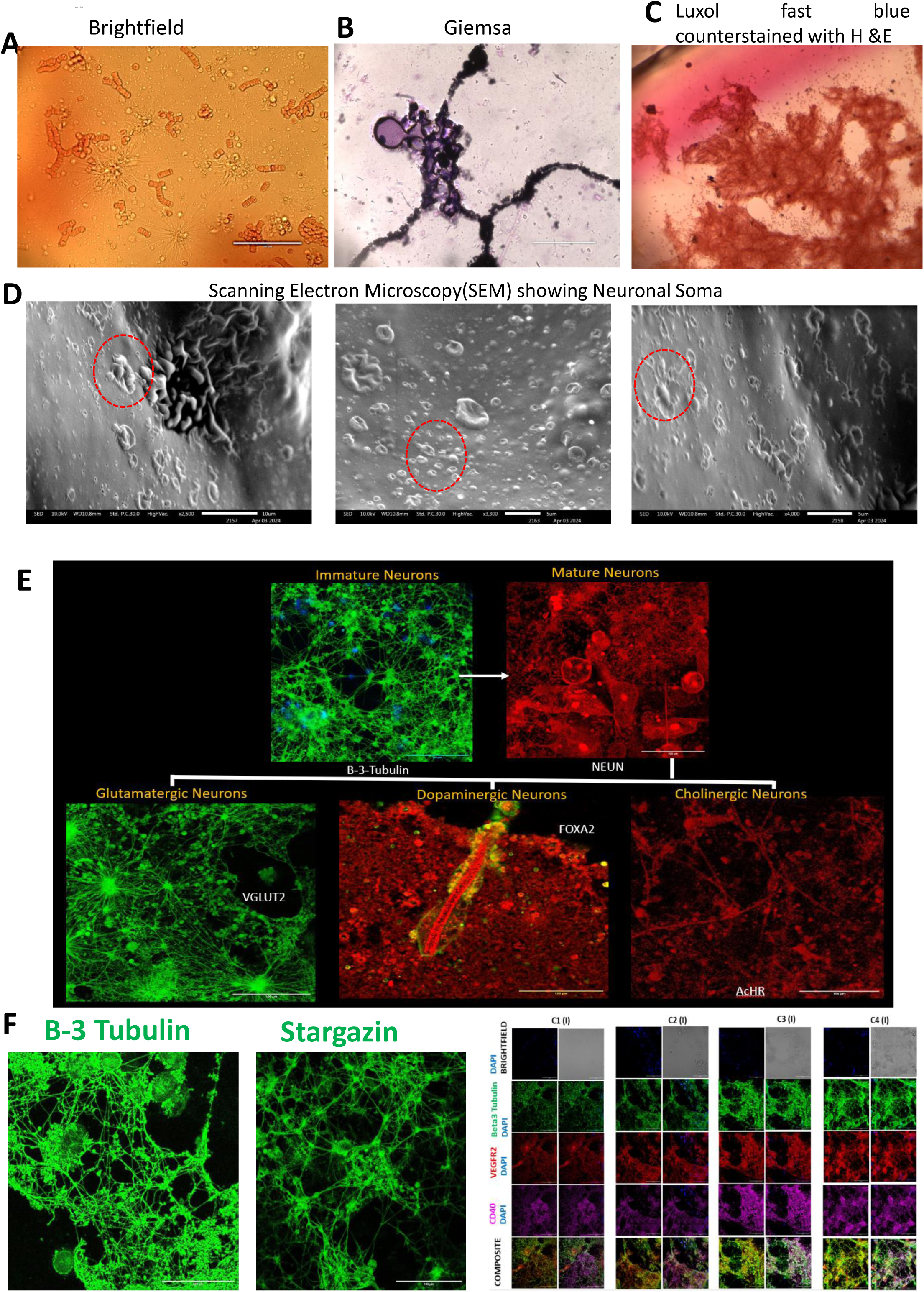
Characterization of Neurogenesis by Scanning Electron Microscopy (SEM), Luxol Fast Blue Staining Counterstained with Hematoxylin & Eosin (H&E), and Confocal Microscopy in the PITTRep Model.

In Panel IV A, brightfield imaging shows neurons developing in the culture. The image captures the early stages of neuronal differentiation, where neuronal cells begin to exhibit structural features typical of developing neurons. These cells appear with distinctive neuronal morphologies, with the growth of neurites extending from the soma (cell body), signifying the early establishment of neuronal networks.

Panel IV B presents Giemsa staining, which clearly visualizes the neuronal somas in culture. The staining highlights the differentiating neuronal cell bodies, which are actively generating multiple neurites. These neurites, the precursor extensions of axons and dendrites, are crucial for synaptic communication, and their appearance signifies the progressive differentiation of neurons in the culture.

In Panel IV C, Luxol fast blue staining, counterstained with Hematoxylin and Eosin (H&E), shows the presence of pyramidal neurons and the developing cerebral cortex. Luxol fast blue indicates the emergence of organized neuronal structures, including the pyramidal neurons that are essential for higher-order brain functions. This panel demonstrates that the PITTRep model supports the maturation of neurons into functional brain-like regions, replicating the cellular organization seen in the cerebral cortex.

Panel IV D features Scanning Electron Microscopy (SEM) images, offering high-resolution views of the neurovascular niches formed in the culture. The SEM captures the intricate details of neuronal soma development at different stages, revealing the emergence of neurites from the soma. These high-magnification images provide structural validation of the neurogenesis observed in the earlier panels, demonstrating the complex three-dimensional architecture of developing neuronal networks within the neurovascular niches.

In Panel IV E, confocal microscopy further validates the developing neurons by using specific molecular markers. The expression of beta-3 tubulin (green) indicates the presence of immature neurons. This protein is commonly expressed during early neuronal differentiation and is crucial for the assembly of the cytoskeleton in developing neurons. NeuN (red) marks postmitotic neurons, indicating neurons that have exited the cell cycle and are in the process of maturation. Together, these markers confirm the differentiation of progenitor cells into neurons, progressing through immature stages to more defined neuronal identities.

Additionally, this panel reveals the presence of specific neuronal subtypes, validated by targeted markers:

- VGLUT2 (vesicular glutamate transporter 2) marks glutamatergic neurons, which are responsible for excitatory neurotransmission in the brain.
- FOXA2 (in red) is a critical transcription factor for the development of dopaminergic neurons, which play a key role in the regulation of movement and reward mechanisms, highlighting the emergence of neurons related to the substantia nigra.
- AcHR, a nicotinic acetylcholine receptor, indicates the presence of a recptor for cholinergic neurons, which are important for neurotransmission related to muscle activation, cognition, and memory.

Panel IV F continues the confocal analysis, providing further insights into the molecular features of developing neurons. The panel shows positive staining for immature neurons alongside the expression of stargazin, a protein crucial for the regulation of calcium channels in glutamatergic neurons. Stargazin is known to increase in active glutamatergic neurons, indicating that these neurons are not only differentiating but are also functionally active. The expression of stargazin is critical for the regulation of synaptic strength and plasticity in glutamatergic circuits.

Furthermore, this panel demonstrates the active development of neurovascular tissues through the expression of specific markers from condition C1 to C4:

- Beta-3 tubulin (green) confirms the ongoing maturation of neurons.
- VEGFR2 (red), a marker for vascular endothelial growth factor receptor 2, indicates the development of vascular components within the neurovascular niches.
- CD40 (magenta), typically associated with microglia activation, may play a role in the regulation of neuronal-vascular interactions during neurogenesis.
- This figure focuses on the proteomic analysis of neurovascular niches generated using the PITTRep model, highlighting the induction of **gliogenesis** and **forebrain development**. The figure underscores the critical role of glial cells in establishing functional neurovascular tissue, a feature often difficult to replicate in vitro. In the PITTRep model, gliogenesis appears to follow physiological processes akin to early human brain development, which is essential for the creation of active neurogenesis and neurovascular niches.
- **Panel VA** presents a brightfield image illustrating a population of **astrocytes** within the culture. Astrocytes are encircled in red, indicating their abundance in the neurovascular niches. Astrocytes are pivotal for maintaining homeostasis, regulating blood-brain barrier permeability, and supporting neurons. Their presence in such significant numbers indicates that the PITTRep model successfully induces glial cell development, an essential factor for neurovascular functionality.
- In **Panel VB**, the brightfield imaging captures the presence of **early radial glial cells**, which are the foundation for brain development in primates. These radial glia serve as scaffolds for newly born neurons, guiding their migration during early brain development. The PITTRep model shows a rich population of these progenitors, signaling that the system replicates key aspects of **human brain development**. The radial glia interact closely with other progenitor cells within the culture, promoting organized neurogenesis.
- **Panel VC** provides a detailed analysis of **high-throughput proteomic data** using the bioinformatics tool **WebGestalt 2024**, revealing the upregulation of pathways associated with gliogenesis. This upregulation indicates the model’s ability to support not only the neuronal aspect of brain development but also the essential glial component. Gliogenesis, the formation of glial cells, is crucial for the functional integration of neurons into neural circuits, providing metabolic support, modulating synaptic activity, and maintaining homeostasis within the nervous system.
- In **Panel VD**, further bioinformatic analysis using the **Enrichr** platform demonstrates the enrichment of pathways related to **forebrain development** and the differentiation of **forebrain neurons**. The data suggest that the neurovascular niches developed through the PITTRep model actively promote the formation of glial progenitors, which are particularly important for forebrain development. The forebrain is a region rich in glial cells, particularly astrocytes, which play key roles in synaptic regulation and neurovascular interactions. The presence of glial progenies in the system thus aligns with the overall goal of replicating **neurogenesis** in vitro.
- **Panel VE** presents a Umap generated through analysis on the **Appyter bioinformatic platform**, displaying the abundance of proteins involved in **gliogenesis** and the regulation of **MSX2(VF)**, a master transcription factor critical for cerebral development. MSX2 regulates key developmental processes, including neural tube formation and forebrain patterning. The high expression of MSX2 in the PITTRep system underscores the model’s ability to replicate early cerebral development, particularly in the forebrain.
- The confocal validation presented in **Panel VG** from condition C1 to C4 provides a closer look at the proteins identified through high-throughput data analysis, confirming their presence in the neurovascular niches. The panel shows strong expression of **GFAP** (glial fibrillary acidic protein) (green), a well-known marker of **astrocytes**, within the neuronal-vascular niches. GFAP-positive cells indicate the robust presence of astrocytic networks, which are essential for neuron-glia interactions and the overall functionality of the neurovascular environment. Additionally, the panel highlights the expression of **TGF-beta** (red), a key regulator in the development of **neuronal-vascular niches**. TGF-beta is known for its role in the differentiation of glial cells and the stabilization of vascular structures, further confirming the development of complex and functional neurovascular networks in the PITTRep model.
- Furthermore, **COL1A1** (collagen type I alpha 1), shown in magenta, provides evidence for the formation of **extracellular matrix (ECM)**, which is crucial for synaptic interaction and the structural support of the neurovascular tissue. The ECM facilitates communication between neurons and glial cells, contributing to synaptic plasticity and the formation of stable synaptic connections. The expression of COL1A1 indicates that the PITTRep system not only promotes the formation of glial and neuronal cells but also supports the **matrix interactions** necessary for long-term functionality of the tissue.

This figure presents an analysis of **microglia development** and **synaptic pruning** within the neurovascular niches developed using the PITTRep model. The importance of gliogenesis, as demonstrated in Figure V, extends to the critical roles of microglia and astrocytes in maintaining healthy brain function by regulating synaptic pruning and supporting the differentiation of new neurons. The processes of synaptic pruning, essential for the maturation and refinement of neural circuits, are actively facilitated by microglia and astrocytes in this system.

**Figure V:**
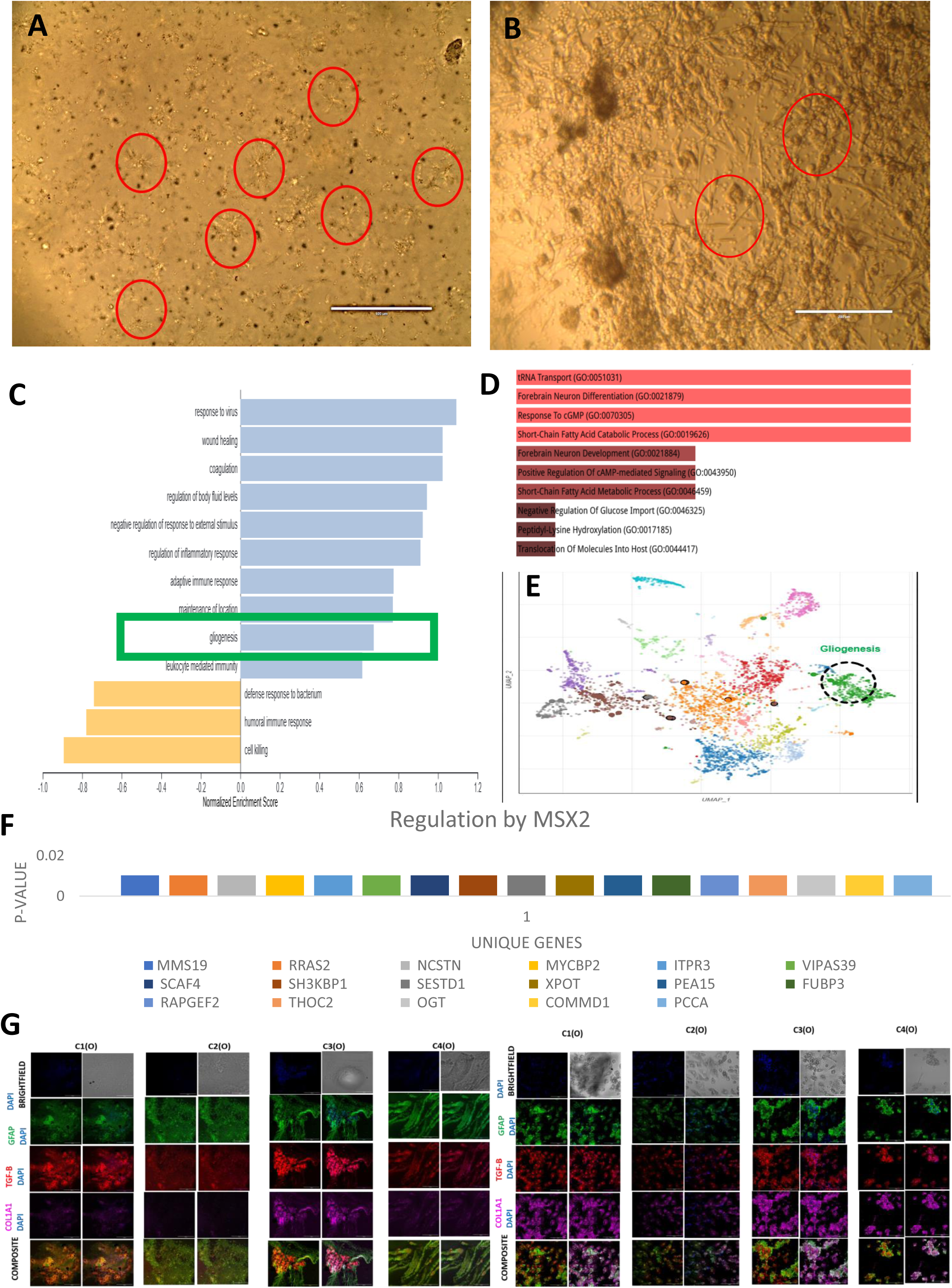
Proteomics Profiling of Neurovascular Niches Indicating Gliogenesis and Forebrain Development.

In **Panel VIA**, confocal microscopy validates the presence of both **microglia** and **astrocytes** in the developing neuronal-vascular tissue across experimental conditions C1 to C4. The specific markers used for this validation include **IBA1** (red), a well-established microglial marker, and **GFAP** (green), an astrocytic marker. The strong expression of these markers across all conditions indicates the robust presence of microglia and astrocytes, essential for synaptic regulation and neurovascular homeostasis. This finding is significant as it highlights the PITTRep system’s ability to mimic critical aspects of in vivo brain development, including the immune and supportive roles of microglia and astrocytes in maintaining neural circuit integrity.

**Panel VIB** provides a proteomic analysis through a **volcano plot**, generated using the Cytoscape platform. The plot shows the upregulation of **complement proteins**, which are key players in synaptic pruning. In the central nervous system, microglia and astrocytes interact with complement proteins by binding to complement receptors, thereby enhancing the process of synaptic elimination. Complement proteins tag weaker synapses for removal, allowing microglia and astrocytes to prune these synapses, a critical process during brain development that ensures efficient and organized synaptic connections. The upregulation of complement proteins in the PITTRep system demonstrates its potential to support proper synaptic pruning dynamics.

**Panel VIC** presents confocal microscopy images that further validate the proteomic data by illustrating the formation of **synaptic boutons**. Synaptic boutons are specialized structures at the terminal ends of axons, where neurotransmitters are released. These boutons are identified using specific markers, including **SOX2** (green), **DBH** (magenta), and **ADRA2A** (red) in the upper image. The presence of these markers indicates the development of functional synapses. In the composite confocal image below, the expression of **Stargazin** (green), **CD40** (magenta), and **HBA1** (red) confirms the formation of the **synaptic cleft** in the presence of an **active extracellular matrix (ECM)**. The ECM plays a crucial role in synapse stabilization and maturation, contributing to synaptic plasticity and the overall development of the neuronal network. This evidence supports the notion that the PITTRep model is capable of generating functional synaptic structures, critical for proper neural communication and circuit formation.

**Panel VID** provides an in-depth analysis of synaptic development using bioinformatic tools, specifically **SynGo**, which is tailored for synapse-related processes. The analysis reveals several key aspects of synaptic formation and regulation within the PITTRep model. The **purple graph** shows the active transportation of molecules across synapses, including **dendritic and axonal transport**, which are essential for synaptic function and neurotransmitter delivery. The **red graph** highlights **synaptic signaling**, including both **presynaptic and postsynaptic signaling**, as well as **anterograde** (from the neuron body to the axon terminal) and **retrograde** (from the synapse back to the neuron body) transport. The **blue graph** demonstrates the organization of synapses, particularly the **remodeling of the actin cytoskeleton**, which is a crucial component of synaptic architecture and plasticity. This data provides further evidence of active synapse formation and maintenance in the PITTRep model, emphasizing the system’s capacity to replicate dynamic synaptic processes in vitro.

In **Panel VIE**, the data are validated once again through confocal microscopy, IBA1 demonstrating the development of active microglia and synaptic pruning across conditions C1 to C3, with more advanced stages observed in C3 and C4. This confocal analysis confirms that as the experimental conditions progress from C1 to C4, there is a corresponding increase in **blood vessel organization** alongside the **neuronal tissue**. The expression of **Stargazin** (green) in these images further corroborates the presence of glutamatergic synapses, which are indicative of mature, functional synaptic structures. The combination of microglial activity and synapse formation suggests that the PITTRep model supports both **neuronal maturation** and **vascular development**, simulating the complexity of neurovascular interactions observed in vivo.

This figure highlights the critical role of the **complement-coagulation system** in neurogenesis, specifically within the PITTRep model. The complement system, traditionally known for its immune function, is revealed to play a significant role in brain development by regulating the differentiation of **meso-ectodermal progenitors** during early human brain formation. The figure underscores how this system not only supports immune responses but also influences key neurodevelopmental processes such as neuronal differentiation, migration, and synaptic pruning.

In **Panel VIIA**, a detailed bioinformatic analysis using the **Enrichr platform** illustrates that the proteomics data from the PITTRep model identifies the complement system as a primary player in **neuronal development**. The analysis outlines that the complement system is involved in several essential processes, including the **differentiation and proliferation of embryonic stem cells**, **hippocampal neurogenesis**, and **synaptic pruning**. These findings suggest that the complement system is intricately linked to the regulation of neural progenitors and their development, supporting the establishment of the brain’s architecture. The pathway analysis indicates a strong relationship between complement activity and the maturation of neurons, particularly in regions like the hippocampus, which is vital for learning and memory.

**Panel VIIB** presents another layer of proteomics analysis conducted via **WebGestalt 2024**, which focuses on the role of **complement-coagulation proteins**. The bar graph in **Panel VIID** quantifies the expression levels of these proteins, indicating their regulatory functions in **neuronal development**. This analysis shows that the complement-coagulation system is actively involved in guiding **axon development**, which is crucial for proper neuronal connectivity in the brain. By facilitating **axon guidance**, the complement system contributes to the proper wiring of neurons and helps ensure that neural circuits are correctly established during brain development.

In **Panel VIIE**, the validation of these proteomic findings is shown through confocal microscopy images that reveal the presence of **VGLUT2**, a marker for glutamatergic neurons, expressed in green. The confocal images depict **axonal spiking**, a clear indicator of **electrical conduction** along the axons. This is further supported by neuronal-specific staining using **Luxol Fast Blue** counterstained with **H&E**, which highlights axonal spikes (encircled in white) that are associated with **electric conduction**. In addition, the third image in this panel shows the development of **myelination** along the axons. Myelination is crucial for the rapid transmission of electrical signals across neurons, and its presence in the PITTRep model underscores the functional maturation of these neurons. The Luxol Fast blue counterstained with H and E images also reveal the formation of **inner and outer tongues** in the synapses, further supporting the model’s ability to replicate physiological myelination and efficient signal conduction in developing neurons.

**Panel VIIF** delves deeper into the proteomic analysis, confirming the presence of **long-term potentiation (LTP)**, a process vital for synaptic strengthening and memory formation. The panel also indicates the presence of various types of synapses, including **glutamatergic**, **dopaminergic**, **cholinergic**, and **GABAergic** synapses. This variety of synaptic types suggests that the neurovascular tissue developed in the PITTRep model is functioning in a **homeostatic state**, with a balanced network of excitatory and inhibitory signals required for maintaining healthy brain function. The inclusion of these synapse types is particularly important for neurogenesis and synaptic plasticity, ensuring that the neural circuits in the model are dynamic and capable of adaptation.

**Panel VIIG** provides a **UMAP analysis** of the developmental stages within the PITTRep model, conducted using the **Appyter platform** (a bioinformatic tool based on R). The UMAP visualization highlights various developmental processes occurring in the system, with a particular focus on **actin cytoskeleton remodeling**, **neurogenesis**, **gliogenesis**, and overall **nervous system development**. The actin cytoskeleton is vital for maintaining synaptic structure and function, and its remodeling is essential for synaptic plasticity. This panel emphasizes that these processes are actively taking place within the PITTRep model, confirming that the system is successfully mimicking key developmental mechanisms present during early brain formation.

Finally, **Panel VIIH** presents the quantitative results of statistical assays measuring the levels of **thrombin** and **plasmin**, two important components of the coagulation system. The assays, conducted across conditions C1 to C4, show significant participation of the **coagulation system** in the neurovascular niches. The active involvement of these proteins in **neuronal regeneration** underscores the role of the complement-coagulation system not only in synaptic pruning and neurogenesis but also in the broader context of brain repair and regeneration. The proteomic and bioinformatic analyses collectively demonstrate how the coagulation system contributes to the maintenance and regeneration of neuronal networks, further solidifying the PITTRep model as a robust platform for studying these complex interactions in vitro.

This figure presents the intricate development of neurovascular tissue through the PITTRep (Plasma Induced Transcriptomics and epi-Transcriptomic reprogramming) approach, demonstrating the complex interactions between neuronal progenitors and vasculature within the developing brain. The figure emphasizes how the transition from hypoxic to normoxic conditions is critical for neuronal progenitor differentiation and maturation, facilitated by the functional flow of blood through an actively developing vasculature.

In **Panel VIIIA**, an illustration created using **BioRender** depicts the development of the **neural tube** under hypoxic conditions. Early in development, the neural tube exists in a hypoxic environment, where the lack of oxygen triggers the upregulation of **HIF-alpha**, a transcription factor that mediates responses to low oxygen levels. As the **perineural vascular plexus** begins to form, it normalizes the hypoxic environment by facilitating active blood flow. This in turn downregulates HIF-alpha, allowing for the progression of neurogenesis. The illustration further shows how, with increasing weeks of gestation, various regions of the neural tube contribute to distinct developmental processes. The **marginal zone** (which eventually forms the white matter) initiates **gliogenesis**, the production of glial cells. The **intermediate zone** plays a major role in **neurogenesis**, giving rise to neurons, while the **ventricular zone** establishes the **choroid plexus** and the **blood-brain barrier**. These zones are populated by various progenitor cells, including **radial glia**, **intermediate progenitors**, **apical and basal progenitors**, all of which contribute to the neuronal network, which becomes increasingly enriched with glial cells. The vascular system releases critical factors that promote neurogenesis, supporting the proliferation and differentiation of progenitors within the developing neural tissue.

**Panel VIIIB** provides experimental validation of the processes illustrated in **Panel VIIIA**. **Scanning electron microscopy (SEM)** reveals the formation of a **neural tube-like structure**, which is essential for brain development. The **marginal zone**, which corresponds to the developing white matter, is further validated through **confocal microscopy** using the **GFAP marker** to highlight the presence of astrocytes. These astrocytes play a key role in supporting neuronal function and maintaining the blood-brain barrier. The **intermediate zone** is also validated through SEM and confocal microscopy, where the presence of **neuronal somas** and neurites is confirmed by the positive staining of **beta-3 tubulin**, a marker for neurons. Additionally, the **ventricular zone**, responsible for generating the ependymal cells that line the brain’s ventricles, is validated by neuronal-specific staining. The confocal images show the expression of **PAX6** (a transcription factor involved in neurogenesis) in green, and **actin cytoskeleton** staining in magenta, indicating the presence of **primitive neuronal progenitors**. The validation is further extended to the developing cortex, where neuronal-specific staining highlights the emerging layers of cortical neurons. The confocal microscopy images on the right side of **Panel VIIIB** show the developing cortex, demonstrating the robust formation of neuronal-vascular tissue within the PITTRep system.

**Panel VIIIC** further validates the development of neurovascular tissue by examining the expression of specific neuronal and vascular markers. **VGLUT2**, a marker for glutamatergic neurons, and **c-Kit**, a marker for hematopoietic stem cells and vascular progenitors, are detected in the in-vitro tissue developed using the PITTRep approach. The lower portion of the panel highlights the presence of the **blood-brain barrier (BBB)**. Confocal images show **GFAP** staining (in green) for astrocytes, which form the **astrocytic end-feet** that surround blood vessels, and **c-Kit** (in red), which marks the **lumen of blood vessels**. These findings, indicated by a red arrow, demonstrate the presence of organized vascular structures within the developing brain tissue, while the white arrow points to the astrocytic feet, providing further evidence of an actively forming BBB. This complex interaction between astrocytes and blood vessels is essential for maintaining the integrity of the neurovascular system and supporting the homeostasis of the brain environment.

The presence of key transcription factors critical for neurogenesis is also confirmed. **5-hmC** (5-hydroxymethylcytosine), shown in green, and **Foxa2**, shown in red, are strongly expressed within the neurovascular tissue, further validating the active **neurogenesis** taking place. These transcription factors are crucial regulators of cortical development, and their expression suggests that the **developing layers of the cortex** are forming in a manner that mirrors early brain development in vivo.

Figure IX presents the detailed stages of neurovascular tissue development achieved through the PITTRep approach, highlighting the structural and molecular differentiation of the developing brain. This figure emphasizes the spatial and temporal patterns of neuronal and vascular development, providing insights into anterior-posterior axis formation and the maturation of specific brain regions.

**Figure VI:**
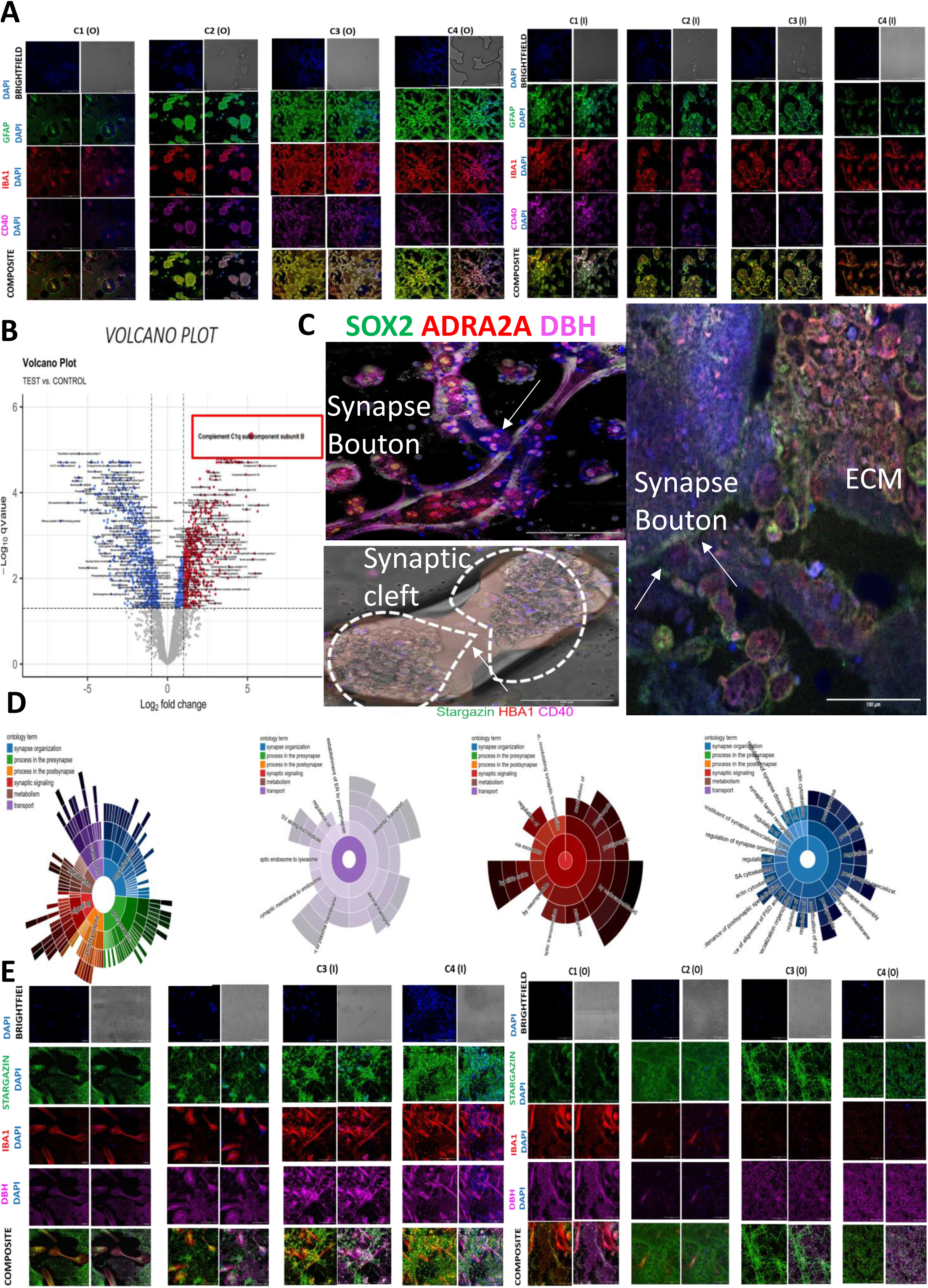
Proteomics Profiling and Confocal Microscopy of Neurovascular Niches Indicating Microglia Development and Synaptic Pruning.

**Figure VII:**
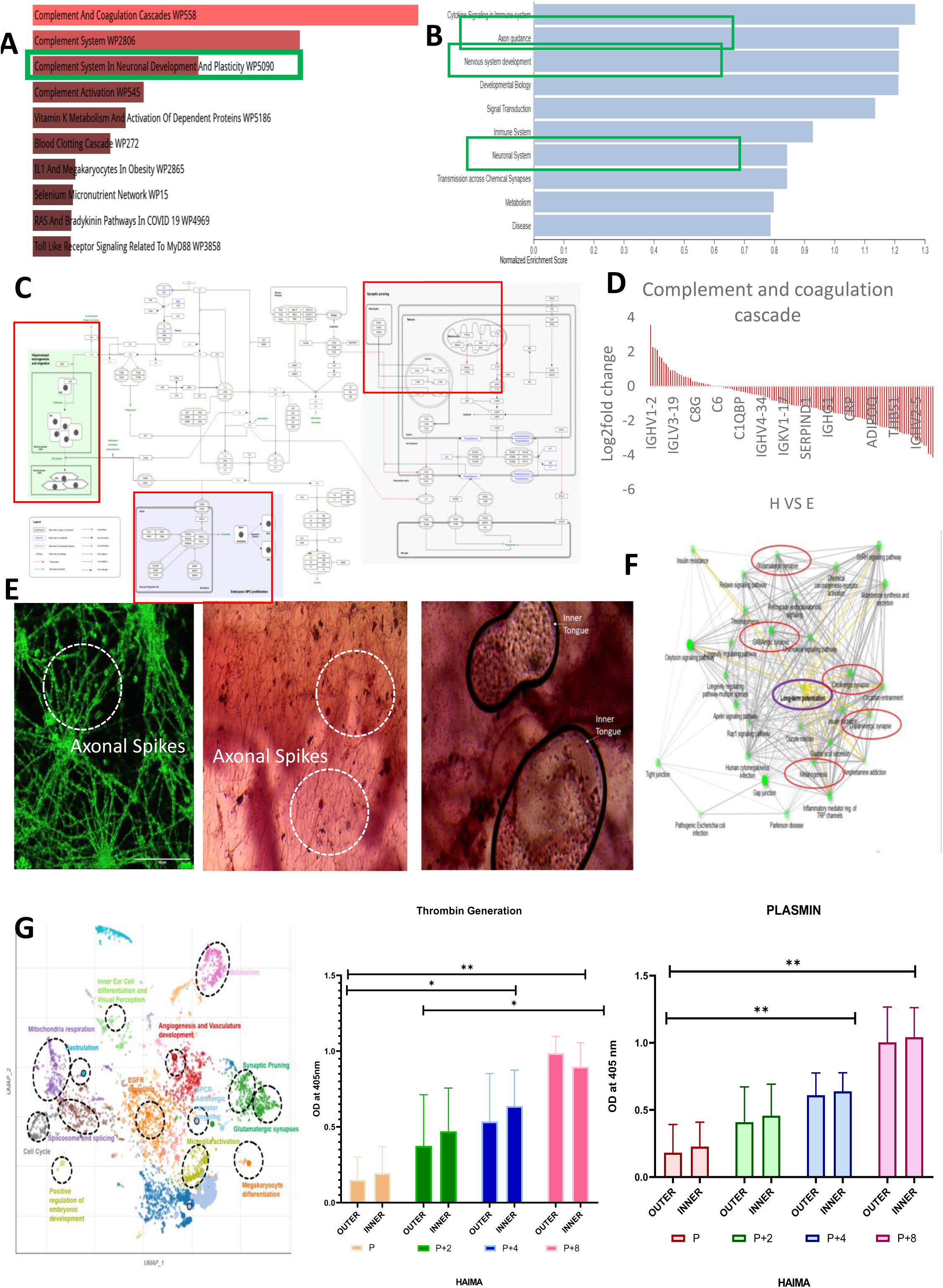
Proteomics Profiling and Confocal Microscopy of Neurovascular Niches Indicating the Role of the Complement System in Neurogenesis.

**Figure VIII:**
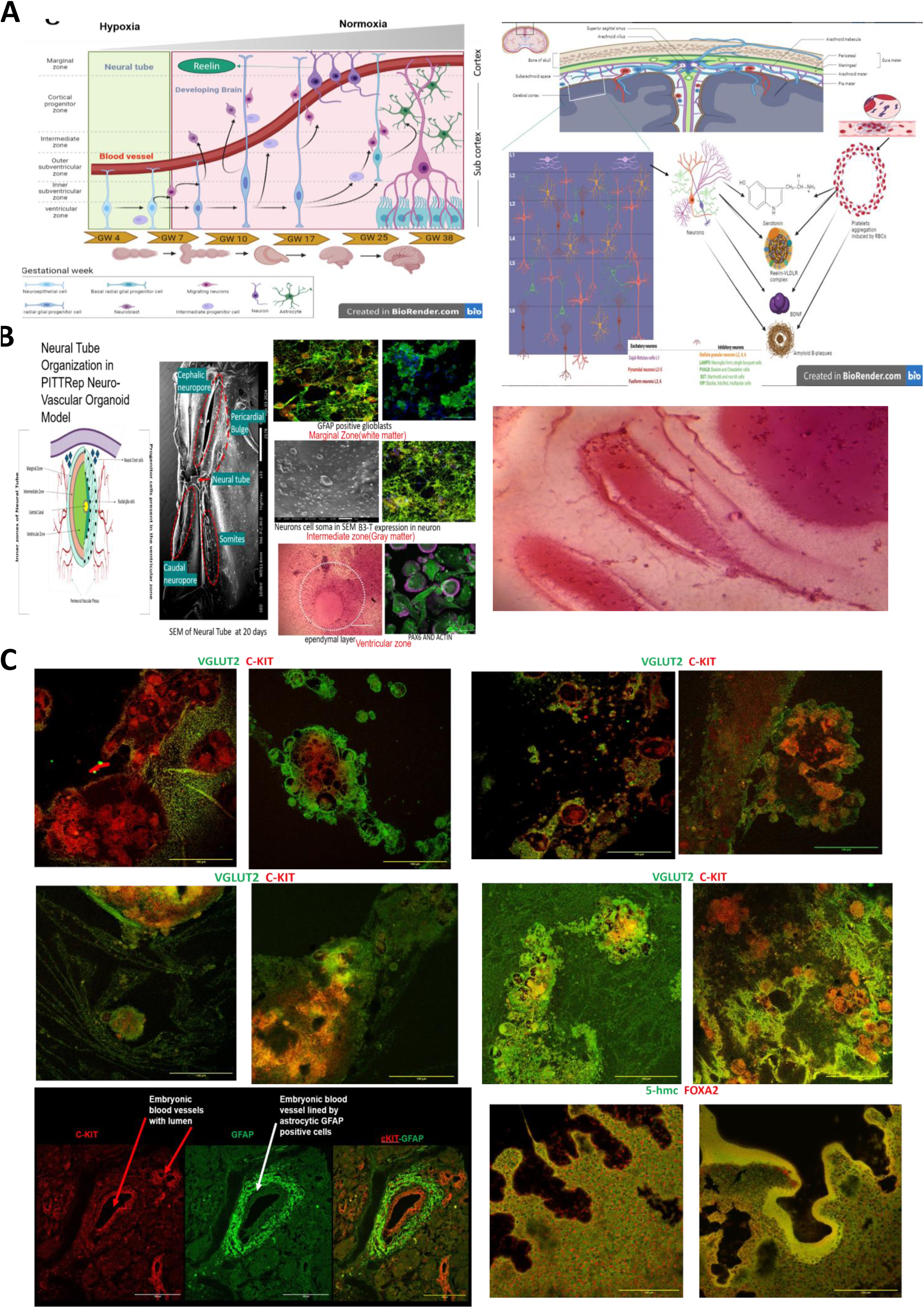
Neurovascular Tissue Development Using the PITTRep Approach.

**Figure IX:**
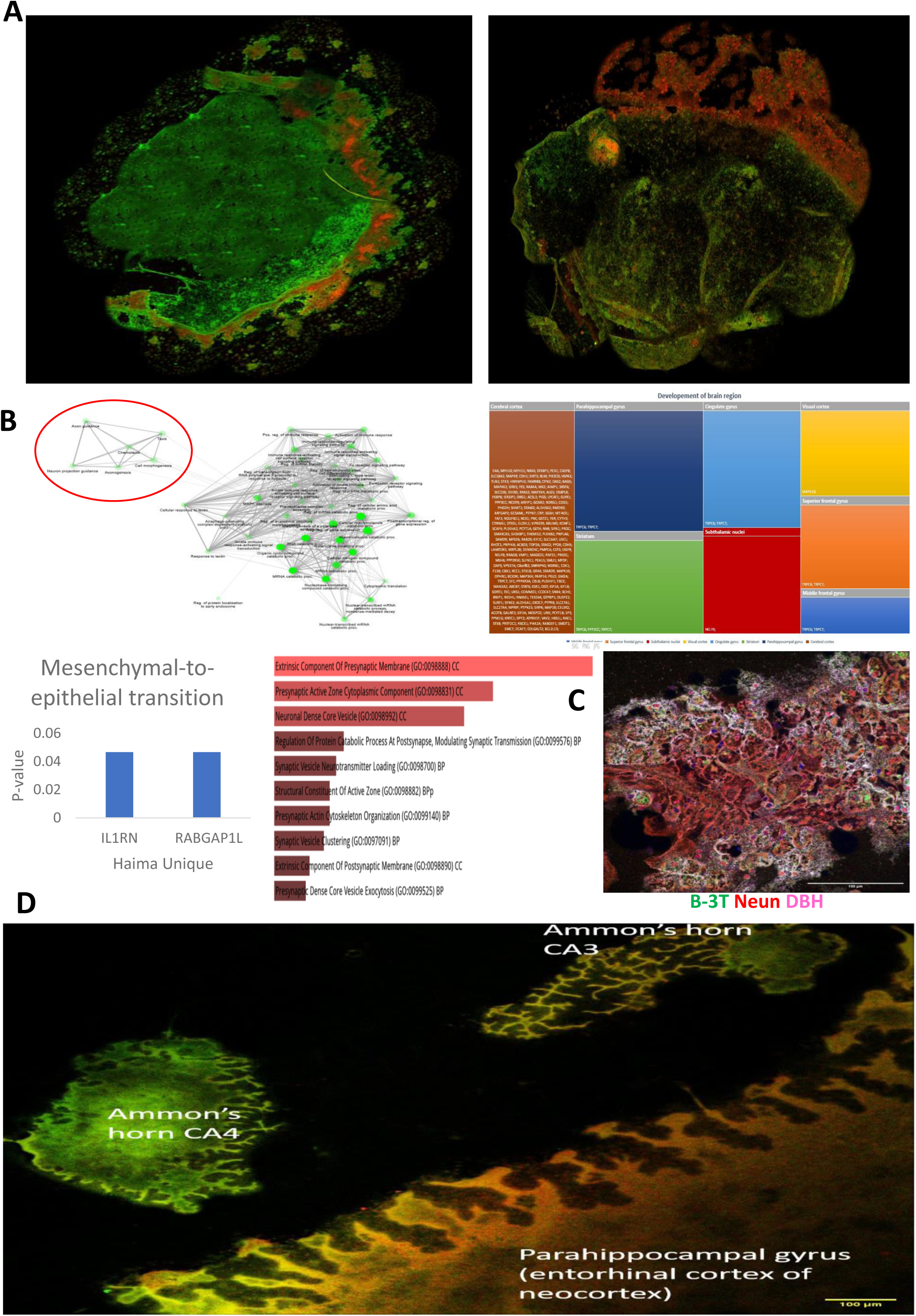
Neurovascular Tissue Development by Employing the PITTRep Approach.

**Figure X:**
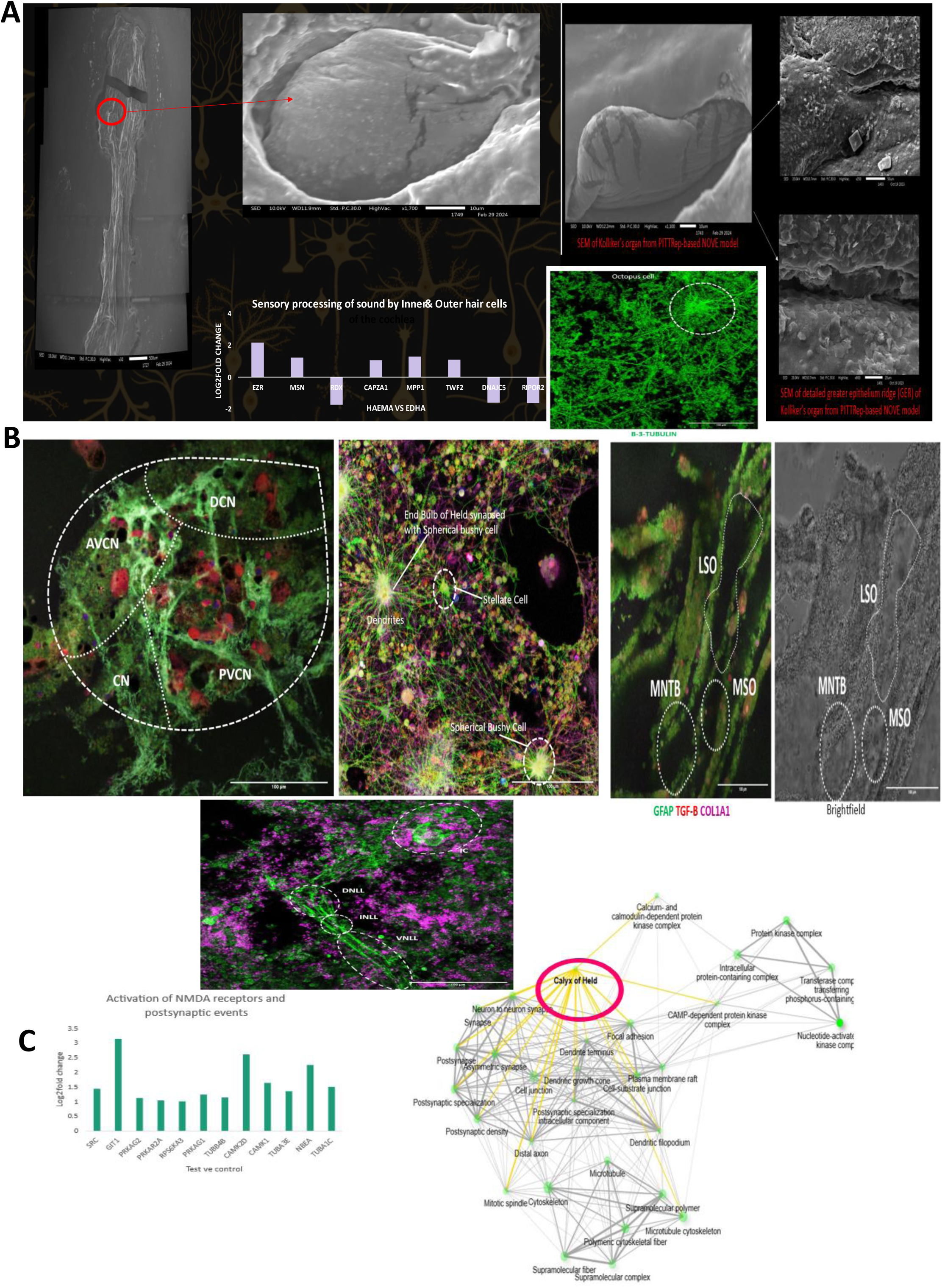
Development and Characterization of Peripheral and Central Auditory Niches by Employing the PITTRep Approach.

**Figure XI:**
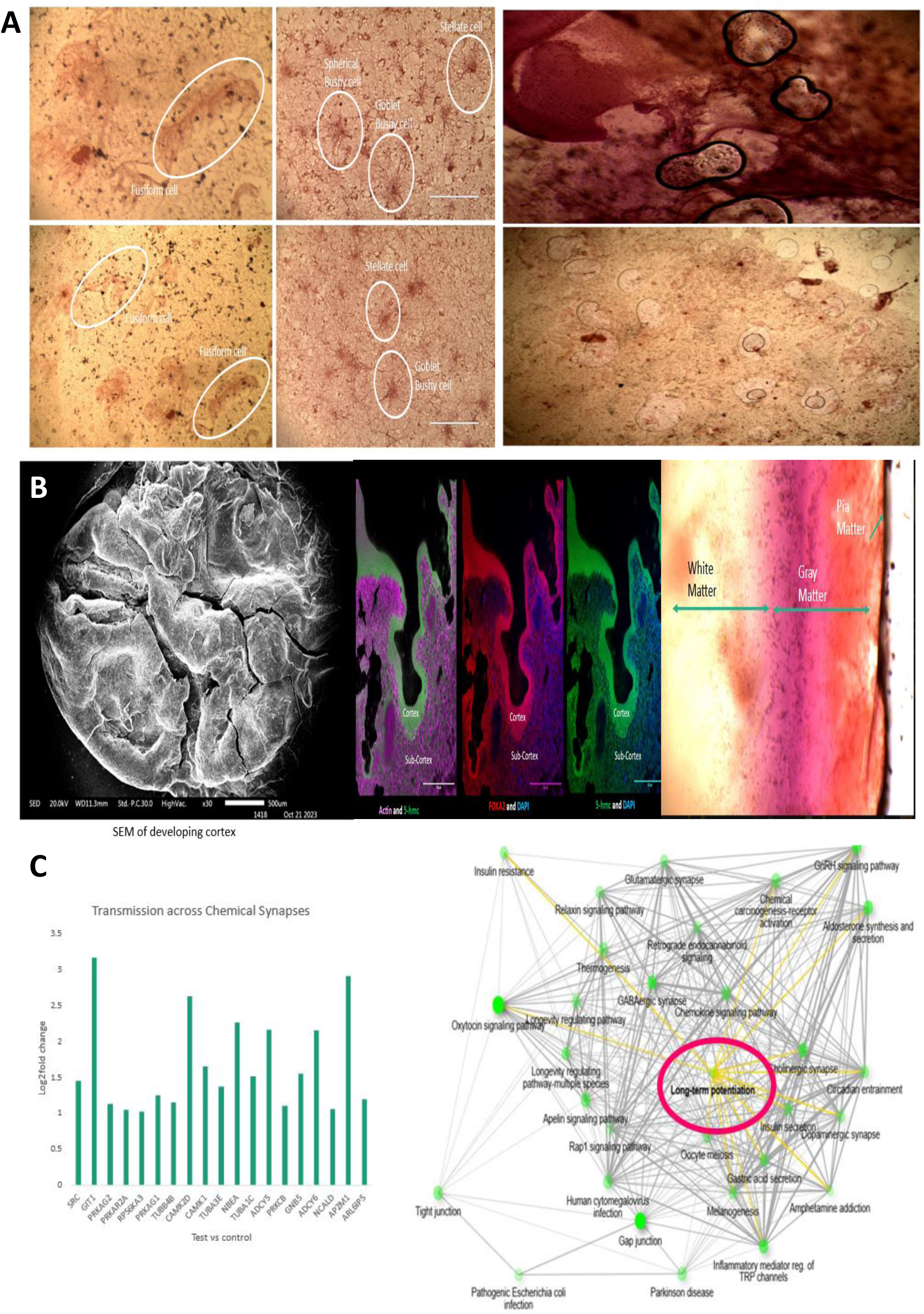
Development and Characterization of Peripheral and Central Auditory Niches by Employing the PITTRep Approach.

**Figure XII:**
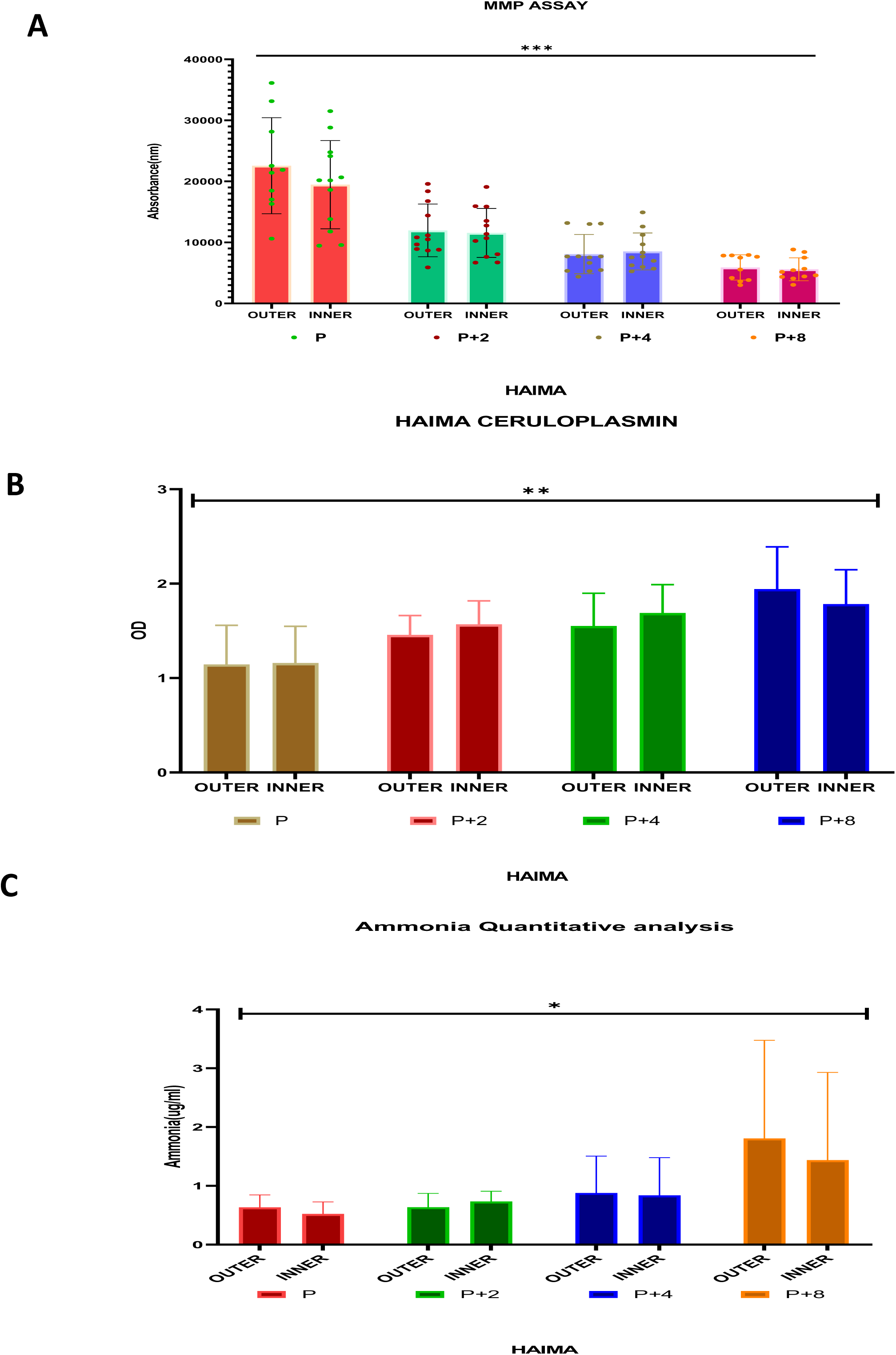
Characterization by functional enzyme assays and ammonia generation assay.

In **Panel IXA**, the confocal microscopy images show the regionalization of the neurovascular tissue. The left side of the panel displays the spatial expression of markers that differentiate between anterior and posterior brain regions. **VGLUT2**, shown in green, marks the **diencephalon**, **midbrain**, and initial parts of the **brainstem**, indicating the anterior regions of the developing brain. The expression of **HOXA4**, shown in red, is localized to the **brainstem**, marking the posteriorization of the tissue. This demonstrates that the PITTRep approach successfully recreates the anterior-posterior axis of brain development in vitro, which is crucial for proper neuronal differentiation and organization.

On the right side of the panel, the development of the **brain tissue**. **MSX2**, displayed in red, is a master transcription factor involved in **osteogenesis** within the brain. It is shown to be strongly expressed in developing regions of the brain crown, where it plays a critical role in promoting **bone formation** and inhibiting the expression of **SOX2**, a marker of pluripotency. The presence of MSX2 suggests the active involvement of **mesodermal differentiation** and osteogenic processes in brain development. The red crown-like induction might indicate skull-like tissue formation. In contrast, **PAX6**, shown in green, is a transcription factor critical for brain development, particularly in regions responsible for **neurogenesis**. PAX6 expression here highlights the emergence of distinct brain regions undergoing neuronal differentiation, complementing the mesodermal-to-neuroepithelial transitions observed during brain tissue maturation.

**Panel IXB** presents the proteomic analysis conducted using various bioinformatics tools. The analysis shows the development of **pre-and postsynaptic structures** as well as **neuronal core vesicles**, which are essential components of synaptic transmission. This proteomic data was analyzed using the **Enrichr** tool, which identified significant pathways involved in **axon guidance** and **neurogenesis** in the developing system. A bar plot highlights the unique proteins implicated in the **mesenchymal-to-neuroepithelial transition**, a critical step in brain development, during which mesodermal progenitors differentiate into neuroepithelial cells. These proteins were further analyzed using the **FunRich database**, which provided insights into the formation of specific brain regions. A **treemap** on the right side of the panel shows the brain regions associated with these proteins, including the **parahippocampal cortex**, **cerebral cortex**, **striatum**, **subthalamic nuclei**, **visual cortex**, **cingulate gyrus**, **superior frontal gyrus**, and **middle frontal gyrus**. This comprehensive analysis underscores the successful regional differentiation of neurovascular tissue, mirroring the complexity of brain organization in vivo.

**Panel IXC** provides validation of the bioinformatic analysis, showing the transition of neurovascular tissue from an immature to a mature stage. The composite confocal microscopy images reveal the expression of critical neuronal markers, including **beta-3 tubulin**, which is essential for cytoskeletal structure in developing neurons; **NeuN**, a marker for mature neurons; and **DBH** (dopamine beta-hydroxylase), which marks **adrenergic and nor adrenergic neurons**. These markers confirm the progression of neurovascular tissue maturation, as immature neurons gradually develop into fully differentiated and functional neuronal populations within the system.

**Panel IXD** further corroborates the bioinformatic analysis through confocal microscopy, 5hmc(green) and FOXA2(red) illustrating the development of the **parahippocampal cortex** or the **entorhinal gyrus** along with the progression of the **Ammon’s horn** (CA3 and CA4 regions) within the hippocampus. These images provide critical insights into the spatial and temporal dynamics of neurovascular tissue development, revealing the intricate interplay between neuronal differentiation, glial cell support, and vascularization that is essential for the establishment of functional brain regions. This particular culture was grown for over 4 weeks.

This figure highlights the development of both peripheral and central auditory niches using the novel PITTRep (Peripheral-central auditory neuronal niches) model.

A major obstacle in auditory research lies in the characterization of otic vesicles in fetal tissues, which presents significant ethical concerns. Hence, developing auditory tissue in vitro becomes essential for studying vestibular and cochlear development. While prior studies have achieved the development of vestibular macula hair cells, no successful model for the in vitro development of cochlear inner and outer hair cells exists. The PITTRep model offers a promising solution, enabling the creation of peripheral and central neurovascular niches for auditory research.

Panel XA: This panel presents the development of the peripheral auditory niche, validated by scanning electron microscopy (SEM). It illustrates the formation of Kolliker’s organ, a structure found in the developing cochlea. Kolliker’s organ consists of the greater and lesser epithelial ridges, with the greater epithelial ridge supporting the differentiation of inner hair cells, eventually leading to the formation of the organ of Corti—critical for auditory function. Additionally, a bar plot shows the proteomic data analysis for sensory processing pathways, focusing on cochlear inner and outer hair cells.

Panel XB: This panel focuses on the development of the central auditory niches. The left side depicts confocal microscopy images of PITTRep-derived neurovascular organoids (NVOEs), showing segments of the cochlear nucleus. The identified regions include the anteroventral cochlear nucleus (AVCN), posteroventral cochlear nucleus (PVCN), and dorsal cochlear nucleus (DCN). The auditory fibers are shown entering the CN (cochlear nucleus), and GFAP staining highlights glial cells along different neuronal sub types, while TGF-Beta staining (red) emphasizes on neurovascular accelerated ascending auditory signaling.

In the right panel, a closer view of the AVCN reveals distinct neuronal types, such as spherical and globular bushy cells. These cells play a pivotal role in auditory signal propagation. Spherical bushy cells receive synaptic inputs from inner hair cells, forming large synapses at the end of Held on the spherical bushy cells. Stellate cells, also present, contribute to sound propagation. Excitatory fibers, marked in green, express VGLUT2, and VMAT2-positive vesicles (red) are shown at active presynaptic terminals, while DBH-positive cells are highlighted in magenta.

The middle section of the panel demonstrates the development of the PVCN, visualized via confocal microscopy. Octopus cells, crucial for processing complex sounds in the dorsal auditory stream of the ascending auditory pathway, are observed.

The lower middle portion presents the lateral lemniscus pathway, including the ventral nucleus of the lateral lemniscus (VNLL), dorsal nucleus of the lateral lemniscus (DNLL), and intermediate nucleus of the lateral lemniscus (INLL). These form a bundle of axons within the ascending auditory central pathway that leads to the inferior colliculus in the midbrain.

Panel XC: The final panel demonstrates the development of the calyx of Held synapse in the PITTRep model, supported by proteomic analysis. The calyx of Held is a large synapse located at the medial nucleus of the trapezoid body (MNTB), which plays a critical role in the inhibition of the contralateral lateral superior olive (LSO) upon receiving excitatory signals from the AVCN. Furthermore, the data indicate long-term potentiation maintenance at the calyx of Held synapse. The accompanying bar plot illustrates the activation of glutamatergic NMDA receptors, vital for regulating auditory pathways.

This figure illustrates the comprehensive development of both peripheral and central auditory niches, following the auditory pathway from the peripheral structures to the midbrain and higher-order cortical regions. It also demonstrates the connection between the descending pathways from the cortex back to the peripheral auditory system. The novel PITTRep (Peripheral-central auditory neuronal niches) model offers a platform for studying these complex auditory circuits.

### Panel XIA

This panel depicts the **Medial Olivocochlear (MOC) system**, a critical component of the descending auditory pathway. The MOC neurons project from the brainstem to the cochlea, with each neuron receiving inputs from 1 to 4 auditory fibers. The upper left section presents a Luxol fast blue counterstained with H&E (Hematoxylin and Eosin), showing the presence of MOC neuron. These neurons are responsible for modulating cochlear function, such as suppressing background noise and protecting the cochlea from acoustic overstimulation, thus enhancing signal detection in complex auditory environments.

### Panel XIB

This panel presents the development of the **cortex** in the PITTRep model as seen by Scanning electron microscopy (SEM). The confocal microscopy reveals the differentiation and maturation of marginal and intermediate zones into the cortex and sub-cortical structures. This differentiation process is closely regulated by active vascular flow, which is essential for proper neuronal development and function. The cortical region is also seen by luxol fast blue staining in the extreme right panel.

Confocal microscopy images show actin cytoskeleton remodeling in these developing cortical regions using the **Phalloidin marker** (magenta). The remodeling of actin filaments is critical for neuronal migration, neurite outgrowth, and synapse formation. Additionally, high expression levels of transcription factors, including **FOXA2** (red) and **5-HMC** (5-hydroxymethylcytosine, green), highlight the upregulation of neurogenesis and cortical development. These factors are crucial for defining cell fate, regulating neuronal differentiation, and maintaining neuroplasticity.

The extreme right panel demonstrates the formation of key cortical structures, including the **meninges**, **gray matter**, and **white matter**, as visualized by neuron-specific staining. This structural development is vital for higher-order brain functions, including auditory processing, memory, and cognition.

### Panel XIC

This panel focuses on the **proteomic profiling** of auditory niches, revealing the molecular mechanisms underpinning the development of long-term potentiation (LTP), a process essential for synaptic plasticity and memory formation. Using bioinformatics analysis with the **ShinyGO v0.80** database, the figure provides insights into the auditory pathways’ functional activity, particularly in glutamatergic neurons, which are crucial for excitatory neurotransmission in the auditory system.

The accompanying bar plot illustrates the presence of **active chemical synapses** in the auditory pathways, specifically highlighting the glutamatergic neurons. These synapses are vital for transmitting auditory signals from the peripheral auditory structures to the cortex and for processing complex auditory information, such as sound localization and speech recognition. The involvement of LTP in these pathways underscores the importance of synaptic plasticity in maintaining auditory function and adapting to auditory stimuli.

This figure presents the metabolic and proteolytic profiles of PITTRep model across different conditions (C1 to C4) associated with brain development, as quantified by various biochemical assays. Each panel provides quantitative insights into the dynamic changes in specific biomolecules, highlighting their roles in neurodevelopmental processes.

**Panel XII A**: This panel demonstrates a **quantitative assay of matrix metalloproteinases (MMPs)**, which are critical enzymes involved in extracellular matrix remodeling during tissue development, including the brain. The assay results show a **significant decrease in MMP levels** as the conditions progress from C1 to C4. This trend indicates a correlation between **brain maturation and reduced proteolytic activity**, as lower MMP levels are associated with stabilized extracellular matrices, reduced tissue remodelling, and increased neuronal integrity. The results suggest that as brain development advances, the demand for MMP-mediated matrix turnover diminishes, favoring the structural consolidation of neuronal circuits.

### Panel XIIB

This panel presents a **quantitative assay of ceruloplasmin levels**, a copper-binding glycoprotein involved in iron metabolism, with a particular focus on its role in the release of ferritin, the iron storage protein. The data show a **significant increase in ceruloplasmin levels** across the conditions from C1 to C4, correlating with advancing brain development. This rise in ceruloplasmin levels is associated with **enhanced ferritin release**, which is crucial for maintaining iron homeostasis during neurodevelopment. Proper iron regulation is essential for processes such as myelination, neurotransmitter synthesis, and mitochondrial function. The increase in ceruloplasmin suggests a growing demand for iron management in the developing brain to support these critical functions

### Panel XIIC

This panel illustrates a **quantitative assay of ammonia levels**, which are known to reflect metabolic processes, including amino acid catabolism and neurotransmitter synthesis. The assay reveals a **significant increase in ammonia levels** from condition C1 to C4, indicating an association between **brain development and rising metabolic activity**. Ammonia is a byproduct of several metabolic pathways, and its accumulation can be linked to heightened energy demands and the biosynthesis of neurotransmitters such as glutamate and GABA during neurodevelopment. While increased ammonia levels are necessary for brain maturation, they also underscore the importance of efficient ammonia clearance mechanisms to prevent neurotoxicity.

The study was approved by the Institute Ethics Committee, PGIMER, Chandigarh 160012, India.

### Neuro-vascularised tissue from autologous blood-an Optoacoustic imaging proof

Optoacoustic imaging was performed on 24 day tissue cultured in different conditions C1-C4. The live tissue was embedded in low melting agarose to prepare phantoms (Figure-1A) and then placed in the iTHERA MSOT inVision optoacoustic imaging device. The results indicate the presence of a defined Hemoglobin (HbO2) and Deoxy-hemoglobin (Hb) signal in the neuro-vascularised tissue (Figure-1 C and D). Some parts of this tissue present more intensified signal than the other, indicating the presence of more abundant vasculature developed in that area (highlighted area in Figure-1 C and D). Although no micro vessels got defined from the Hb and HbO2 spectra scans. The signal strength is more in C4 condition tissue indicating towards more robust development of neuro-vascular tissue with respect to other conditions.

Link to high resolution figures: https://ln5.sync.com/dl/63b1e0c30/26ds74s7-xv5aj2di-9pzwr9uc-33mjeies https://ln5.sync.com/dl/4c5b50af0/ihvz5cwd-yf92ckb4-8mbskgxt-635hfa7e

Note: The link for raw data will be uploaded soon.

## Discussion

We have developed a human in vitro model of autologous blood-derived neurovascular tissue that is free from exogenous genetic modification, external growth factors, and induced pluripotent stem cell (iPSC) derivation. This model uniquely integrates functional vasculature and neurogenesis. To our knowledge, it represents the first adult blood-derived neuro-vascular tissue model capable of generating functional vasculature, as evidenced by haemoglobin signal detection through optoacoustic imaging, while mimicking a blood-brain barrier (BBB)-like environment with diverse cell populations, including microglia and astrocytes. All this has been achieved in a self-patterning system which is devoid of any morphogen guided patterning.

Our findings demonstrate the formation of vasculature independent of any established methods, such as transplantation into host animals, endothelial cell co-culture, mesodermal progenitor co-differentiation, mechanical stimulation, or the use of engineering techniques like microfluidic chambers or scaffold materials (e.g., Matrigel). The vasculature that forms in our system is functional, as indicated by the results of optoacoustic imaging, and develops in tandem with neuronal progenitors, supporting the differentiation and survival of neurons within a complex neurovascular niche. The autologous nature of this model system increases its relevance for developing patient specific neuro-vascular tissue that can be used for investigating the molecular basis of neurosensory, neuro-vascular and neurodegenerative disorders. One of the most persuasive implications of this system lies in its potential for developing autologous neuro-regeneration therapies, which offers a patient-tailored approach for treating neurological diseases through the transplantation of patient-specific neurons derived from their own blood. Furthermore, the formation of BBB like structure suggests it use for studying anatomy and physiology of a healthy and diseased BBB essential for the development of neurotherapeutic drugs and subsequent drug delivery systems. PITTRep methodology derived neuro-vascular tissue is a fully physiologic, highly reproducible, and cost-efficient model which enhances its appeal for conducting comprehensive brain investigations and formulating therapeutic interventions for neurological disorders.

Brain activity-dependent synaptic potentiation (SP) has long been proposed to represent the subcellular substrate of learning, memory, and for integrating complex sensory inputs. Long-term potentiation (LTP) is a key mechanism underlying synaptic plasticity and is closely linked to learning and memory processes. It refers to the persistent strengthening of synaptic connections based on recent patterns of activity (Abraham et al., 2019). The study presented by Kotak and colleagues in 2007, has demonstrated that sensorineural hearing loss (SNHL) induces alterations in synaptic strength within the auditory cortex using gerbils. These changes include a shift towards long-term depression (LTD) rather than LTP, suggesting that developmental hearing loss eliminates long-term potentiation in the auditory cortex and normal auditory experience is essential for the maturation of synaptic plasticity mechanisms (Kotak et al., 2007). Most adults with prior hearing experience can comprehend speech via a cochlear implant well enough for phone conversations without lip-reading because of the auditory LTP developed earlier during normal language acquisition. However, in cases of congenital hearing loss where the brain’s pattern-recognition system has not been developed, the central auditory system must adapt to a new spectrum of peripheral inputs. Hence poor speech perception is observed in pre-lingually deafened but later implanted children (Moore et al., 2009). The mechanism of developing auditory long term potentiation using cochlear implants is not yet deciphered which once understood can help in understanding the vast outcomes in the patients who have received this implant.

The challenge is understanding how cochlear implants (CIs) can induce auditory Long Term Potentiation (LTP), which is vital for effective auditory processing and speech perception. Neuroplasticity, the brain’s ability to change and adapt, is most active during critical and sensitive periods early in life (Citri & Malenka, 2007; Kral & Sharma, 2012). These periods are essential for sensory and cognitive development, with early auditory experiences being crucial for proper hearing and language skills (Pedrosa et al., 2022; Polley et al., 2013). While early CI implantation greatly benefits language and auditory development, the exact mechanisms through which CIs enhance neuroplasticity are not well understood (Kral & Sharma, 2012; Kotak et al., 2007). Mixed results in studies about LTP induction in CI users point to the need for further research (Barlow, 2014; Lei et al., 2017). More comprehensive studies are needed, combining EEG, fMRI/fNIRS, and other methods with molecular research, to better understand and improve CI outcomes (Zaehle et al., 2007; Jia et al., 2024).

Considering the complexity of auditory processing and speech perception in human brain and the fact that CI performance is thought to be influenced by genetic variables and brain plasticity it is essential to perform in vitro studies for determining critical molecular and genetic cross talk which regulates these patient specific outcomes in CI. Stem cell derived brain organoids have emerged as powerful tool for understanding the brain development and modelling neurological disorders. Since brain is a highly functional tissue, neuronal activity is the key feature to determine the functional aspect of the developed brain organoids (Mulder et al., 2023). Neural circuit formation and integration reflects about the complex activity developing in the brain organoids that is very essential for recapitulating any disease physiology. Emerging studies have been successful in recording the complex activity and short term plasticity/potentiation in cerebral organoids (Osaki et al., 2024). But due to insufficient maturation and lack of long term potentiation/synaptic plasticity, the current brain/cortical organoids fail to completely replicate the functionality of brain tissue, and dissuade investigative brain-activity (sensory-evoked neural activity) based studies. The major reason for lack of mature neurons and obtaining LTP synapses in the cerebral organoids is the lack of active vasculature along with absence of supporting cells such as microglia and astrocytes which promote NVC. NVC is vital for synaptic strengthening and brain activity as it ensures sufficient oxygen and nutrients are delivered to support the increased metabolic demands associated with synaptic strengthening and plasticity in the brain.

Given the limitations of current brain organoid protocols, which include the absence of a functional neuro-vascular unit, incomplete mimicking of the brain’s cytoarchitecture, complexity and cellular diversity, physiological functioning, and the lack of holistic recapitulation of complex neural circuitry and the microenvironment, they are unsuitable for studying neurosensory disorders such as SNHL. Although bioengineered models of the neurovascular unit, blood brain barrier, and brain on chip device address some of these limitations, they also have their own set of challenges, such as delicate protocols, the use of artificial fabrication materials, the presence of incomplete cell types, scalability, and cost effectiveness, all of which impede the use of such brain investigative studies. Therefore it is imperative to develop a model system which is more physiologic and incorporates a better functional integrity of the brain tissue and brings us a step closer to understand the complexity of neural circuitry and mechanism of neural plasticity.

For addressing this problem we incorporate a novel protocol of haemodynamic reprograming, by pivoting around our previous reprograming protocols (**Sharma and Panda, 2020; Sharma et al., 2022, Sharma et al., 2023; Arora et al., 2024;** https://dst.gov.in/new-prototype-developed-generate-neurovascular-tissuesorganoids-autologous-blood-can-help-precision), which takes into account the fluid mechanic feature of blood erythrocytes as it flows not only via blood vessels of smaller/larger diameter, but also under abnormal flow states, such as in the presence of stenosis, aneurysm, and thrombosis. Pathophysiology of ischemic stroke (also experimentally induced in animals by photo-thrombosis in cerebral areas) lies at the interface of cerebrovascular deregulation that is characterized by (physiologically intriguing yet clinically meaningful) neurological recovery with a non-linear and dynamic logrithmatics. Remarkably enough, post-stroke heightened neuroplasticity marked by dendrites, synapses, astrogliosis, microglia activation, and growth factor upregulation (neuroregeneration and compensation) is accompanied by adult neurogenesis (otherwise non-resilient phenomenon). The current study attempted to develop a functional neurovascular tissue from PITTRep methodology based on the principle of hemodynamic reprograming from autologous blood.

Overall our findings suggest that the current approach to generating neurovascular tissue from autologous blood provides a highly efficient model for studying the pathophysiology of a wide range of neurological, neurosensory, and neurodevelopmental conditions. It enables the path for developing novel therapeutic approaches, particularly in neuro-regeneration, by facilitating autologous transplantation of functional neurons tailored to the individual patient.

C1 to C4 present increasing concentration of RBC in the culture. Where C2 serves as baseline added concentration of RBC, C3 has twice the concentration and C4 has three times the concentration than C2.

### Methodology

#### Neuro-vascular tissue culture condition

Blood sample was obtained from healthy subject (devoid of any underlining health condition at the time of withdrawing blood) and processed further using PITTRep methodology. Briefly, autologous blood components were first separated using centrifugation and then were mixed together with increasing RBC concentration for obtaining the conditions C1, C2, C3 and C4. Where C2 serves as baseline added concentration of RBC, C3 has twice the concentration and C4 has three times the concentration than C2. The autologous blood components were cultured at optimal culture conditions in a CO_2_ incubator for obtaining the neuro-vascular tissue for different timepoints (maximum being 30 days).

#### Routine staining

Neuron-specific Luxol fast blue counterstaining with Hematoxylin and Eosin was performed on the cells. The cells were fixed with 4% paraformaldehyde (PFA) followed by incubation with Luxol fast blue. For differential staining lithium carbonate was used, subsequently it was followed by counterstain with Mayer’s hematoxylin and Eosin on the cells.

#### Immunofluorescence

Cells and tissue were fixed with 4% paraformaldehyde (PFA) followed by treatment by 0.3% Triton X-100 and blocked using a freshly prepared blocking solution (2.5% BSA). Cells were then incubated with primary antibodies at the following dilutions: Kit (rabbit, Cell Signaling Technology, 3074s, 1:300), Stargazing (mouse, Invitrogen, MA5-27654, 1:300), Iba-1 (rabbit, Invitrogen, MA5-29012, 1:300), Dopamine Beta Hydroxylase (DBH) (donkey, Invitrogen, PA3925, 1:300), Beta −3 Tubulin (mouse, Invitrogen, MA119187, 1:300), VEGFR2 (rabbit, Invitrogen, PA1-16613, 1:300), CD40 (donkey, Invitrogen, PA5-142944, 1:300), GFAP (mouse, Invitrogen, MA515086, 1:300), Neun (rabbit, Invitrogen, PA5-78639, 1:300), VGLUT2 (mouse, Invitrogen, MA527613, 1:300), Tissue Growth Factor-beta1 (TGF-β1) (rabbit, Bioss, Bs-0086R,1:300), COL1A1 (donkey, Santa Cruz Biotechnology, sc-25974, 1:300), Myh-7A (donkey,Santa Cruz Biotechnology, sc-168678, 1:300), VEGFR3 (mouse, Invitrogen, 14-5988-82, 1:300), HLA-DR (mouse, Invitrogen, 14-9956-82, 1:300), FoxA2 (rabbit, Invitrogen, 1:300,701698), HB-Z (mouse, Invitrogen, MA5-27754, 1:300), GATA4 (rabbit, Cell Signaling Technology, 36966s, 1:300), LYVE-1(mouse, Invitrogen, 14-0443-82, 1:300), GOLPH4 (rabbit, Bioss, bs-13490R, 1:300), PDGFR-Alpha (mouse, Santa Cruz Biotechnology, sc-398206, 1:300), CD34 (mouse, Invitrogen, MA1-10202, 1:300), TCF 21 (LEF) (rabbit, NeoBiotechnologies, 51176-RBM1-P1, 1:300), CD 31(mouse, Invitrogen, MA5-13188, 1:300), VE-Cadherin (rabbit, Invitrogen, PA5-17401, 1:300), FGF-2 (mouse, Santa Cruz Biotechnology, sc-271847, 1:300), Phospho-FGFR2 (rabbit, Invitrogen, PA5-105880), MAOB (rabbit, Invitrogen, PA528338, 1:300), VMAT2 (rabbit, Invitrogen, PA5-22864, 1:300), ACE-2 (mouse, Invitrogen, MA531395,1:300), ADRA B1 (rabbit, Invitrogen, PA5-28808,1:300), Ki67 (rabbit, Invitrogen, PA5-19462, 1:200), Pax6 (mouse, Santa Cruz Biotechnology, sc-32766, 1:200), MSX2 (rabbit, Invitrogen, PA5-106670, 1:300), HoxA4 (rabbit, Invitrogen, PA5-112574 1:300), PPAR-Alpha (rabbit, Santa Cruz Biotechnology, sc-9000, 1:300), Sox-2 (mouse, Santa Cruz Biotechnology,sc-365823, 1:200), mTOR (mouse, Invitrogen, AHO1232, 1:300), AMPKp (rabbit, Invitrogen, MA5-35486, 1:300), HBA-1 (rabbit, Invitrogen, PA5-102943, 1:300). Secondary antibodies used were goat Alexa Fluor 488 and 568 and donkey Alexa Fluor 647 conjugates (Invitrogen, 1:200) and Alexa Fluor 647 Phalloidin (Invitrogen, 1:500).

#### Scanning Electron Microscopy

For understanding the morphology and structural of the neuro-vascular tissue, the tissue was fixed with 4% paraformaldehyde (PFA) followed by dehydration in different grades of ethyl alcohol. For sticking the sample on the aluminum stub, critical point drying was done after which the sample was coated with gold particle of the size 10^-6^ mm for viewing in the electron microscope.

#### LC-MS/MS and data processing

To prepare the sample, the cell pellet was centrifuged at 3000g for 15 minutes at 4°C and stored at −80°C. For in-solution digestion, 100µL of the cell pellet was lysed with DTT, treated with 6M guanidinium hydrochloride, and heated at 90°C for 10 minutes. The protein concentration was determined using the BCA/Bradford test. Following reduction and alkylation with 10mM DTT and 50mM iodoacetamide, the sample was digested with LC-MS grade trypsin at 37°C for 12-16 hours. Digested samples were kept at −80°C for LC-MS/MS analysis. An LTQ-Orbitrap Velos mass spectrometer was connected to an Agilent 1200 nano liquid chromatography system to undertake proteomics analysis. Peptides were separated on a C18 analytical column with a linear gradient of 7-35% solvent B over 60 minutes. MS/MS data were acquired at a resolution of 15000 in the orbitrap. Data analysis was performed using multiple search engines, including Proteome Discoverer 1.3, Sequest, and Mascot, with searches against the NCBI RefSeq human protein database. A 1% false discovery rate was applied. In-silico analysis of pathways and networks was conducted using tools such as STRING, DAVID, KEGG, and Perseus.

#### Hemoglobin typing

Analytical HPLC quantification of haemoglobin tetramers and individual globin chains was performed using IE and RP columns on an HPLC system, as recently described by Mayuranathan et al. (2023). The relative amounts of haemoglobin or individual globin chains were calculated from the area under the 418-nm peak and normalized based on the dimethyl sulfoxide control. The percentage of respective developmental haemoglobin was calculated, e.g., HbF was calculated from IE-HPLC as follows: percentage HbF = (HbF/(HbA + HbF)) × 100. The percentage of γ-globin haemoglobin subunits was calculated from reverse-phase HPLC as follows: percentage γ-globin = ((Gγ-chain + Aγ-chain)/β-like chains (β+ Gγ+ Aγ)) × 100. 4o

#### Optoacoustic imaging

For preforming Optoacoustic Imaging on the neuro-vascular tissue, the cultured tissue was embedded in phantoms prepared from low melting agarose (SigmaAldrich A9414). The phantoms were scanned using a preclinical iTHERA optoacoustic device at wavelengths from 660 to 980nm, with 5nm steps. Spectral un-mixing was done using all wavelengths comprised between 660 and 870nm with (43 wavelengths).

## Acknowledgement

We acknowledge SERB media cell for featuring our work on its official portal https://dst.gov.in/new-prototype-developed-generate-neurovascular-tissuesorganoids-autologous-blood-can-help-precision; and constant funding support-SERB-SPR/2019/1447; SERB-CRG/2019/6755; SERB-CRG/2022/4946).

---

**Figure 1.**
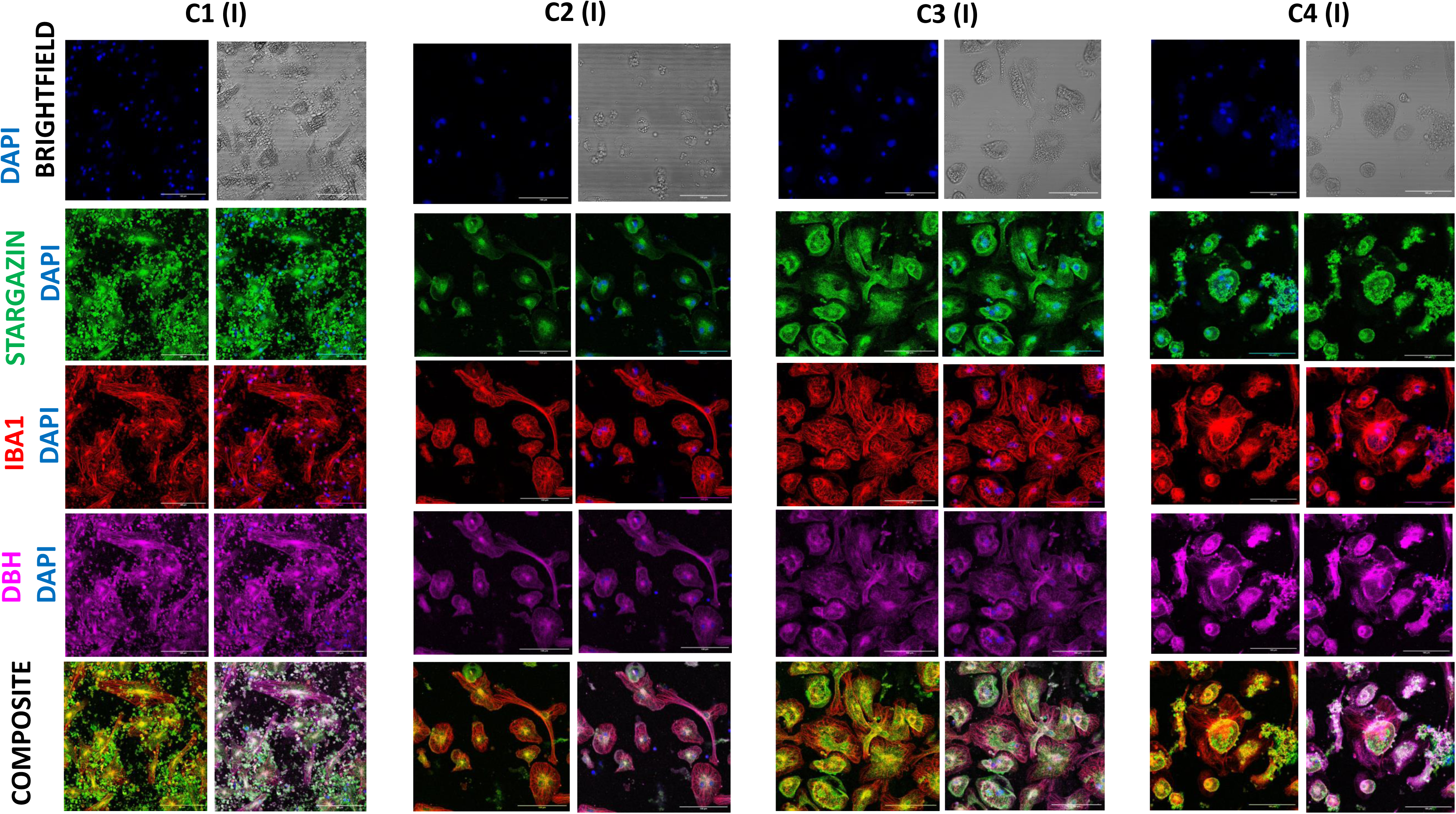
Pruning of synapses by Microglia: The figure depicts the expression Stargazin (green), a marker of synaptic functionality, Iba-1 microglia marker and DBH (pink) a marker of sympathetic (norepinephrine) neurons across different conditions at day 15. The panel shows the data of the cellular field for the culture done in the wells identified as inwardly positioned (I). From C1-C4, the expression of Stargazin, Iba-1 and DBH is increasing. Notice the presence of filamentous actin in the microglia (red) indicating the activated cells involved in pruning of the synapses, there appearance reduces in C4 condition. Scale bars-100 µm.

**Figure 2.**
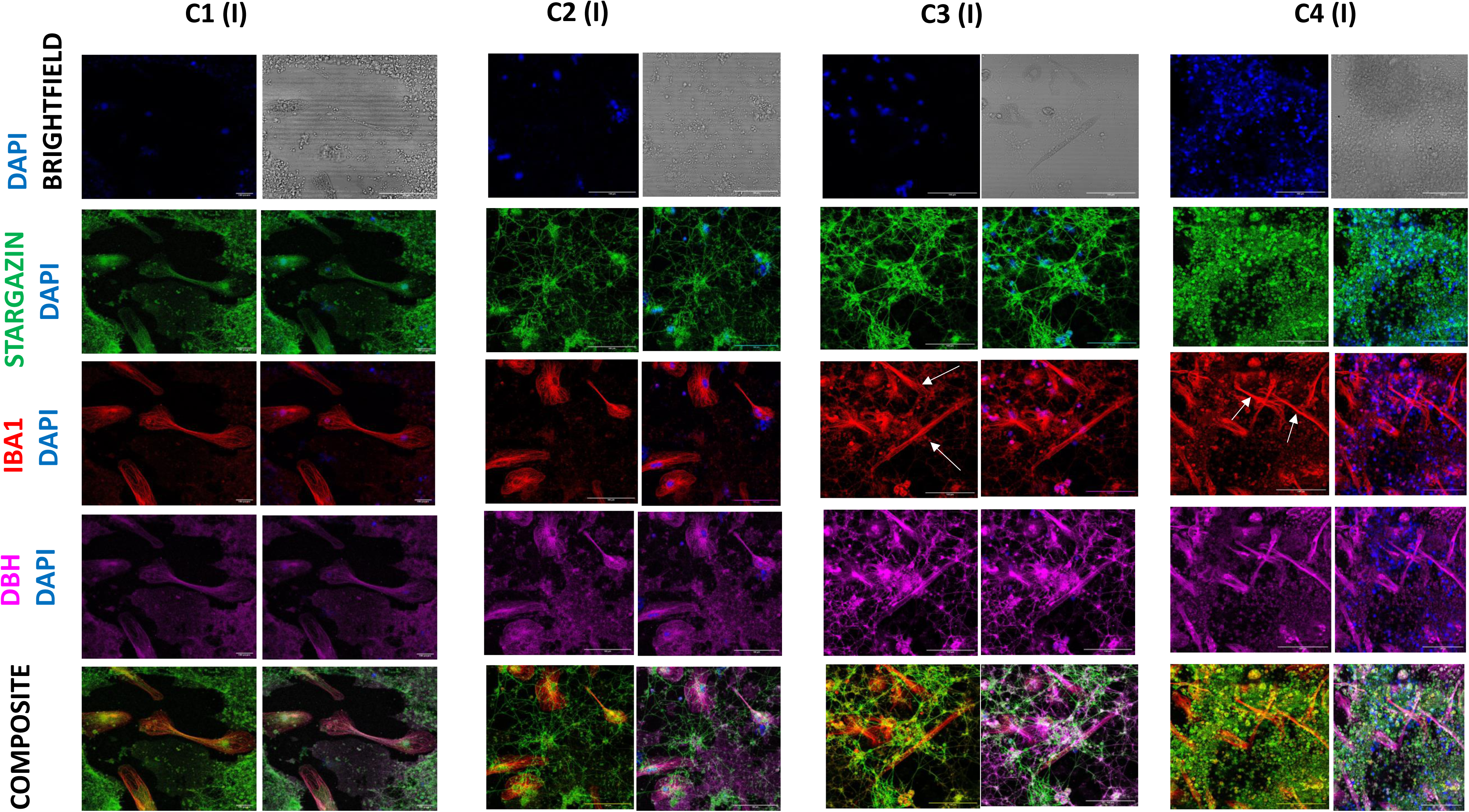
Pruning of synapses by Microglia: The figure depicts the expression Stargazin (green), a marker of synaptic functionality, Iba-1 microglia marker and DBH (pink) a marker of sympathetic (norepinephrine) neurons across different conditions at day 15. The panel shows the data of tissue field for the culture done in the wells identified as inwardly positioned (I). From C1-C4, the expression of Stargazin, Iba-1 and DBH is increasing. Notice the presence of tubular structures (white arrows) stained by Iba-1 (red) entangled with Stargazin and DBH positive neurons, these structures are micro blood vessels indicating the neurovascular crosstalk. Scale bars-100 µm

**Figure 3.**
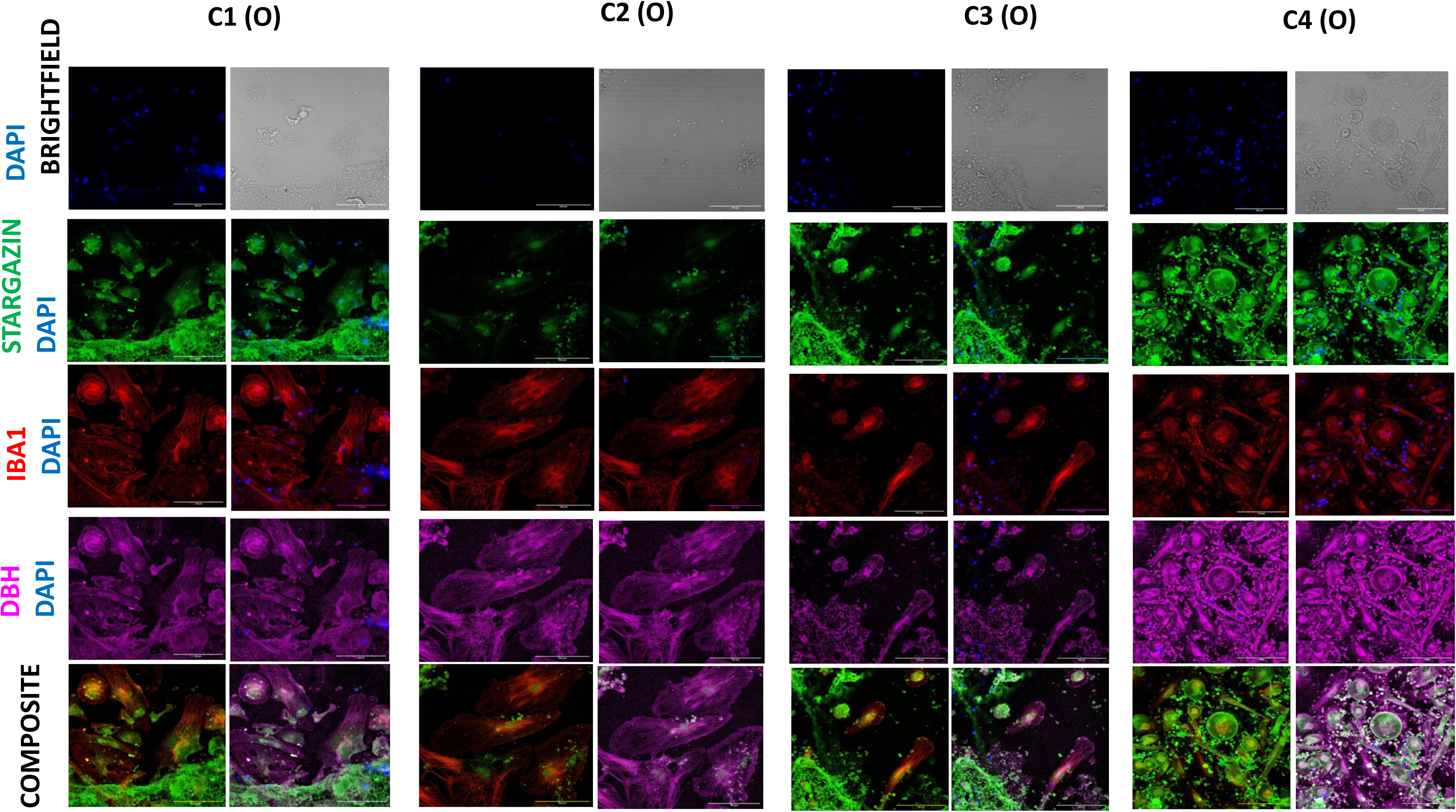
Pruning of synapses by Microglia: The figure depicts the expression Stargazin (green), a marker of synaptic functionality, Iba-1 microglia marker and DBH (pink) a marker of sympathetic (norepinephrine) neurons across different conditions at day 15. The panel shows the data of cellular field for the culture done in the wells identified as outwardly positioned (O). From C1-C4, the expression of Stargazin and DBH is increasing but that of Iba-1 is decreasing. The composite indicates the reduction in the size of activated microglia as the culture condition varies from C1 to C4 corresponding to the formation of more mature synapses. Scale bars-100 µm

**Figure 4.**
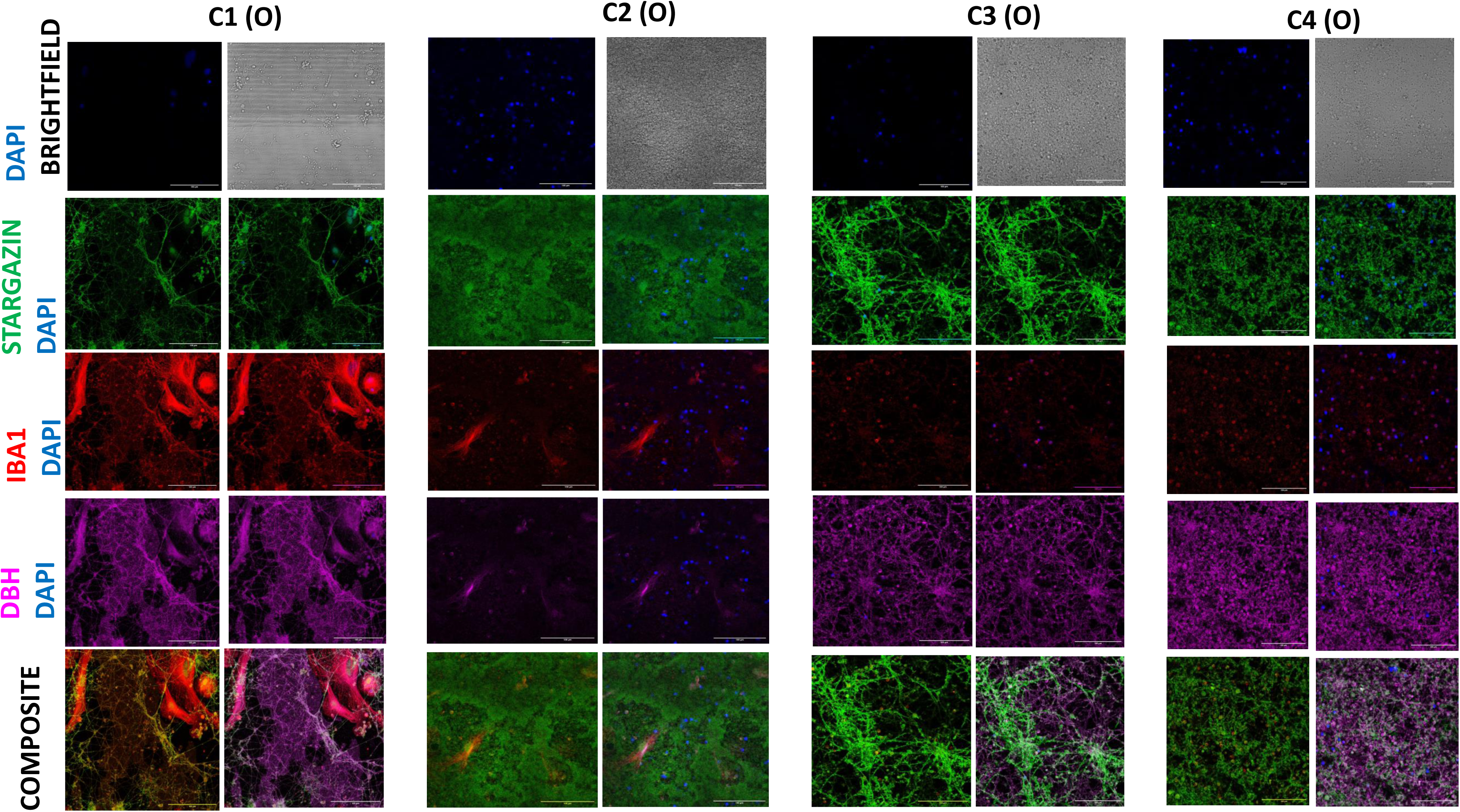
Pruning of synapses by Microglia: The figure depicts the expression Stargazin (green), a marker of synaptic functionality, Iba-1 microglia marker and DBH (pink) a marker of sympathetic (norepinephrine) neurons across different conditions at day 15. The panel shows the data of neuronal field for the culture done in the wells identified as outwardly positioned (O). From C1-C4, the expression of Stargazin and DBH is increasing but that of Iba-1 is decreasing. Scale bars-100 µm

**Figure 5.**
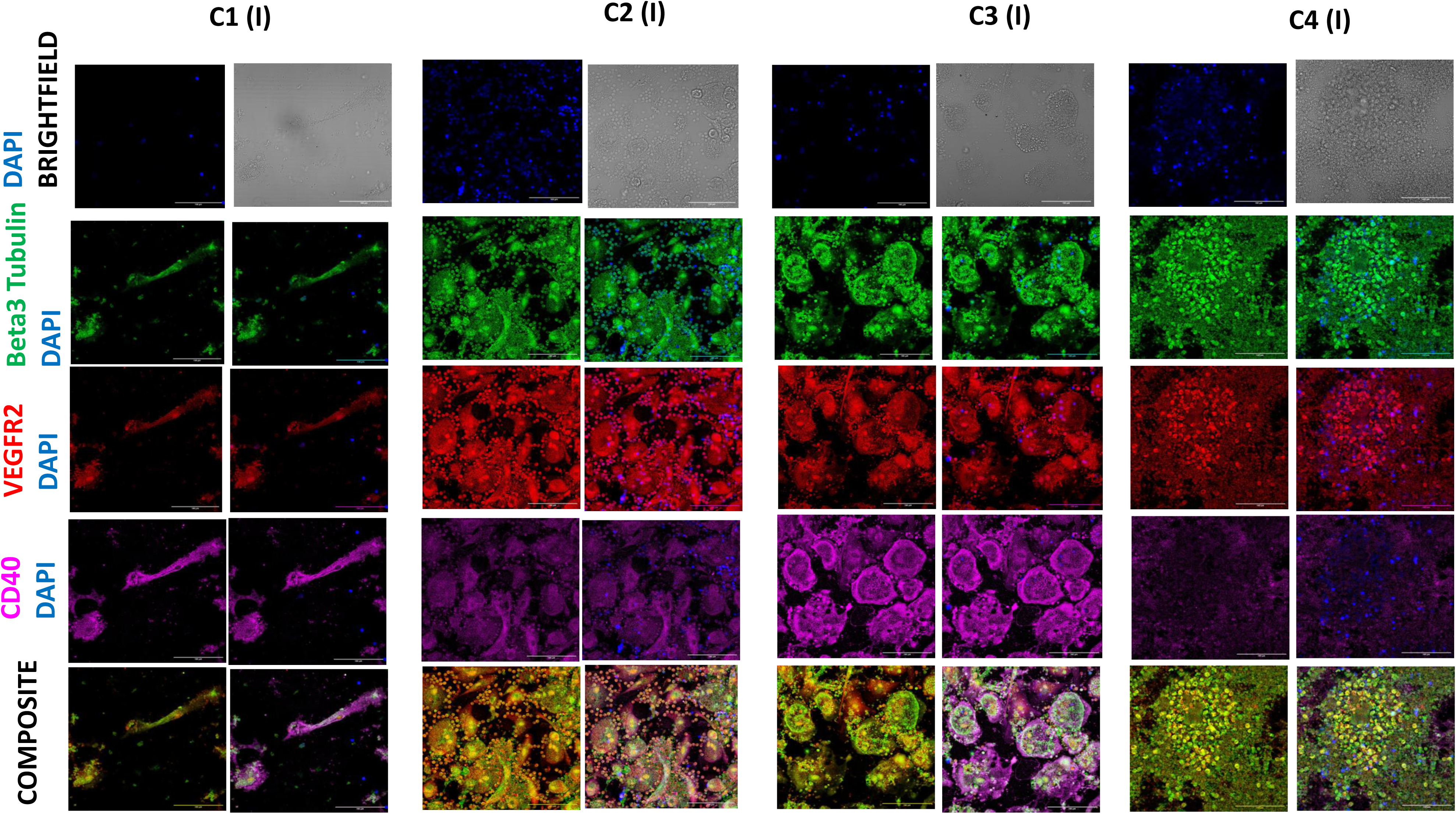
Neuro-vascular unit formation: The figure depicts the expression Beta3 Tubulin (green), a marker of early formed neurons, VEGFR2 a marker predominantly expressed by endothelial cells which is primarily involved in angiogenesis CD40 (pink), a co stimulatory protein expressed by activated macrophages across different conditions at day 15. The panel shows the data of the cellular field for the culture done in the wells identified as inwardly positioned (I). From C1-C4, the expression of Beta3 Tubulin and VEGFR2 is increasing however the expression of CD40 is decreasing. Notice the composite images of VEGFR2 and Beta3 Tubulin that depicts that coexistence of neural and vascular progenies. Scale bars-100 µm

**Figure 6.**
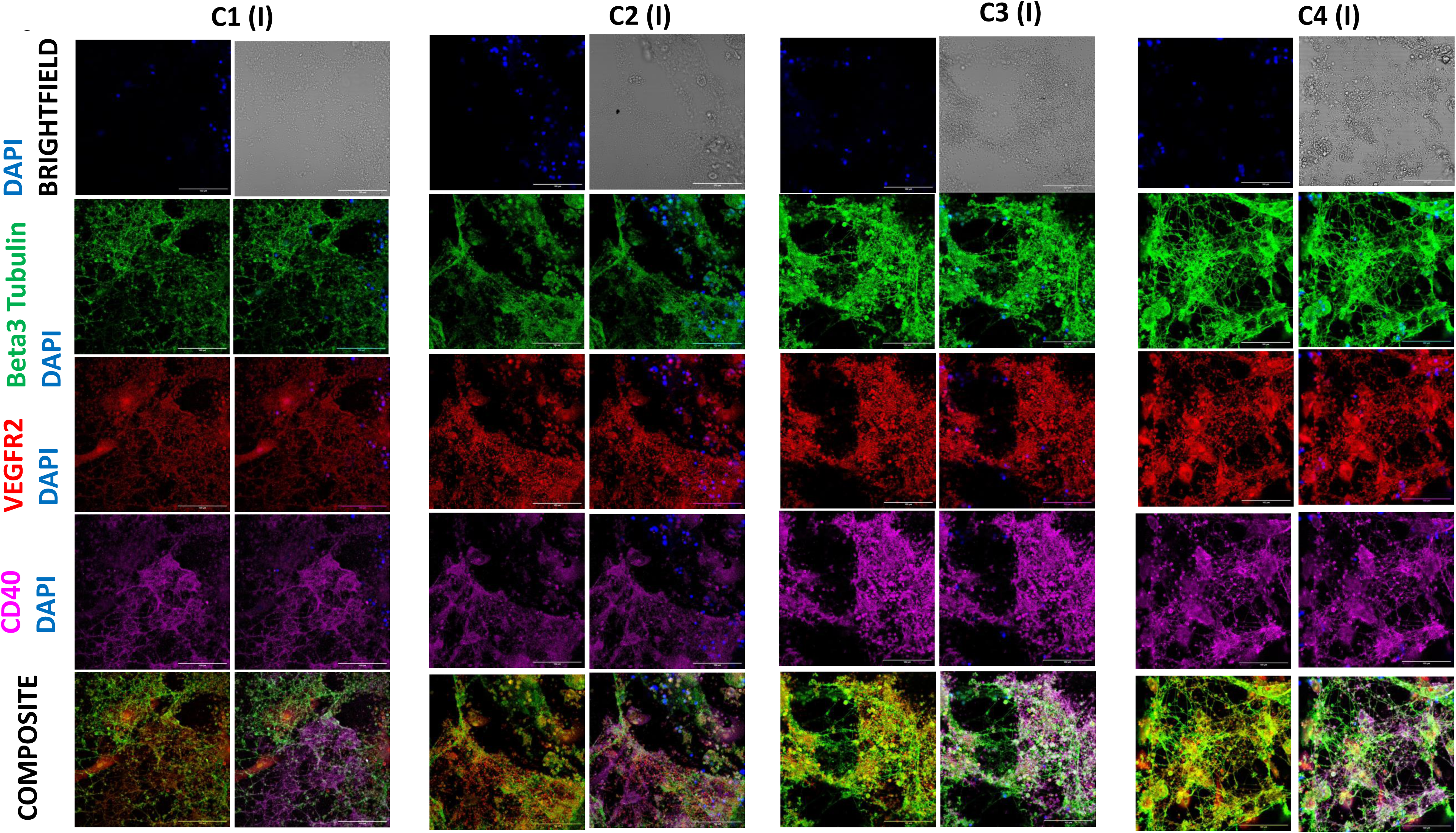
Neuro-vascular unit formation: The figure depicts the expression Beta3 Tubulin (green), a marker of early formed neurons, VEGFR2 a marker predominantly expressed by endothelial cells which is primarily involved in angiogenesis CD40 (pink), a co stimulatory protein expressed by activated macrophages across different conditions at day 15. The panel shows the data of the neuronal field for the culture done in the wells identified as inwardly positioned (I). From C1-C4, the expression of Beta3 Tubulin and VEGFR2 is increasing however the expression of CD40 is decreasing. Notice the composite images of VEGFR2 and Beta3 Tubulin that depicts that coexistence of neural and vascular progenies. Scale bars-100 µm

**Figure 7.**
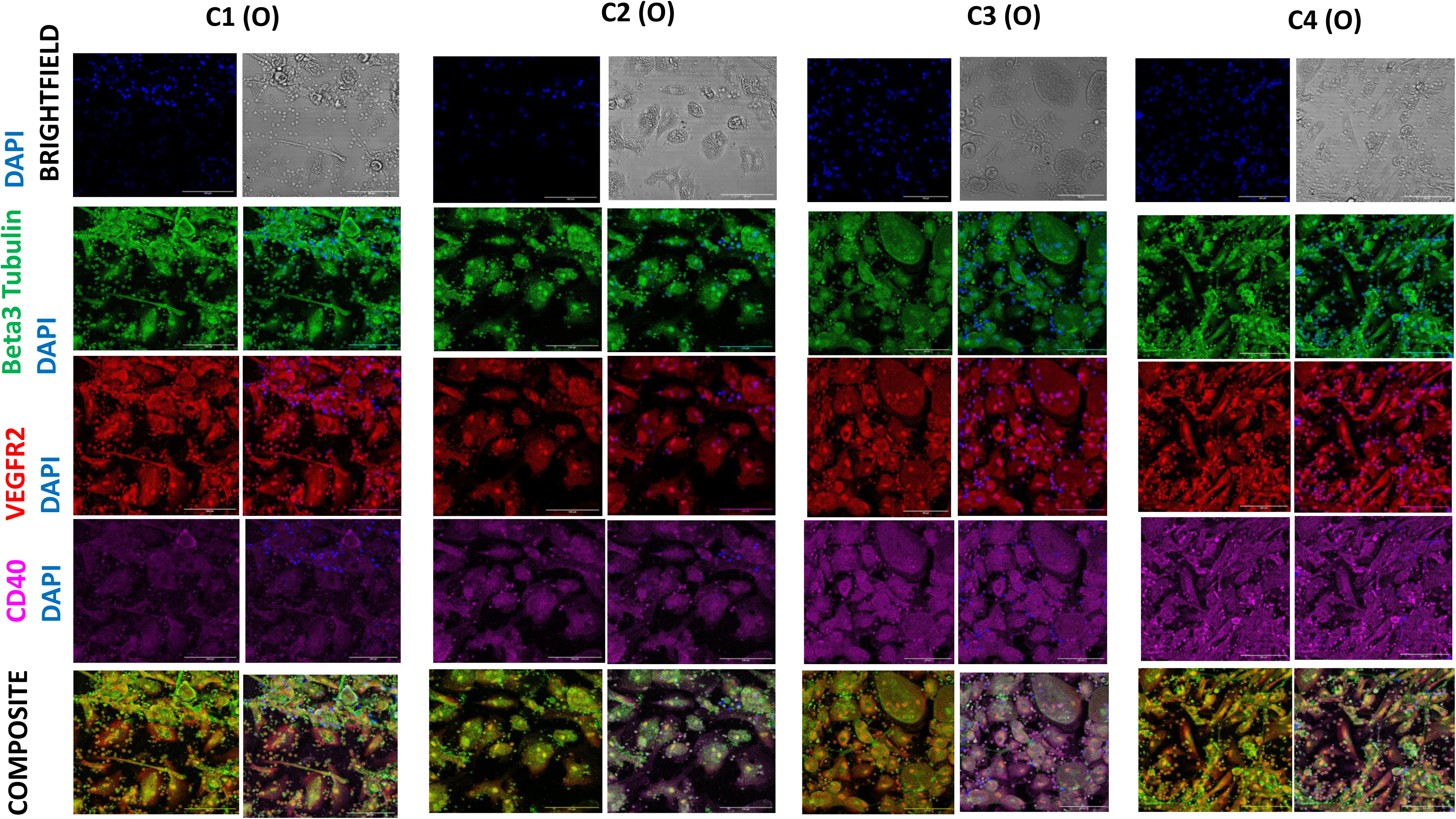
Neuro-vascular unit formation: The figure depicts the expression Beta3 Tubulin (green), a marker of early formed neurons, VEGFR2 a marker predominantly expressed by endothelial cells which is primarily involved in angiogenesis CD40 (pink), a co stimulatory protein expressed by activated macrophages across different conditions at day 15. The panel shows the data of the cellular field for the culture done in the wells identified as outwardly positioned (O). From C1-C4, the expression of Beta3 Tubulin, VEGFR2 and CD40 is. Notice the composite images of VEGFR2 and Beta3 Tubulin that depicts that coexistence of neural and vascular progenies. Scale bars-100 µm

**Figure 8.**
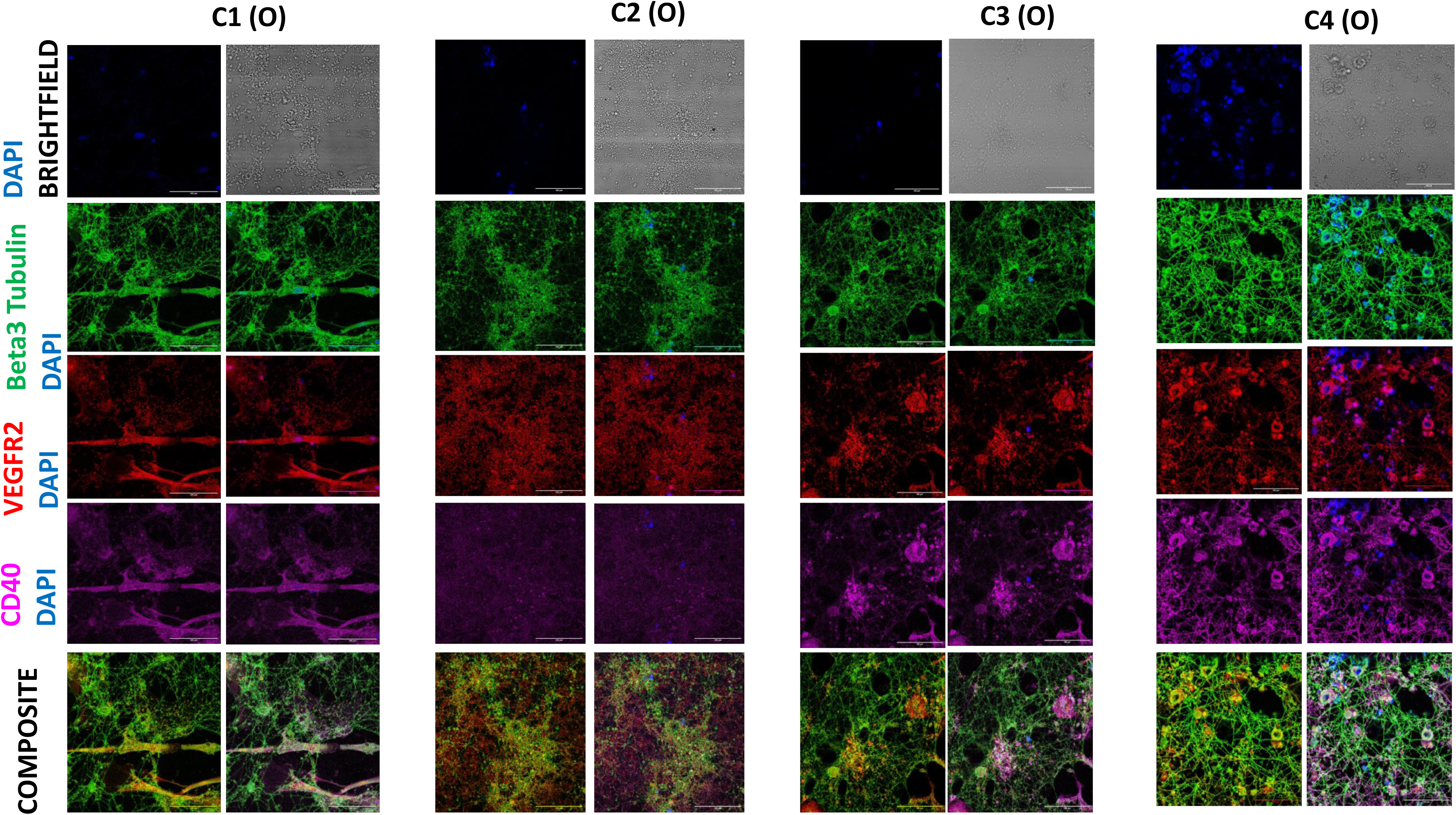
Neuro-vascular unit formation: The figure depicts the expression Beta3 Tubulin (green), a marker of early formed neurons, VEGFR2 a marker predominantly expressed by endothelial cells which is primarily involved in angiogenesis CD40 (pink), a co-stimulatory protein expressed by activated macrophages across different conditions at day 15. The panel shows the data of the neuronal field for the culture done in the wells identified as outwardly positioned (O). From C1-C4, the expression of Beta3 Tubulin, VEGFR2 and CD40 is. Notice the composite images of VEGFR2 and Beta3 Tubulin that depicts that coexistence of neural and vascular progenies. Scale bars-100 µm

**Figure 9.**
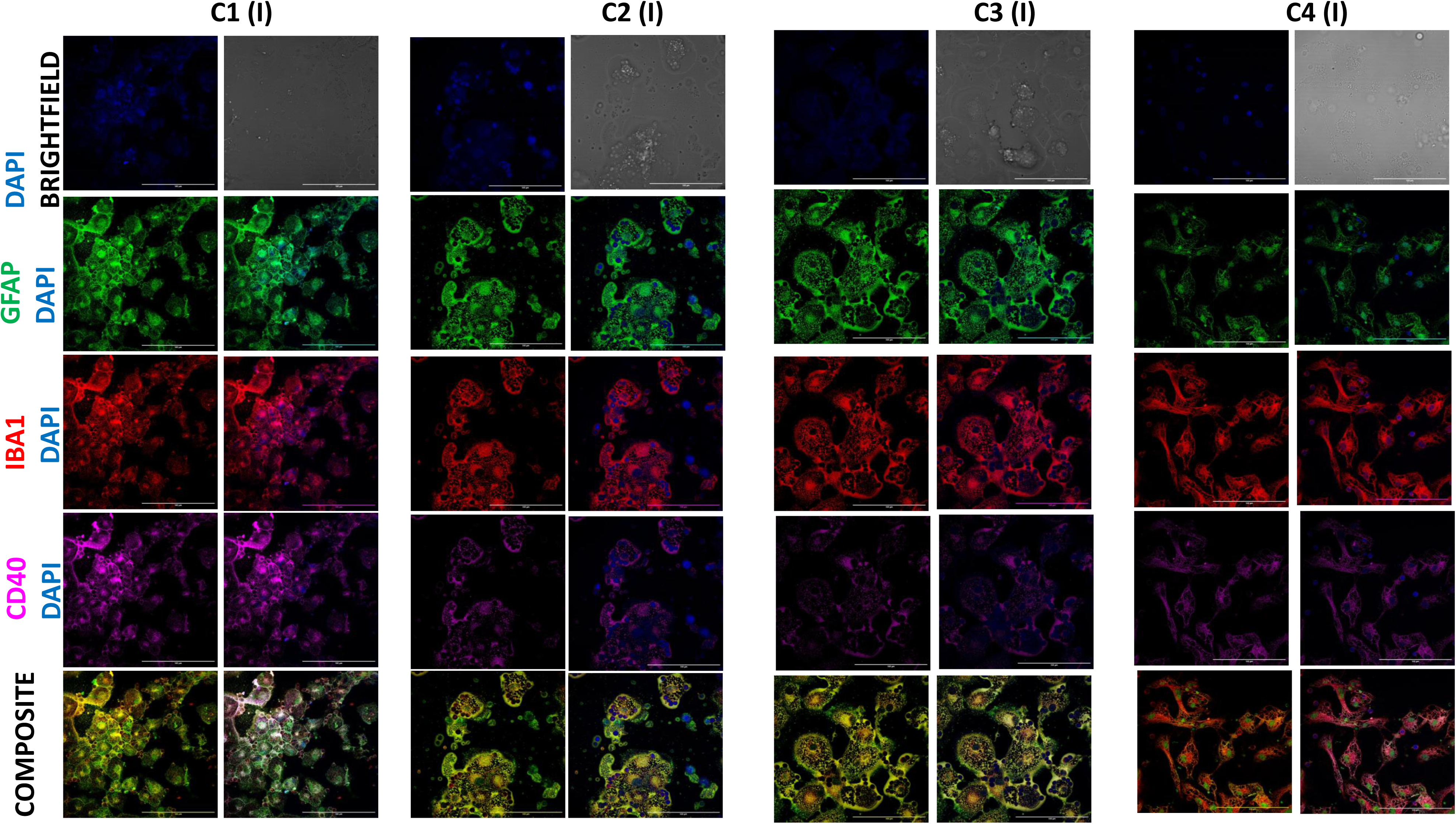
Astrocytic and Microglia niches: The figure depicts the expression GFAP (green), a marker of astrocytes, Iba-1 a microglia marker CD40 (pink), a co stimulatory protein expressed by activated macrophages across different conditions at day 15. The panel shows the data of the cellular field for the culture done in the wells identified as inwardly positioned (I). From C1-C4, the expression of GFAP and CD40 is decreasing and Iba-1 remains consistent. Despite the general trend notice the localization of GFAP is decreasing from C1C4 and its expression reduces to a small circle surrounded by the microglia with filamentous appearance in C4. Scale bars-100 µm

**Figure 10.**
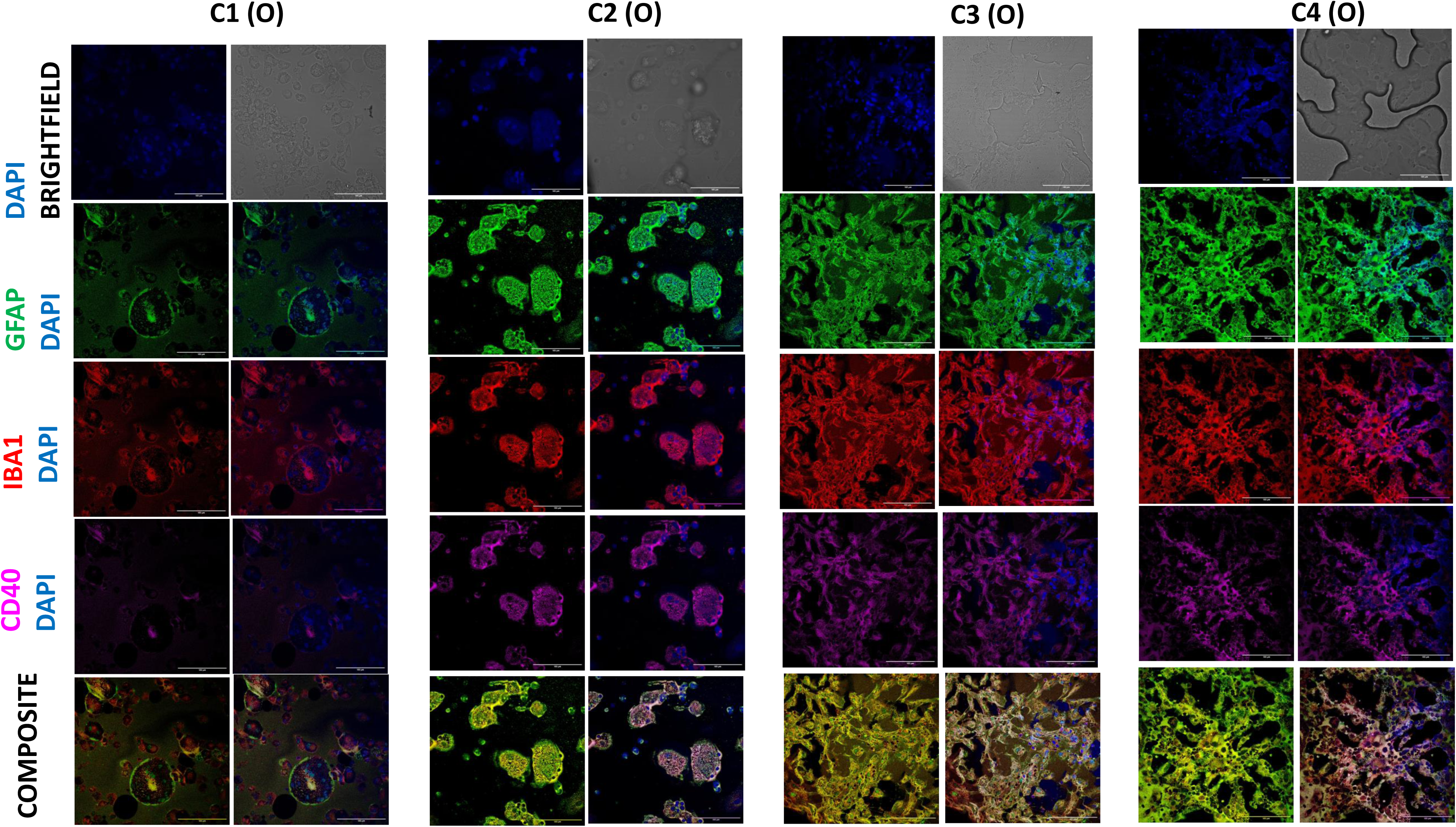
Astrocytic and Microglia niches: The figure depicts the expression GFAP (green), a marker of astrocytes, Iba-1 a microglia marker CD40 (pink), a co stimulatory protein expressed by activated macrophages across different conditions at day 15. The panel shows the data of the cellular field for the culture done in the wells identified as outwardly positioned (O). From C1-C4, the expression of GFAP and CD40 is increasing and Iba-1 remains consistent. Scale bars-100 µm

**Figure 11.**
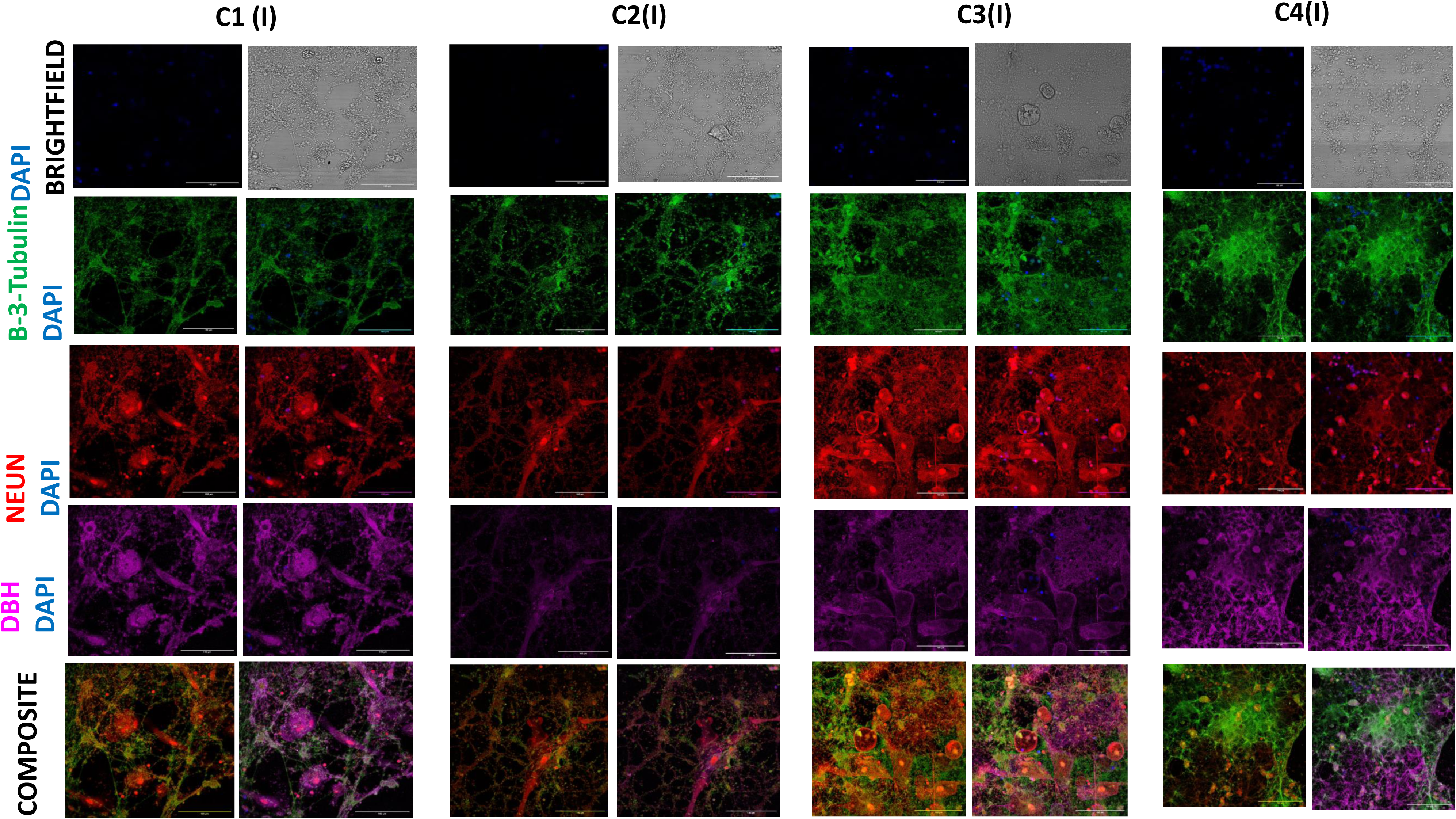
Neuronal differentiation and maturation: The figure depicts the expression Beta 3 tubulin(green), a marker of early neuron differentiation, Neun (red) a early neuron marker and DBH (pink) a marker of sympathetic (norepinephrine) neurons across different conditions at day 15. The panel shows the data of cellular field for the culture done in the wells identified as inwardly positioned (I). From C1-C4, the expression of Beta 3 tubulin, Neun and DBH increases. The composite indicates that the culture is more mature in C4 condition with the neurons forming more tissue like appearance. Scale bars-100 µm

**Figure 12.**
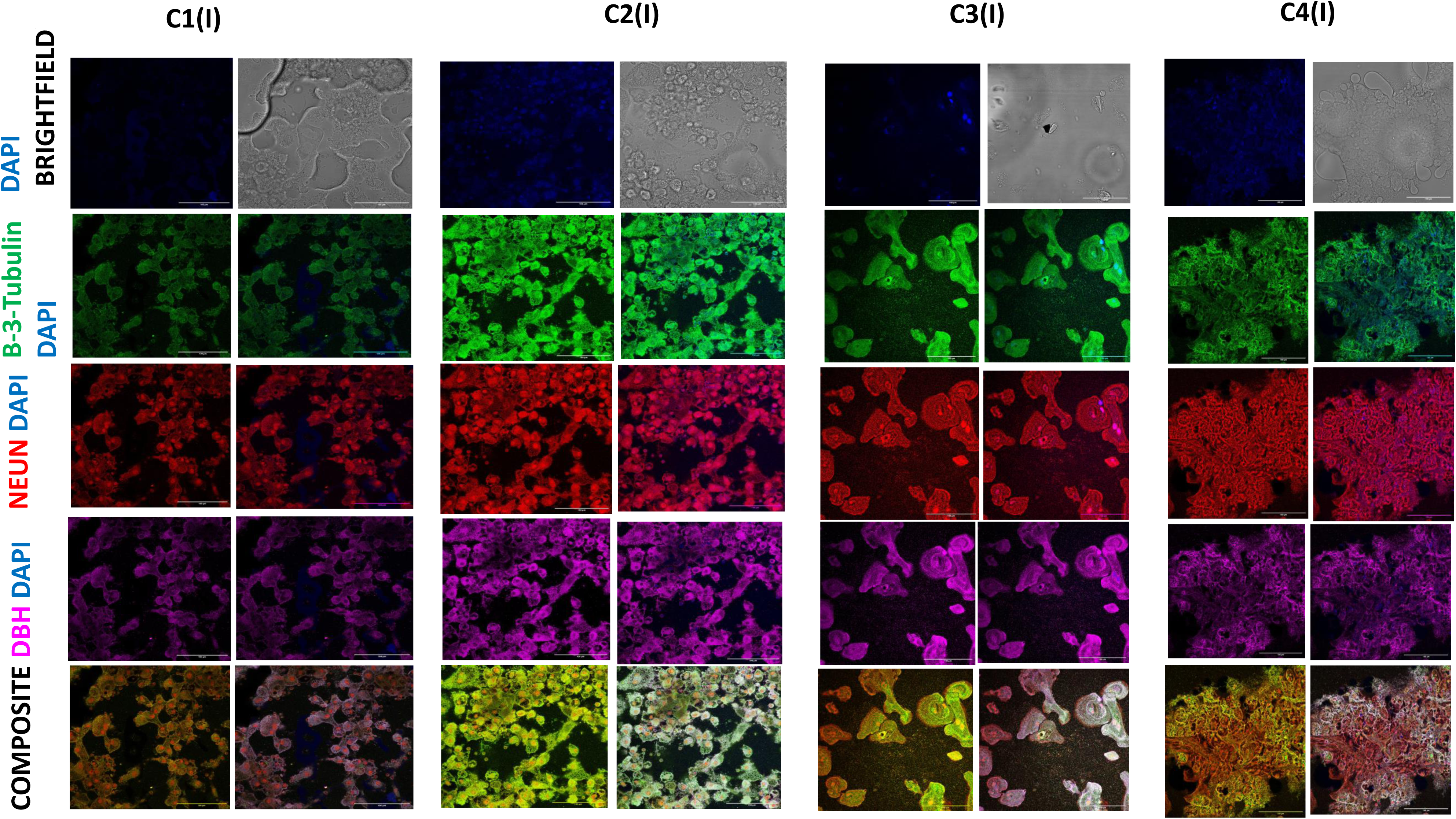
Neuronal differentiation and maturation: The figure depicts the expression Beta 3 tubulin(green), a marker of early neuron differentiation, Neun (red) a early neuron marker and DBH (pink) a marker of sympathetic (norepinephrine) neurons across different conditions at day 15. The panel shows the data of tissue containing field for the culture done in the wells identified as inwardly positioned (I). From C1-C4, the expression of Beta 3 tubulin, Neun and DBH increases. The composite indicates that the culture is more mature in C4 condition with the neurons forming more tissue like appearance. Scale bars-100 µm

**Figure 13.**
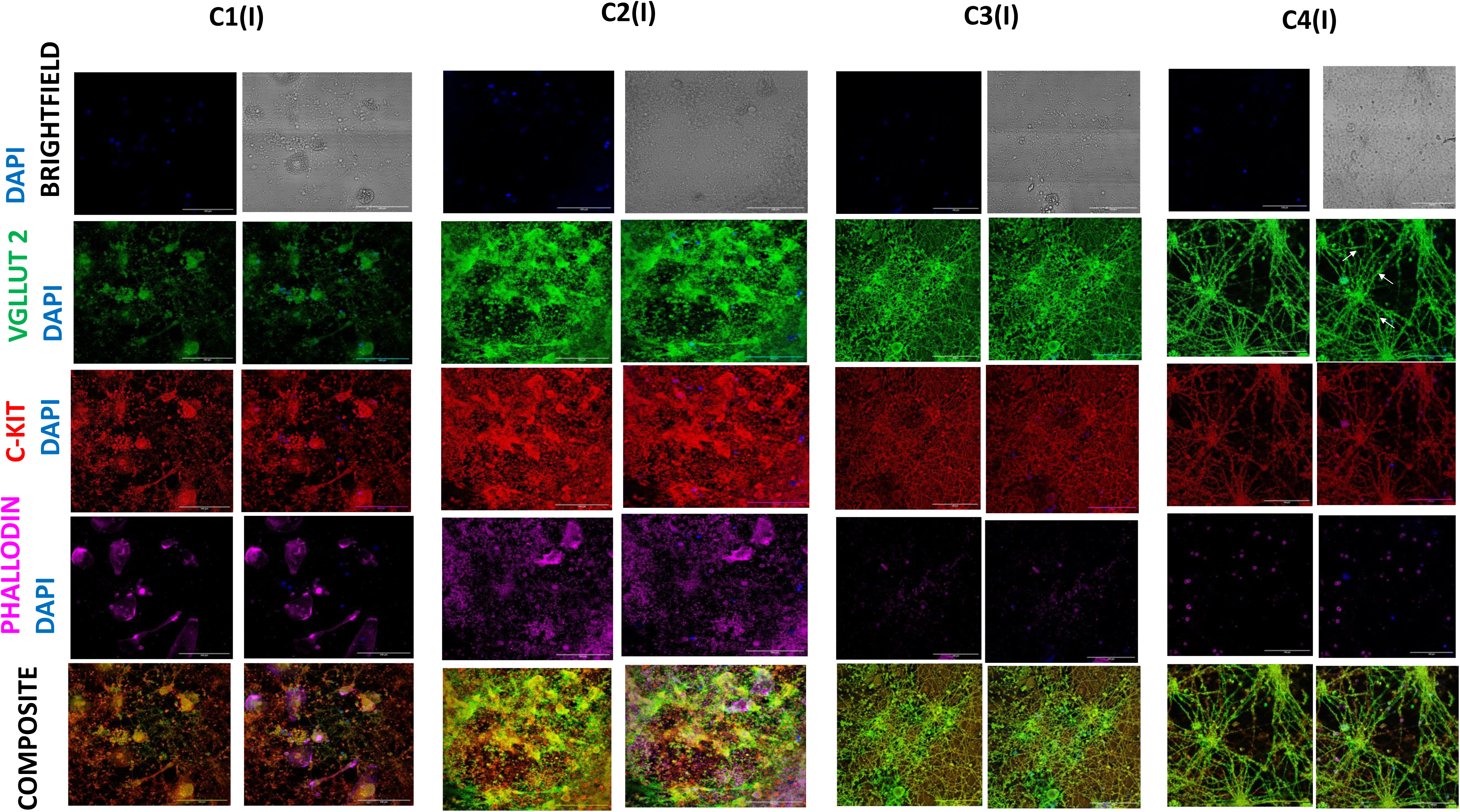
Neuro-vascular cross talk: The figure depicts the expression VGLUT2 (green), a marker of glutamatergic neurons, C-Kit (red) hematopoietic stem cell marker and Phalloidin (pink) used to stained actin filaments across different conditions at day 15. The panel shows the data of cellular field for the culture done in the wells identified as inwardly positioned (I). From C1-C4, the expression of VGLUT2 and C-Kit increases and Phalloidin decreases. The composite indicates that hematopoietic progenitor are promoting neurogenesis. Scale bars-100 µm

**Figure 14.**
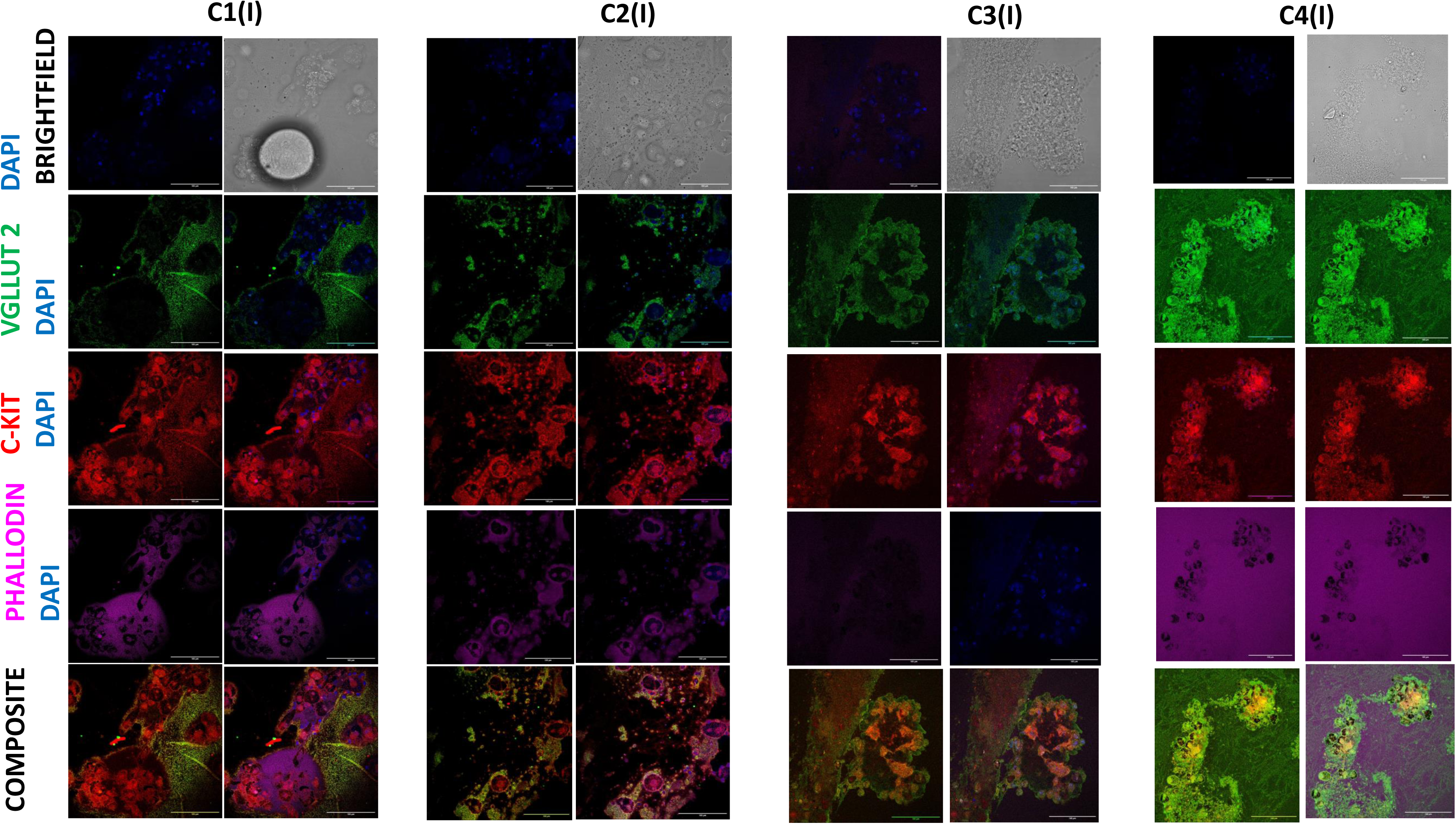
Neuro-vascular cross talk: The figure depicts the expression VGLUT2 (green), a marker of glutamatergic neurons, C-Kit (red) hematopoietic stem cell marker and Phalloidin (pink) used to stained actin filaments across different conditions at day 15. The panel shows the data of tissue containing field for the culture done in the wells identified as inwardly positioned (I). From C1-C4, the expression of VGLUT2 and C-Kit increases and Phalloidin decreases. The composite indicates that hematopoietic progenitor are promoting neurogenesis. Notice as the extent of tissue formation increases, the phalloidin expression decreases. Scale bars-100 µm

**Figure 15.**
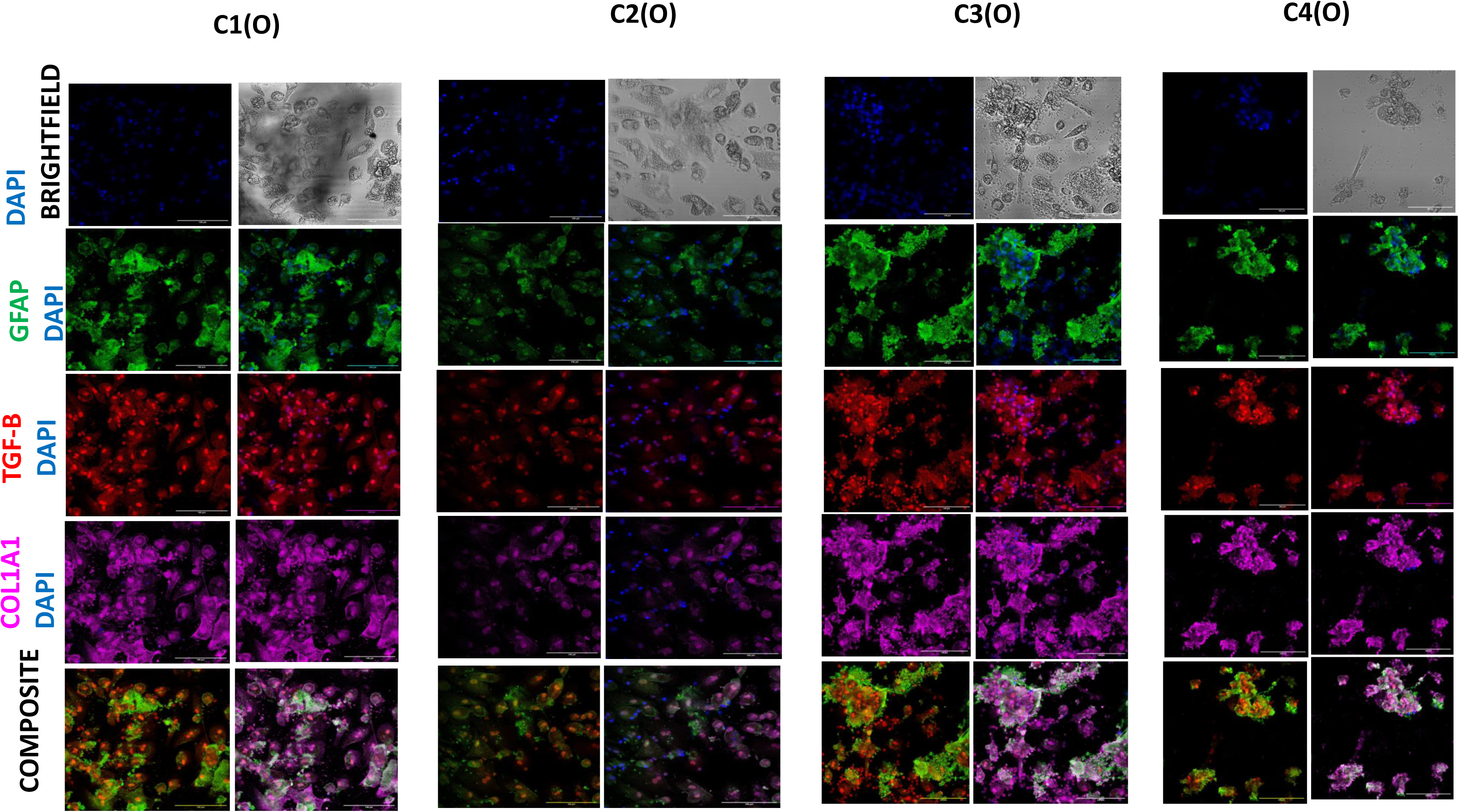
Role of TGF-Beta signalling and developing vasculature: The figure depicts the expression GFAP (green), astrocytic marker, TGF-beta (red) signalling which is involved in cerebral angiogenesis, Col1A1 (pink)a collagen marker to check the structural integrity of developing vasculature all these markers is across different conditions at day 15. The panel shows the data of cellular field of the culture done in the wells identified as outwardly positioned (O). From C1-C4, the expression of GFAP, TGF-beta and Col1A1 increases. Scale bars-100 µm

**Figure 16.**
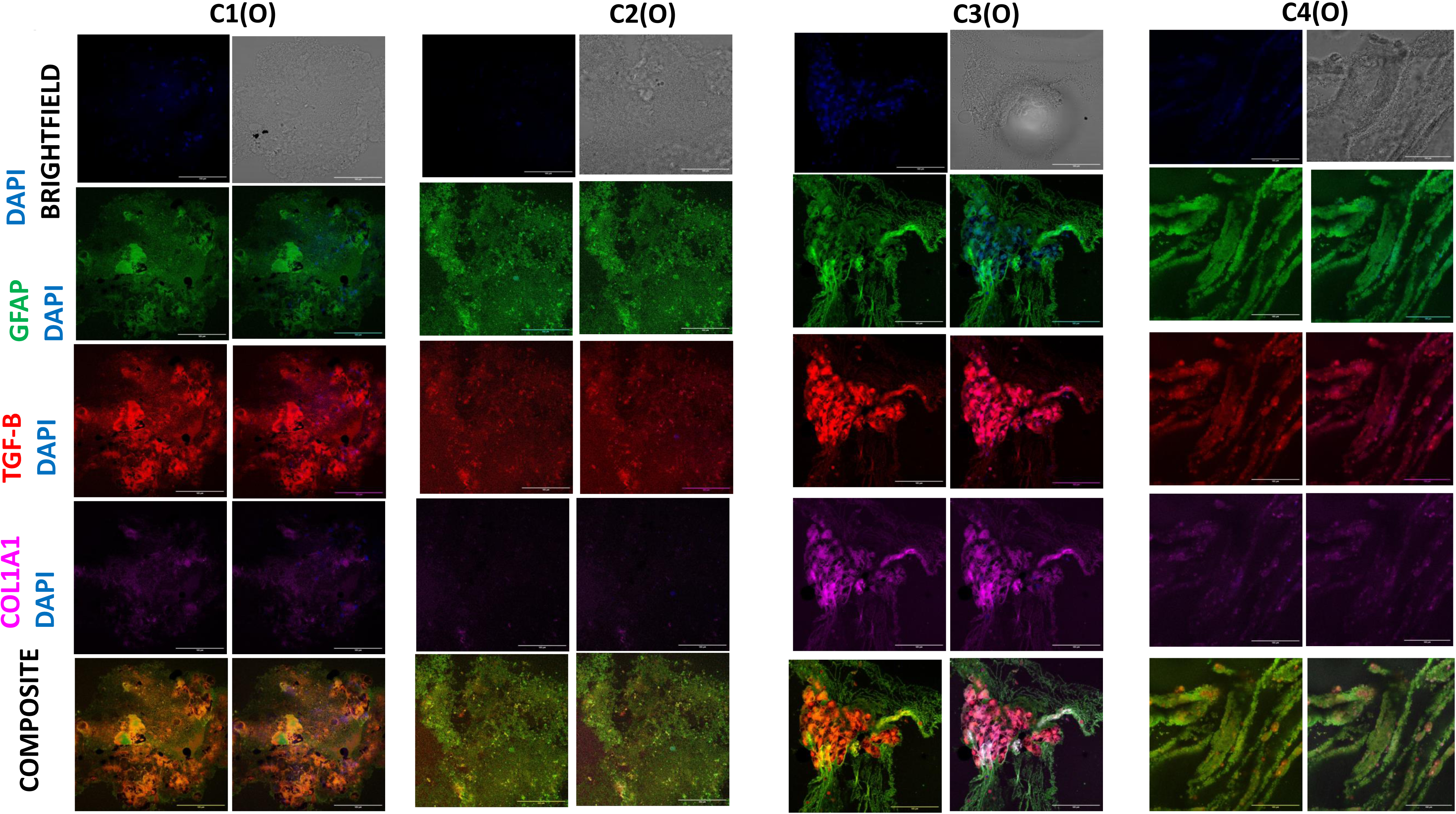
Role of TGF-Beta signalling and developing vasculature in astrogenesis: The figure depicts the expression GFAP (green), astrocytic marker, TGF-beta (red) signalling which is involved in cerebral angiogenesis, Col1A1 (pink)a collagen marker to check the structural integrity of developing vasculature all these markers is across different conditions at day 15. The panel shows the data of tissue field of the culture done in the wells identified as outwardly positioned (O). From C1-C4, the expression of GFAP, TGF-beta and Col1A1 increases. Despite the general trend notice that the expression of TGF-Beta is comparatively more than GFAP initially but GFAP’s expression gradually increases with the incresing expression of TGF-beta signal across the various conditions indicating the role of vasculature in promoting astrogenesis. Scale bars-100 µm

**Figure 17.**
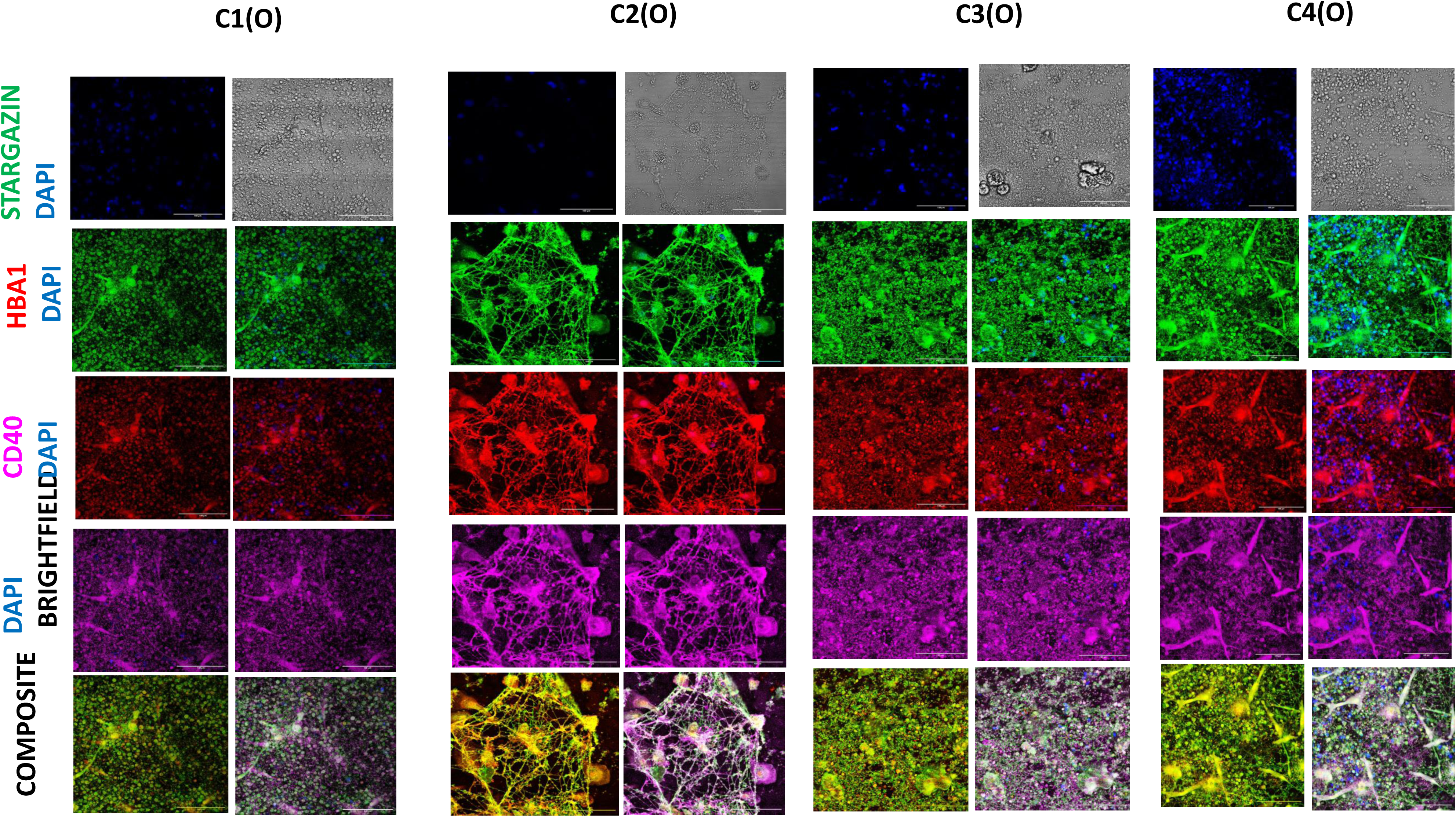
Neuro-inflammatory niches: The figure depicts the expression Stargazin (green), a marker of synaptic functionality, Hba-1 expressed by differentiating erythroid cells marker and CD40 (pink) a co stimulatory protein expressed by activated macrophages across different conditions at day 15. The panel shows the data of neuronal field for the culture done in the wells identified as outwardly positioned (O). From C1-C4, the expression of Stargazin,Hba-1 and CD40 is increasing. Increasing Hba-1 levels from C1-C4 indicates maximum erythropoiesis in C4 condition. Rising expression of CD40 from C1-C4 highlights the presence of active immune niches in the system. Scale bars-100 µm

**Figure 18.**
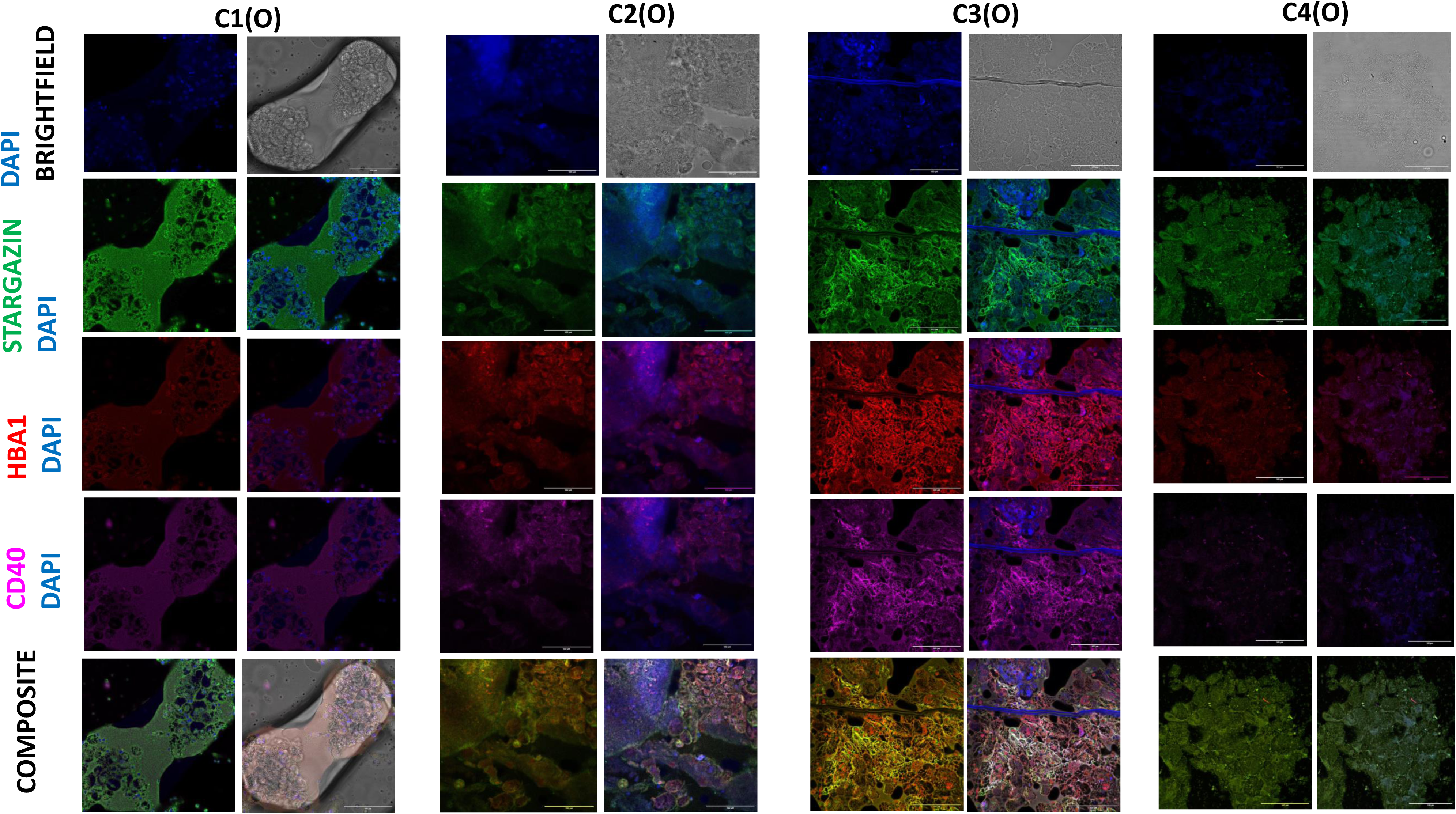
Neuro-inflammatory niches: The figure depicts the expression Stargazin (green), a marker of synaptic functionality, Hba-1 expressed by differentiating erythroid cells marker and CD40 (pink) a co stimulatory protein expressed by activated macrophages across different conditions at day 15. The panel shows the data of tissue field for the culture done in the wells identified as outwardly positioned (O). From C1-C4, the expression of Stargazin, Hba-1 and CD40 is increasing. There is more formation of neuro vascular tissue in C4 condition. Scale bars-100 µm

**Figure 19.**
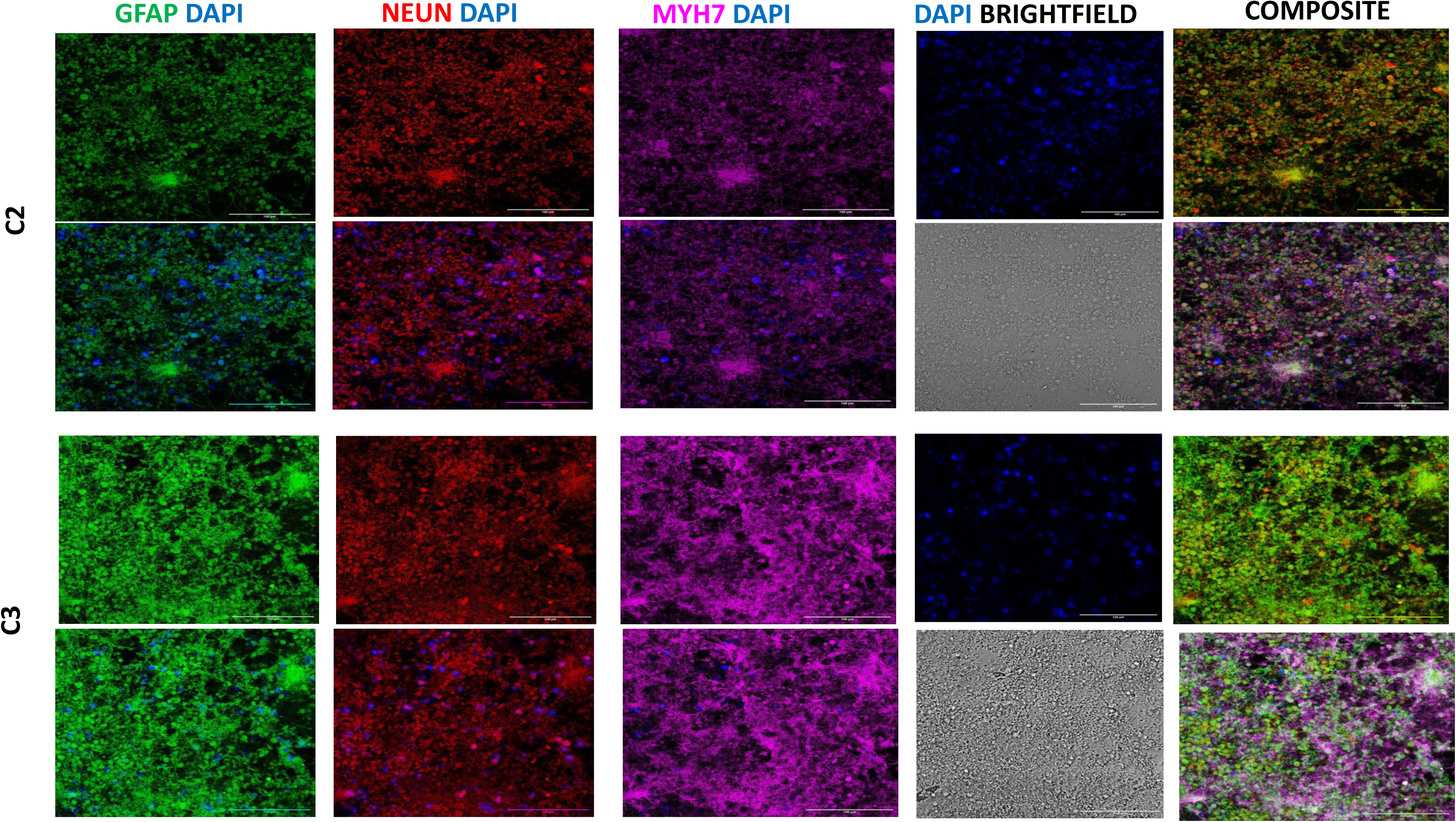
Presence of mesodermal progenitors with neurons: The figure depicts the expression GFAP (green), astrocytic marker, Neun (red) expressed a early neuron marker and MYH 7 (pink) a crucial component of cardiac muscles here it indicates mesodermal progenitors in the culture across different conditions at day 15. The expression of GFAP, Neun and Myh7is increasing in C3 condition. Scale bars-100 µm

**Figure 20.**
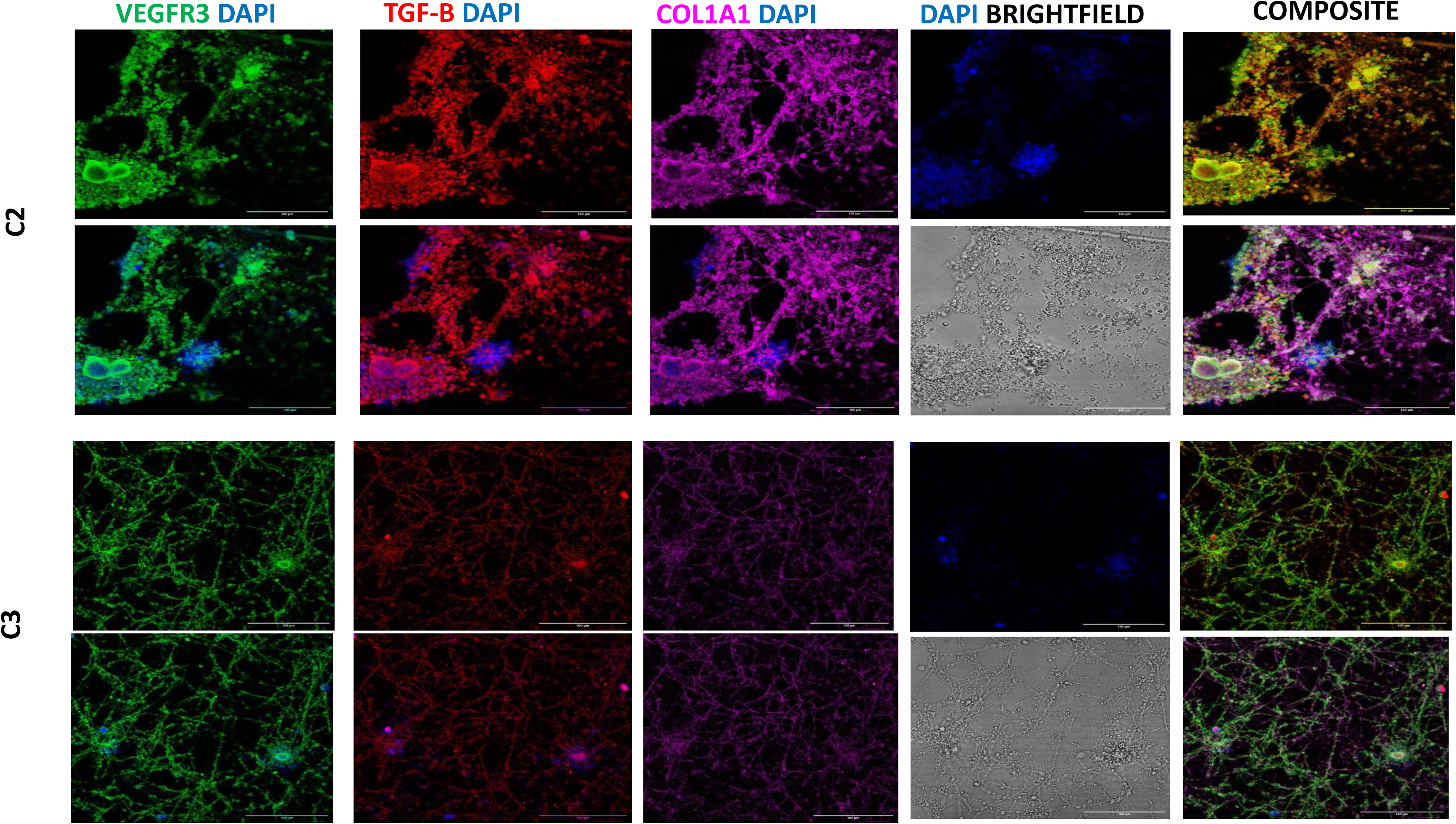
The figure depicts the expression VEGFR3 (green),known for its role in lymphangiogenesis and vascular function, TGF-beta (red) signalling which is involved in cerebral angiogenesis, Col1A1 (pink)a collagen marker to check the structural integrity of developing vasculature all these markers across different conditions at day 15. Scale bars-100 µm

**Figure 21.**
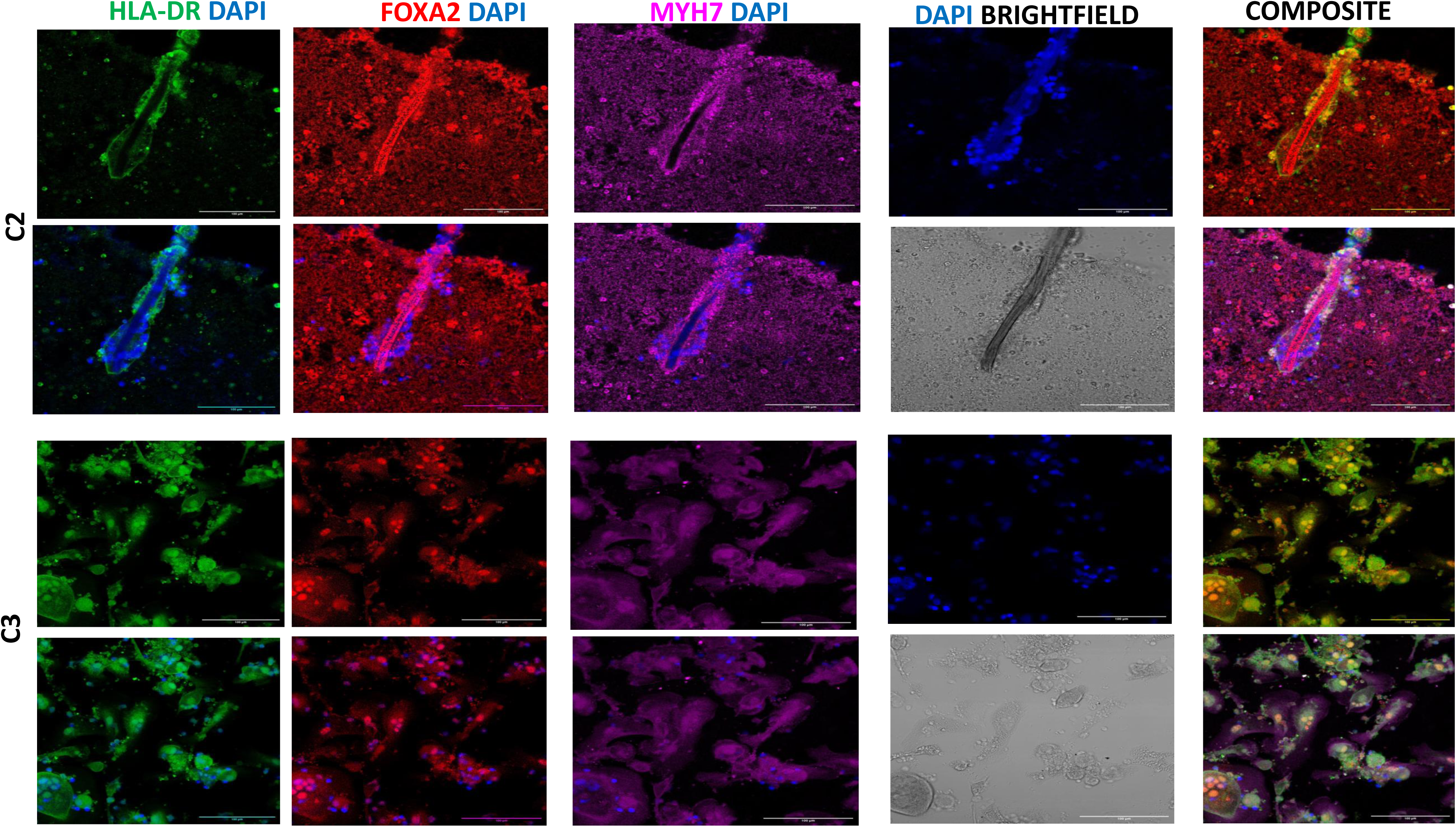
The figure depicts the expression HLA-DR (green) expressed by activated microglia, FoxA2 (red) marker of promoting mid brain neurogenesis and MYH 7 (pink) a crucial component of cardiac muscles here it indicates mesodermal progenitors in the culture across different conditions at day 15. Scale bars-100 µm

**Figure 22.**
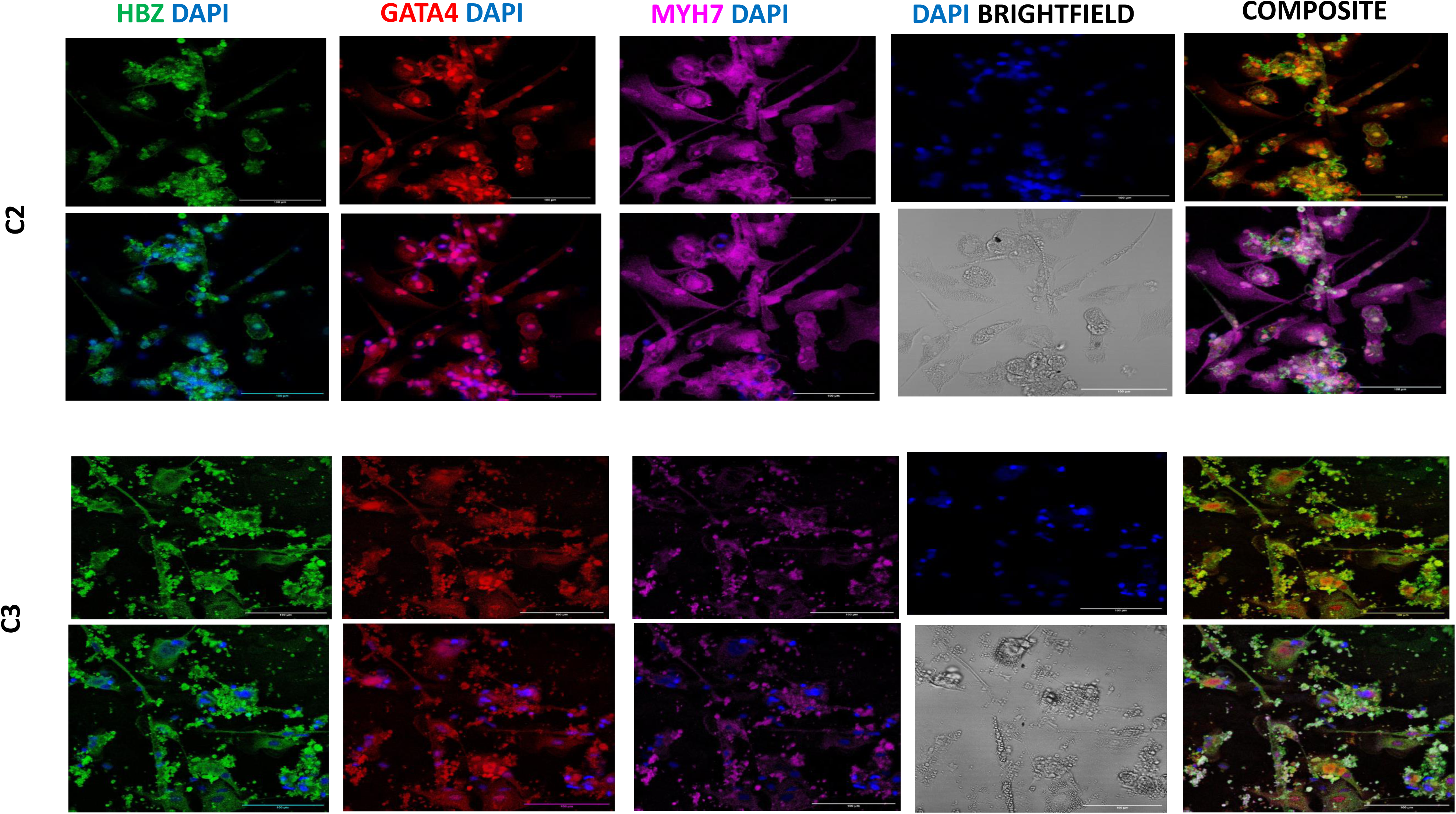
The figure depicts the expression HBZ (green) expressed by activated microglia, GATA4 (red), a transcription factor involved in neural crest and craniofacial skeleton development and MYH 7 (pink) a crucial component of cardiac muscles here it indicates mesodermal progenitors in the culture across different conditions at day 15. Scale bars-100 µm

**Figure 23.**
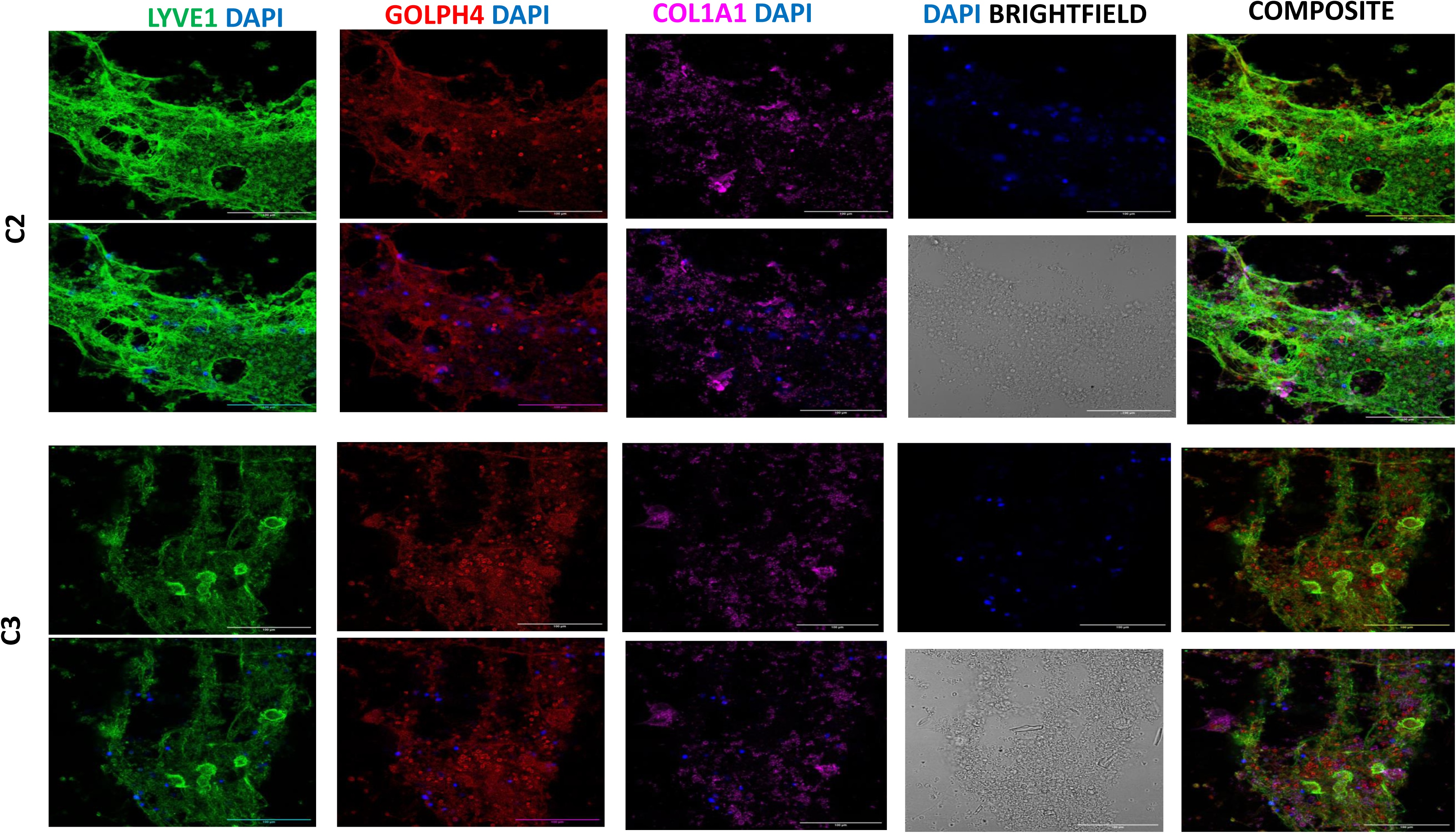
The figure depicts the expression LYVE1 (green) receptor involved in lymphatic endothelial cells, GOLPH4 (red), is a Golgi-associated protein involved protein sorting and trafficking and Col1A1 (pink) a collagen marker to check the structural integrity of developing vasculature in the culture across different conditions at day 15. Scale bars-100 µm

**Figure 24.**
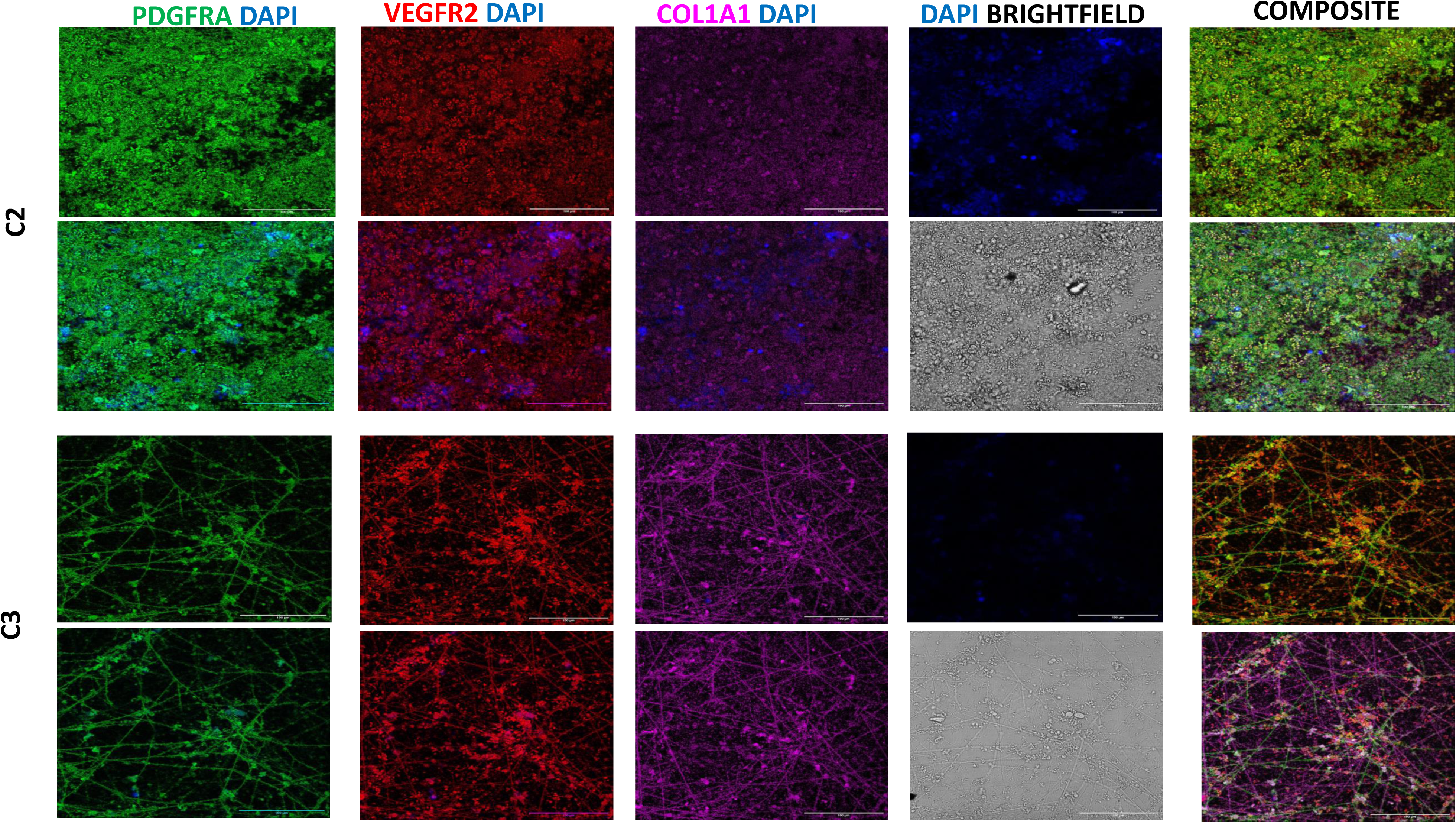
The figure depicts the expression PDGFR Alpha (green) receptor involved in lymphatic endothelial cells, VEGFR2 (red) a marker predominantly expressed by endothelial cells which is primarily involved in angiogenesis, and Col1A1 (pink) a collagen marker to check the structural integrity of developing vasculature in the culture across different conditions at day 15. Scale bars-100 µm

**Figure 25.**
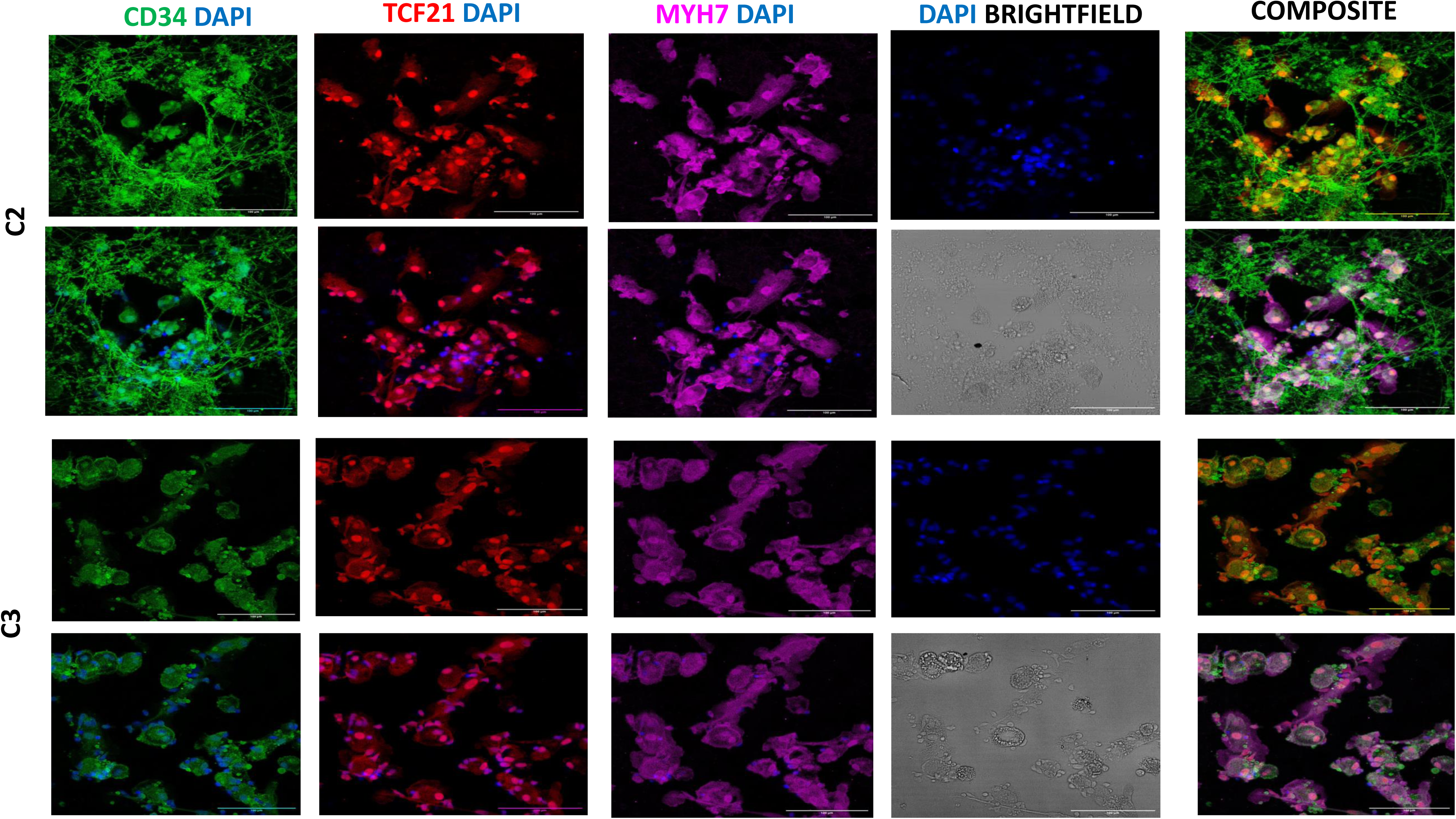
The figure depicts the expression CD34 (green) expressed by hematopoietic stem cells, TCF 21(red) marker of mesenchymal stem cells and MYH 7 (pink) a crucial component of cardiac muscles here it indicates mesodermal progenitors in the culture across different conditions at day 15. Scale bars-100 µm

**Figure 26.**
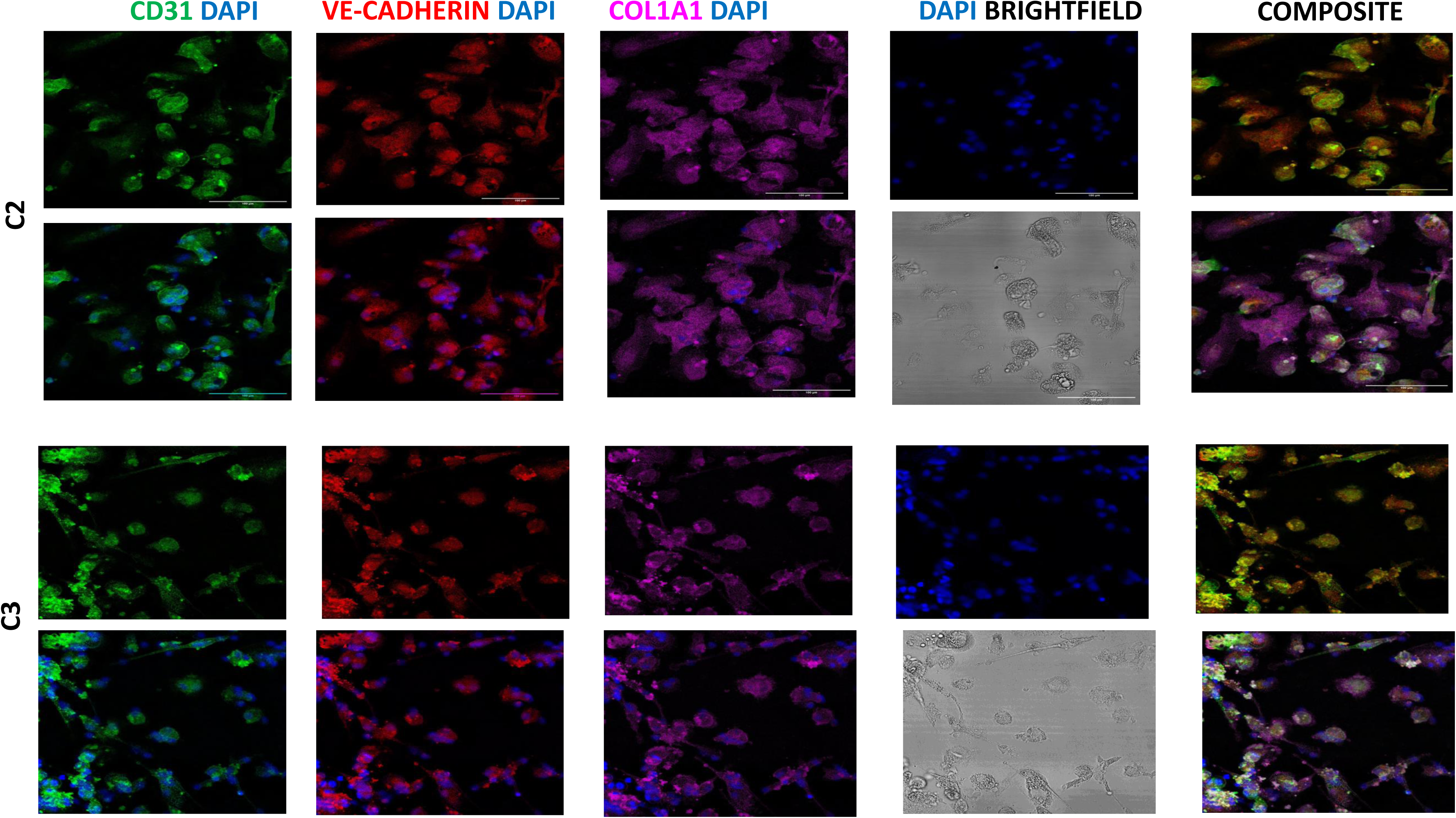
The figure depicts the expression CD31 (green) expressed by endothelial cells,VE cadherin (red) expressed in endothelial cells and are vital for maintaining the integrity of the endothelial barrier and Col1A1 (pink) a collagen marker to check the structural integrity of developing vasculature in the culture across different conditions at day 15. Scale bars-100 µm

**Figure 27.**
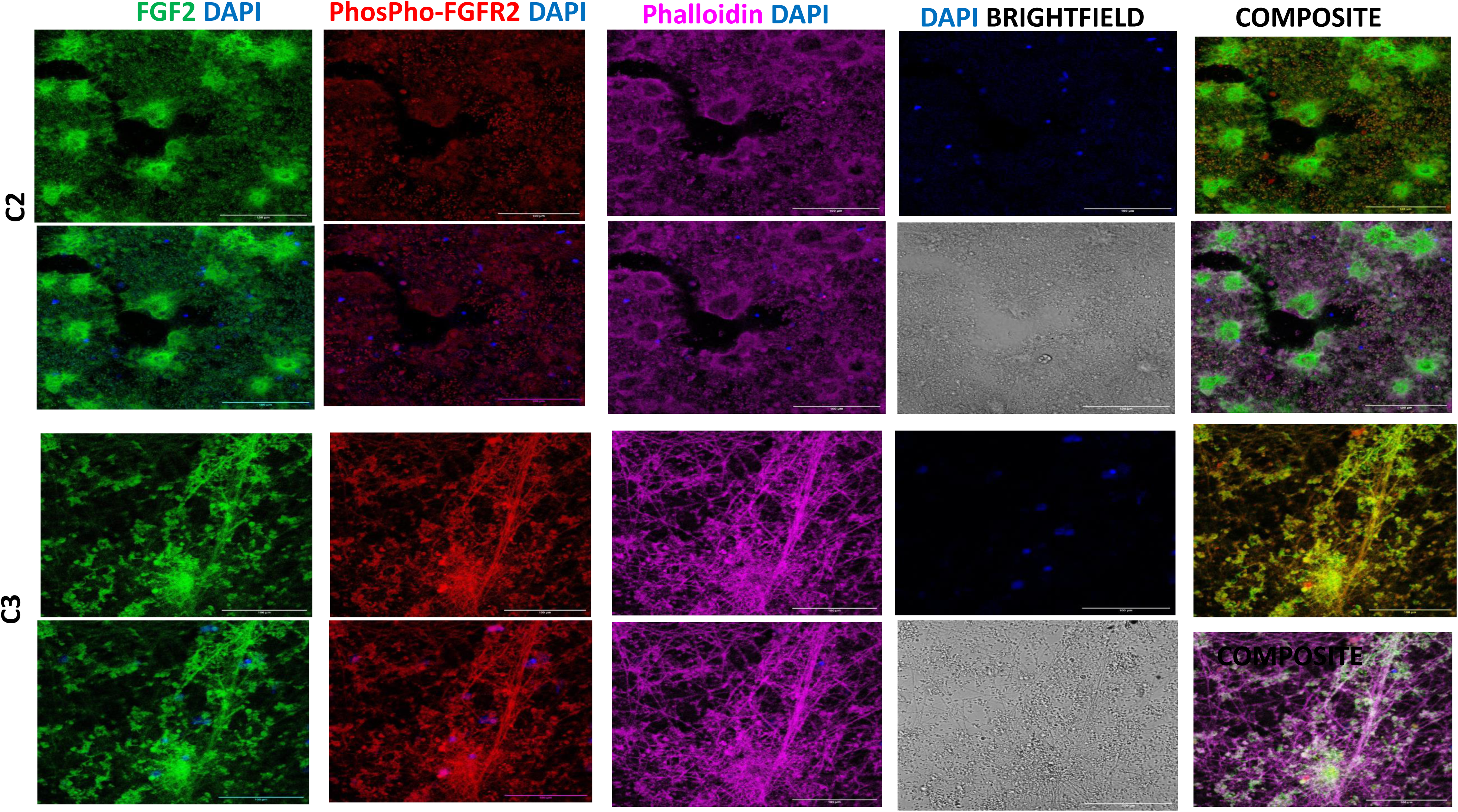
Active FGF Signalling: The figure depicts the expression FGF2 (green),involved in angiogenesis, Phospho-FGFR2 (red) is the active form of FGF2 receptor and Phalloidin (pink) used to stained actin filaments across different conditions at day 15. The FGF2 signalling is more active in C3 condition. Scale bars-100 µm

**Figure 28.**
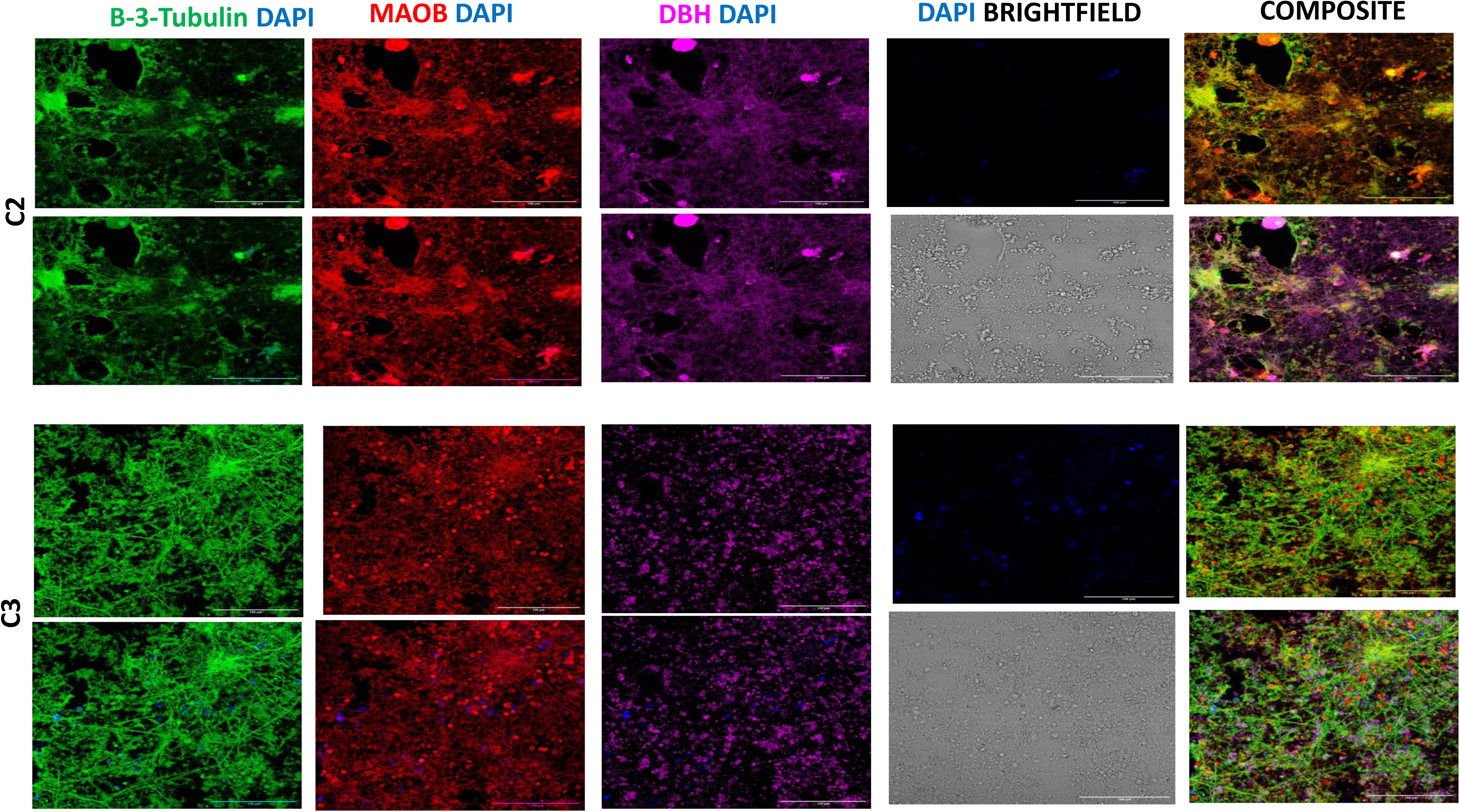
The figure depicts the expression Beta3 Tubulin (green), a marker of early formed neurons, MAOB (red) astrocytic activation marker and DBH (pink) a marker of sympathetic (norepinephrine) neurons across different conditions at day 15. The FGF2 signalling is more active in C3 condition. Scale bars-100 µm

**Figure 29.**
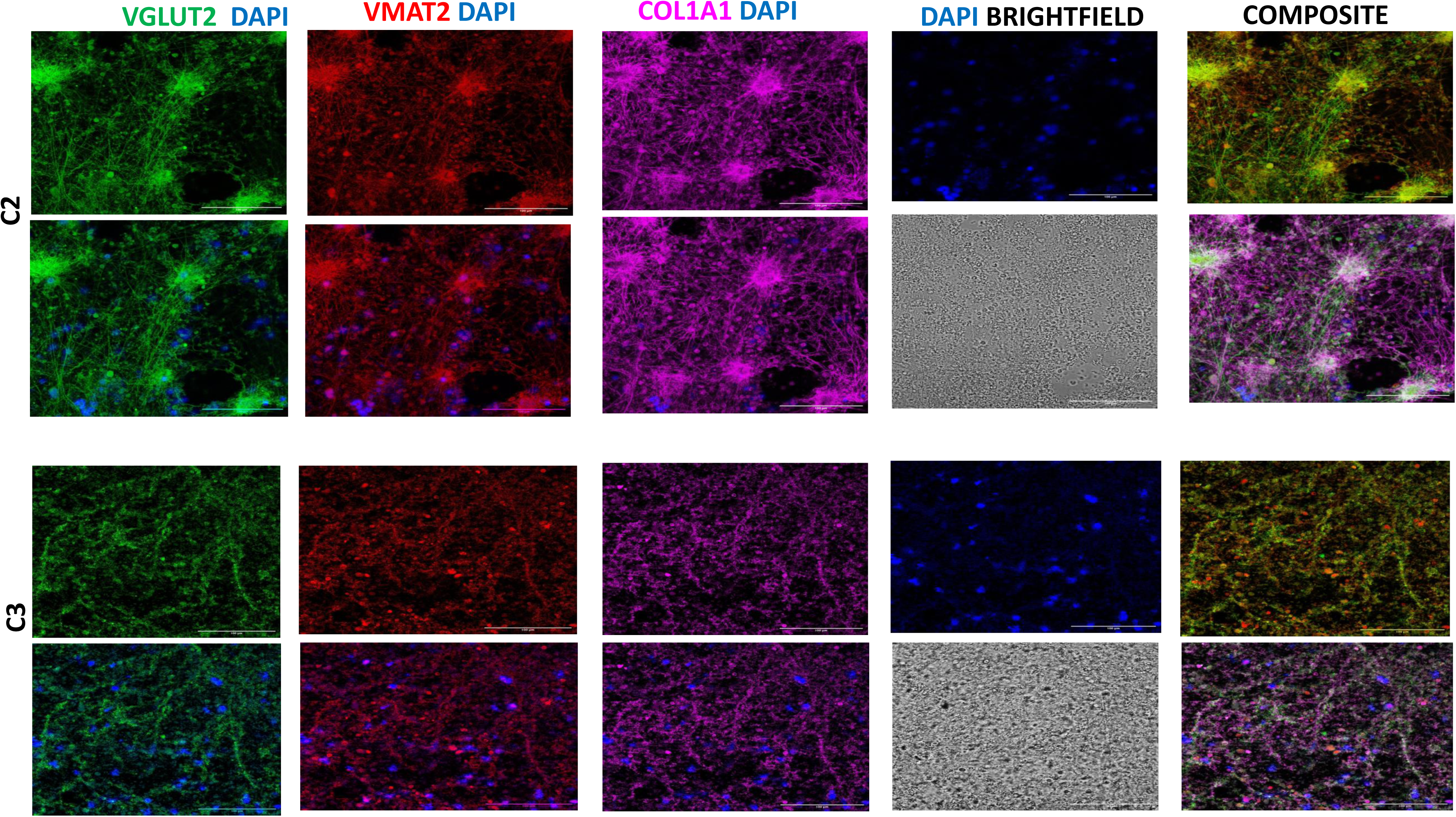
The figure depicts the expression VGLUT2 (green), a marker of glutamatergic neurons, VMAT 2 (red) plays a crucial role in the transport of monoamines (such as dopamine, serotonin, and norepinephrine) to the into synaptic vesicles for release during neurotransmission and Col1A1 (pink) a collagen marker to check the structural integrity of developing vasculature in the culture across different conditions at day 15. The positive signal of VMAT2 in the culture indicates the presence of neuromodulatory neurons. Scale bars-100 µm

**Figure 30.**
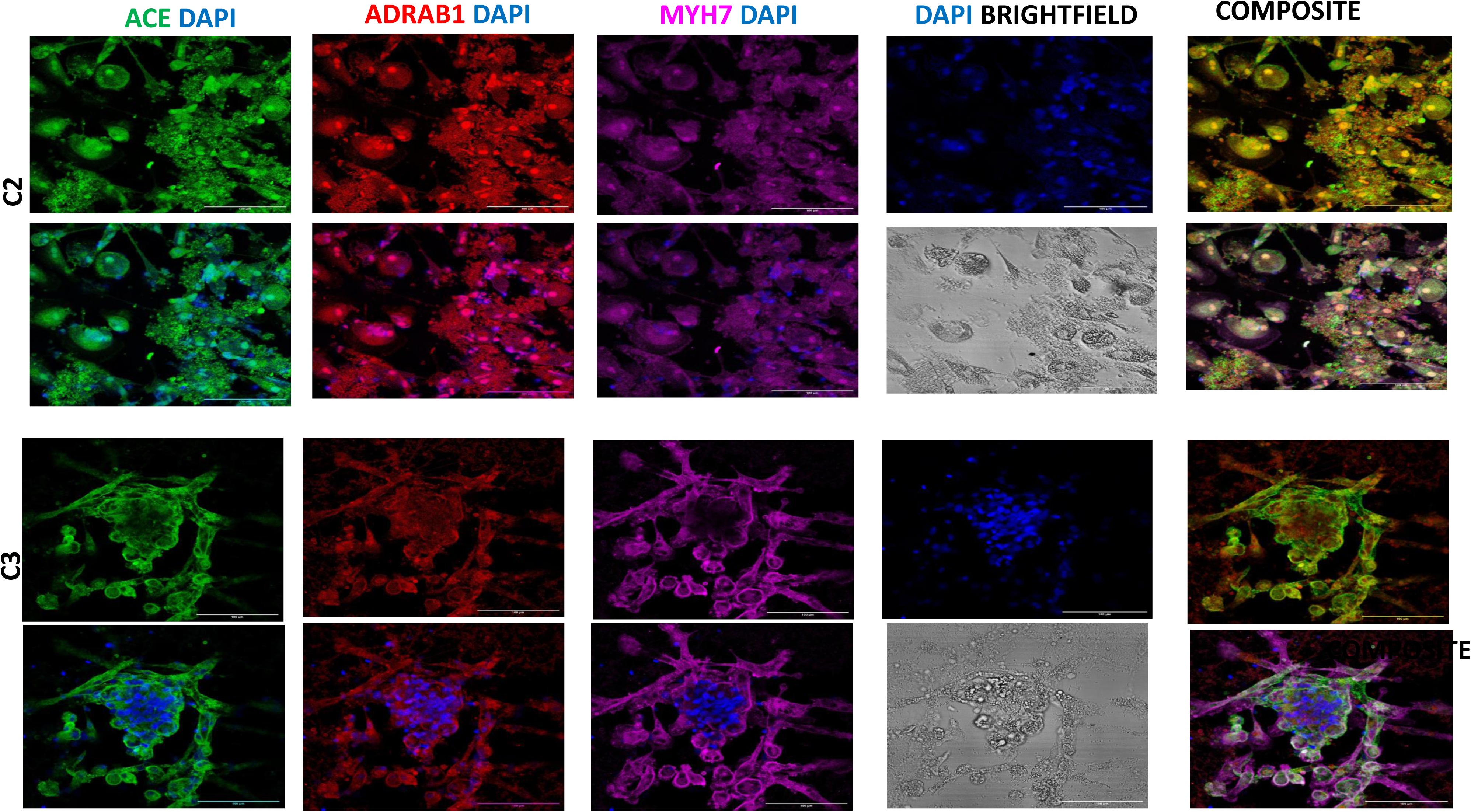
The figure depicts the expression ACE (green), is involved in the local production of angiotensin II within the brain, which can affect cerebral blood flow and vascular reactivity, ADRAB1(red) plays a crucial role in the in mediating the effects of norepinephrine and epinephrine on neuronal excitability and synaptic transmission in cardiac tissue and MYH 7 (pink) a crucial component of cardiac muscles here it indicates mesodermal progenitors in the culture across different conditions at day 15. Scale bars-100 µm

**Figure 31-63.**
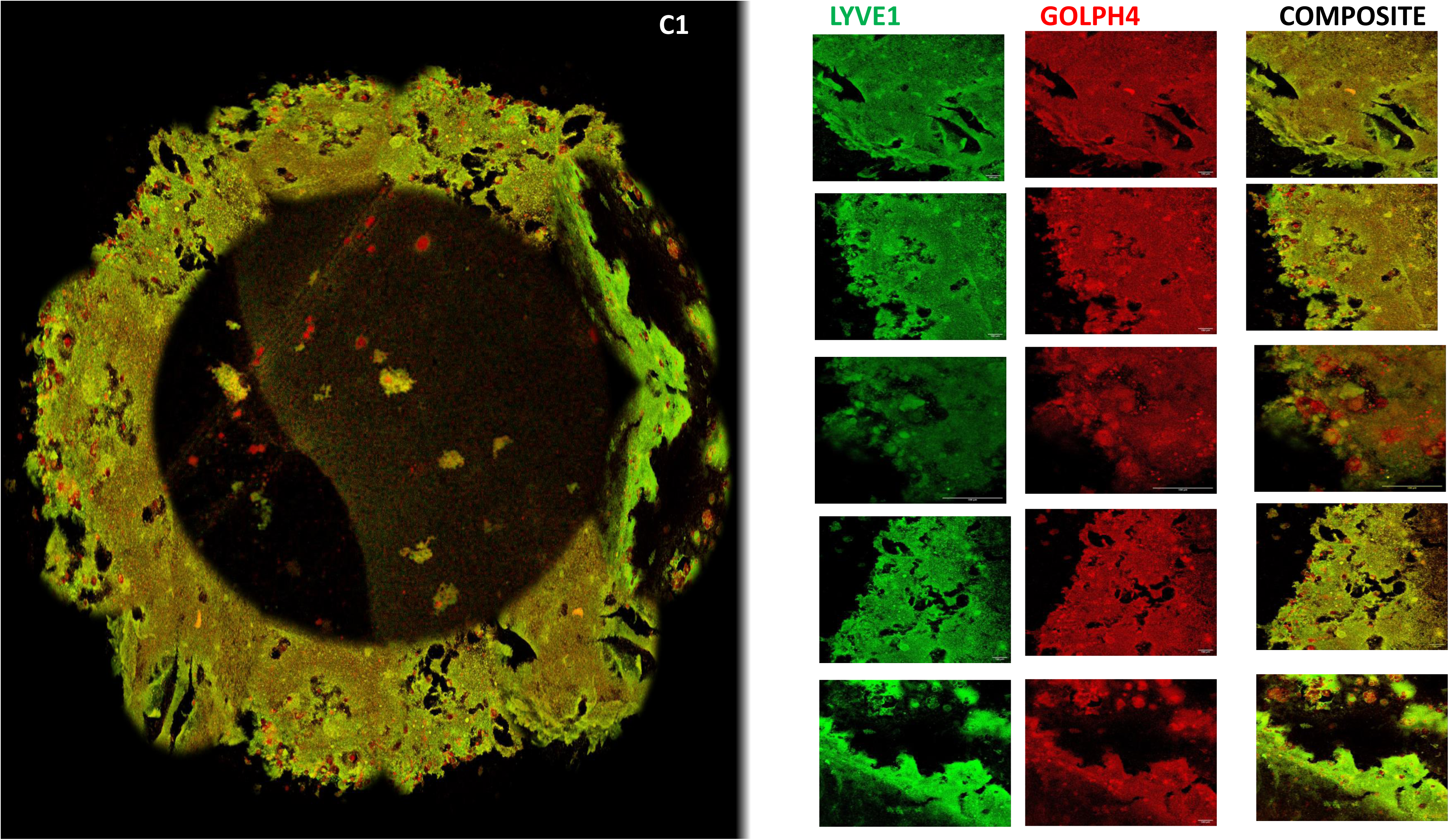
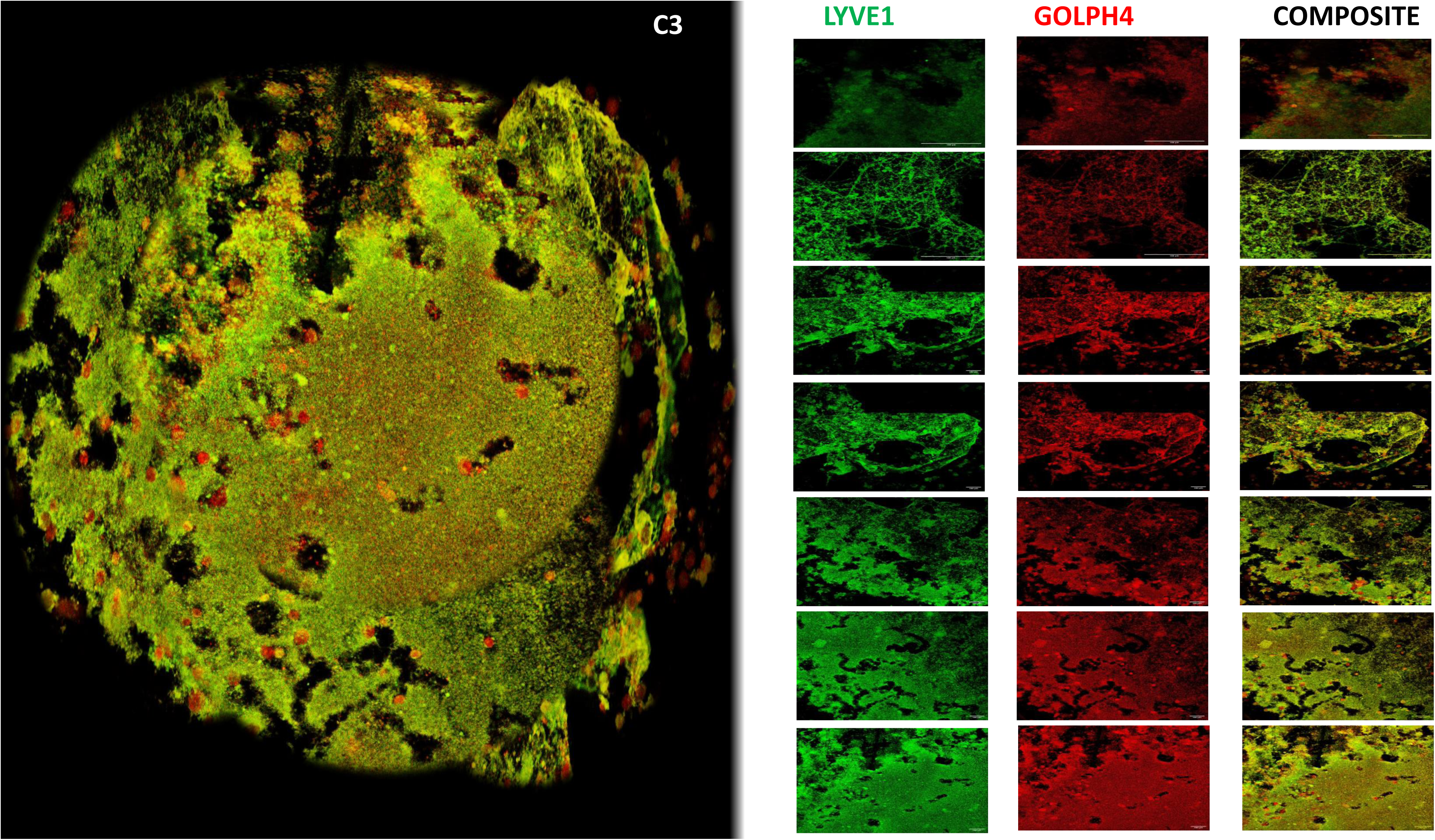
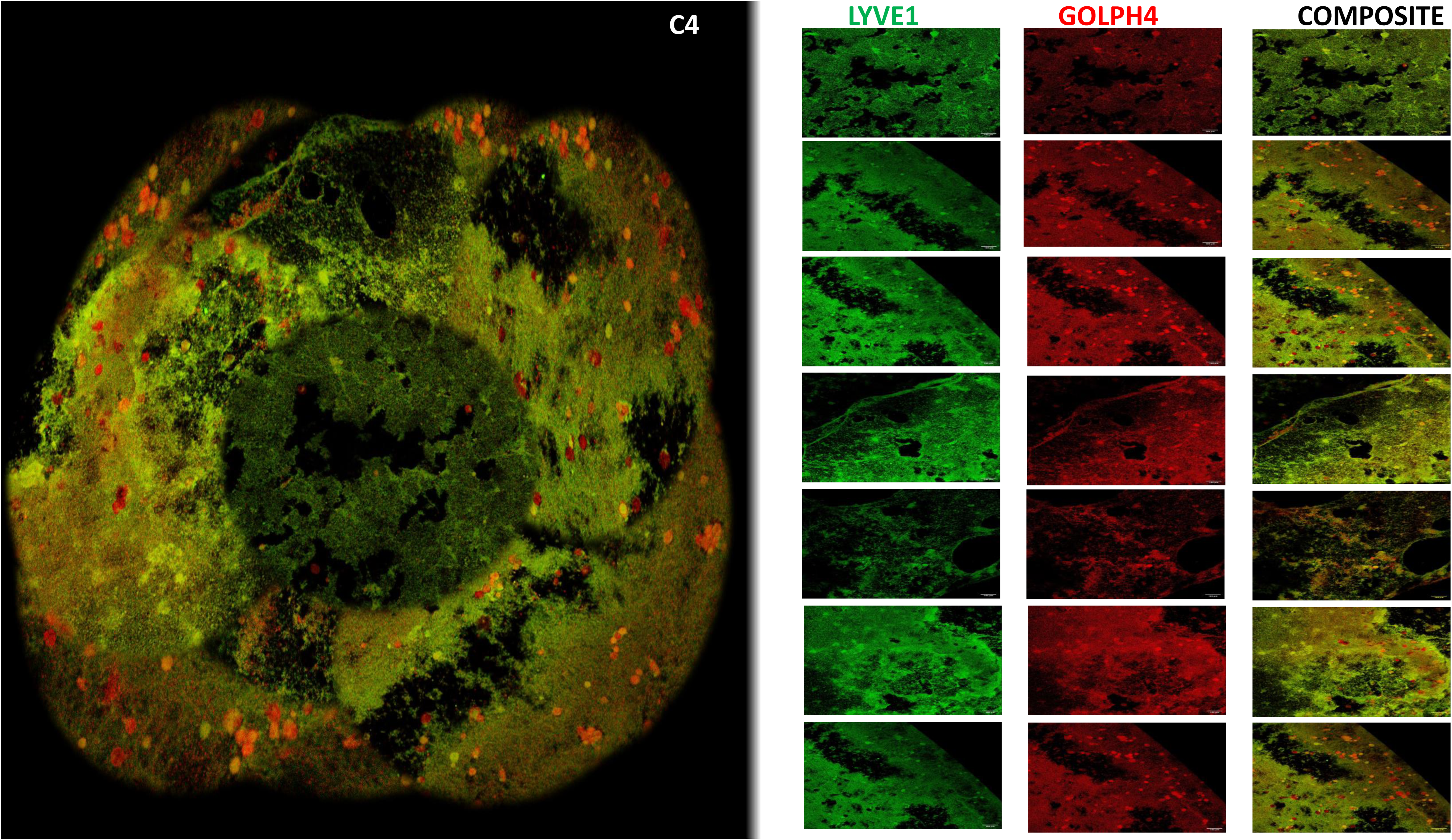
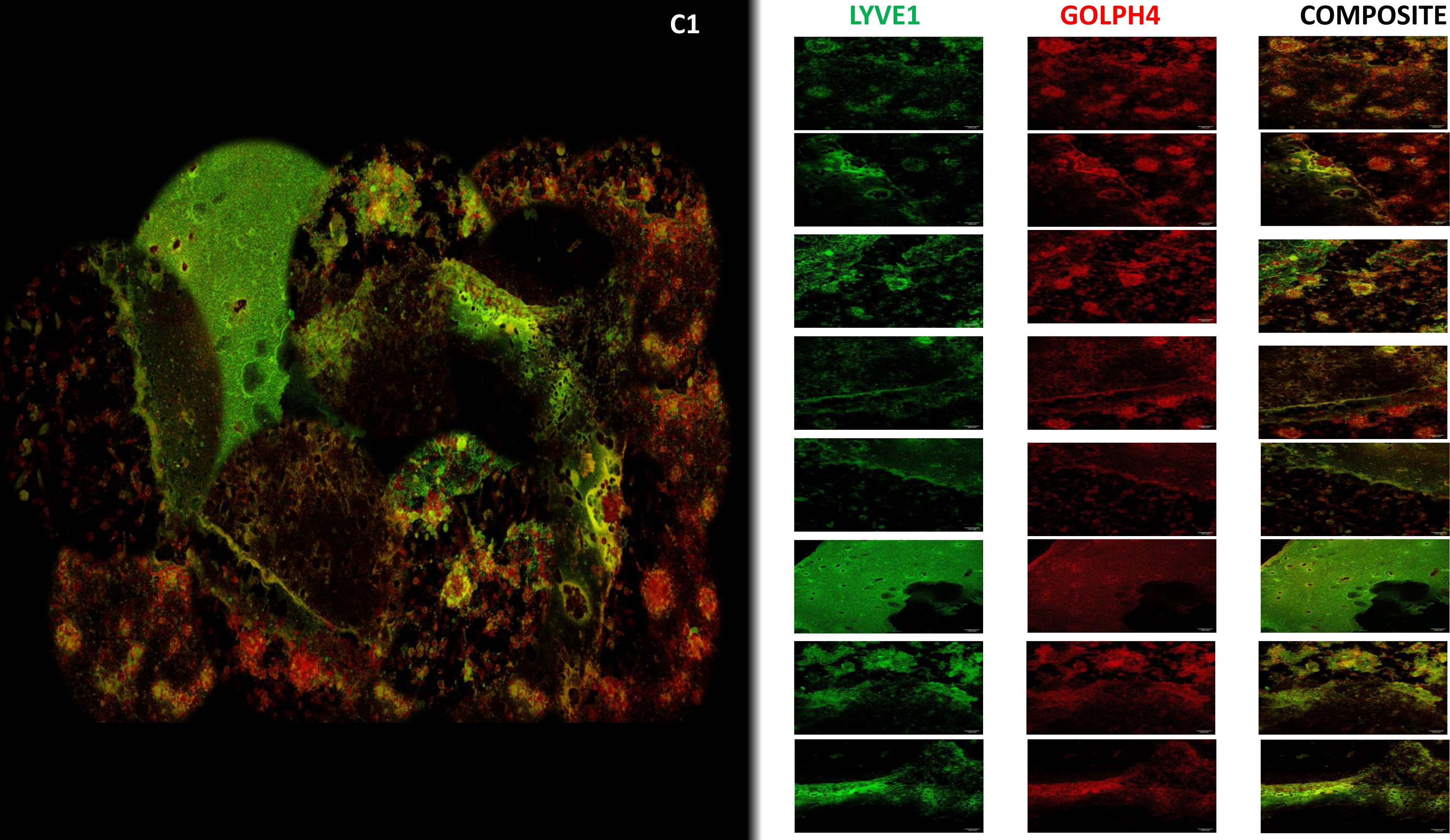
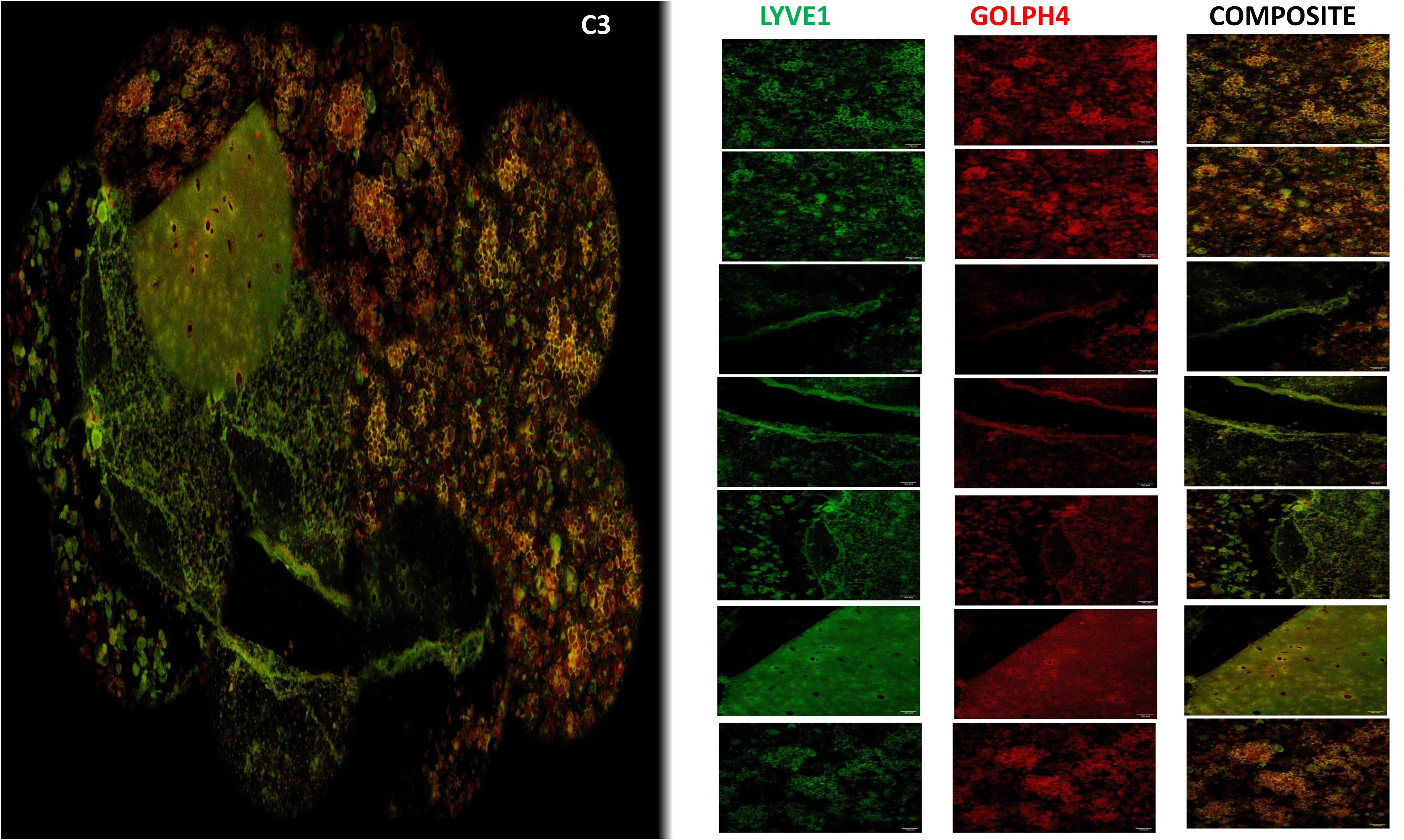
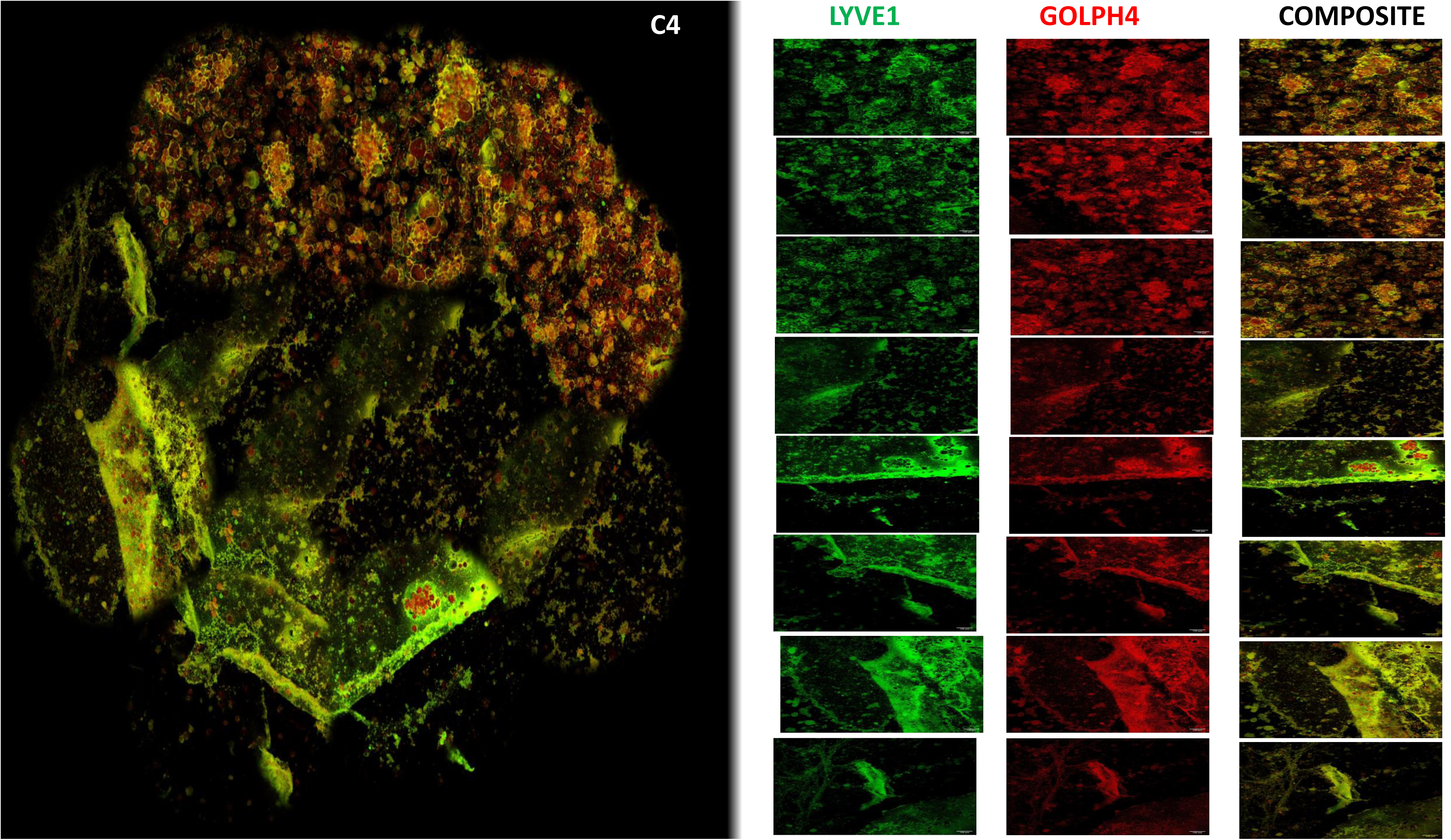
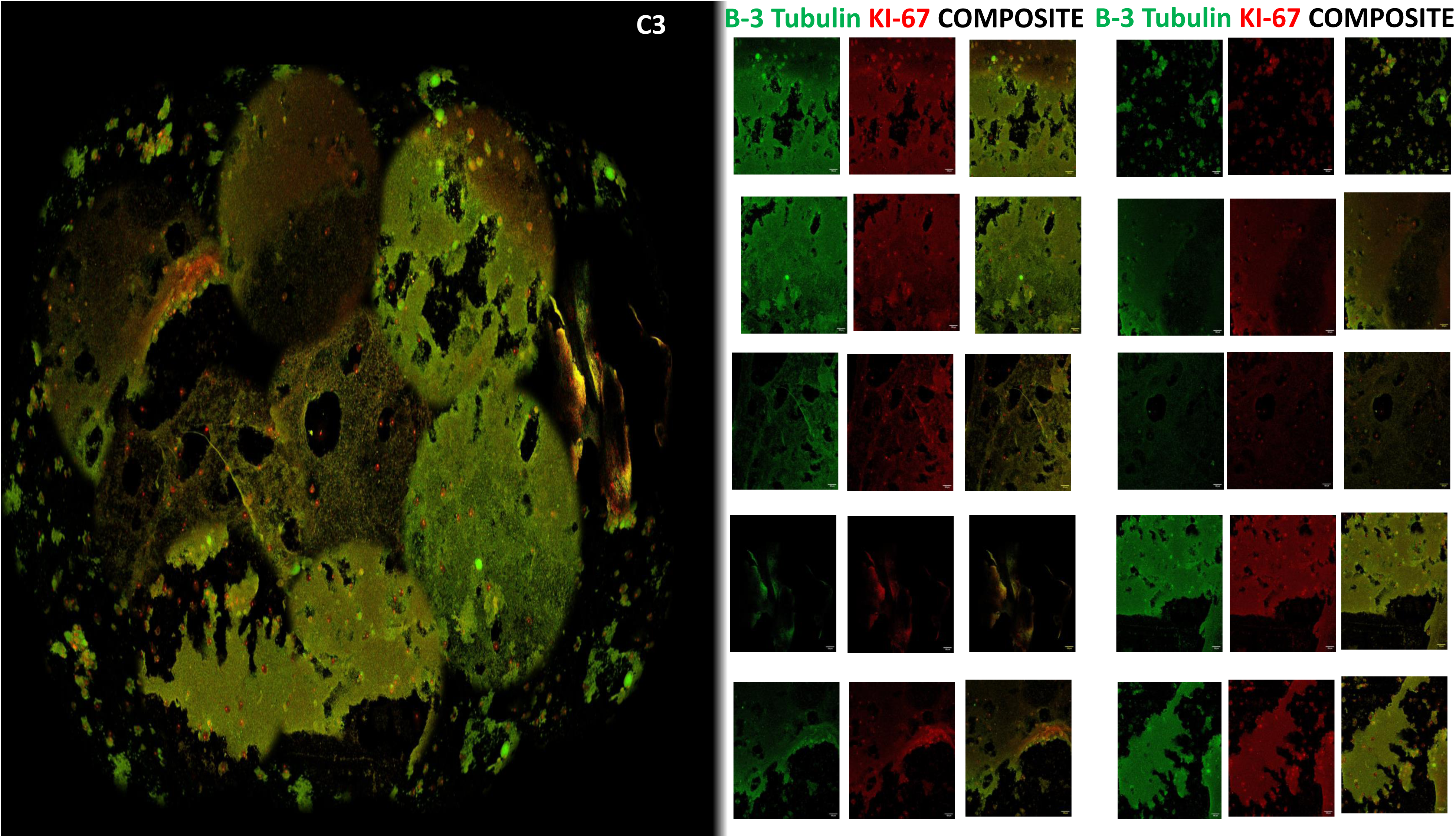
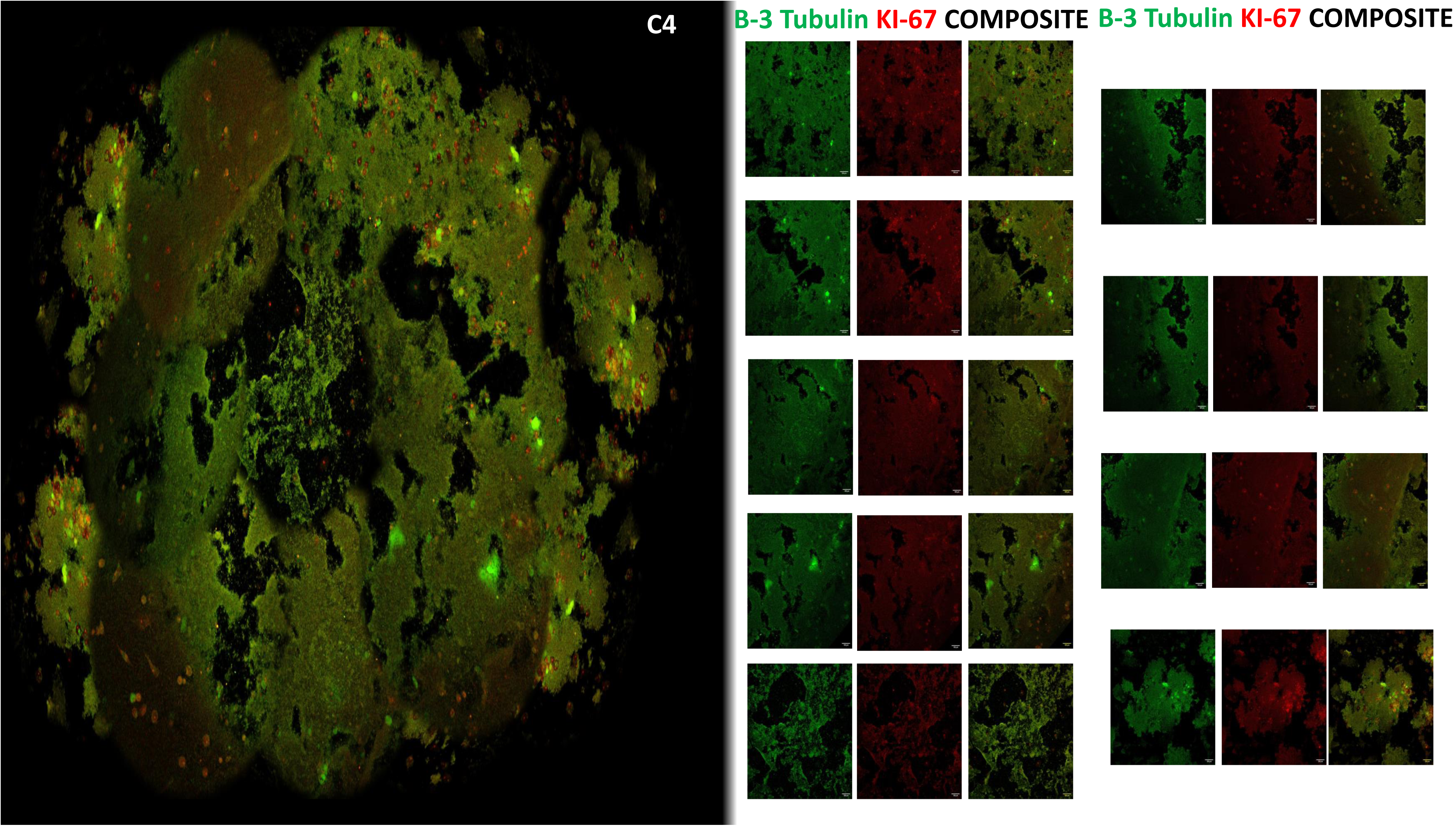
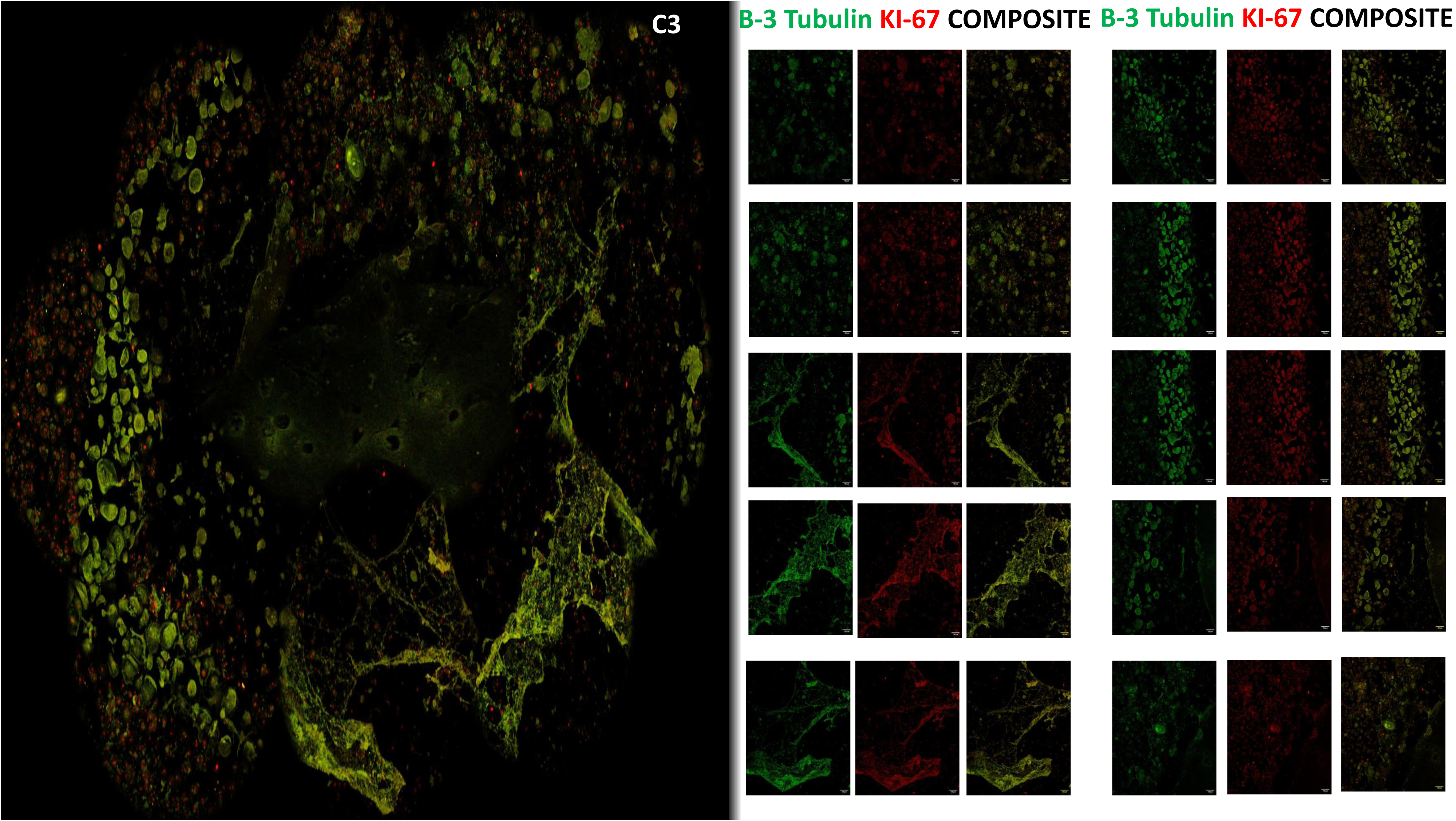
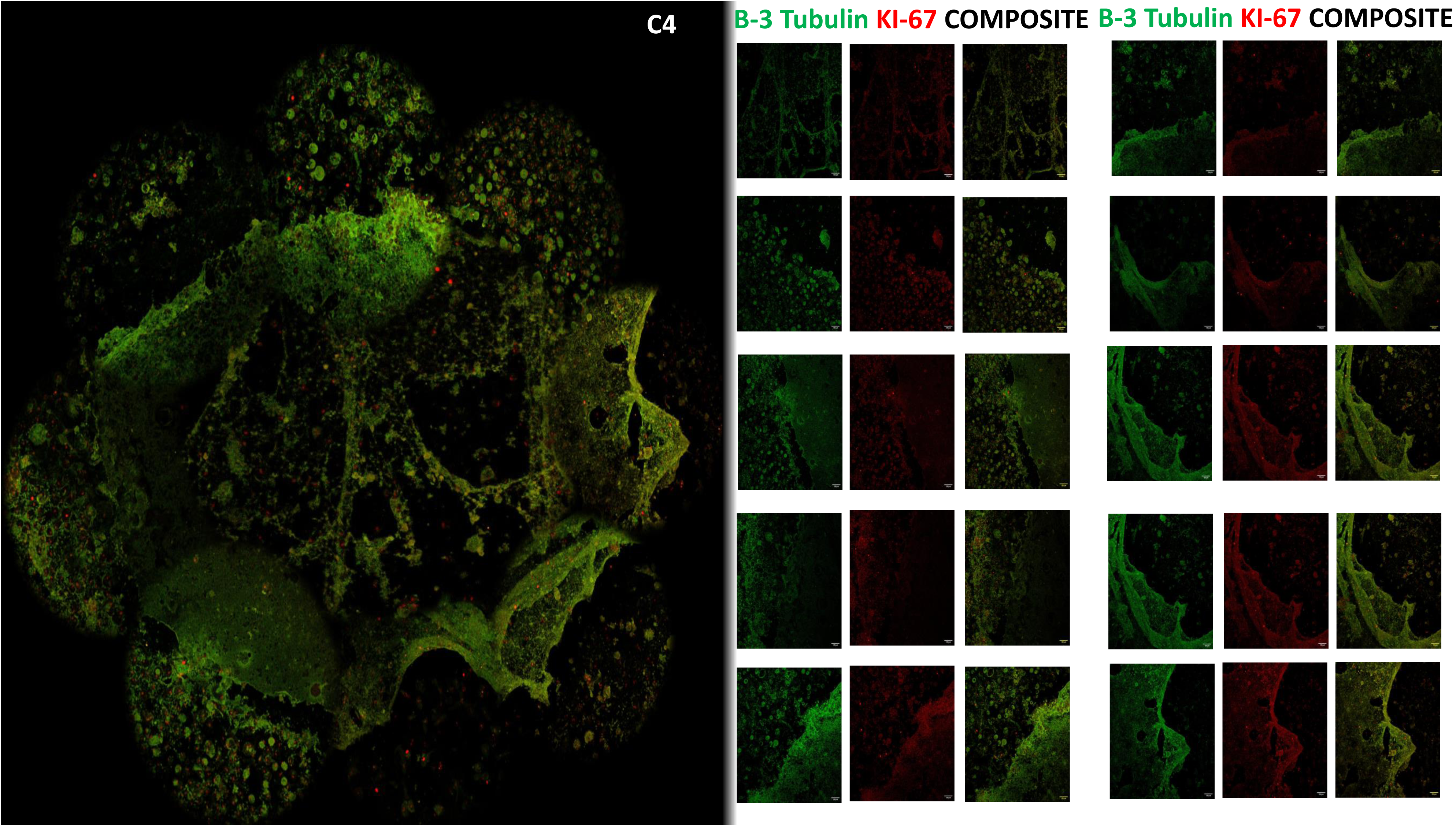
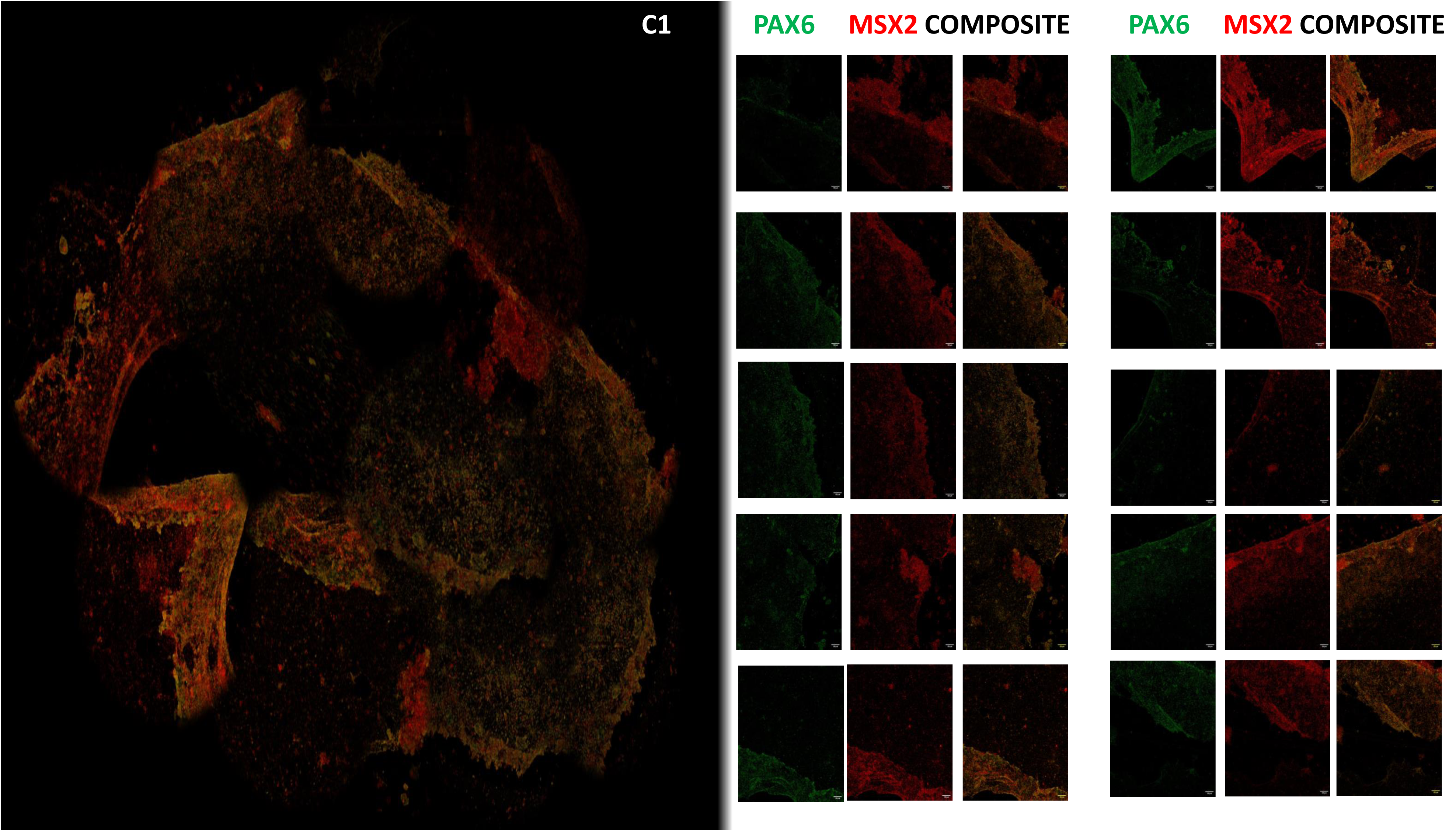
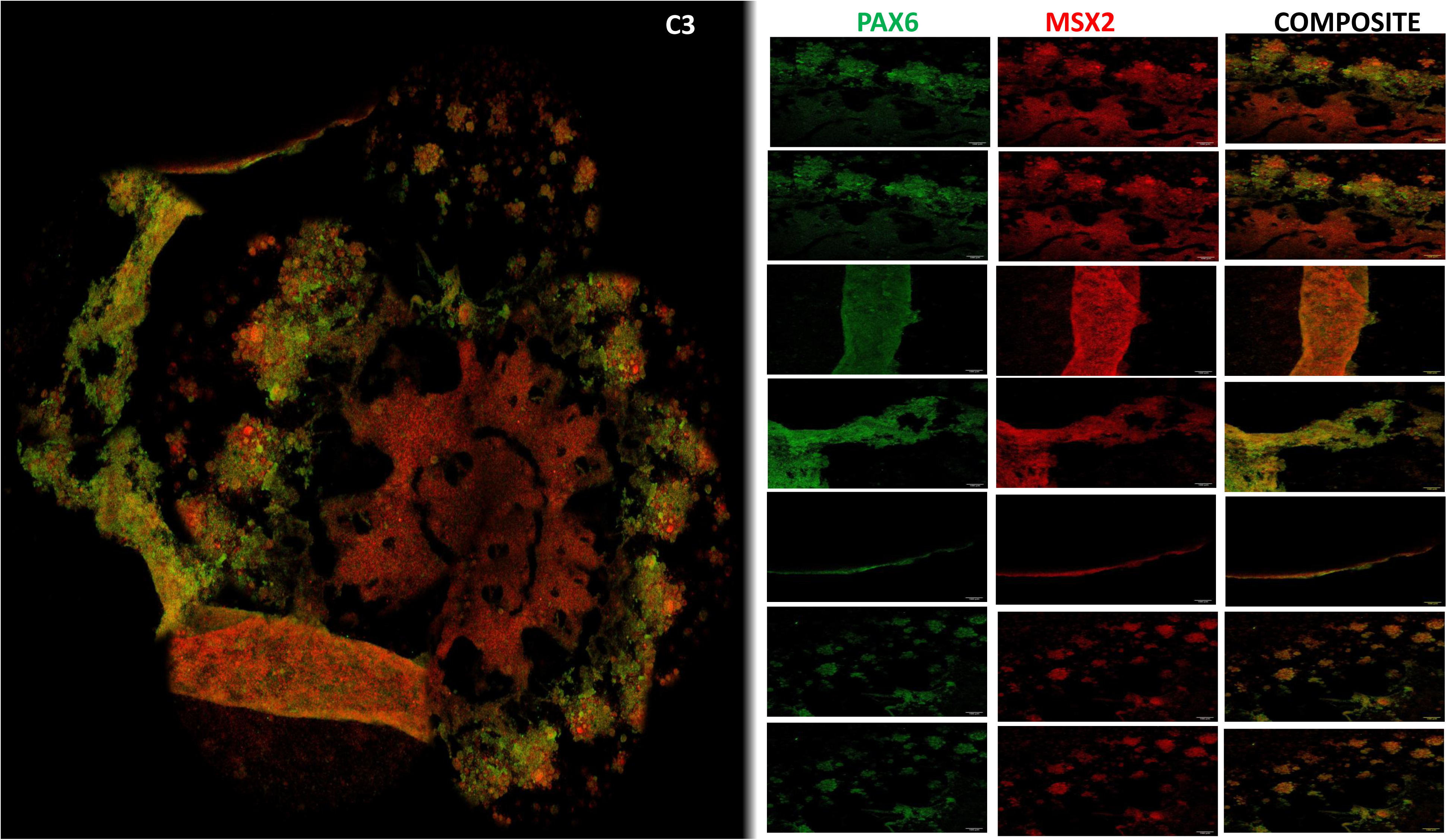
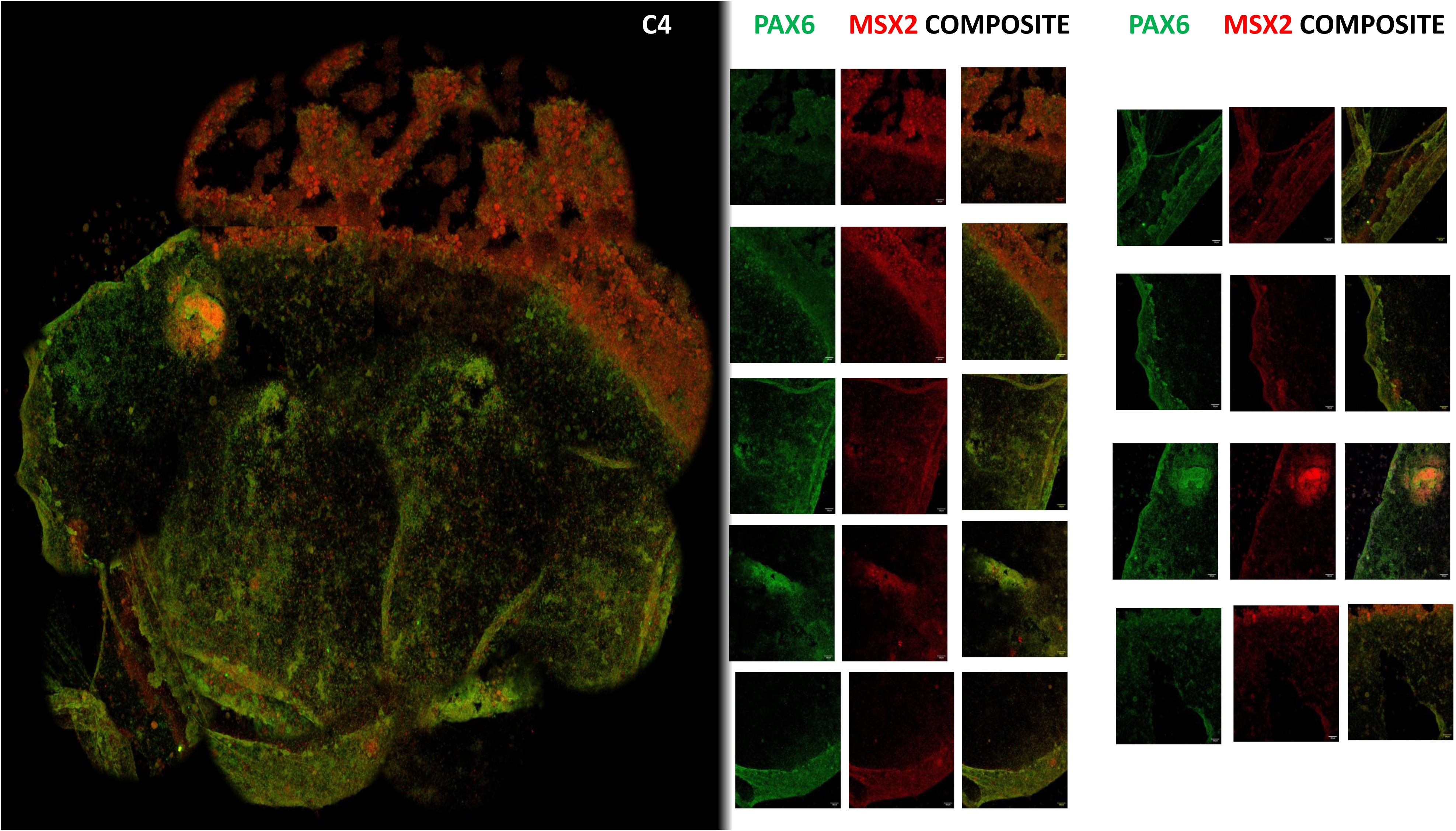
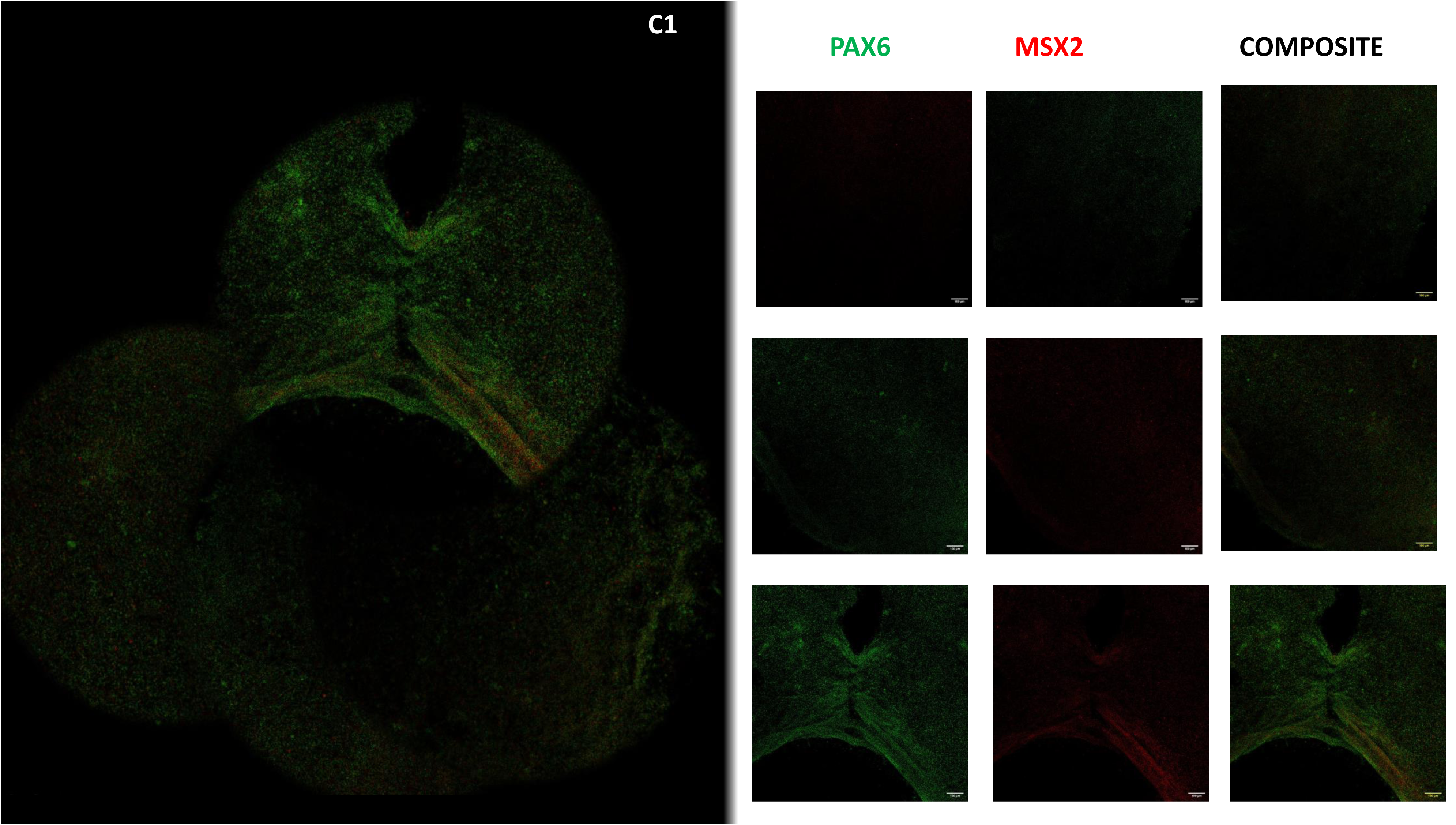
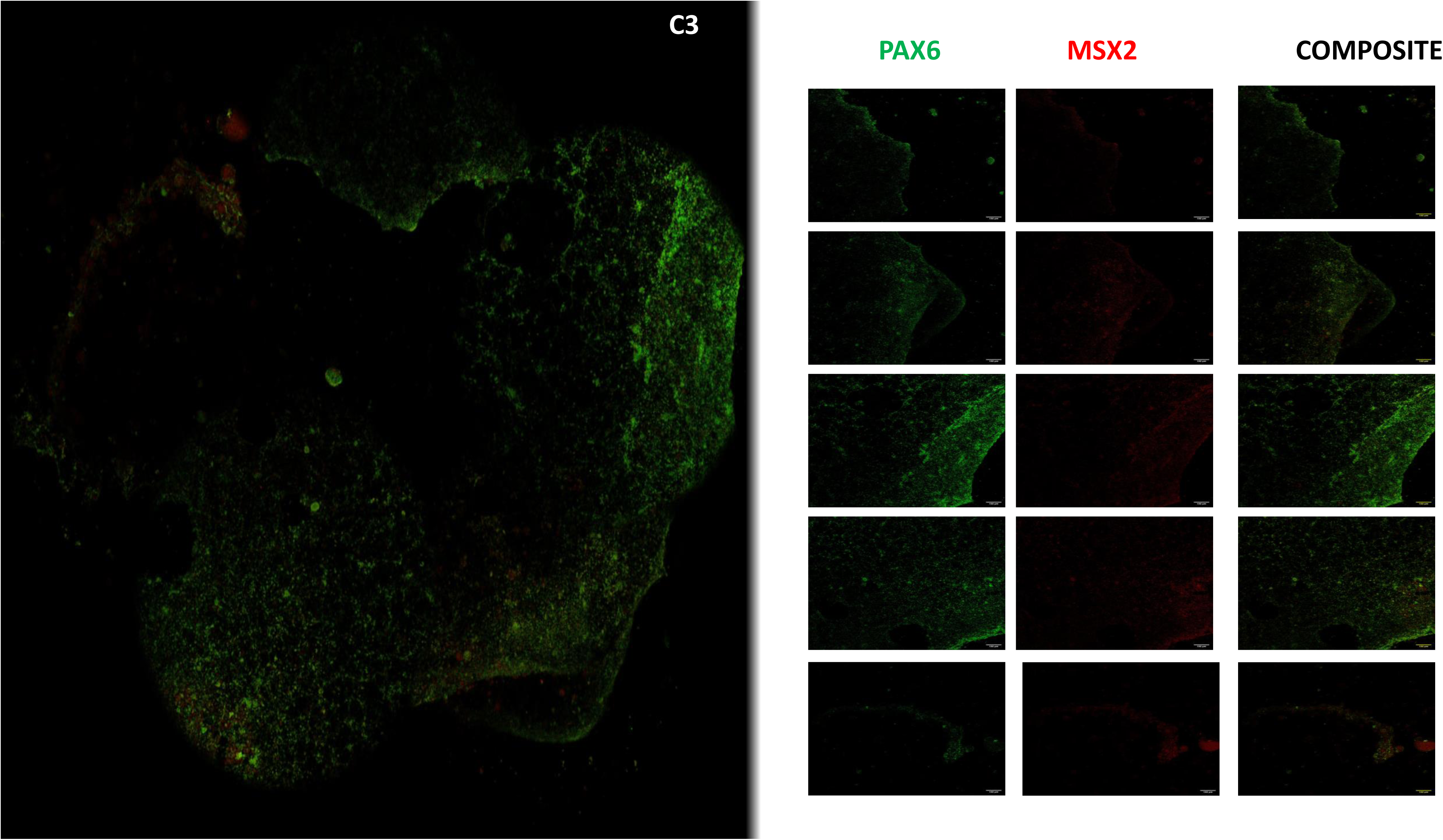
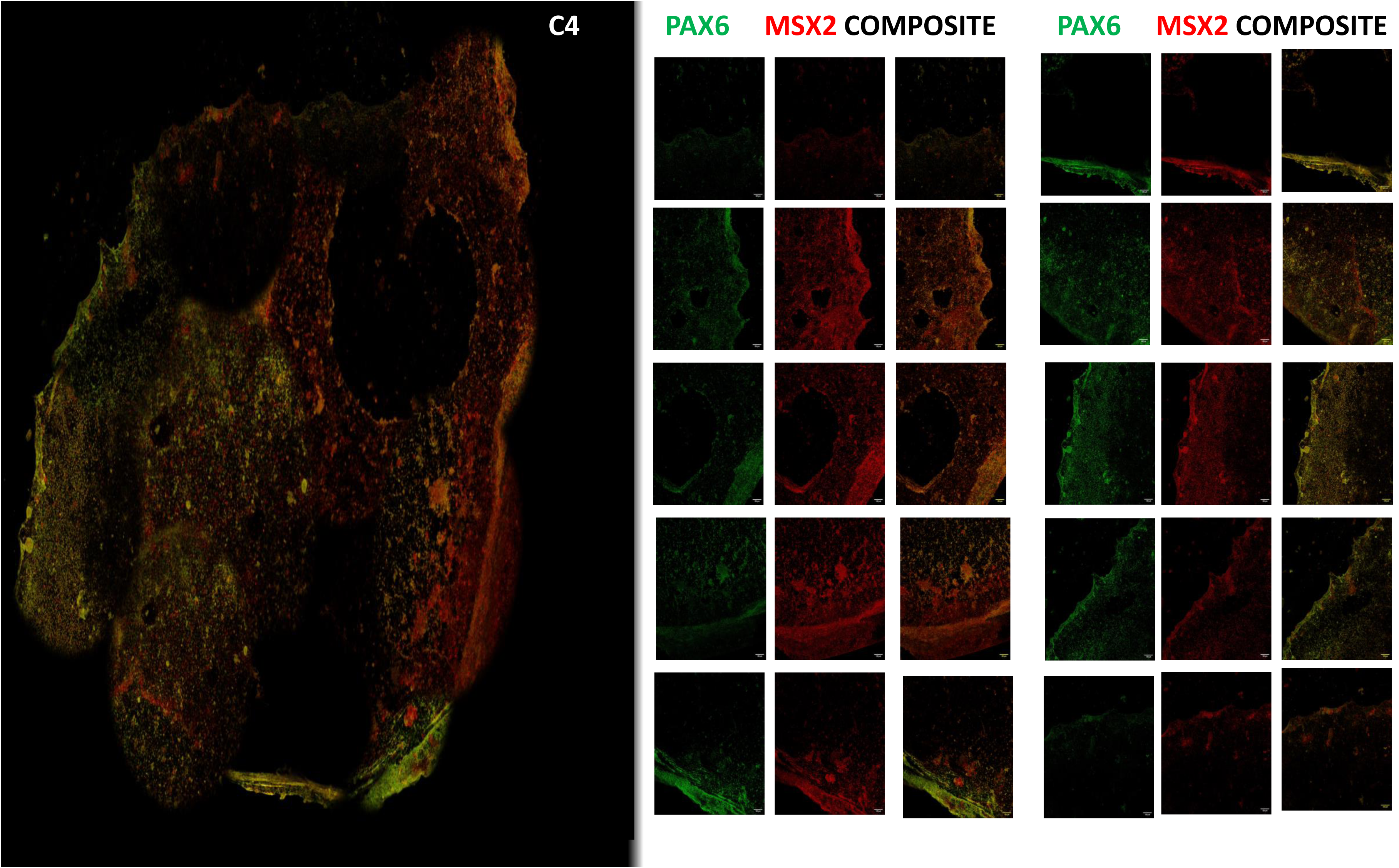
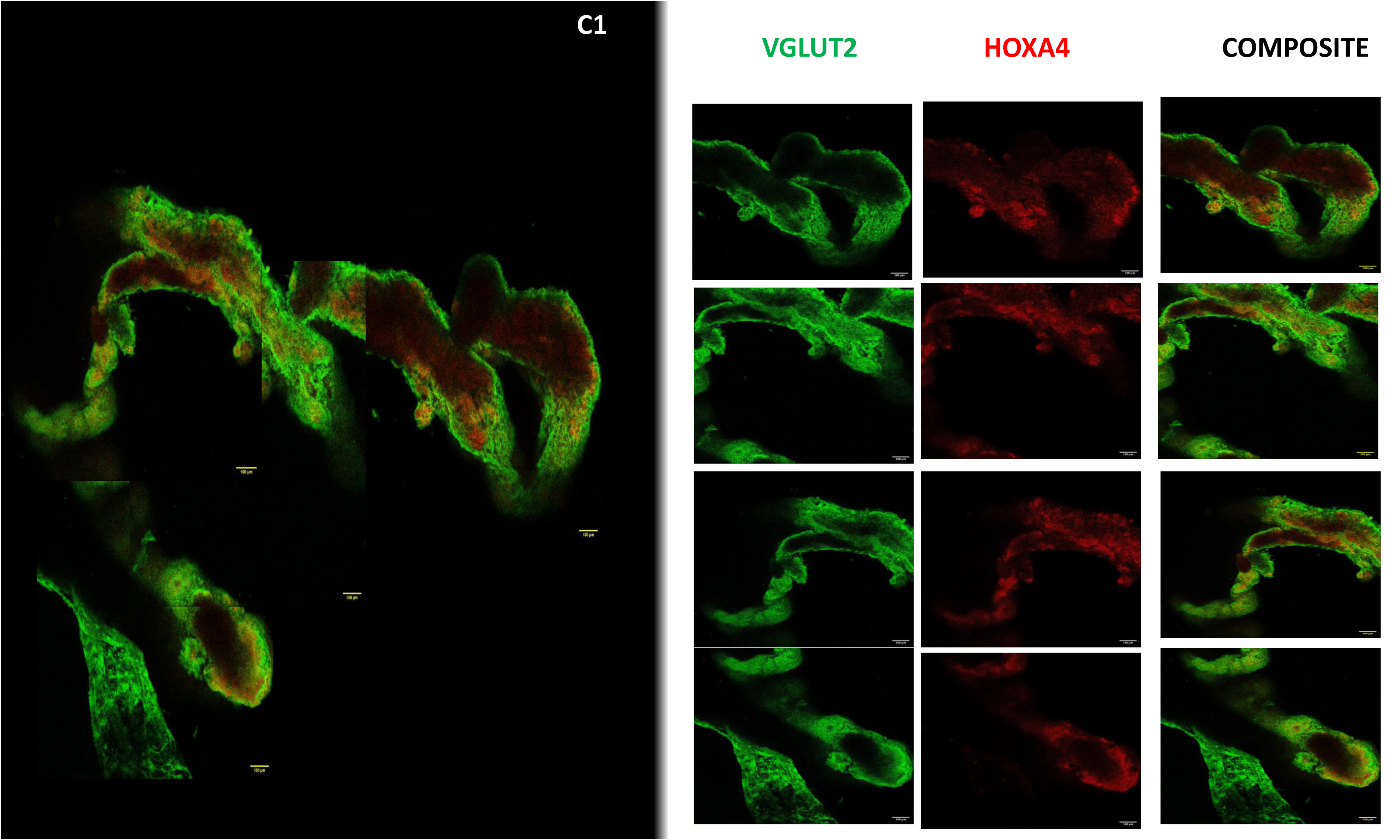
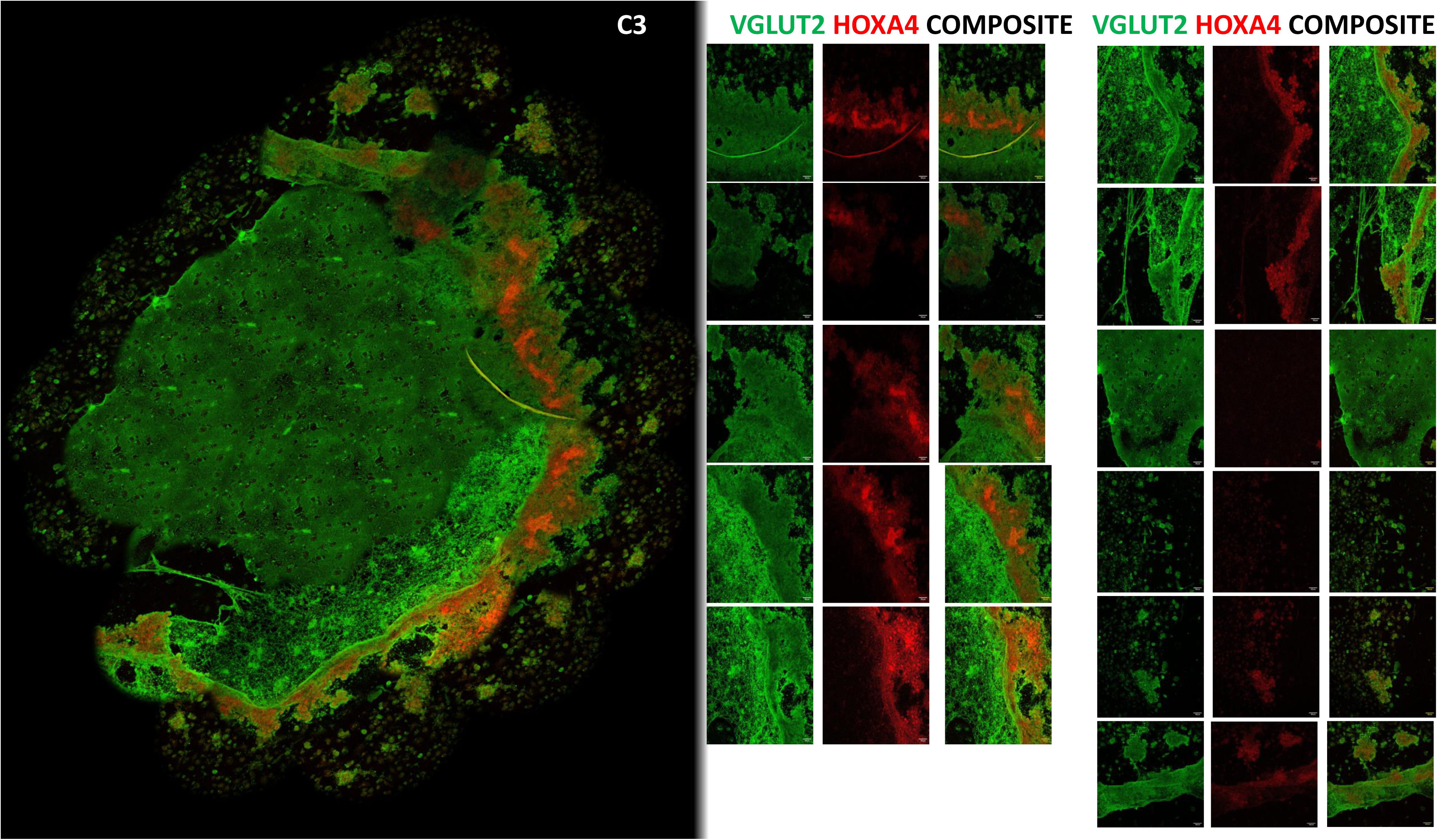
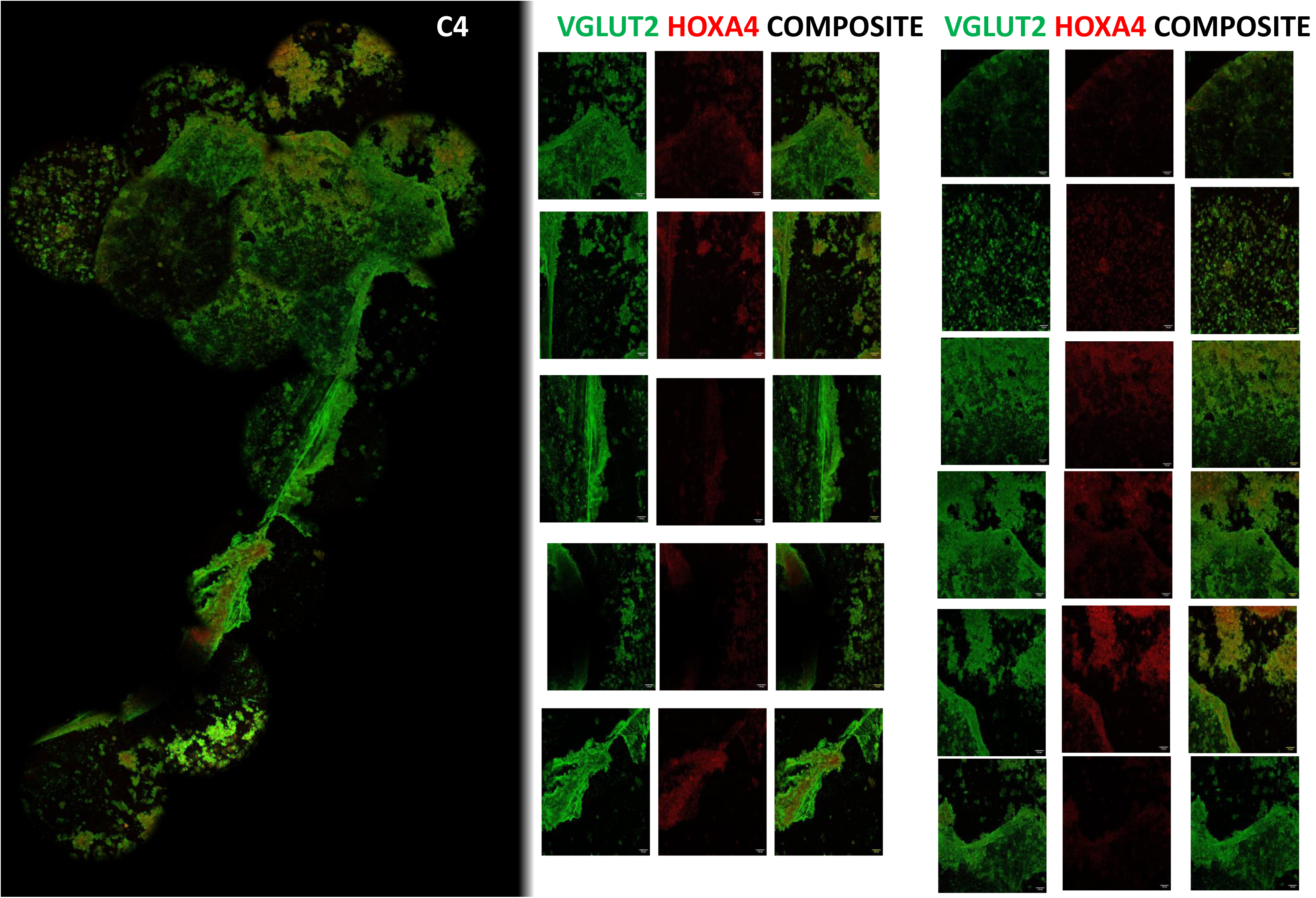
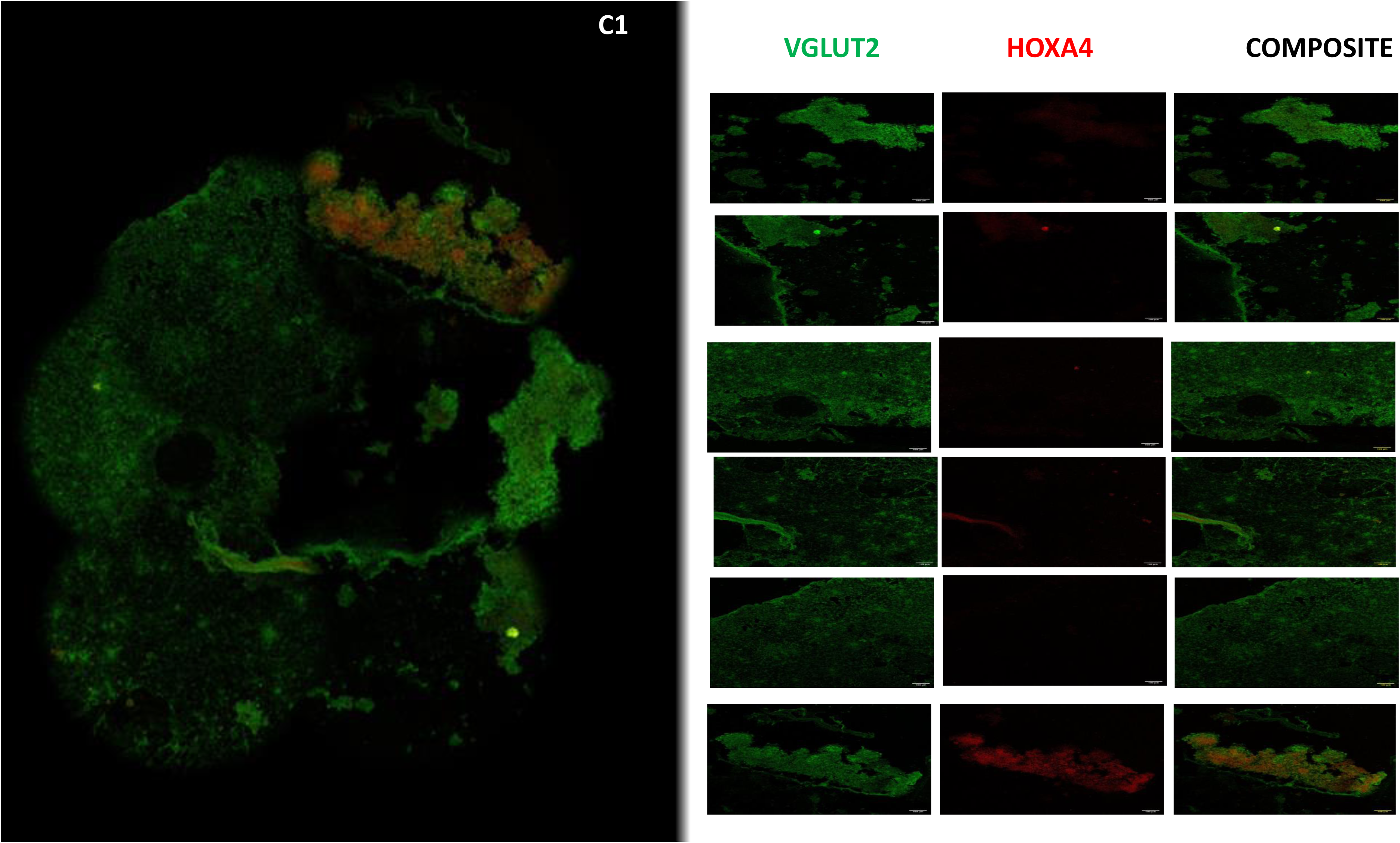
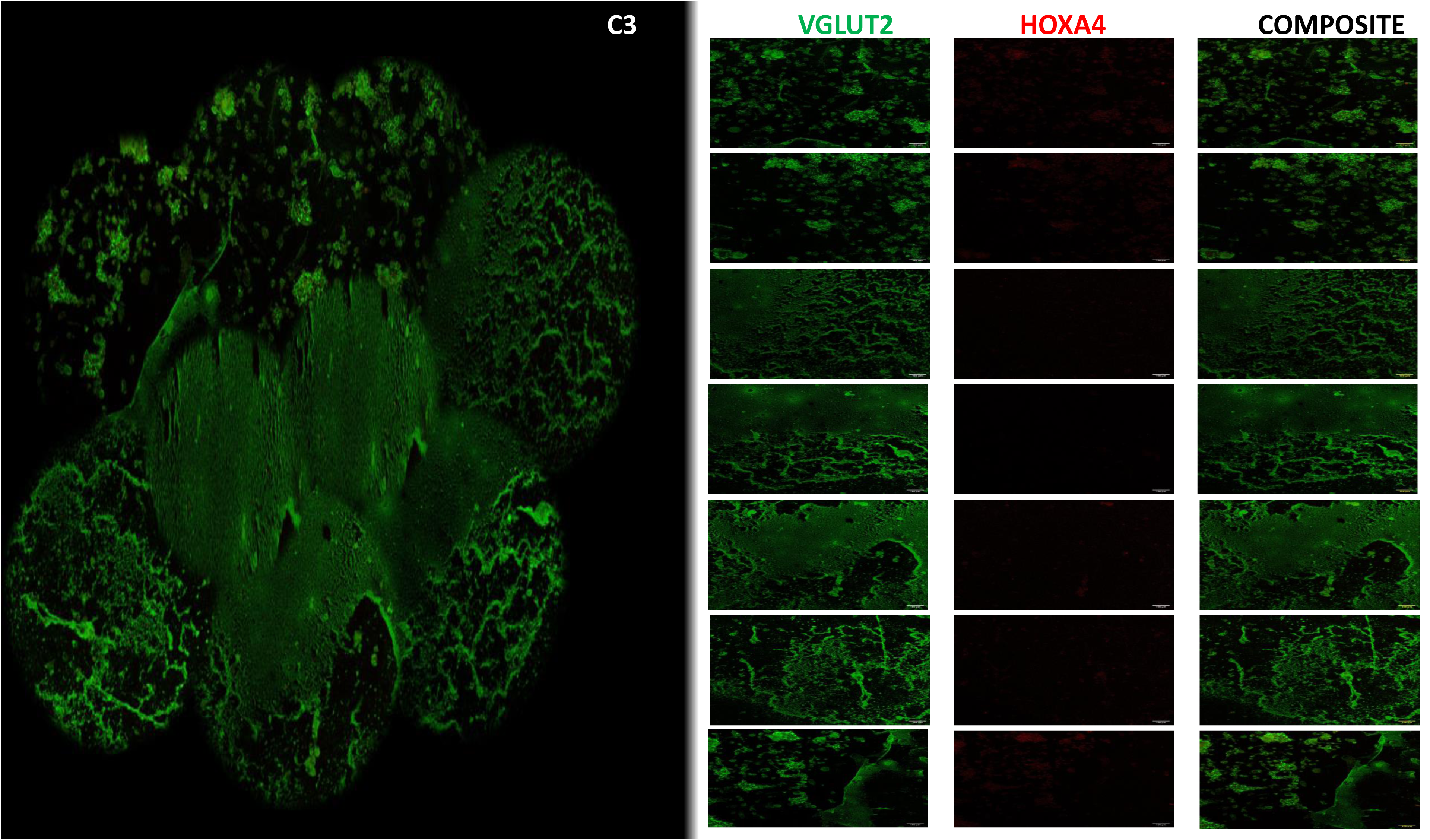
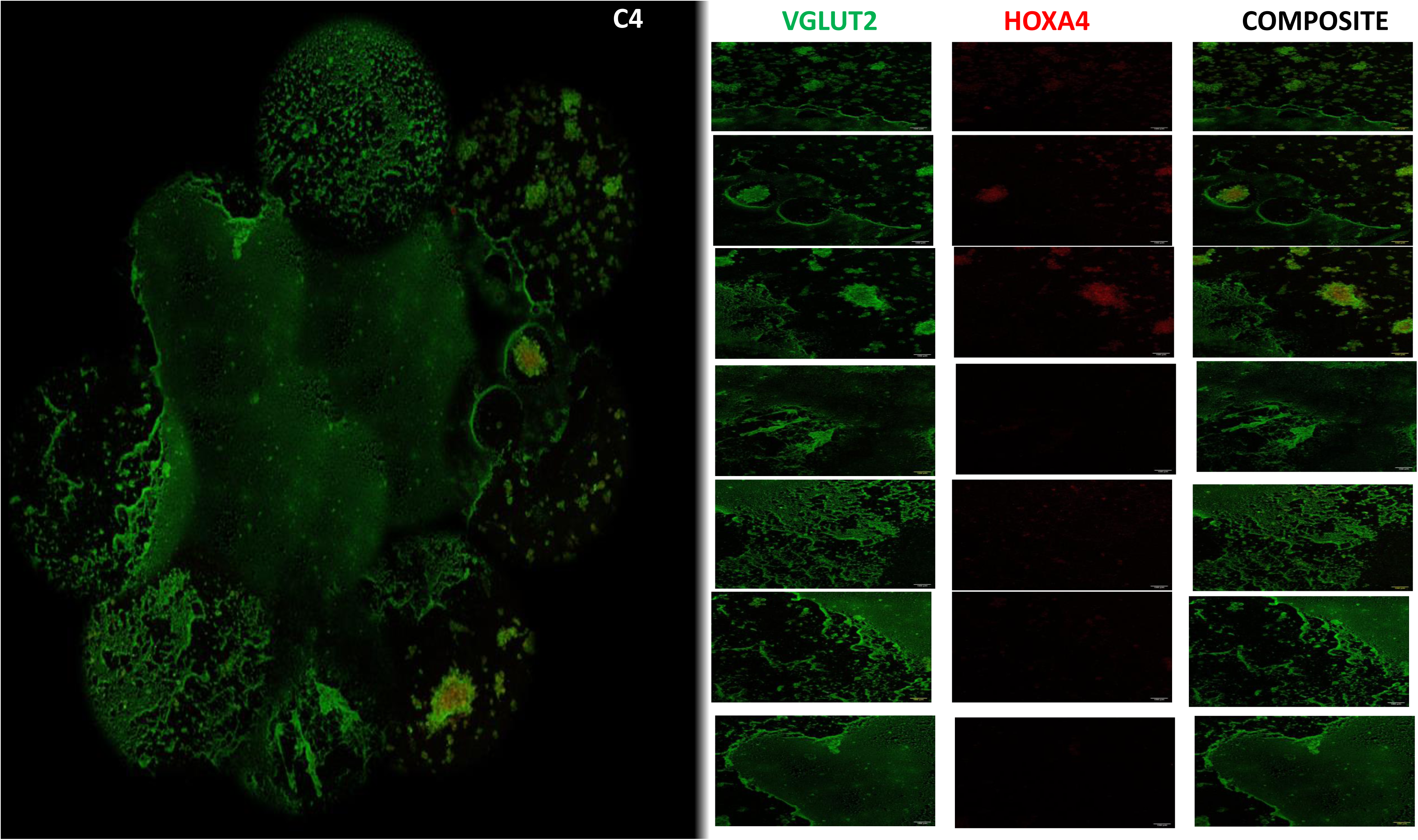
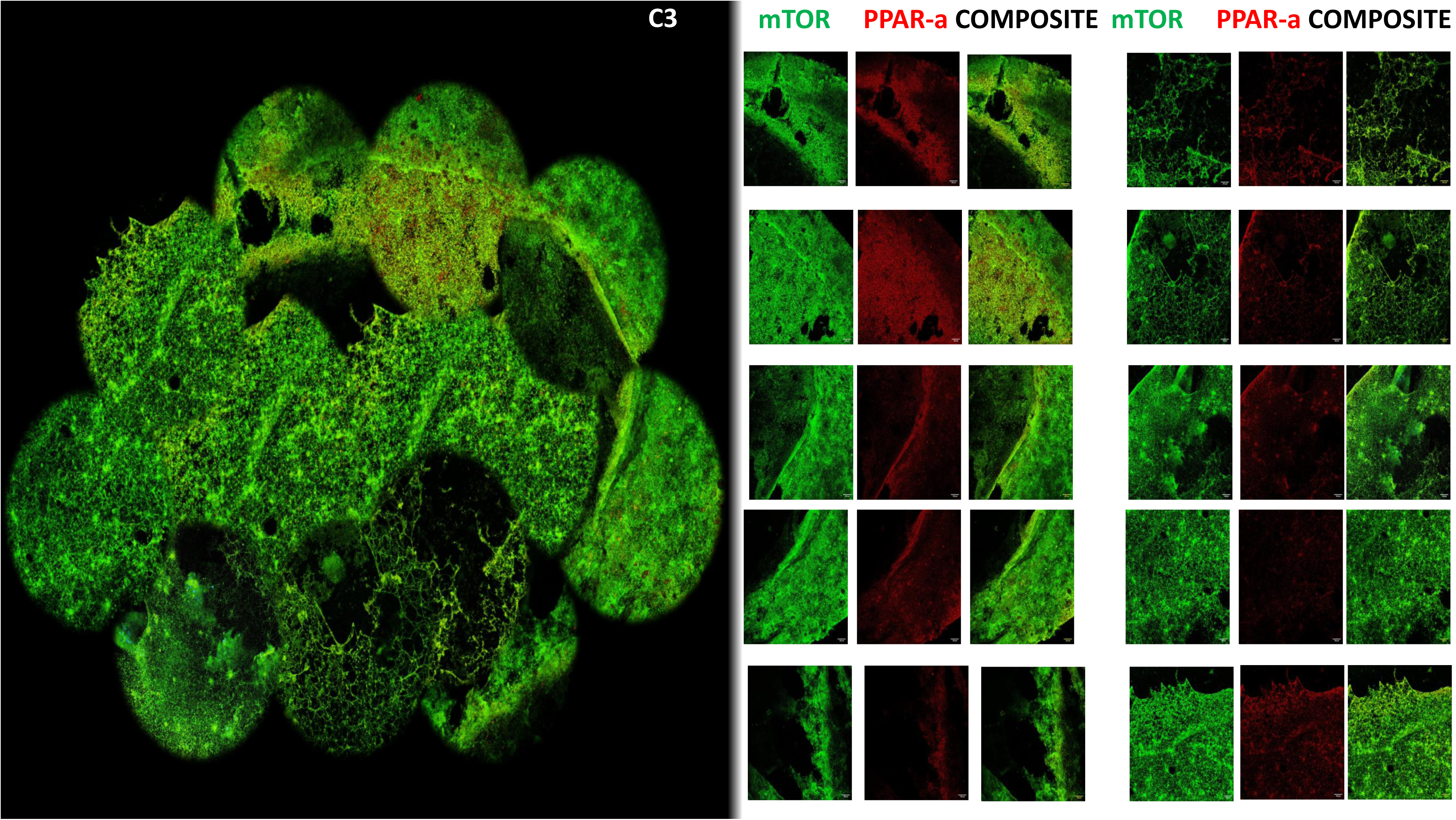
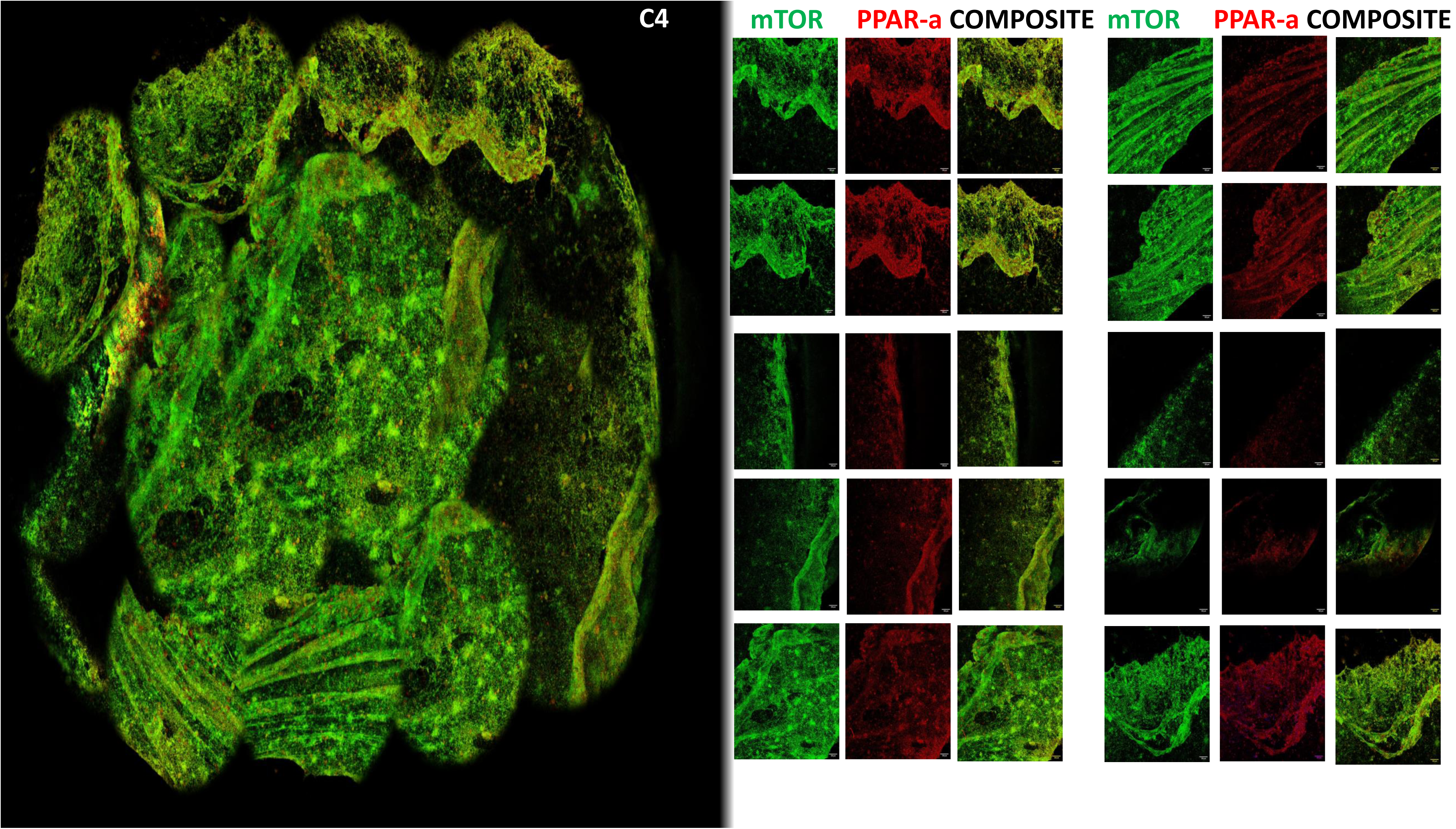
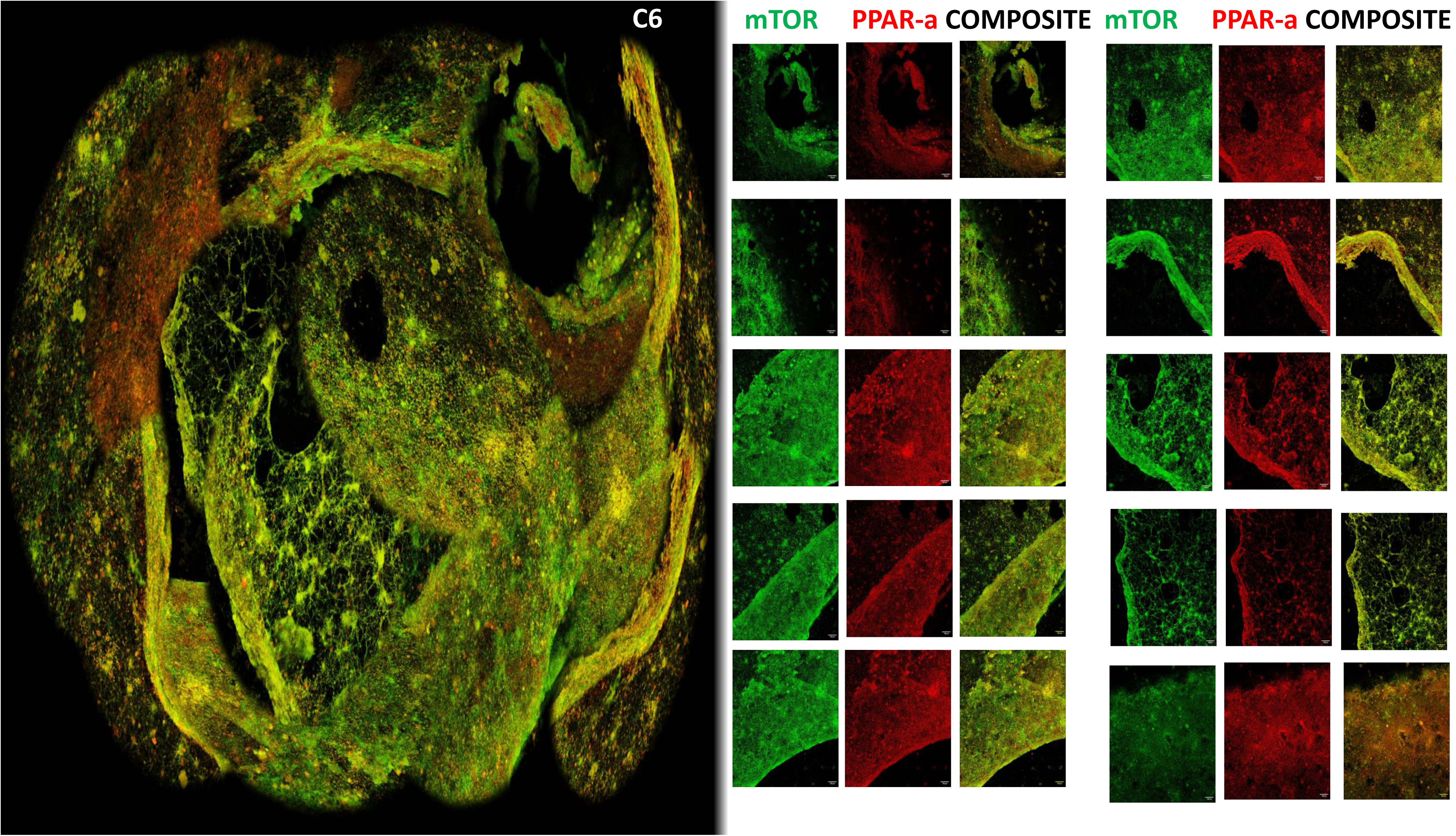
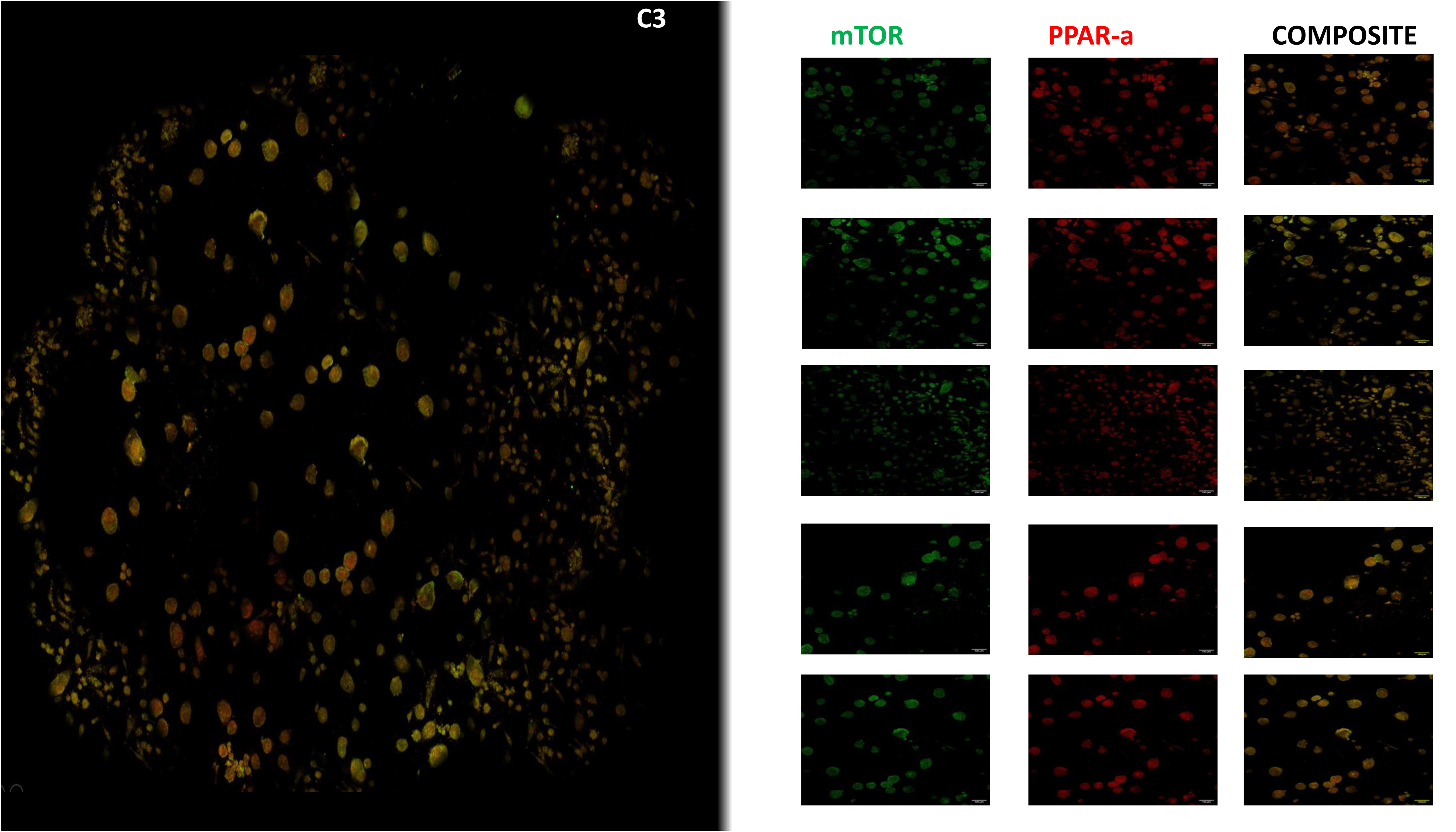
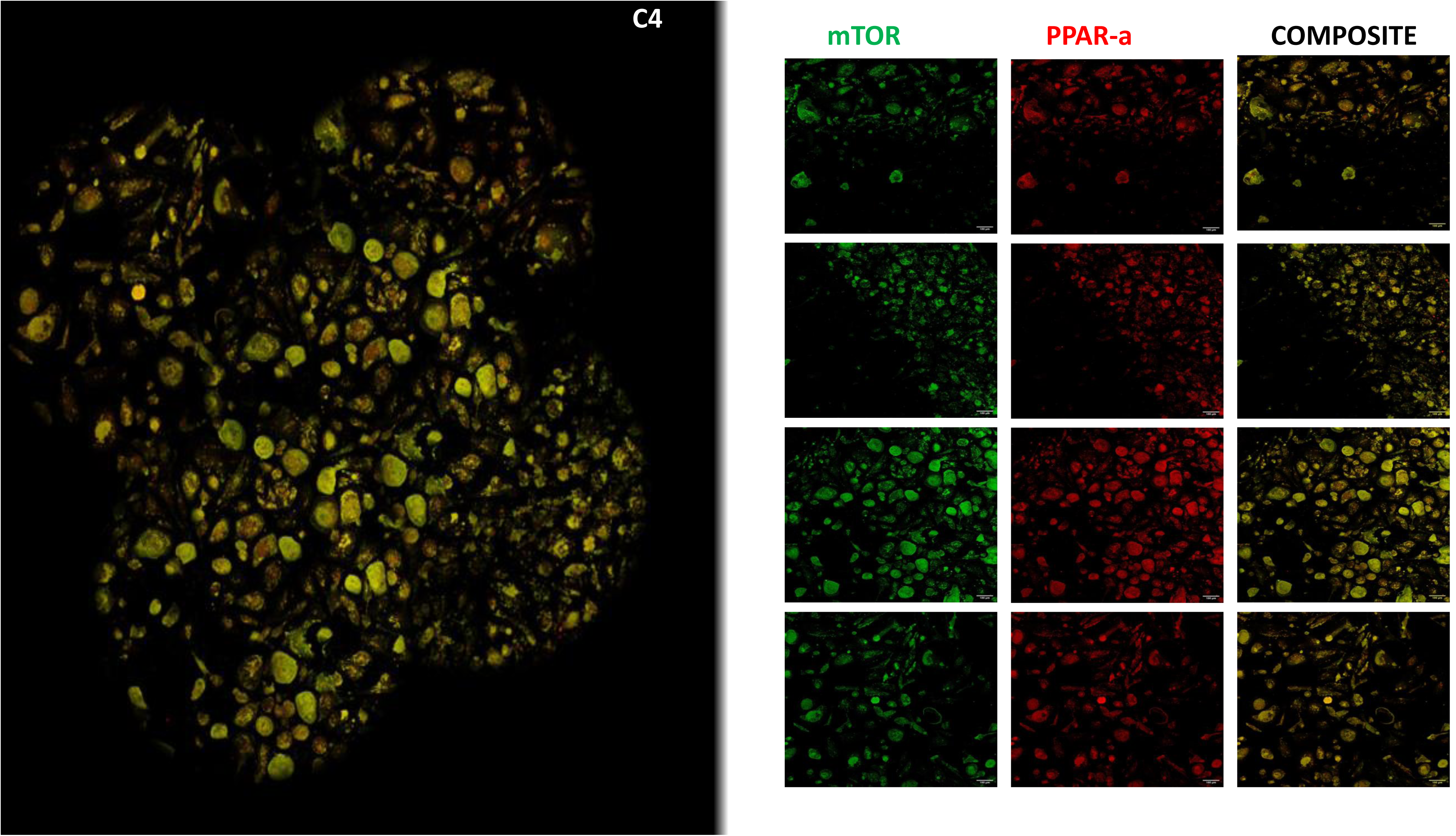
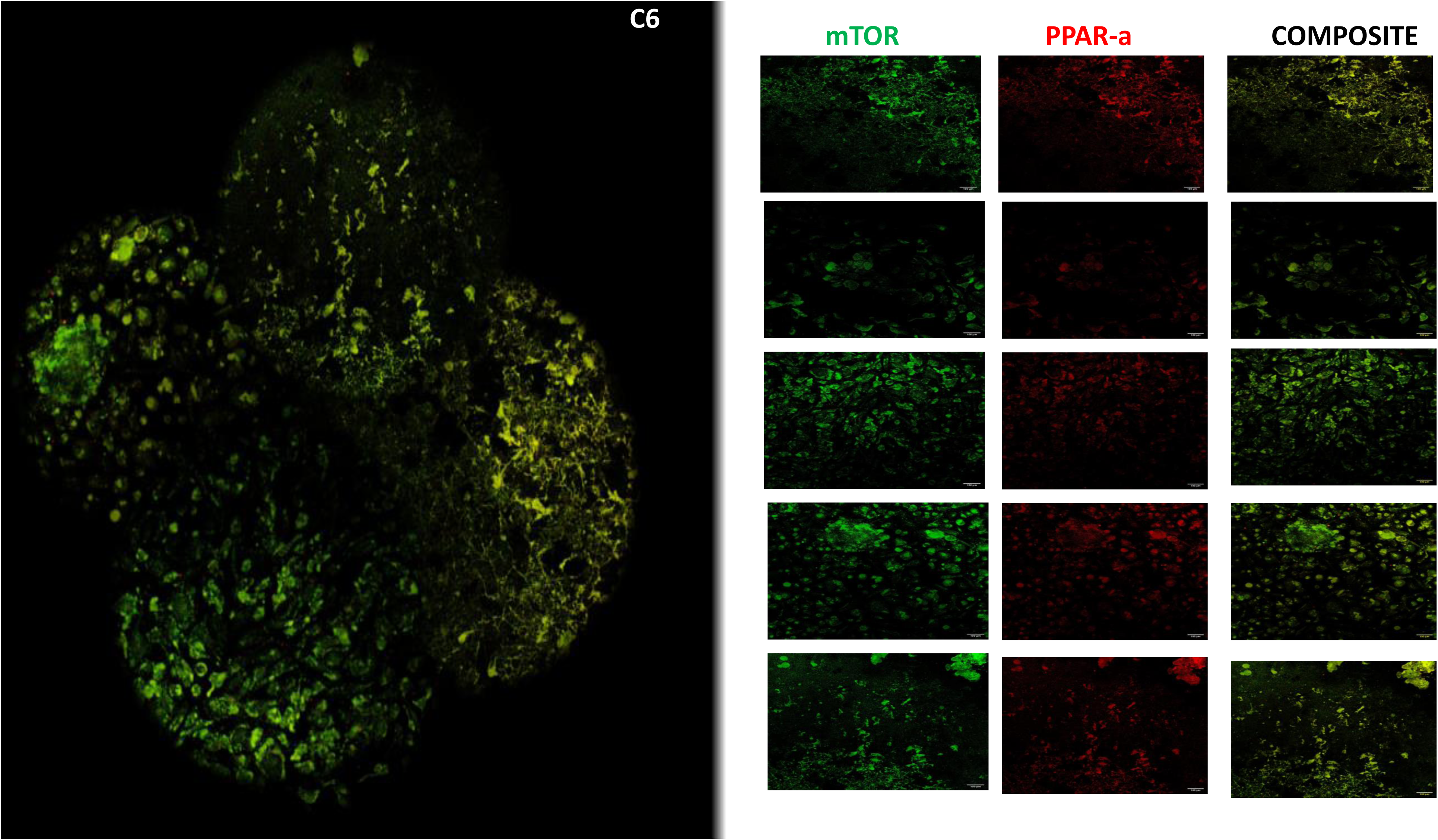
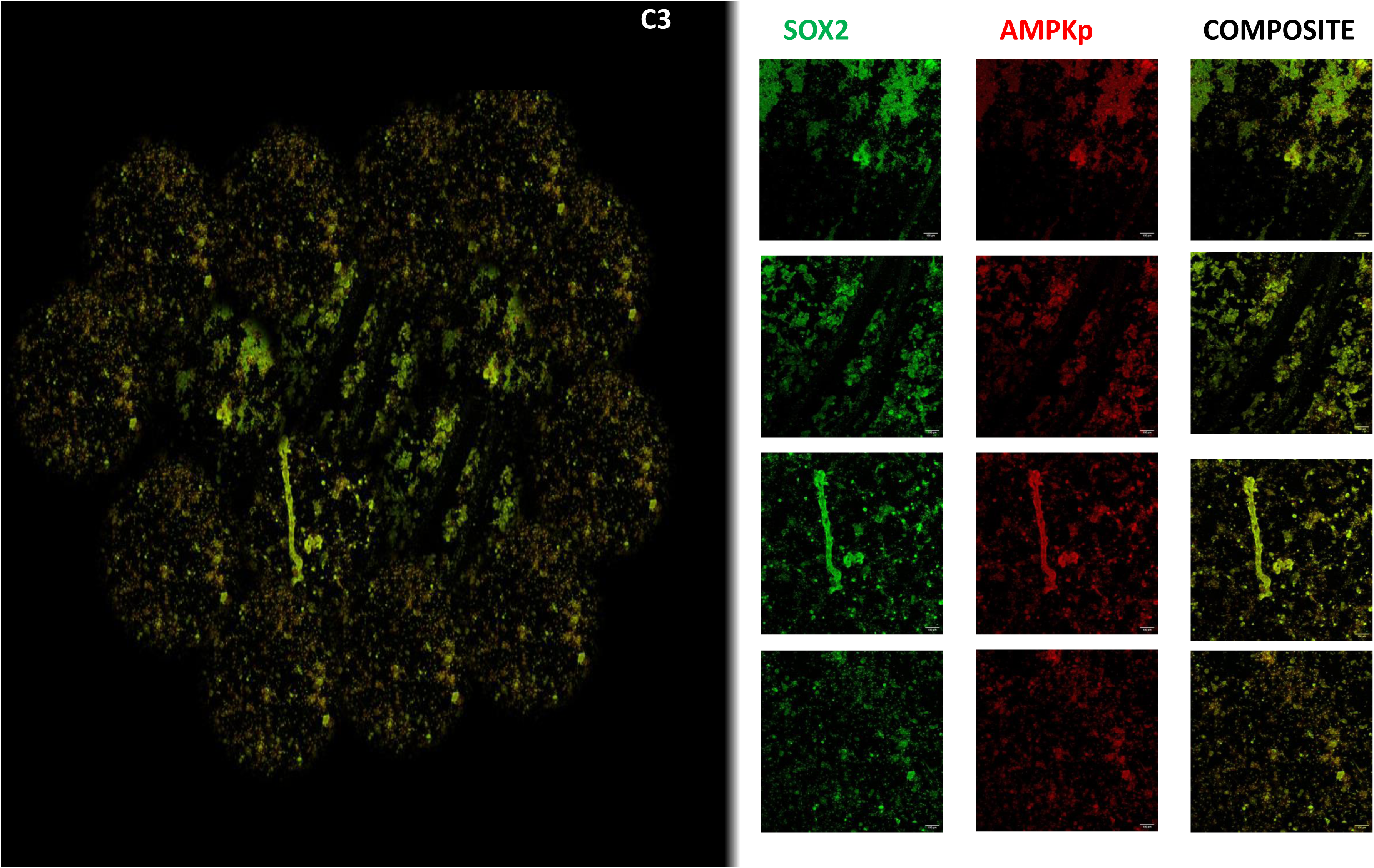
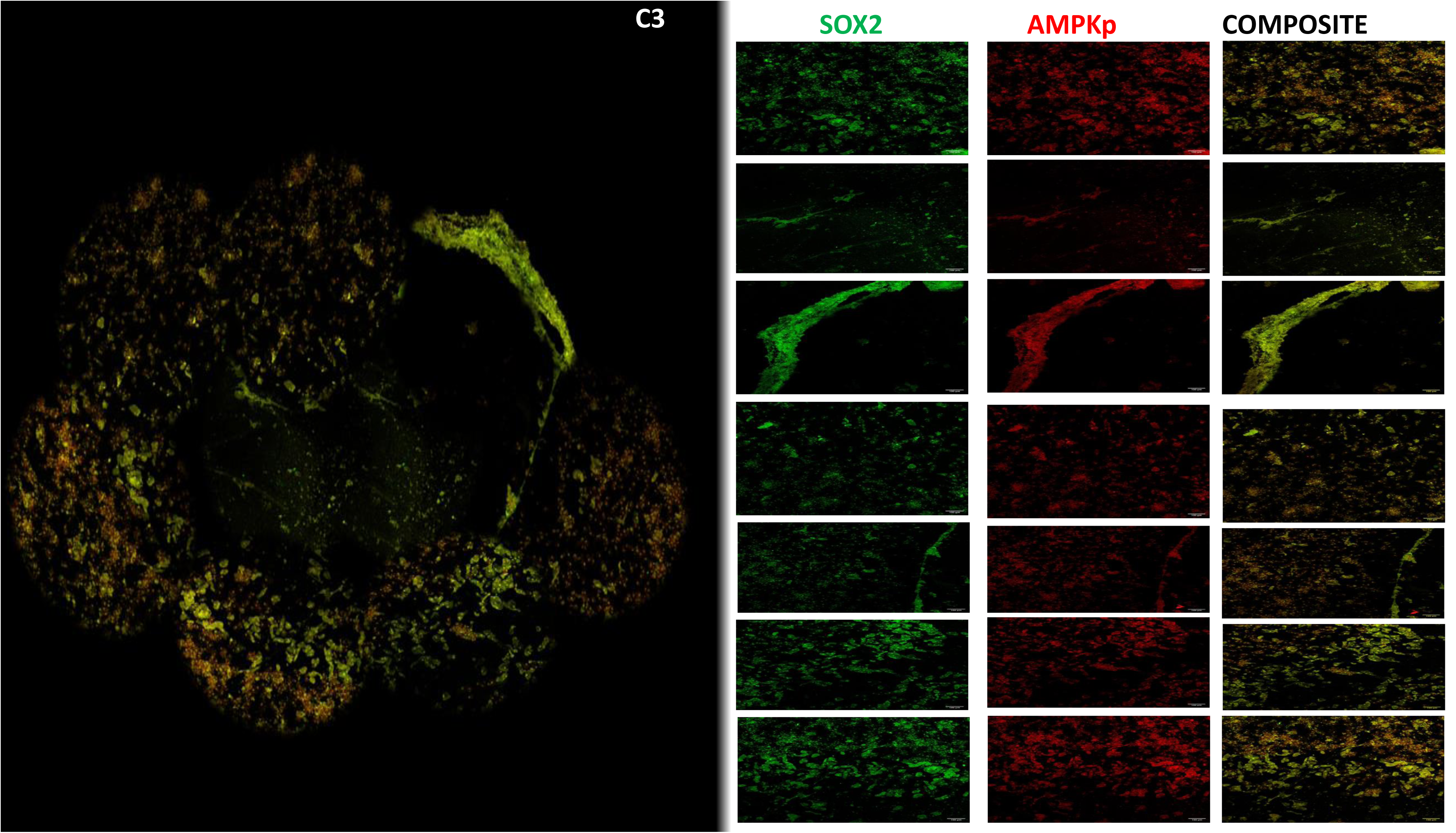
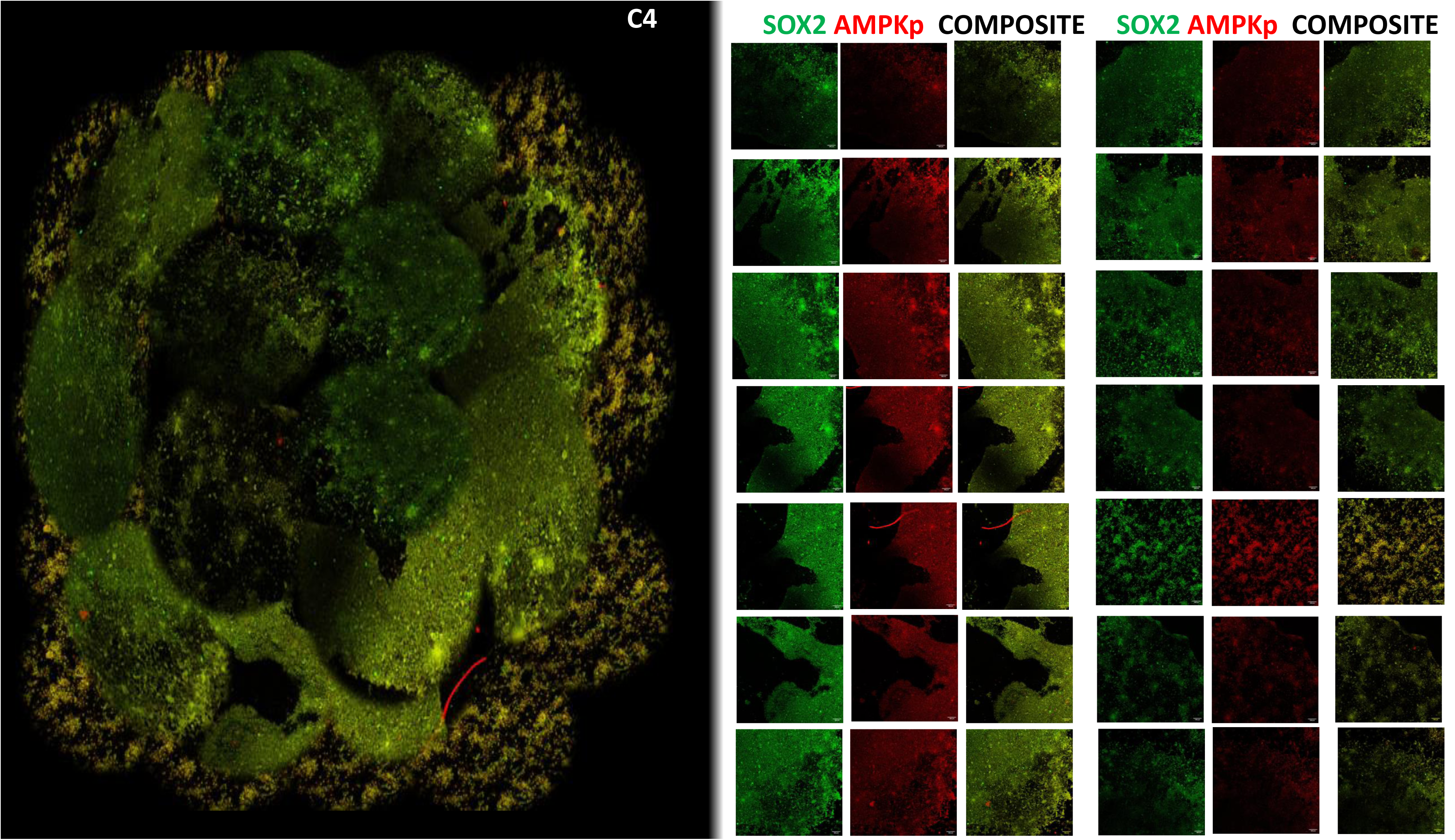
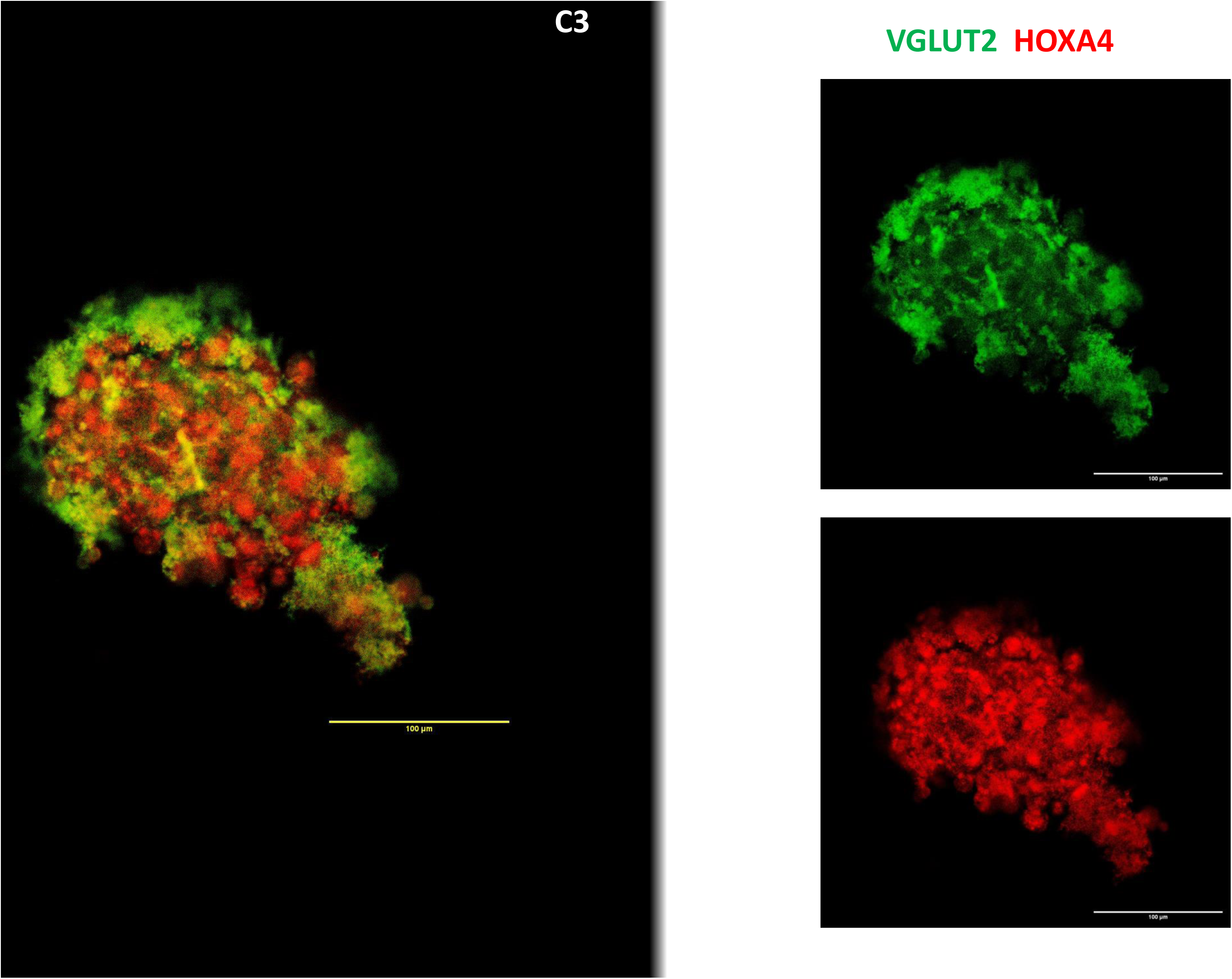
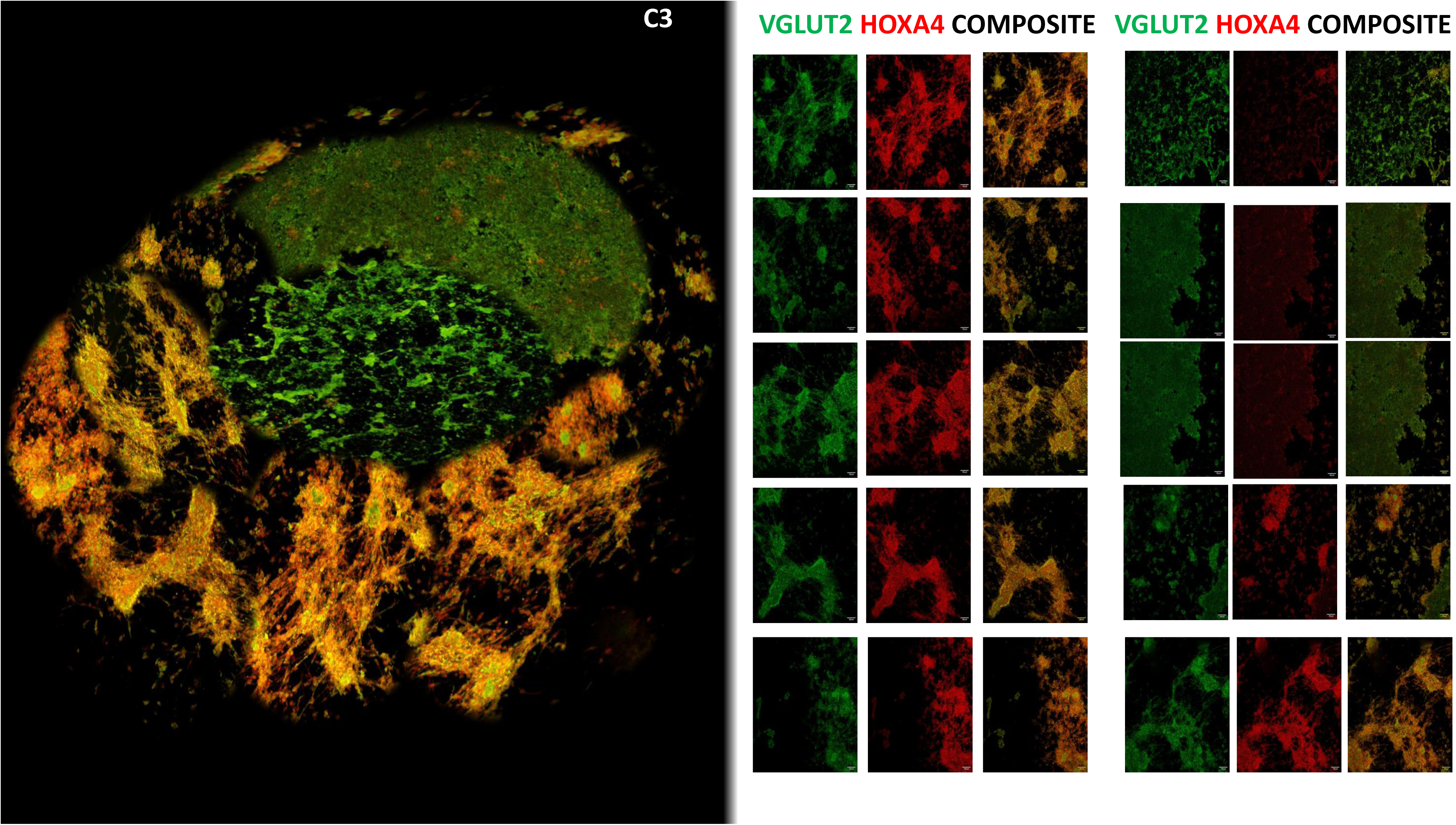
Stitched composite of abutting images of the neuro-vascularized tissue generated using PITTRep methodology.

**Figure 64.**
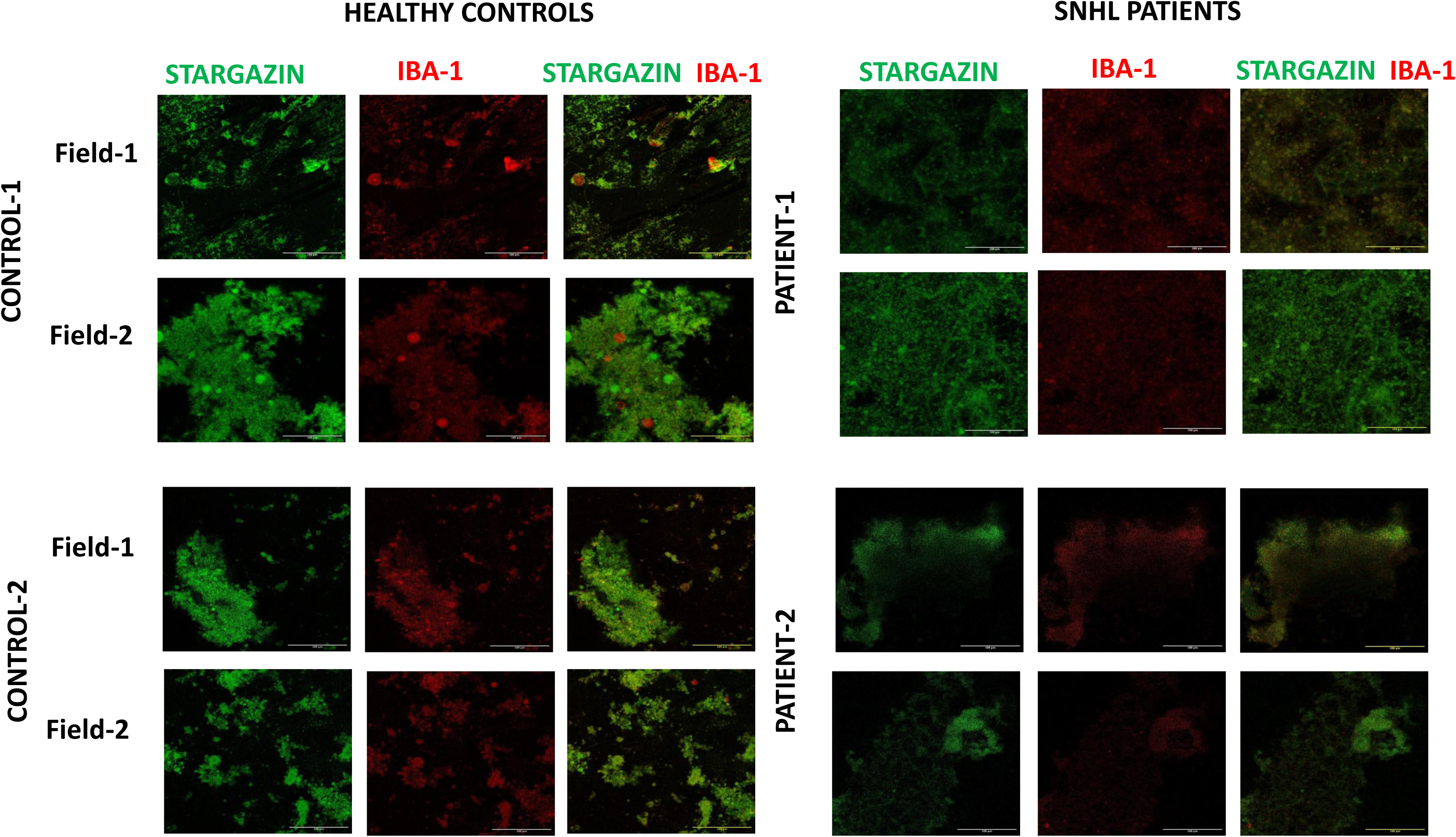
The figure shows comparison between the expression of Stargazin (green) a marker of synaptic functionality and Iba-1 (red), a microglia marker responsible for pruning day 14 of culture between Sensory neural hearing loss (SNHL) patient and healthy control with normal hearing (isogenic controls were taken in this case). The patient group depicts decreased expression of Stargazin and Iba-1 as compared to healthy control indicating the absence of mature synaptic connections due to less activation of microglia. Scale bars: 100 µm

**Figure 65.**
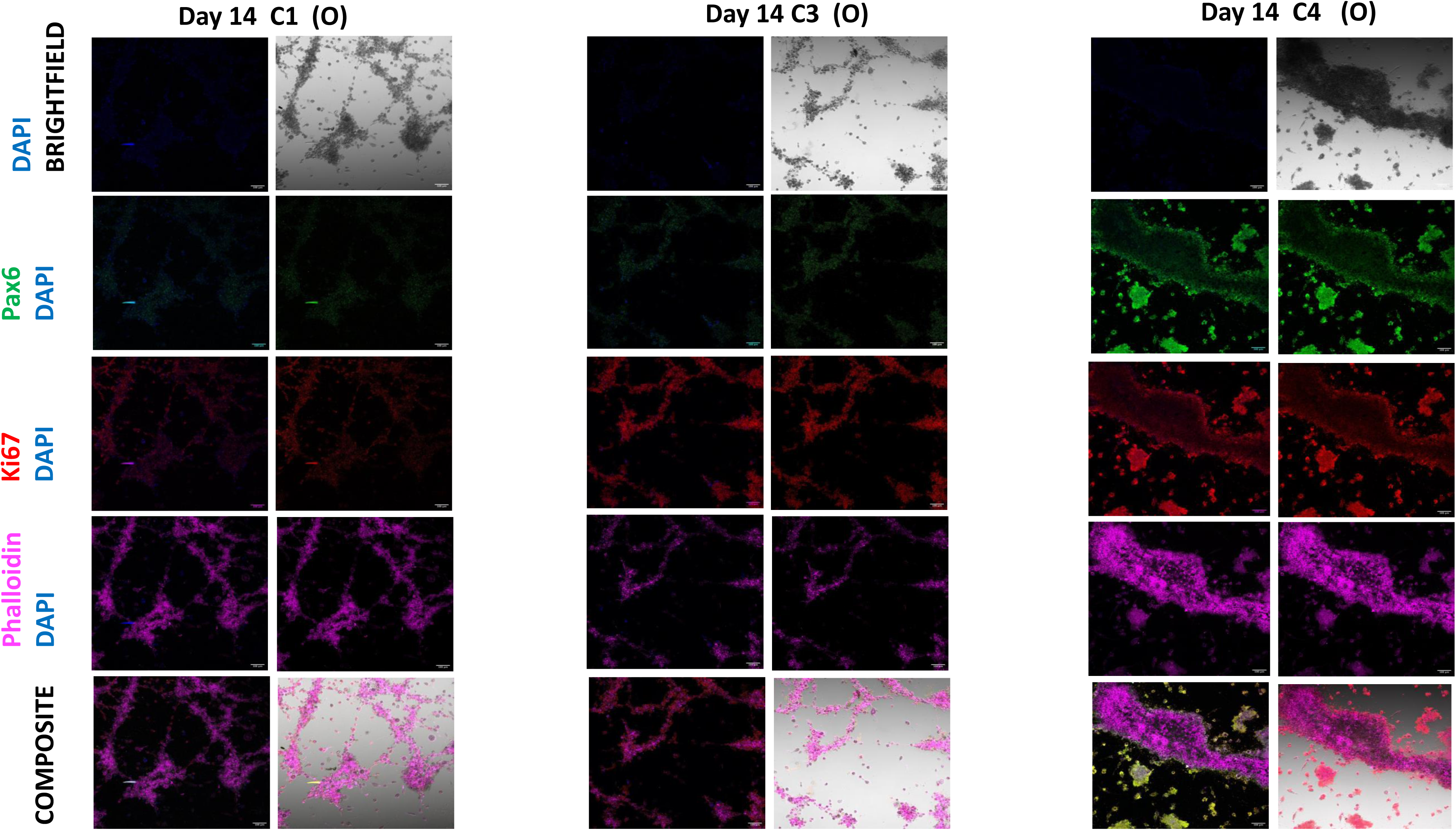
Emerging radial glia: Pax6 (green), which is a radial glia marker and Ki67 (red)which is a cell proliferation marker. At day-14, the expression of Pax6 and Ki67 increases from the condition C1-C3 indicating the increase in the proliferation of radial glia cells as the amount of RBC’s increase. Scale bars: 100 µm

**Figure 66.**
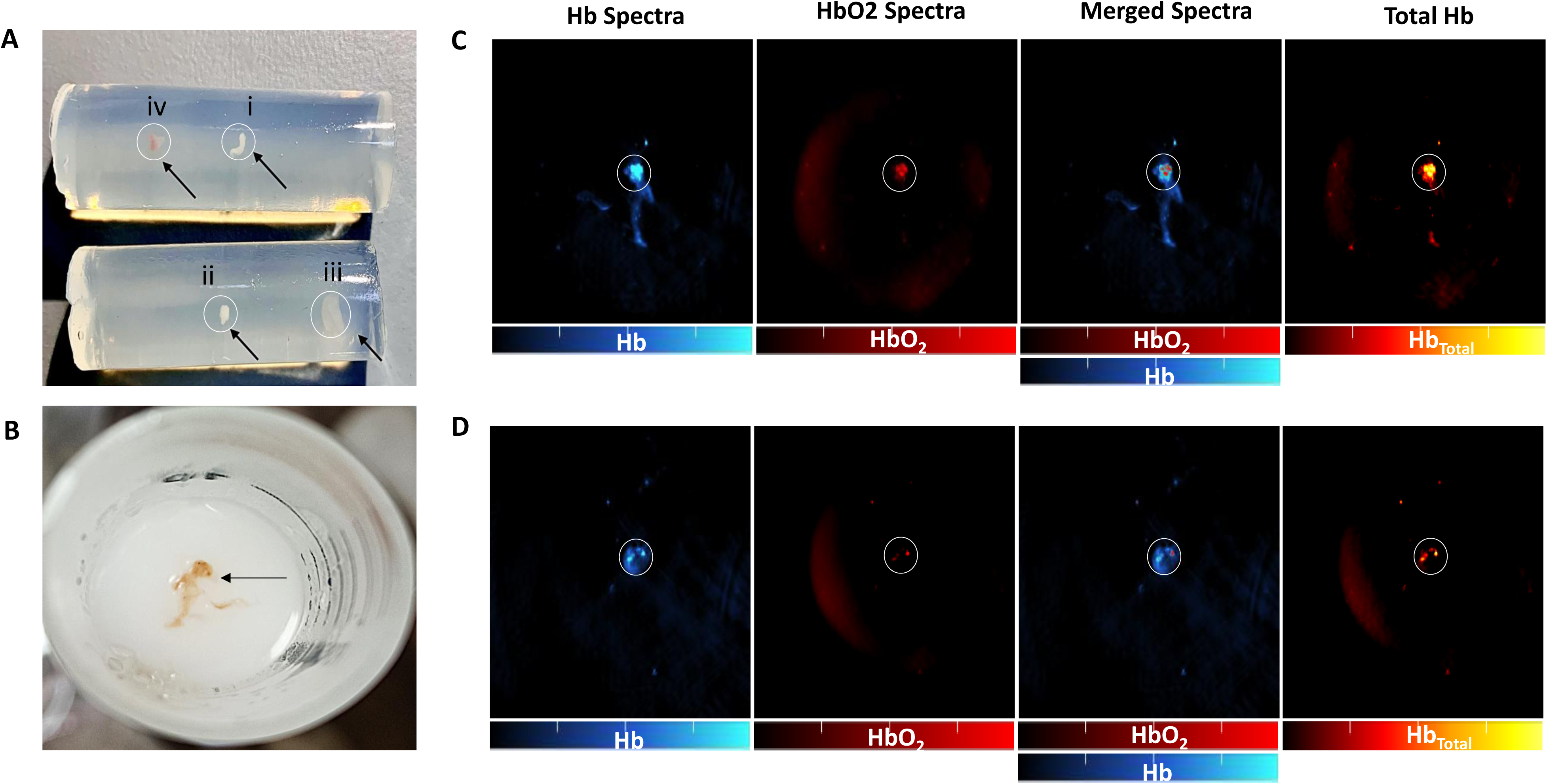
Optoacoustic imaging indicating the formation of neuro vascularised tissue: A) Neuro vascularized tissue (indicated by black arrows) of the conditions C1-C4 embedded in cylindrical phantoms prepared from low melting agarose where i= C4 condition, ii = C2 condition, iii= C3 condition and iv= C4 condition of the cultured tissue. B) A closer view of iv (C4 condition) neuro vascularised tissue placed in low melting agarose for optoacoustic imaging. C) and D) are scans of the neuro vascularised tissue (iv and iii respectively) obtained from the iTHERA MSOT inVision optoacoustic imaging device. Notice the defined Hb and HbO2 signal in localised in the boundary of neuro vascularised tissue. Some portions of this neuro vascularised tissue are richer in signal assigned to Hb and HbO_2_ (highlighted in circle) than the other parts. There is more intensified Hb-HbO2 signal in the scan of C4 condition (C) than C3 condition (D) neuro vascularised tissue.

