## Supplemental Data 1 for "Building Neurovascular tissue from autologous blood for modeling brain activity"

C1

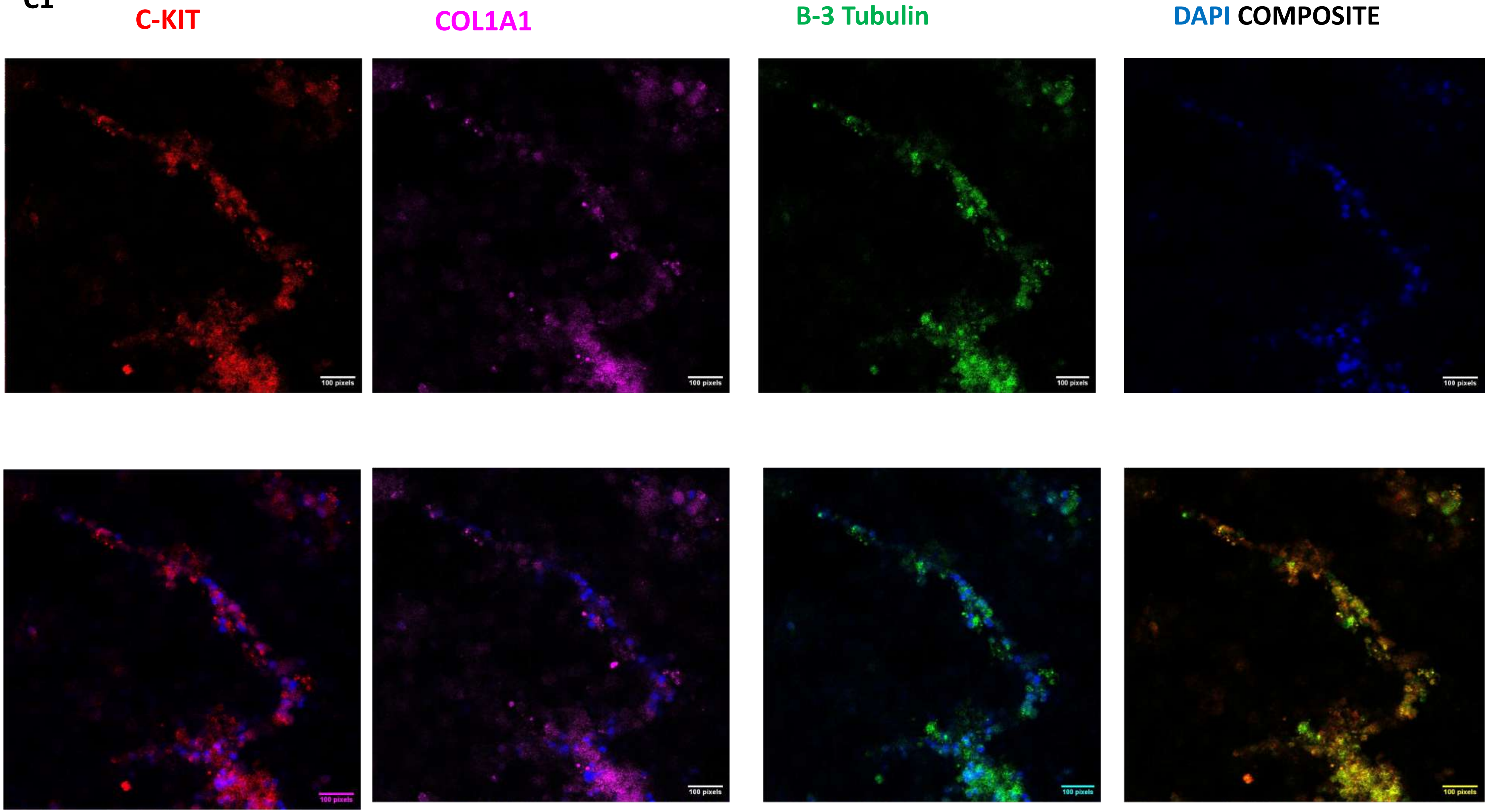
C-KIT
COL1A1
B-3 Tubulin
DAPI COMPOSITE
100 pixels
100 pixels
100 pixels
100 pixels
100 pixels
100 pixels
100 pixels
100 pixels

C1

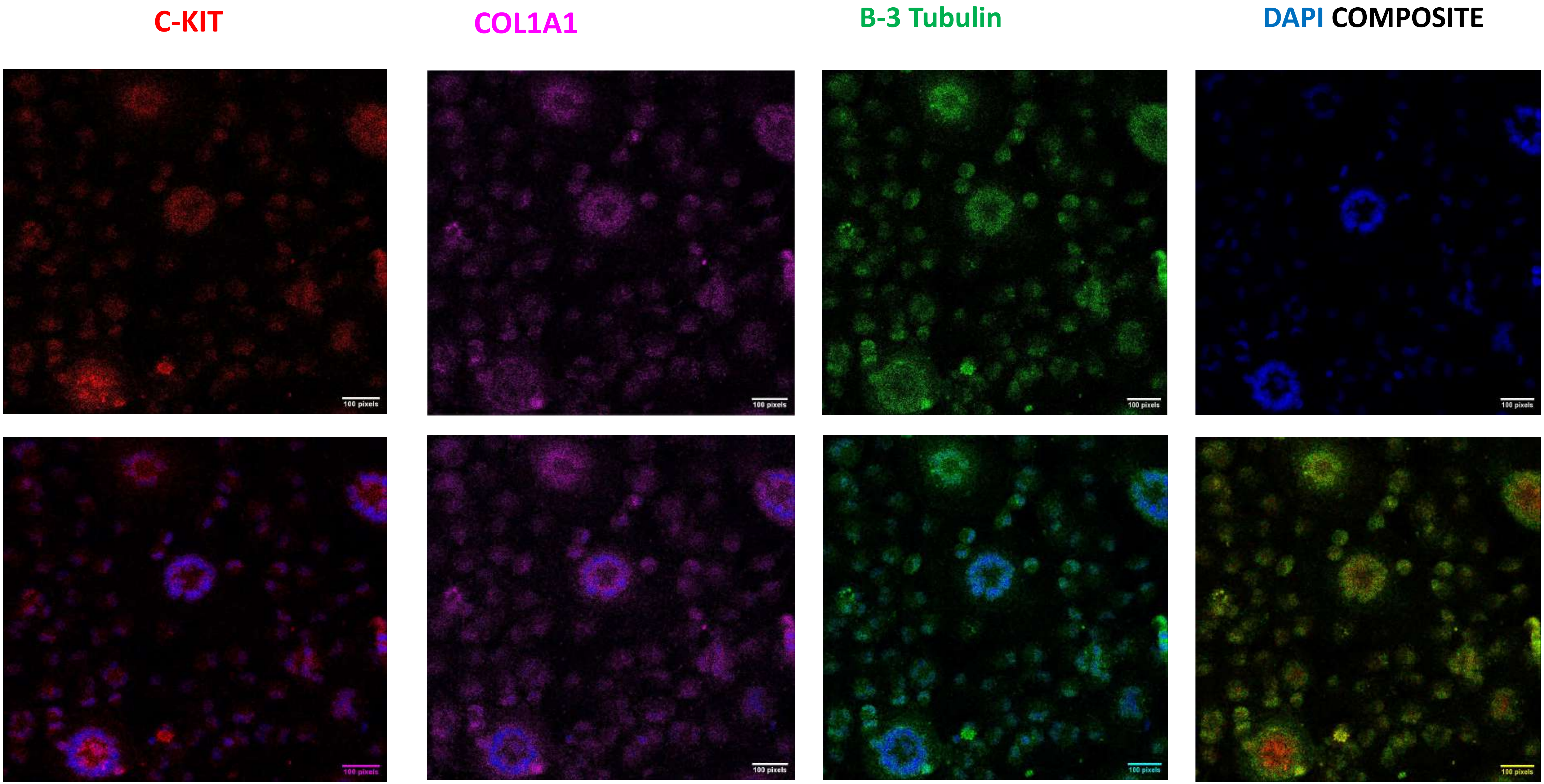

C1

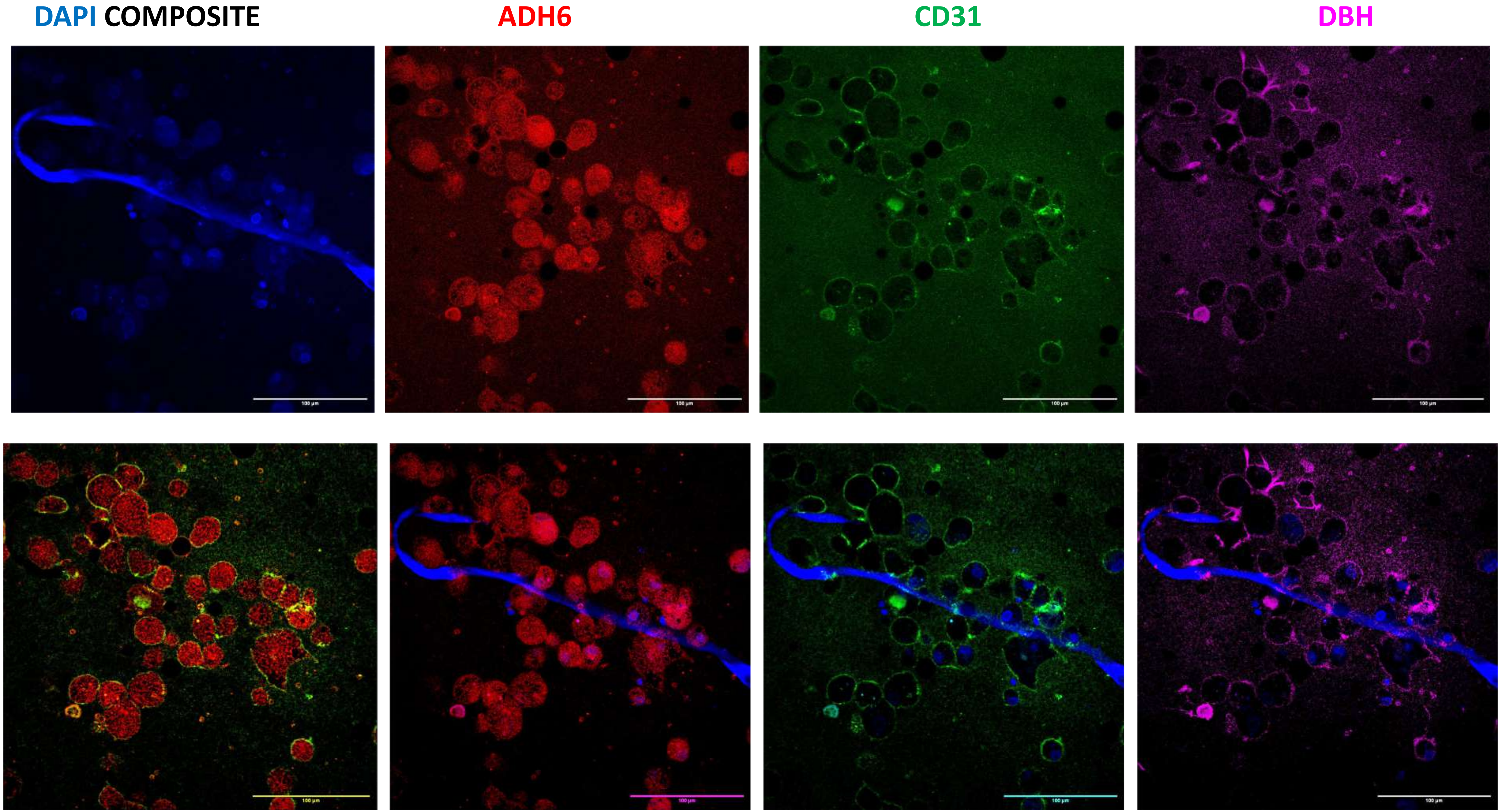

C1

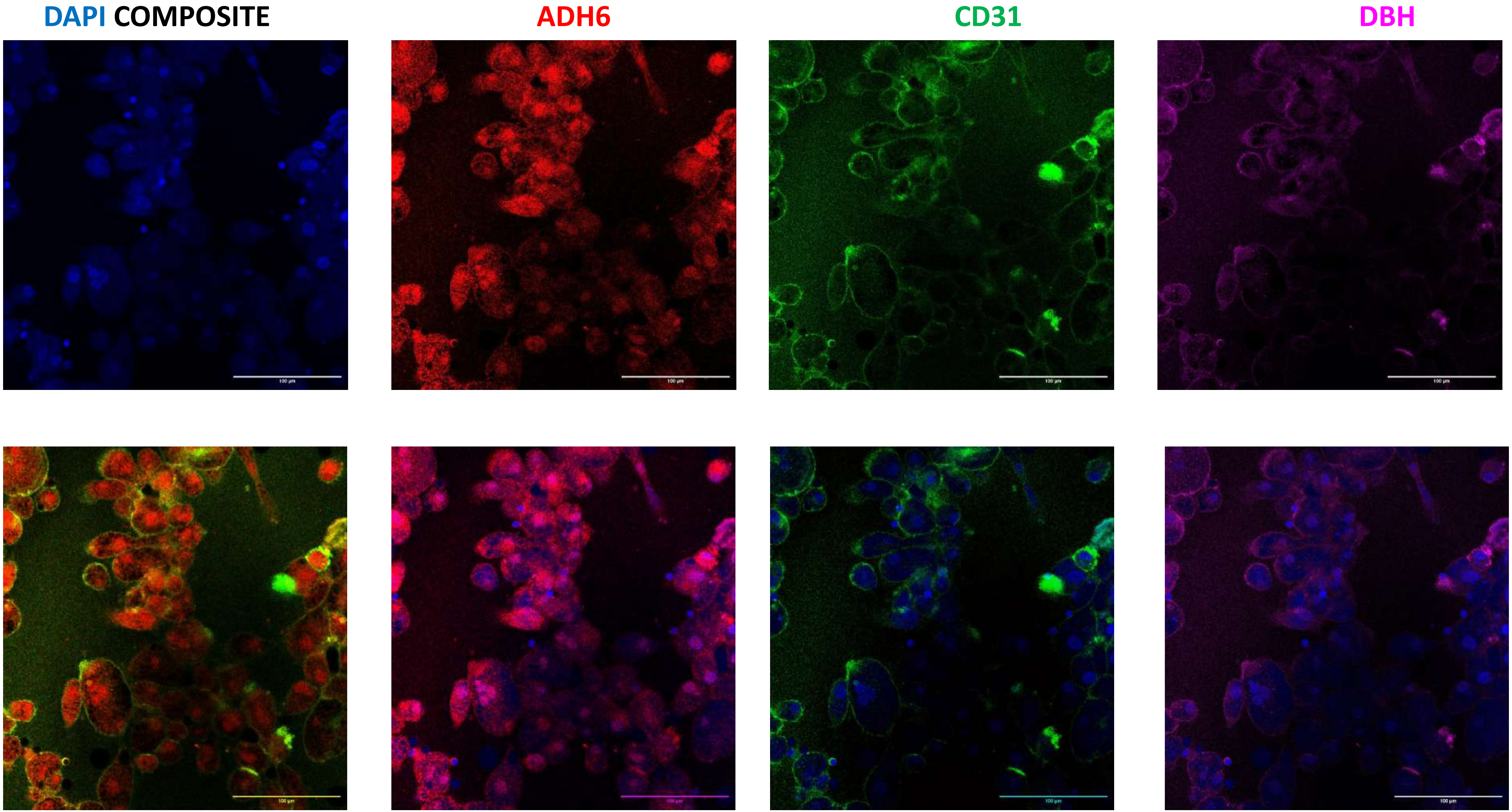

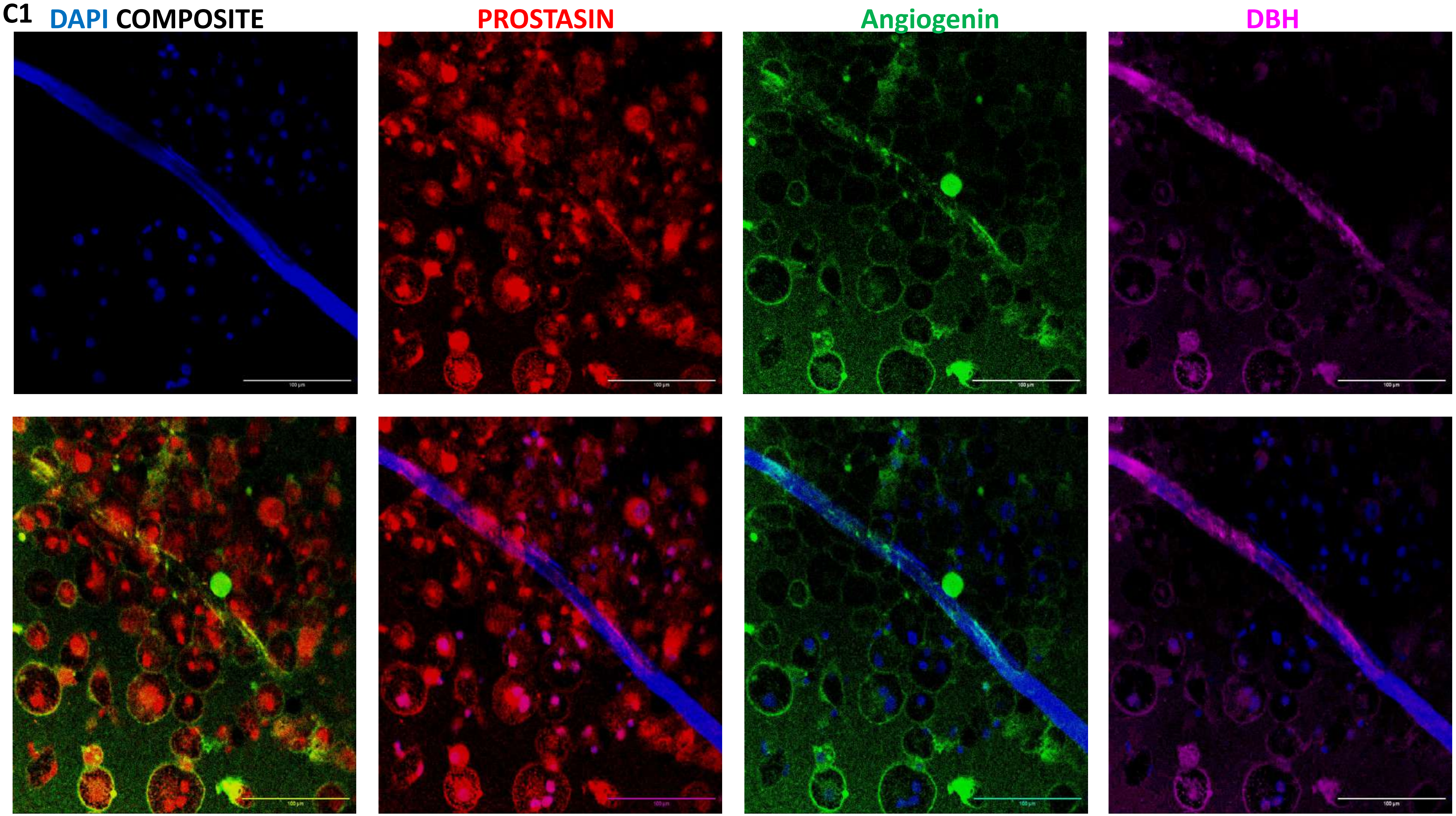
C1
DAPI COMPOSITE
PROSTASIN
Angiogenin
DBH

C1

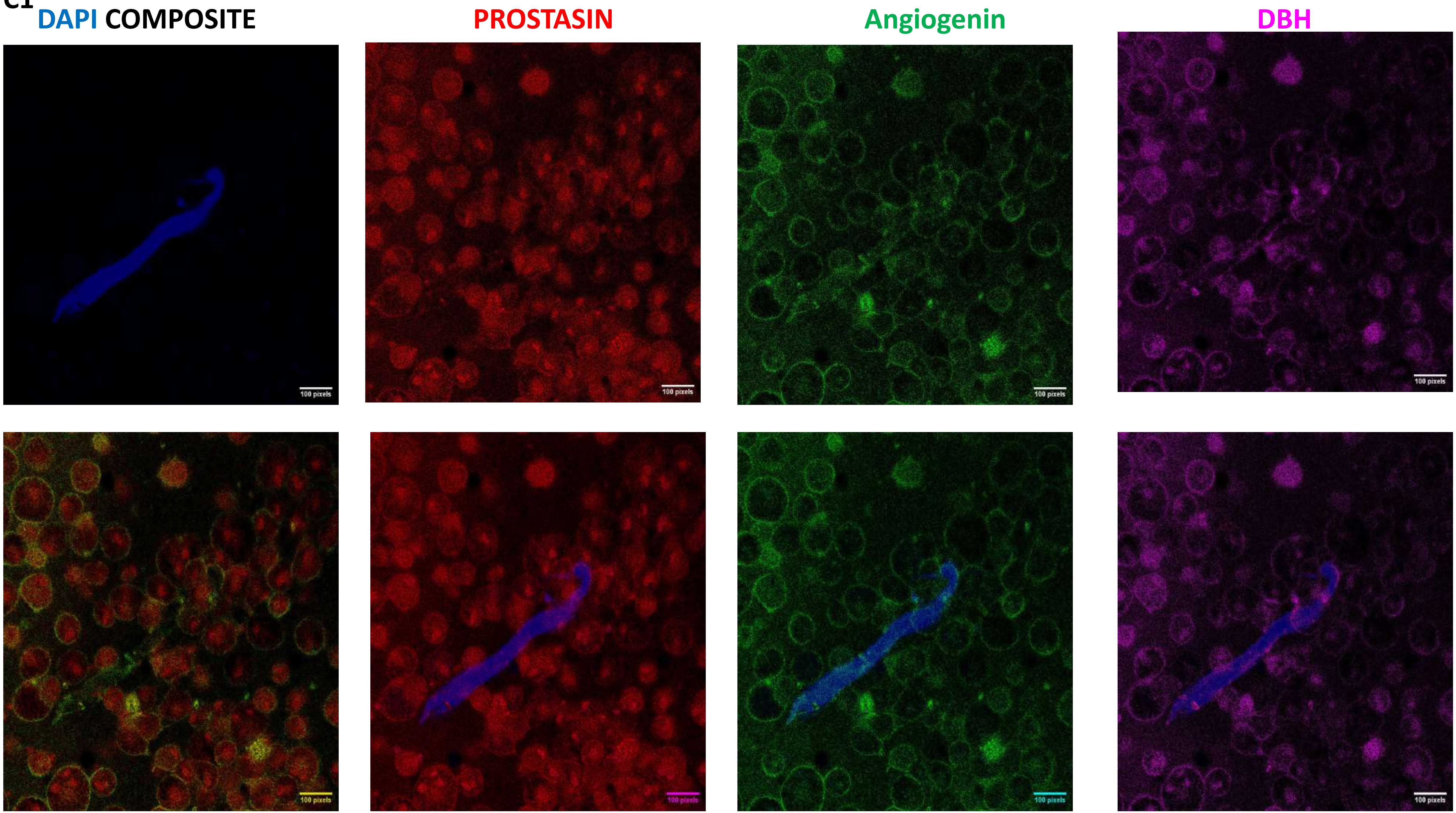
DAPI COMPOSITE
PROSTASIN
Angiogenin
DBH
100 pixels
100 pixels
100 pixels
100 pixels
100 pixels
100 pixels
100 pixels
100 pixels

C1

DAPI COMPOSITE | PROSTASIN | Angiogenin | DBH

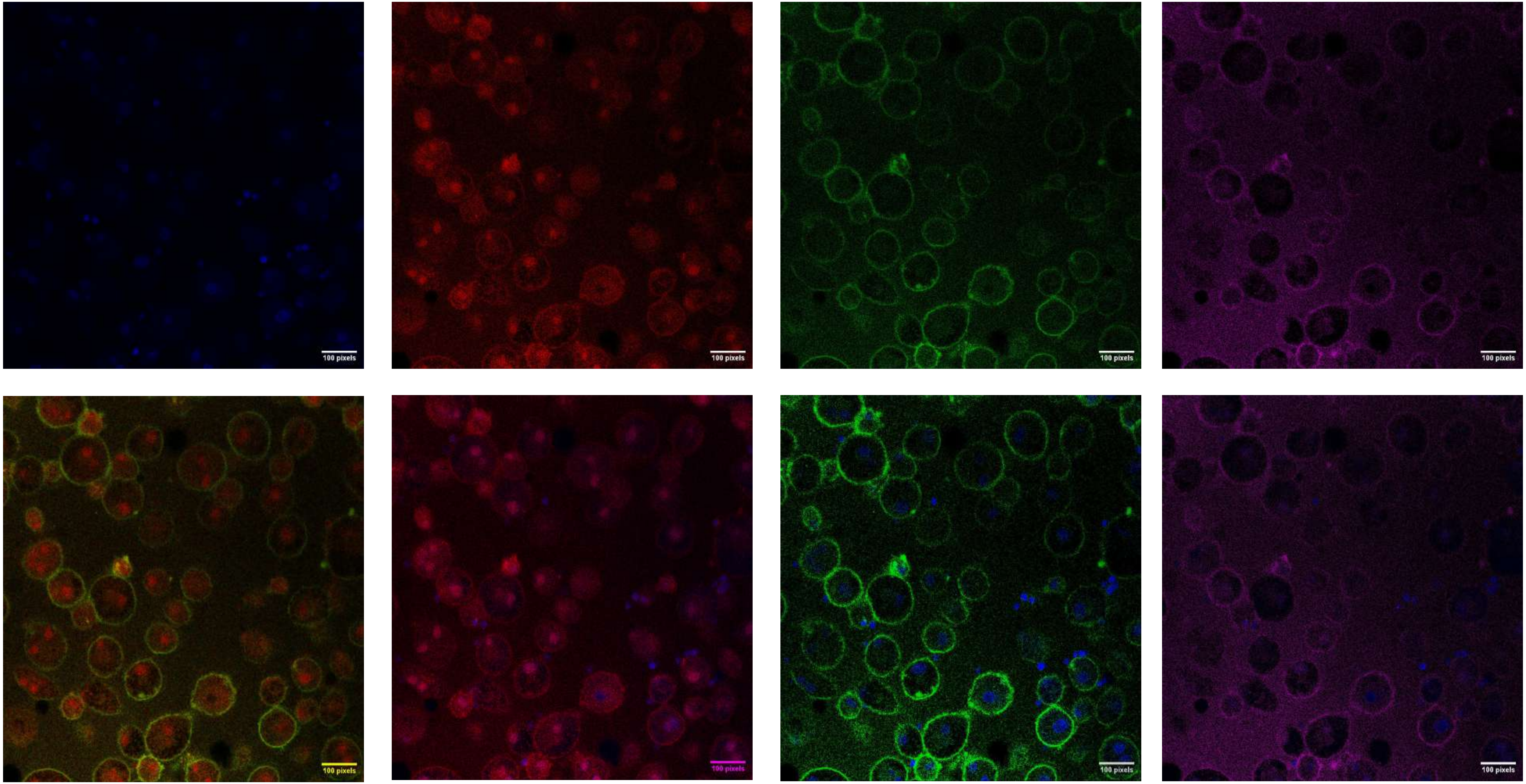

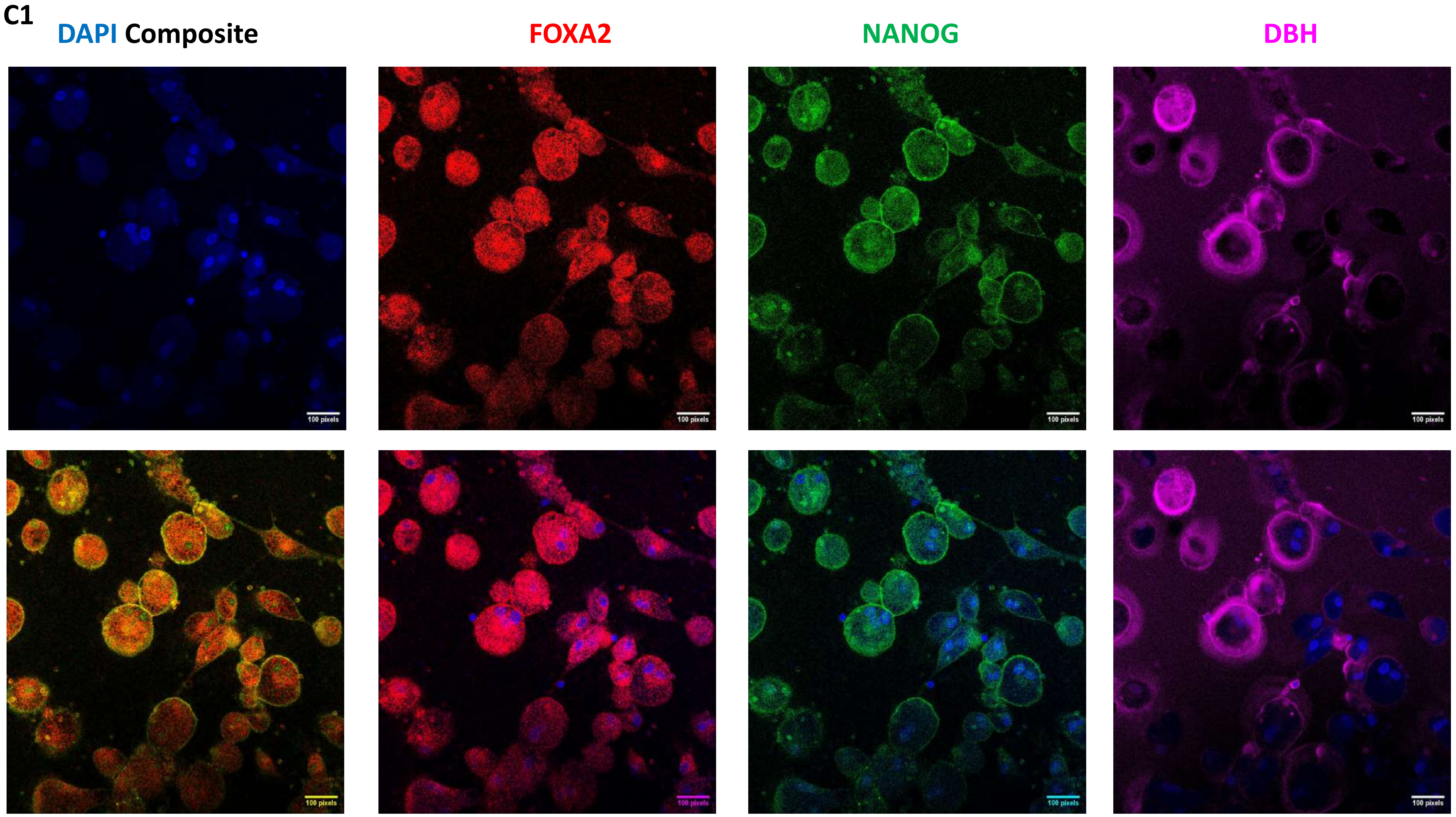
C1
DAPI Composite
FOXA2
NANOG
DBH
100 pixels
100 pixels
100 pixels
100 pixels
100 pixels
100 pixels
100 pixels
100 pixels

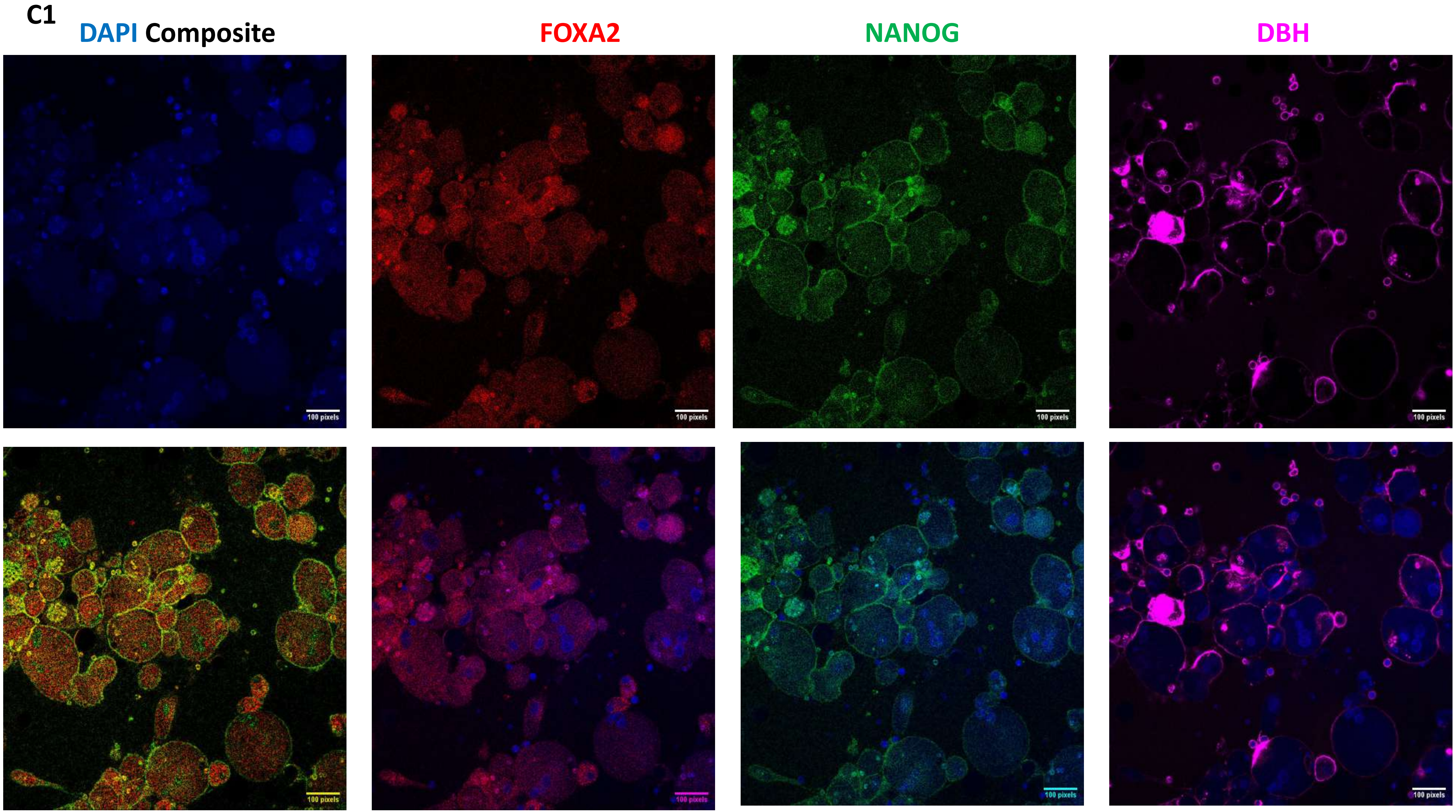
C1
DAPI Composite
FOXA2
NANOG
DBH
100 pixels

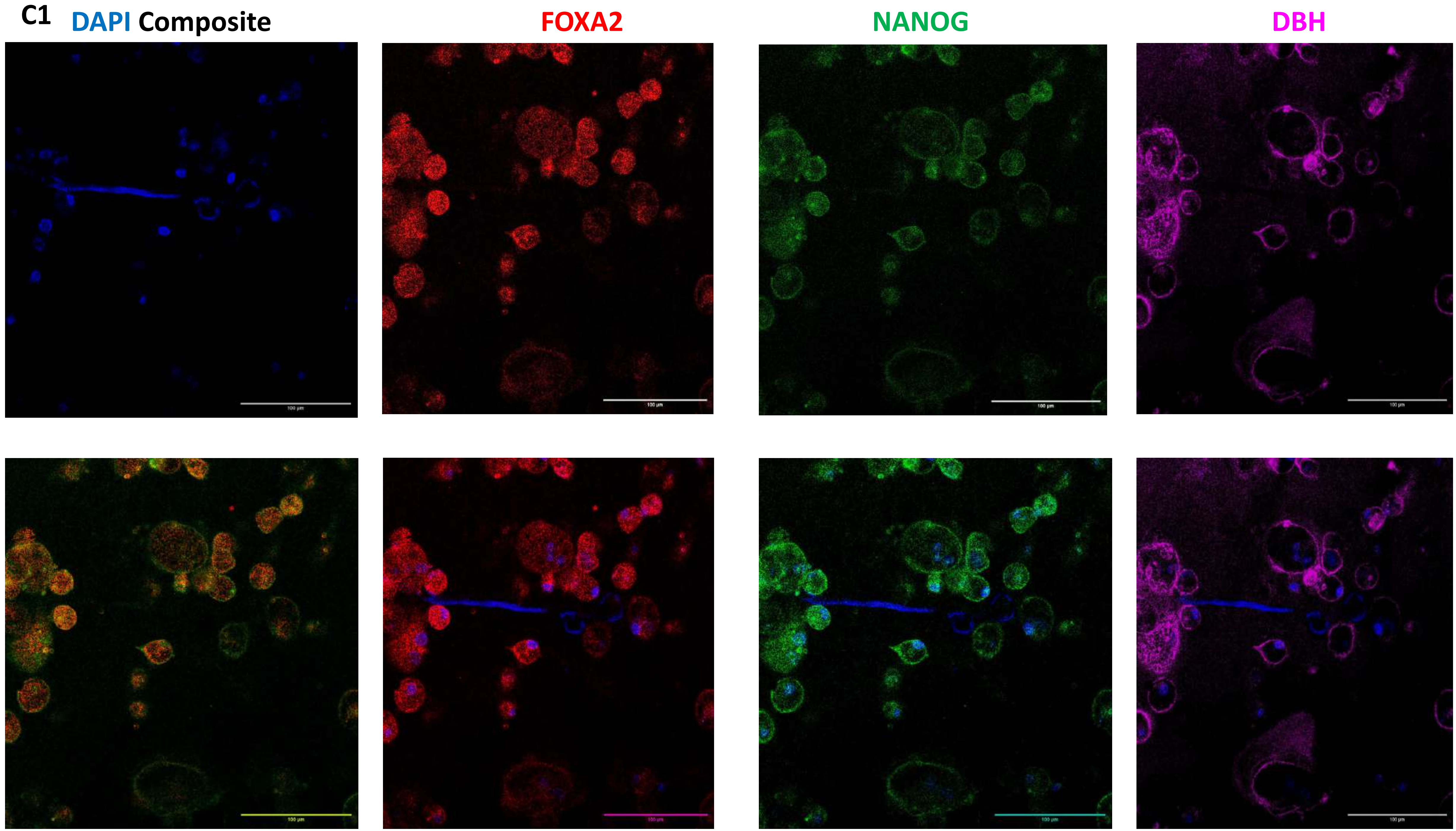
C1
DAPI
Composite
FOXA2
NANOG
DBH

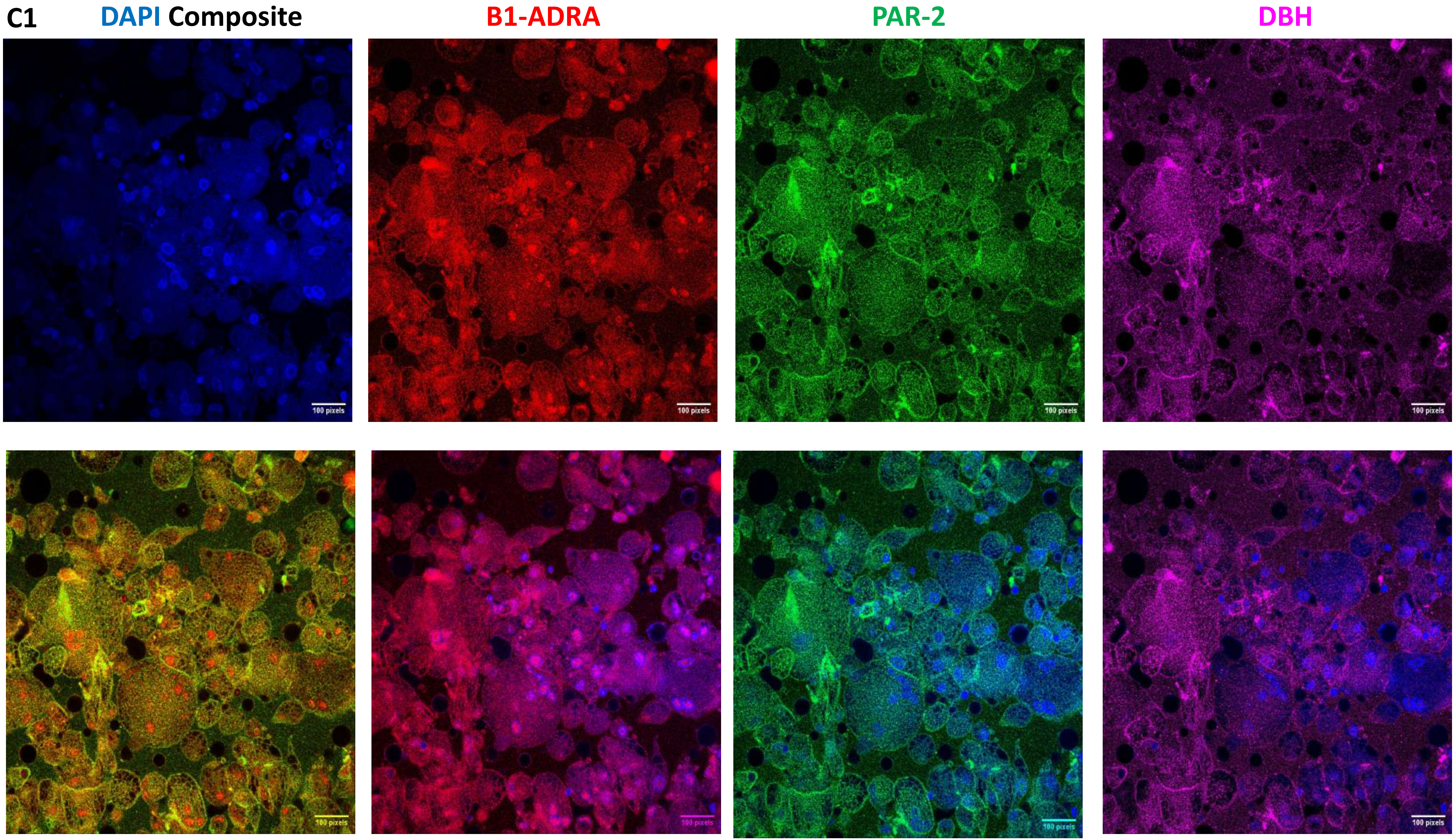
C1
DAPI Composite
B1-ADRA
PAR-2
DBH
100 pixels

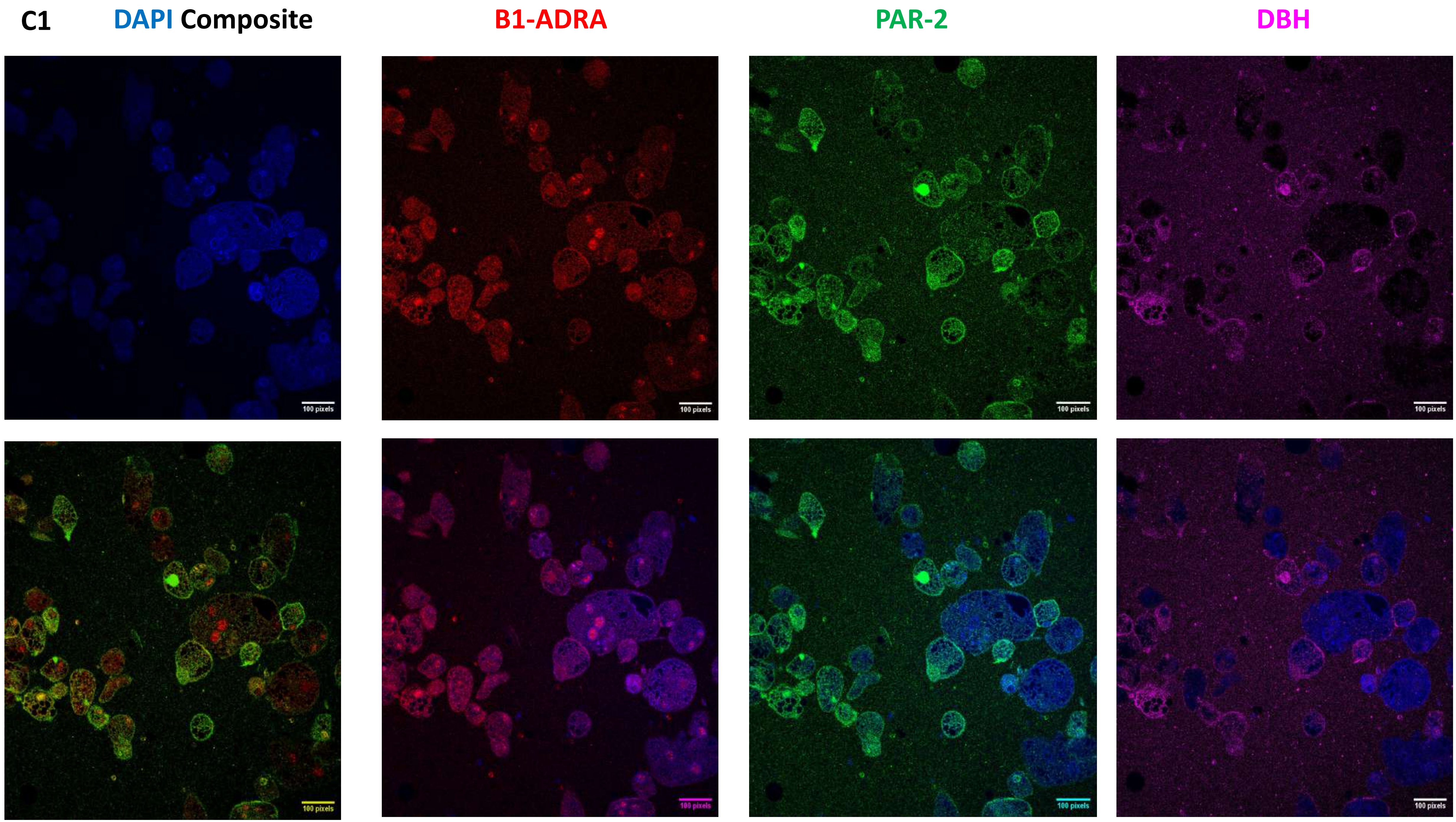
C1
DAPI Composite
B1-ADRA
PAR-2
DBH
100 pixels

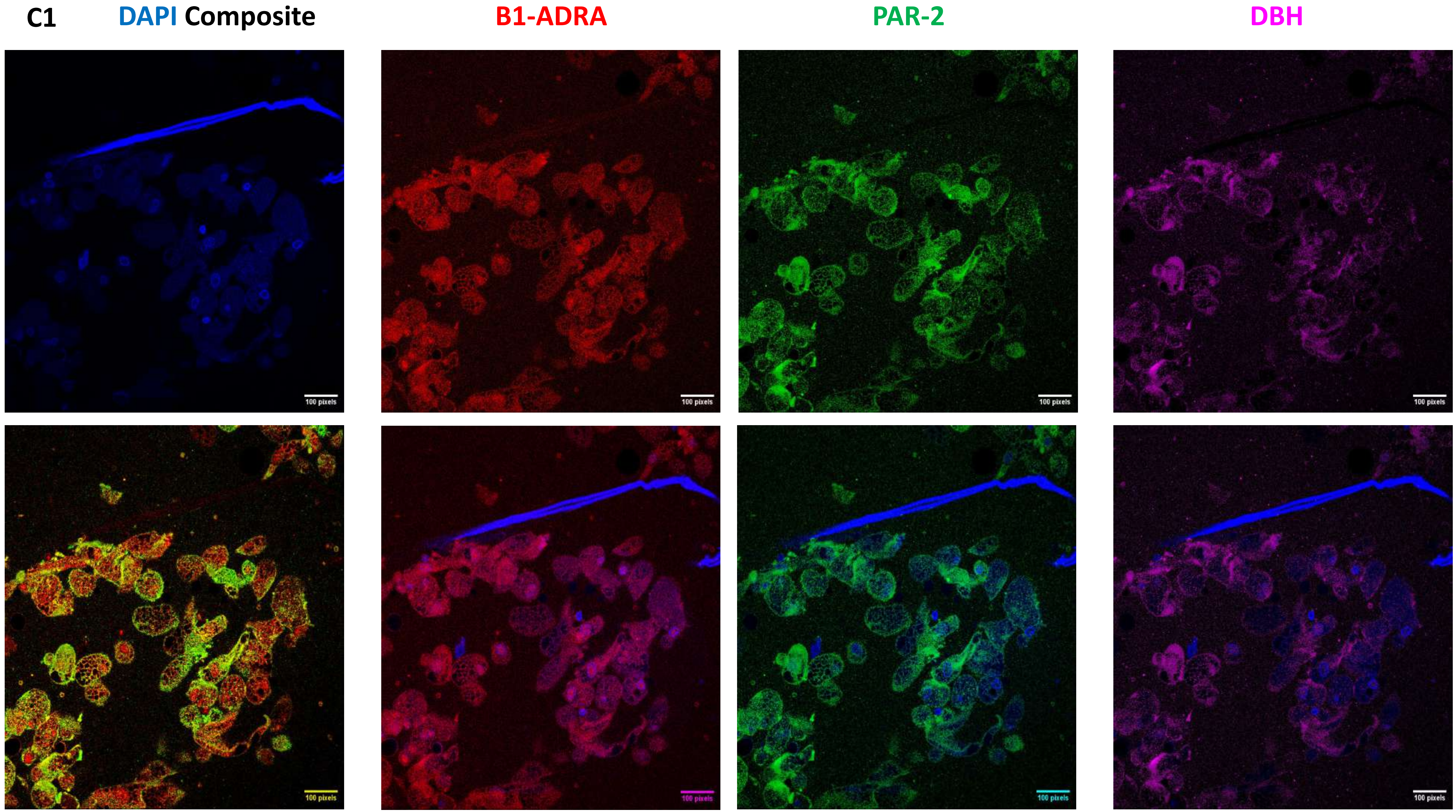
C1
DAPI Composite
B1-ADRA
PAR-2
DBH
100 pixels
100 pixels
100 pixels
100 pixels
100 pixels
100 pixels
100 pixels
100 pixels

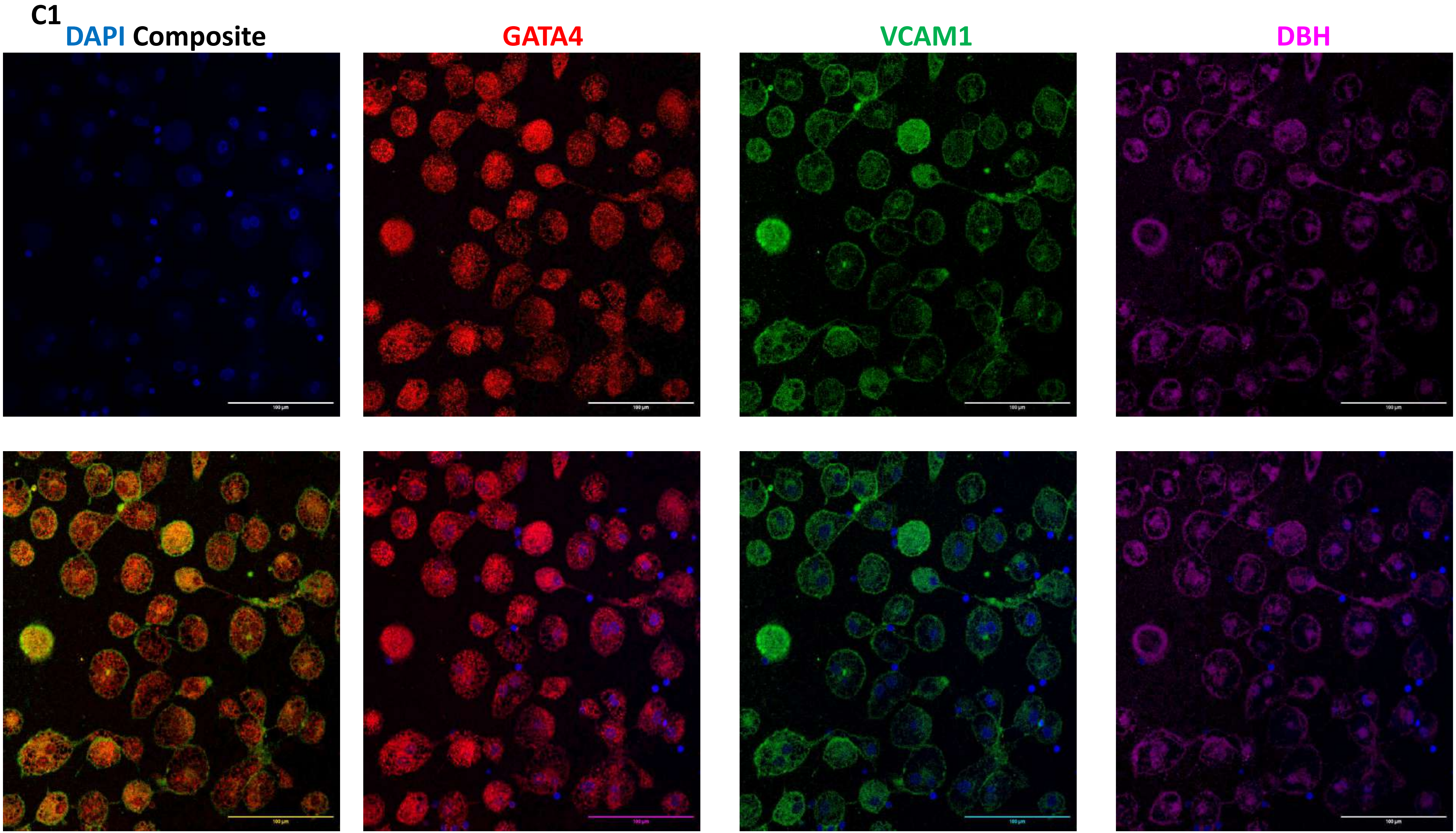
C1
DAPI Composite
GATA4
VCAM1
DBH

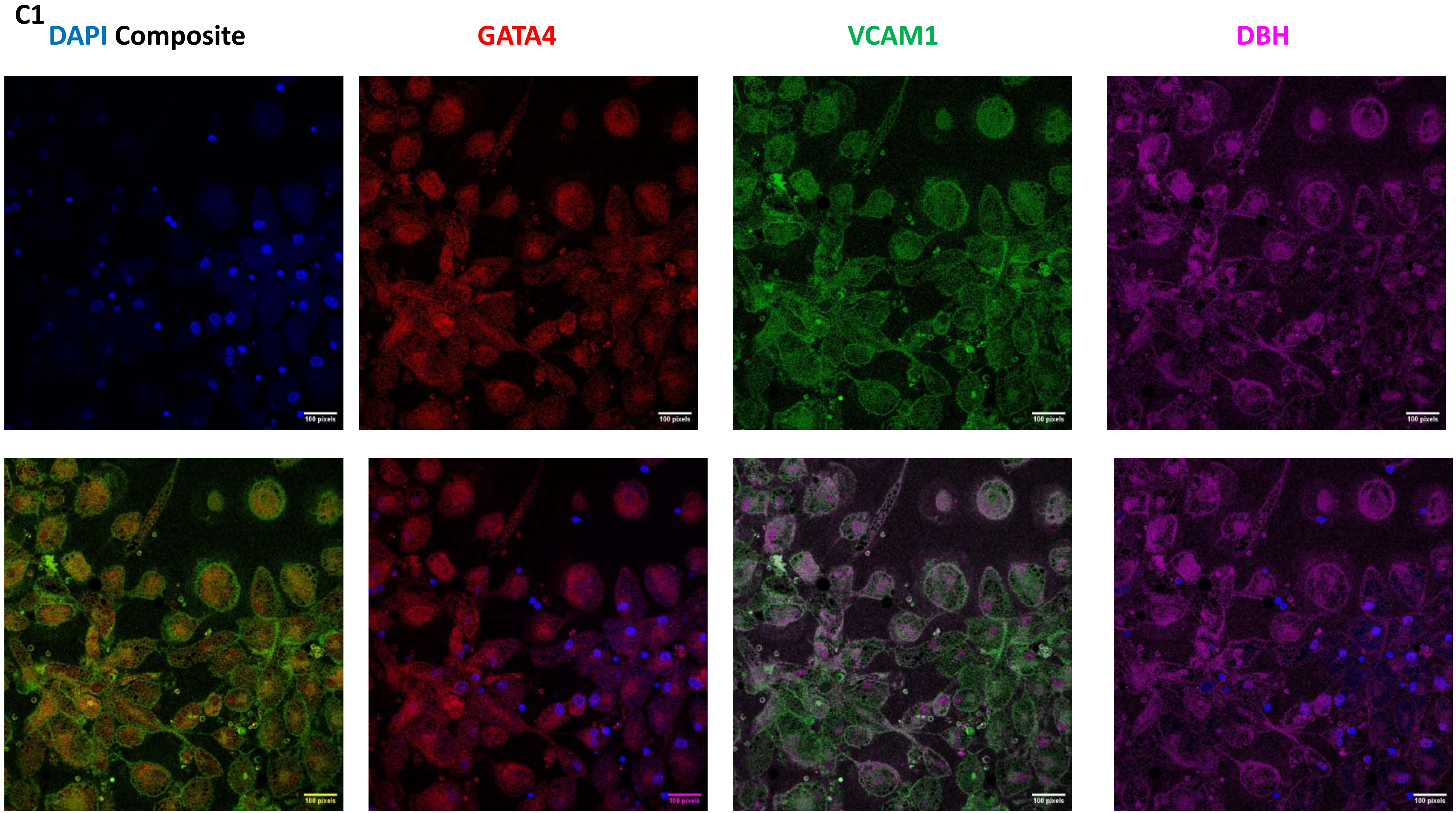
C1
DAPI Composite
GATA4
VCAM1
DBH
100 pixels

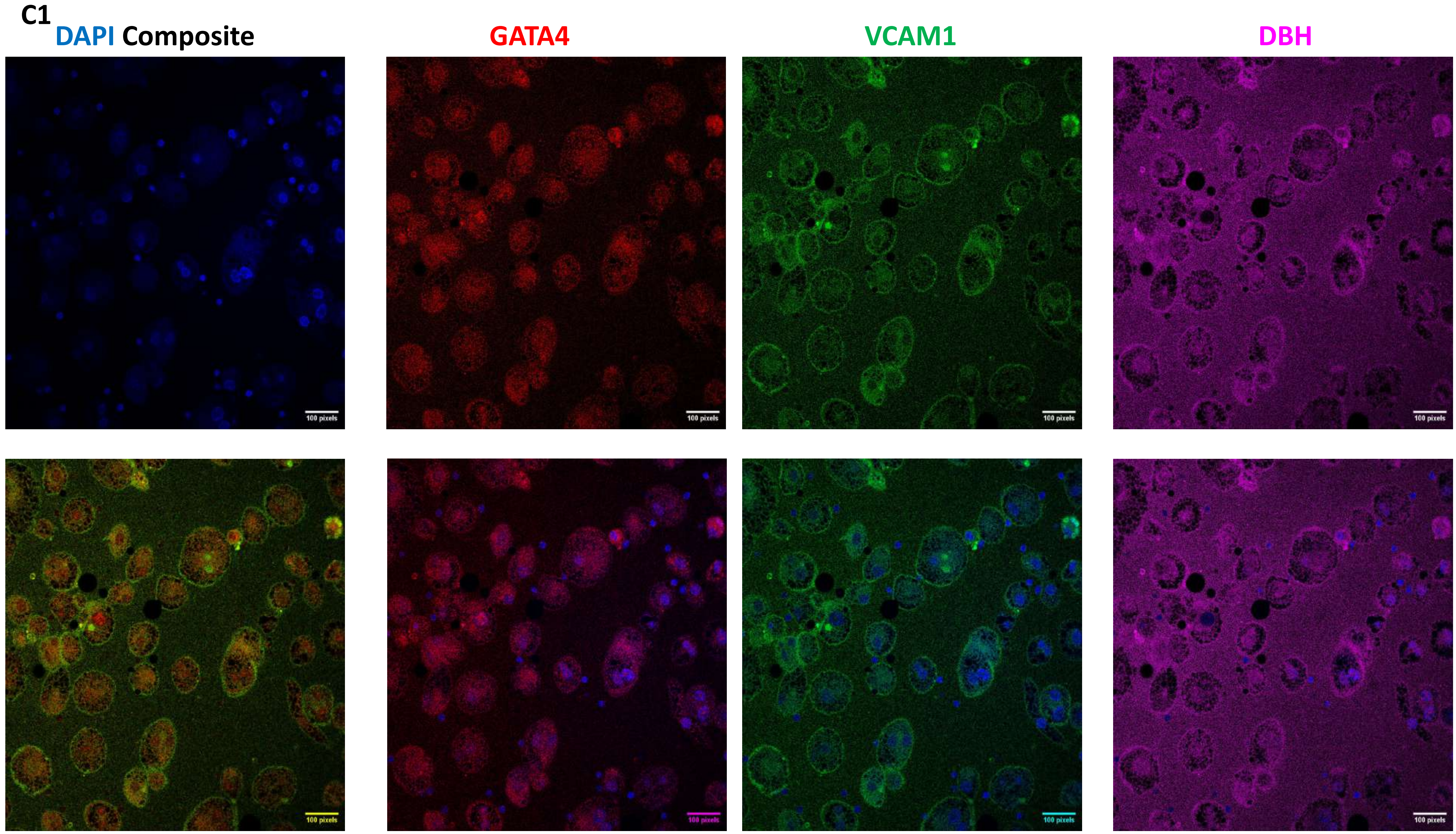
C1
DAPI Composite
GATA4
VCAM1
DBH
100 pixels

C1

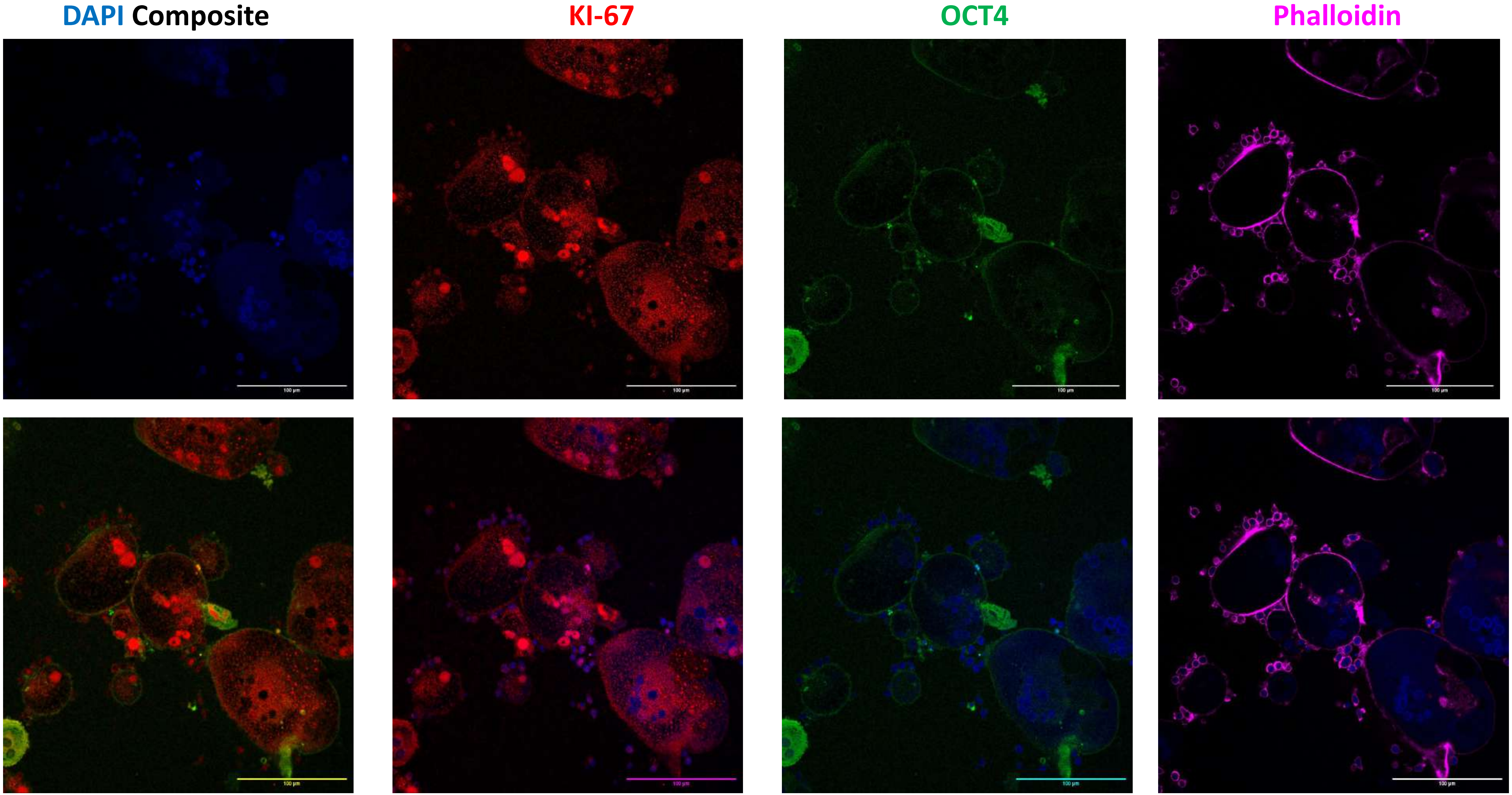

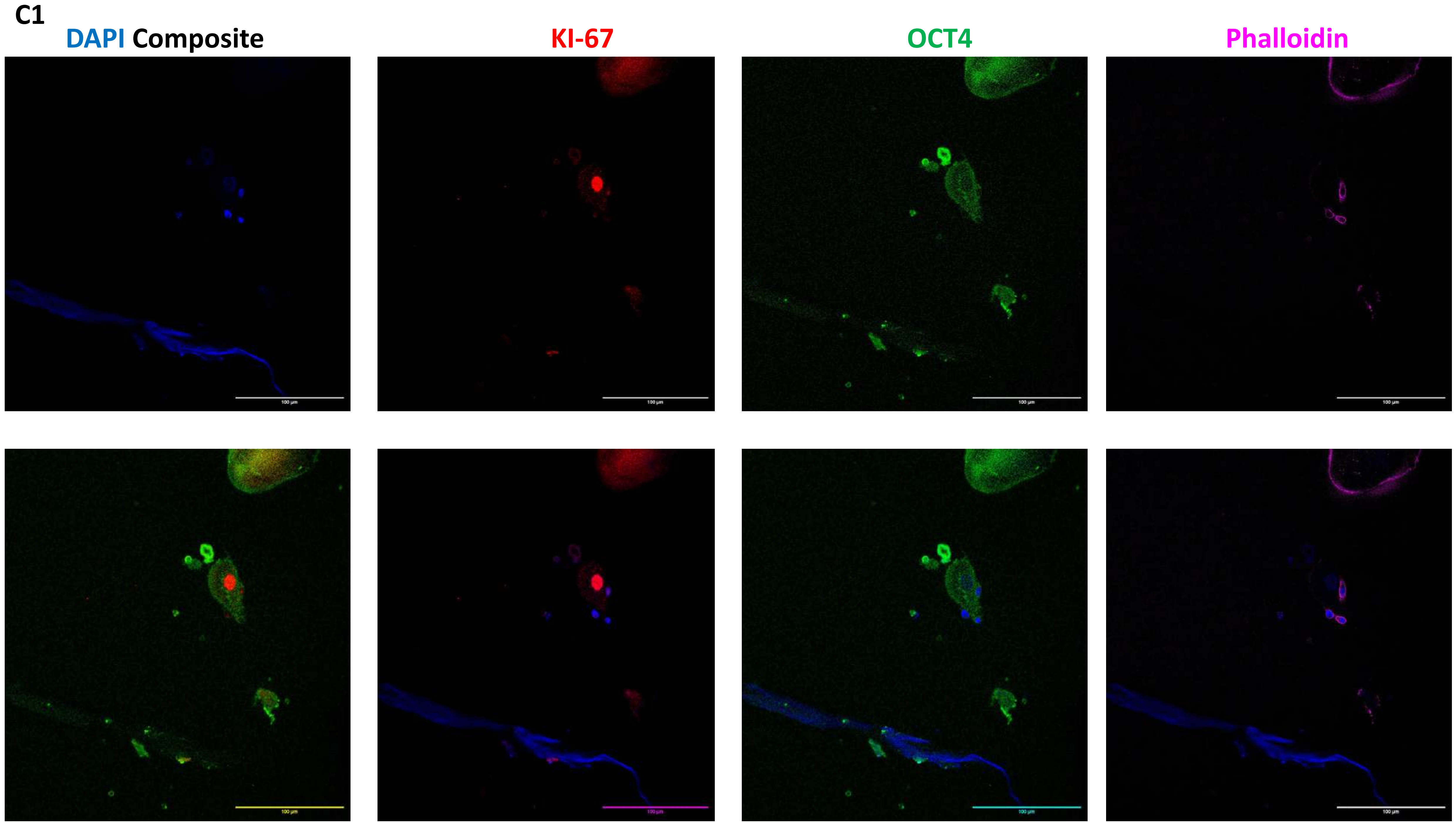
C1
DAPI Composite
KI-67
OCT4
Phalloidin
100 µm

# C1(160)

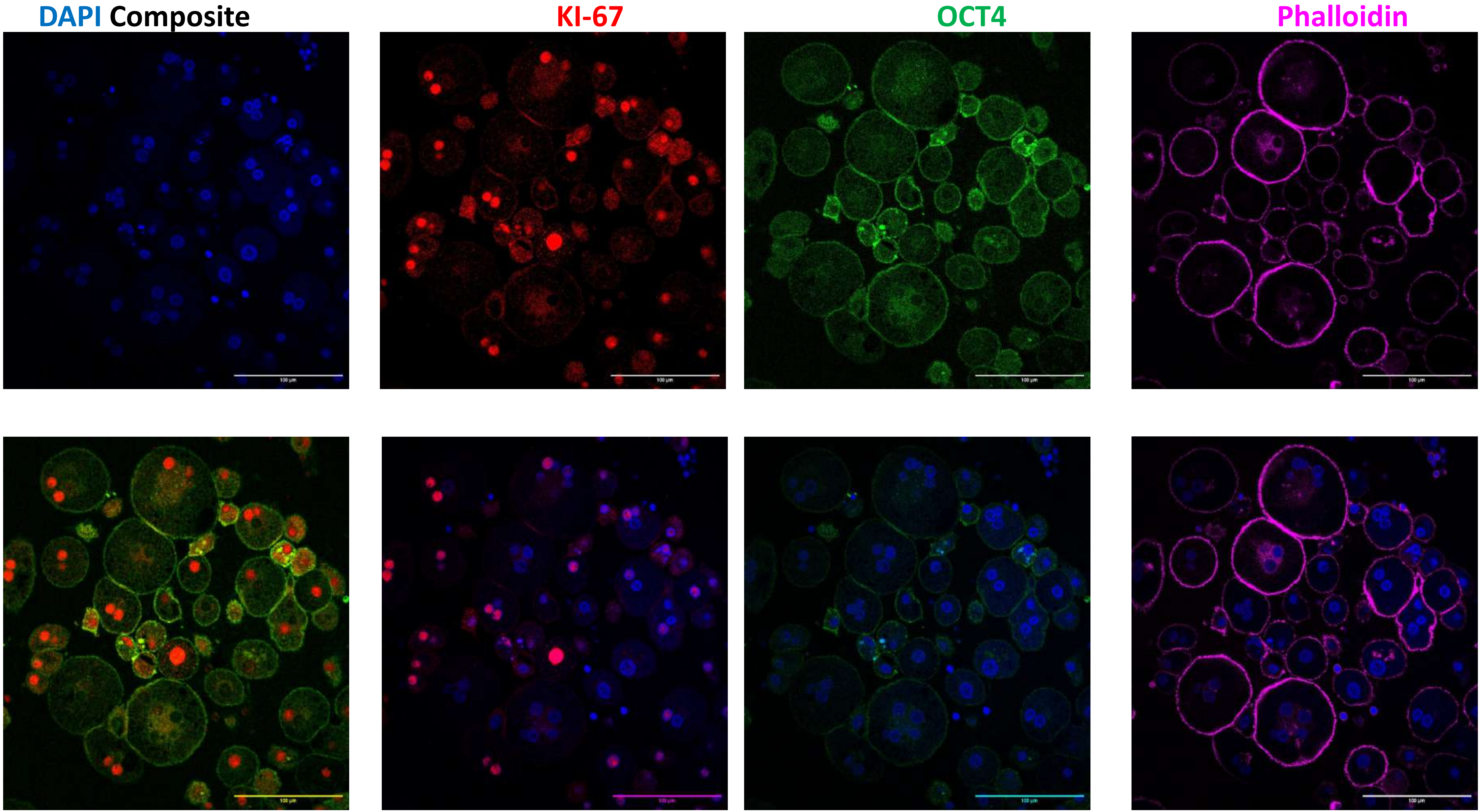

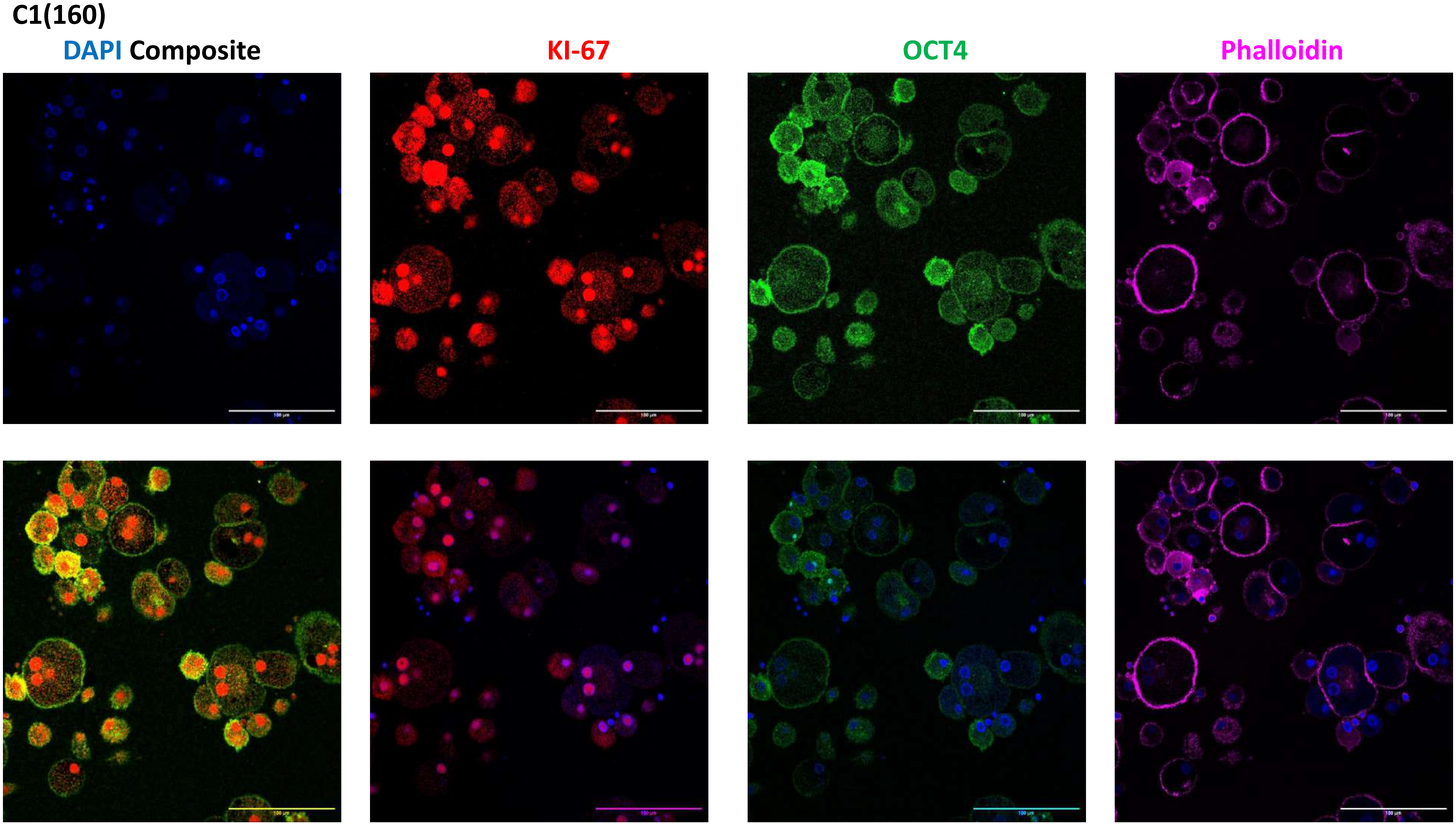
C1(160)
DAPI Composite
KI-67
OCT4
Phalloidin

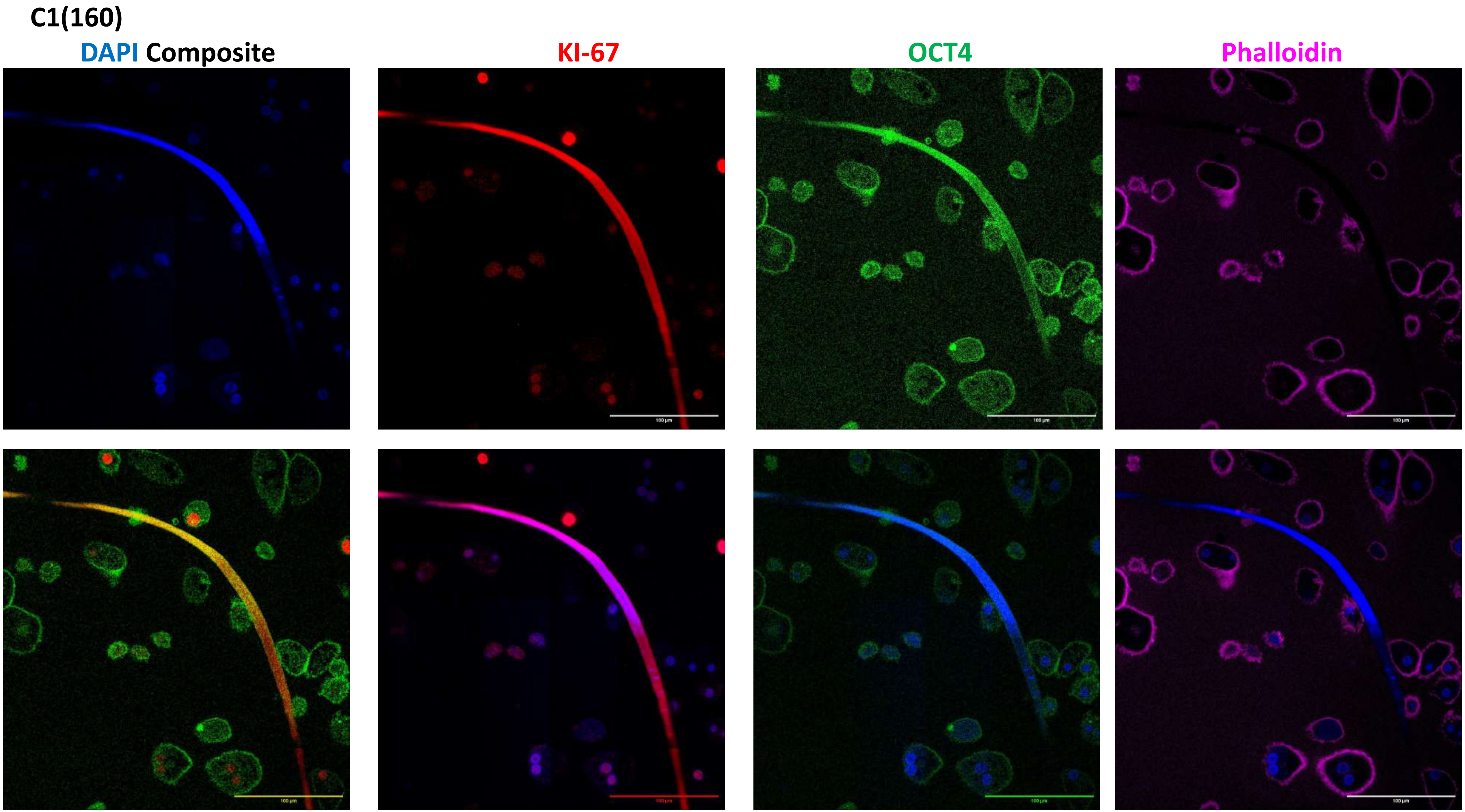
C1(160)
DAPI Composite
KI-67
OCT4
Phalloidin

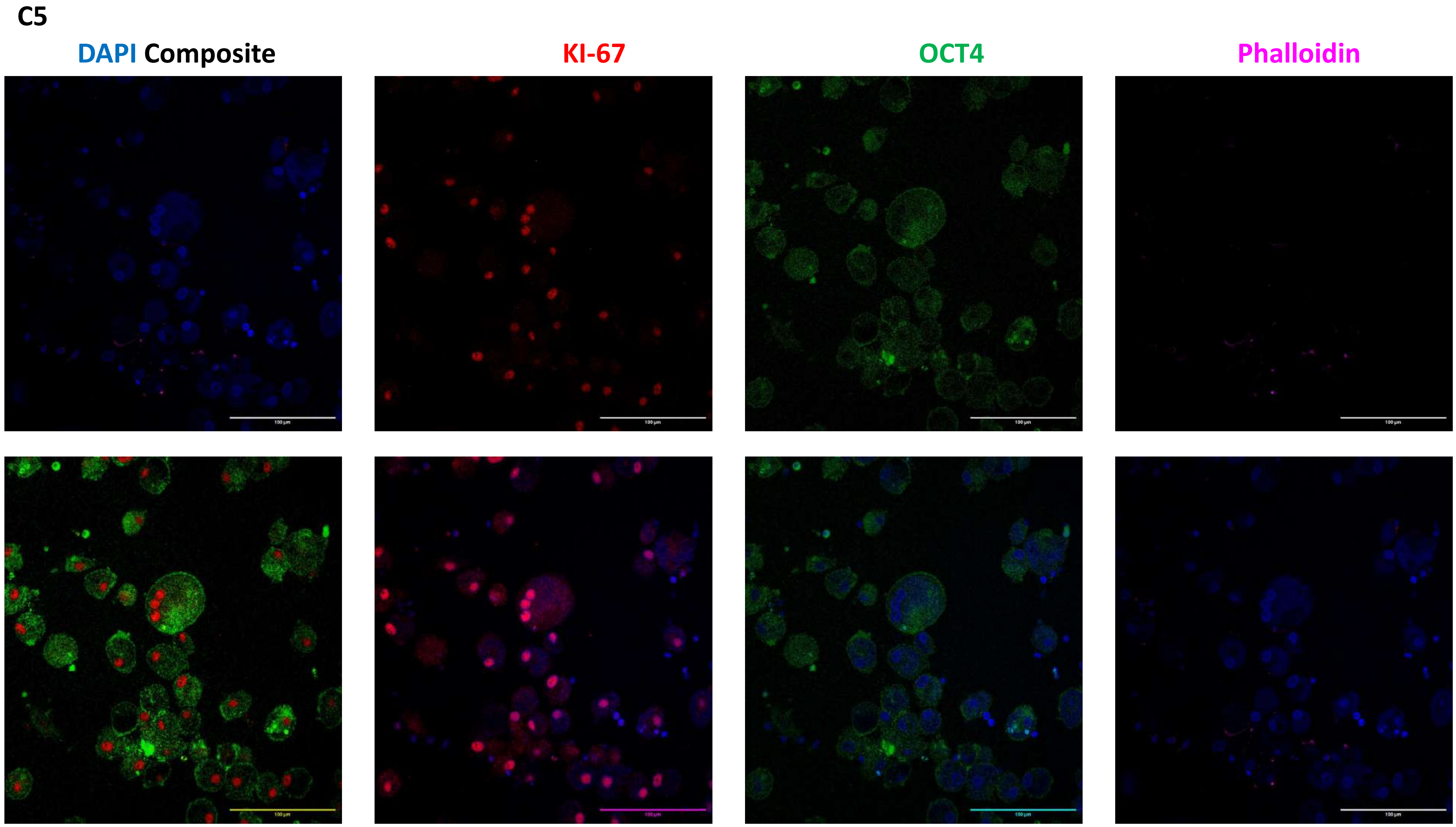
C5
DAPI Composite
KI-67
OCT4
Phalloidin

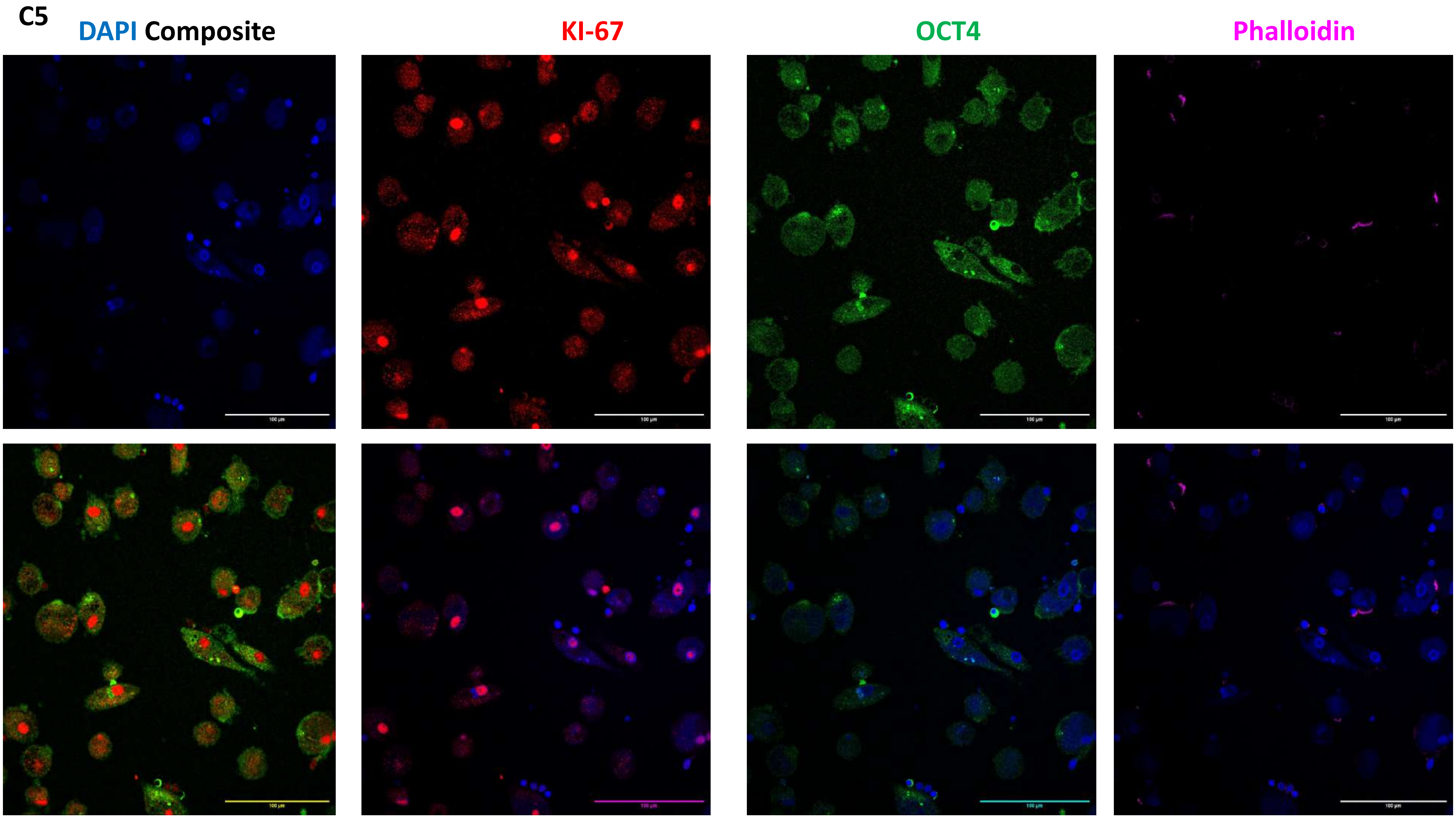
C5
DAPI Composite
KI-67
OCT4
Phalloidin

C5

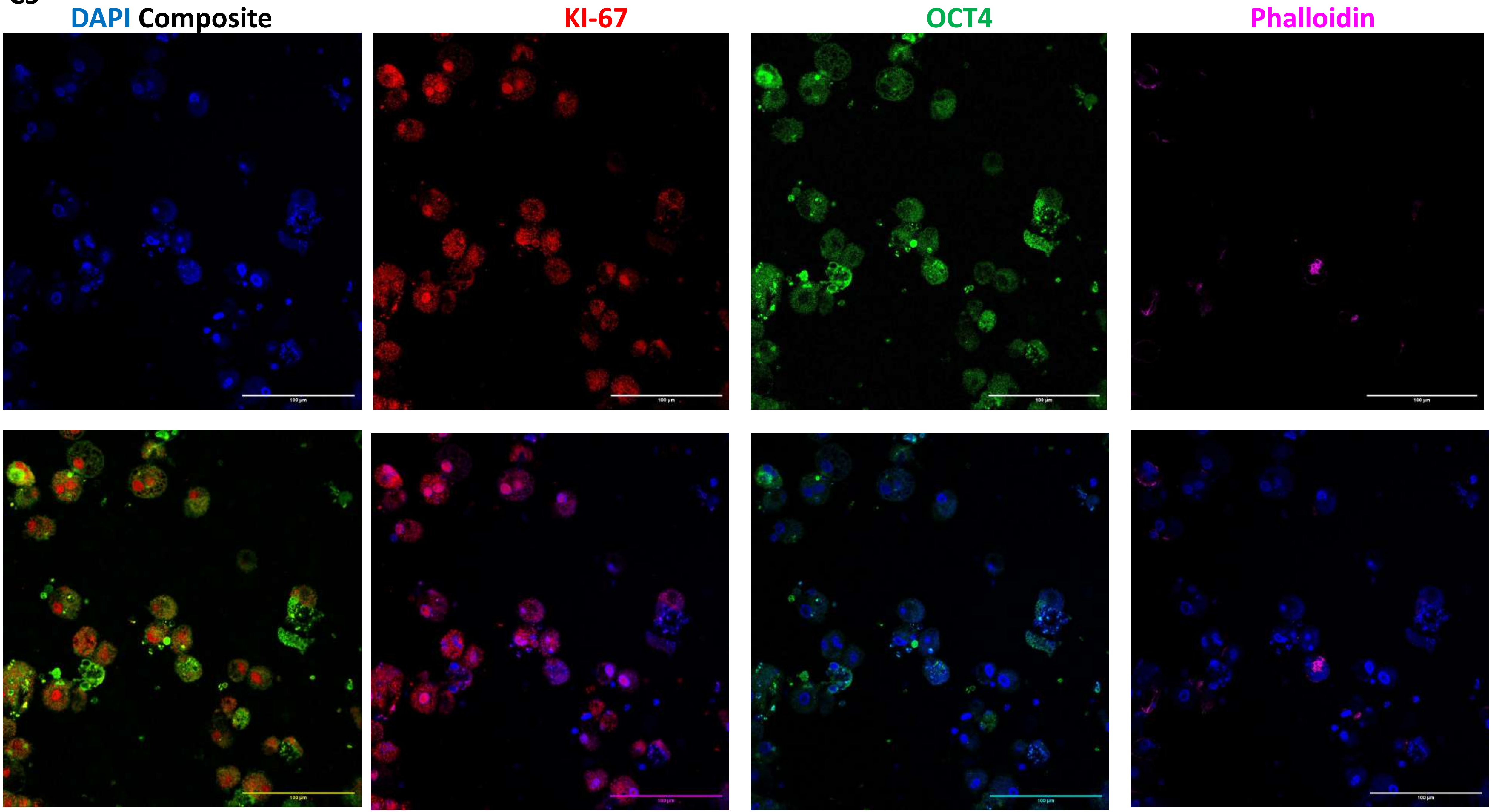

C5(160)

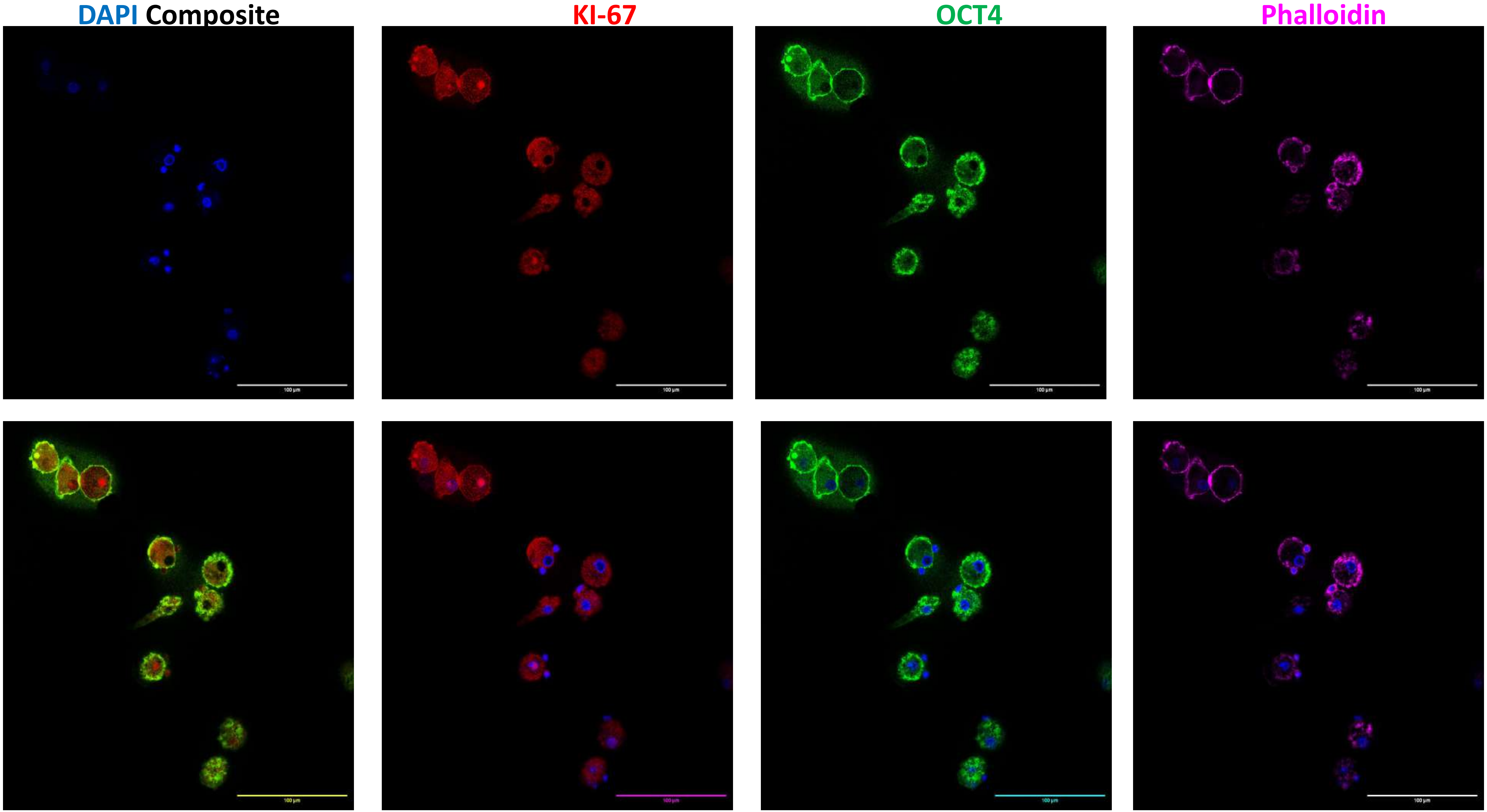

C5(160)

DAPI Composite KI-67 OCT4 Phalloidin

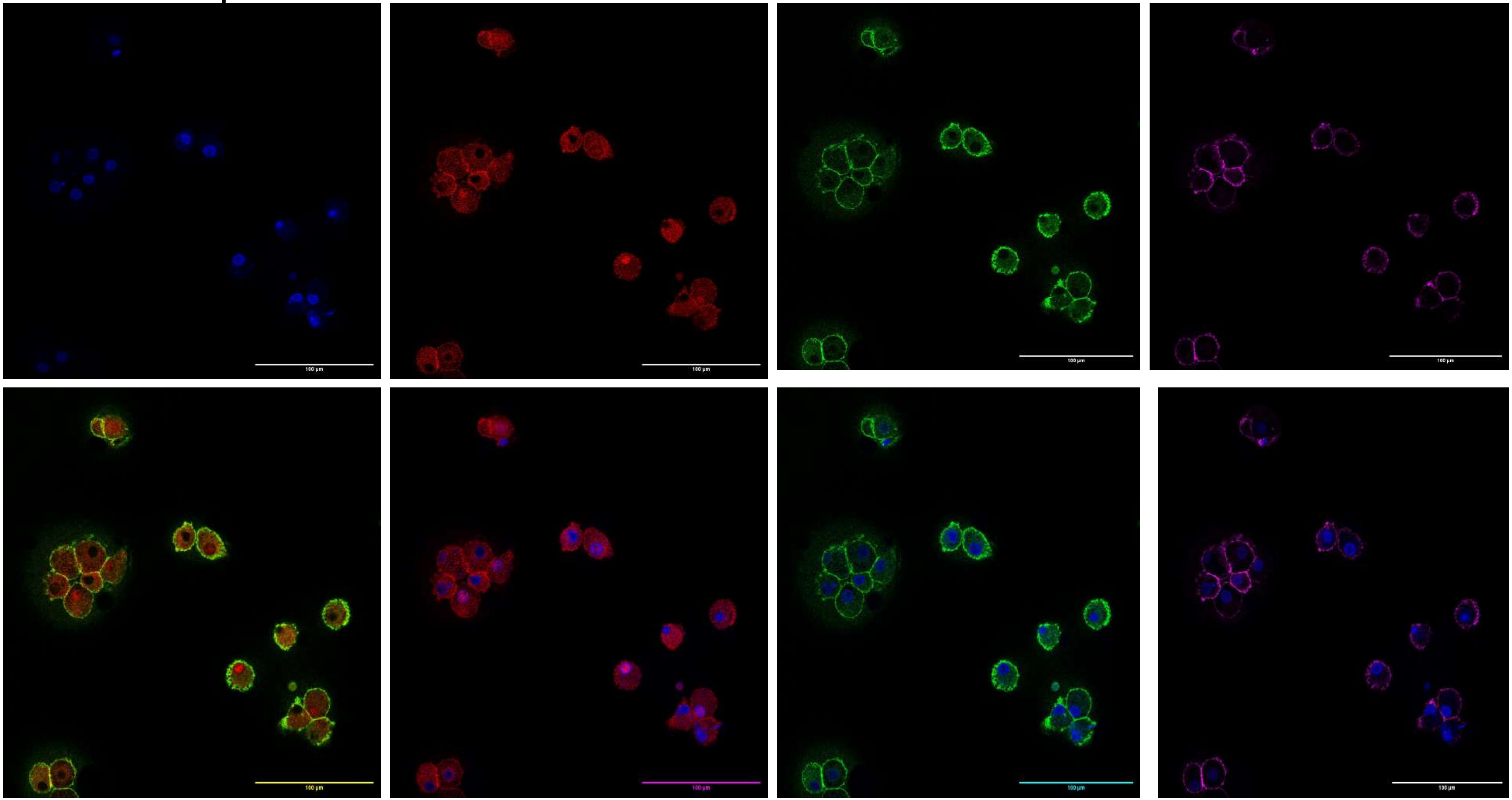

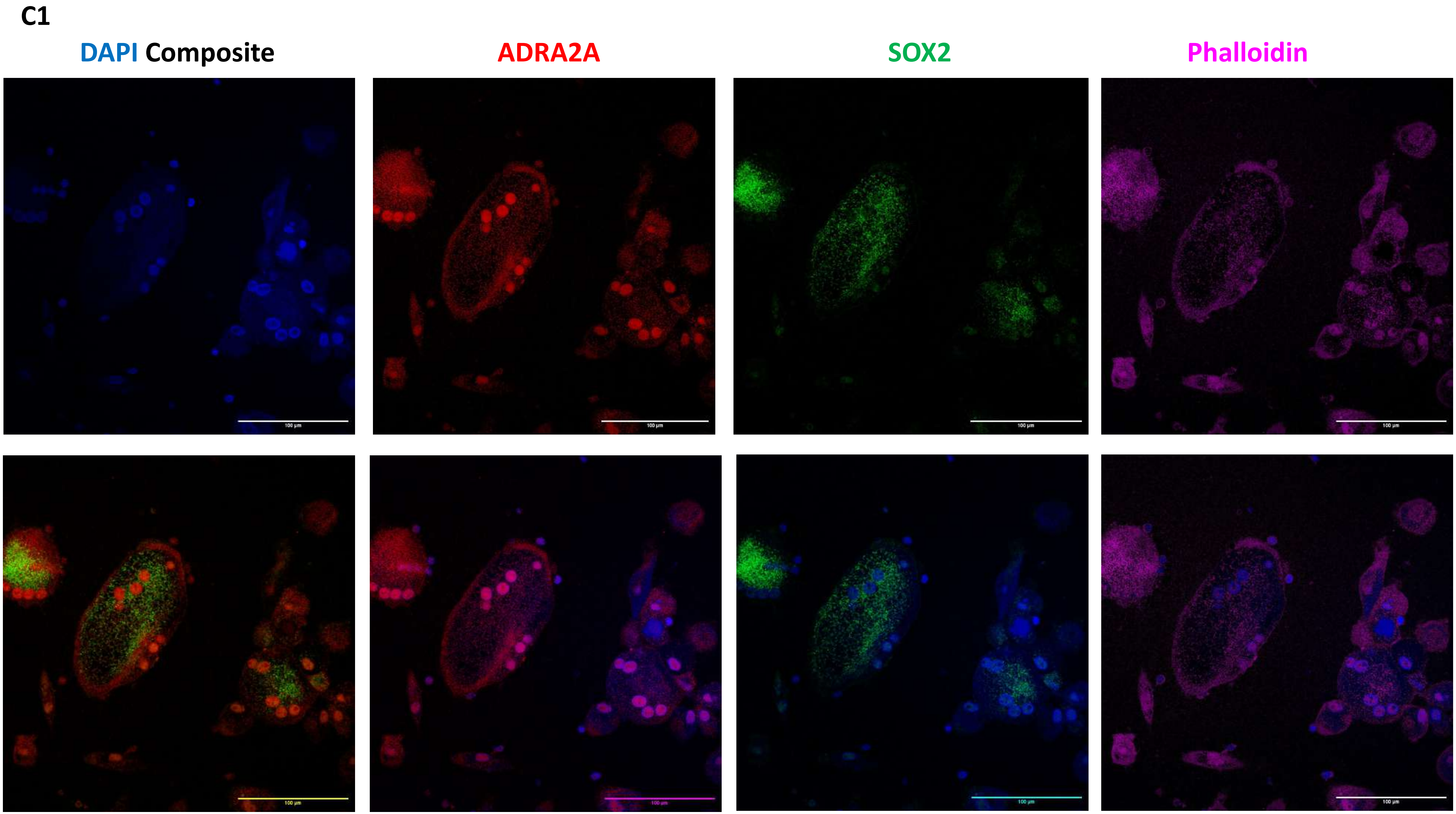
C1
DAPI Composite
ADRA2A
SOX2
Phalloidin
100 µm

C1

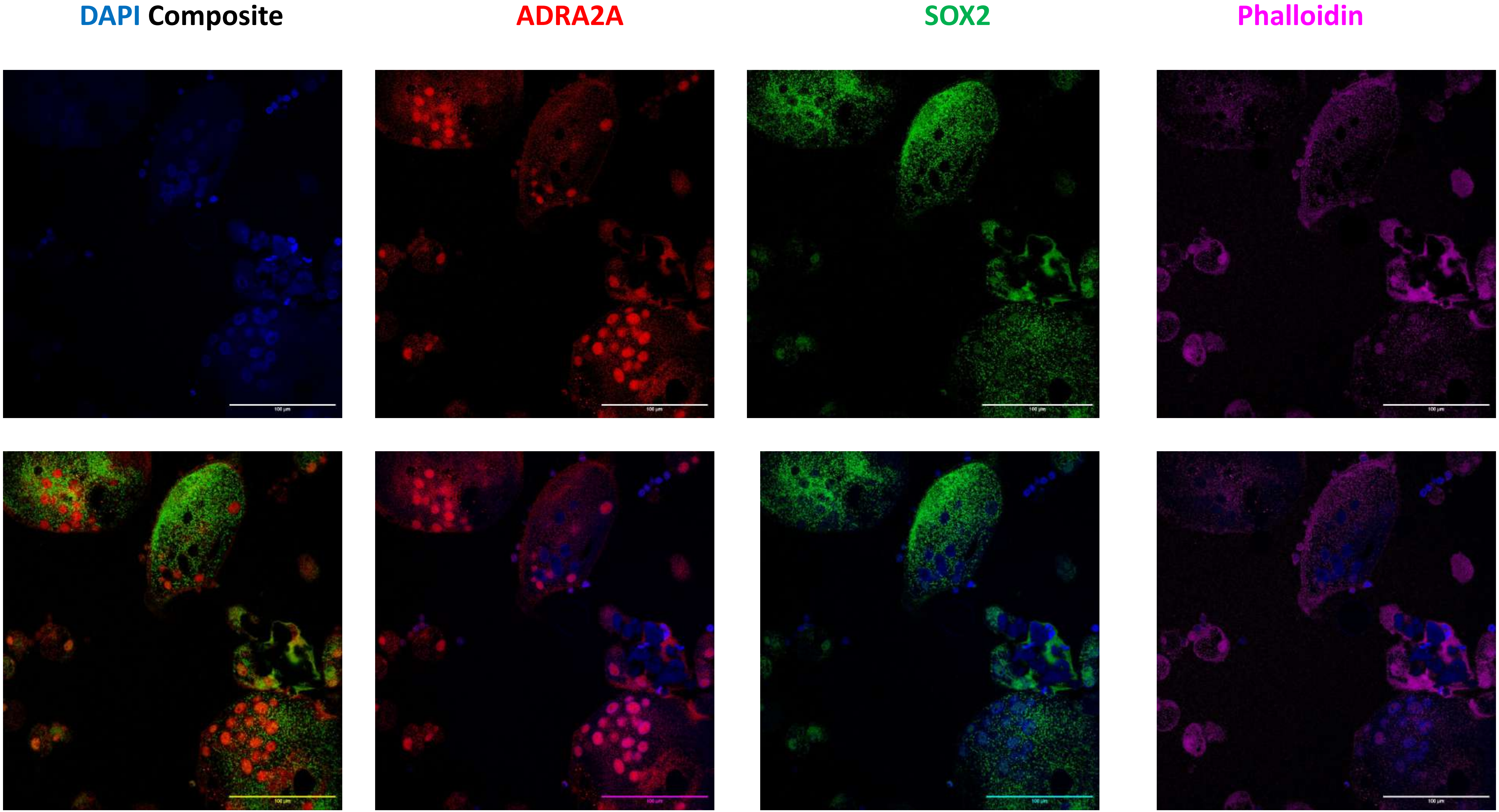

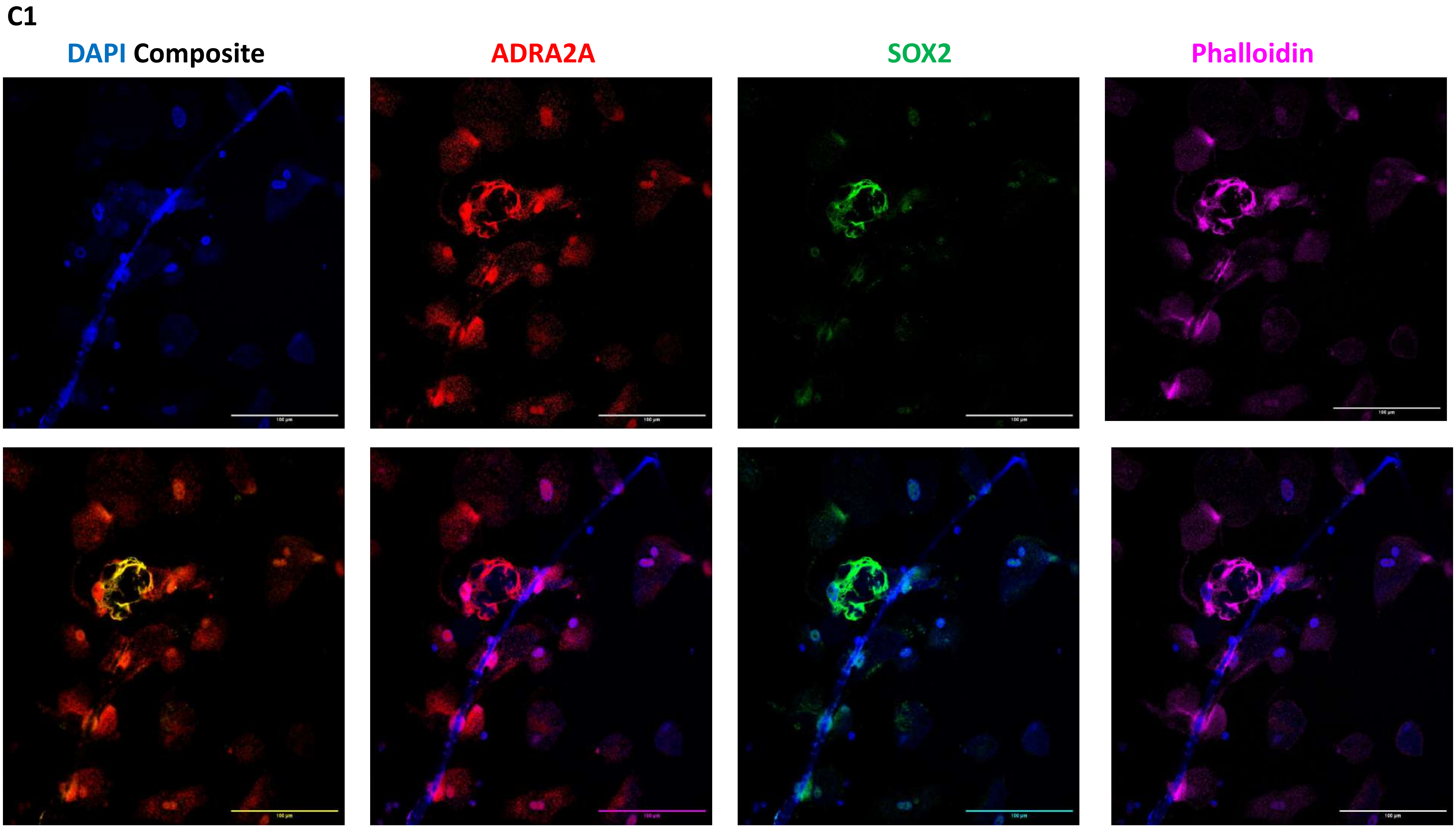
C1
DAPI Composite
ADRA2A
SOX2
Phalloidin

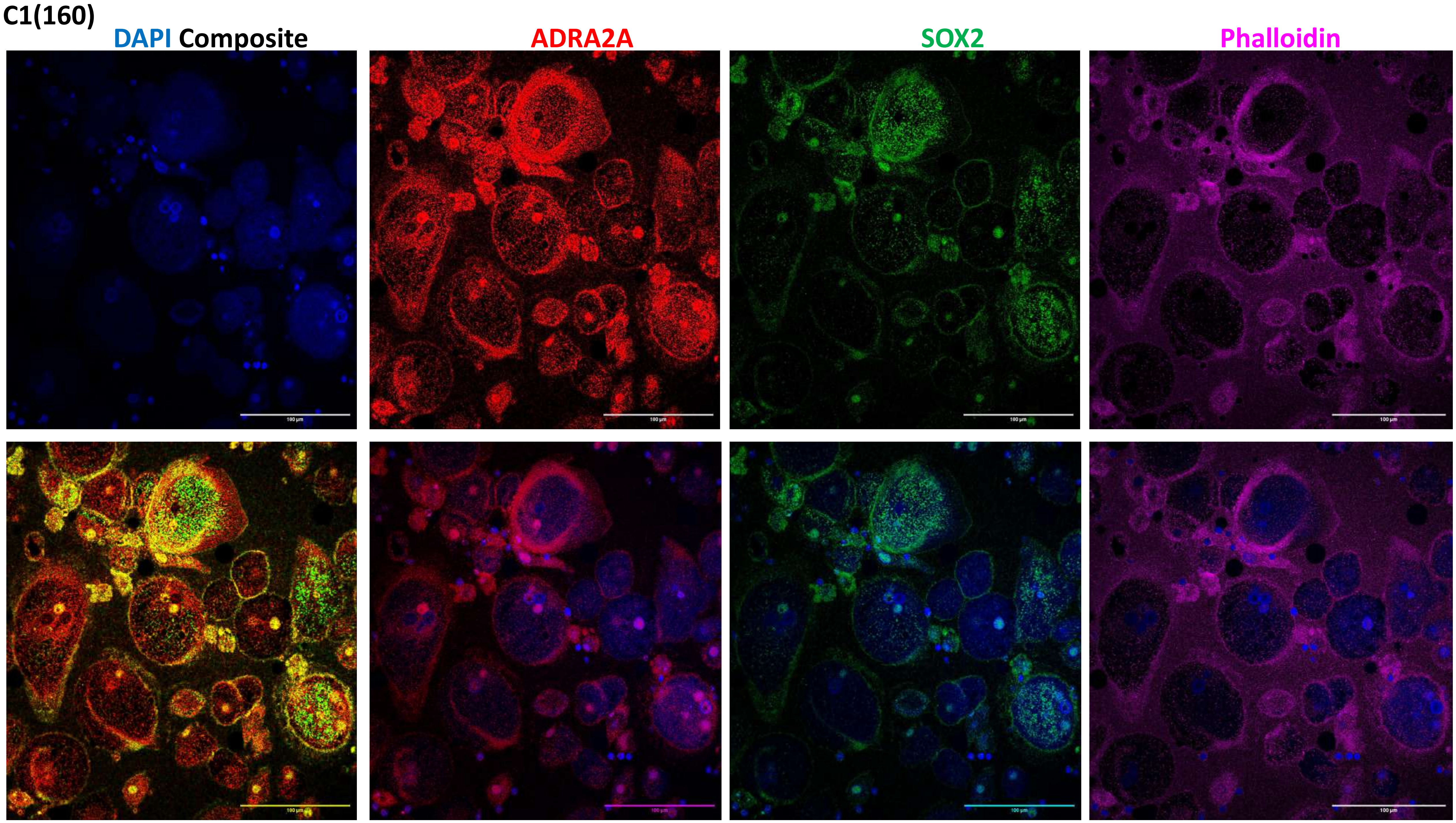
C1(160)
DAPI Composite
ADRA2A
SOX2
Phalloidin
100 µm

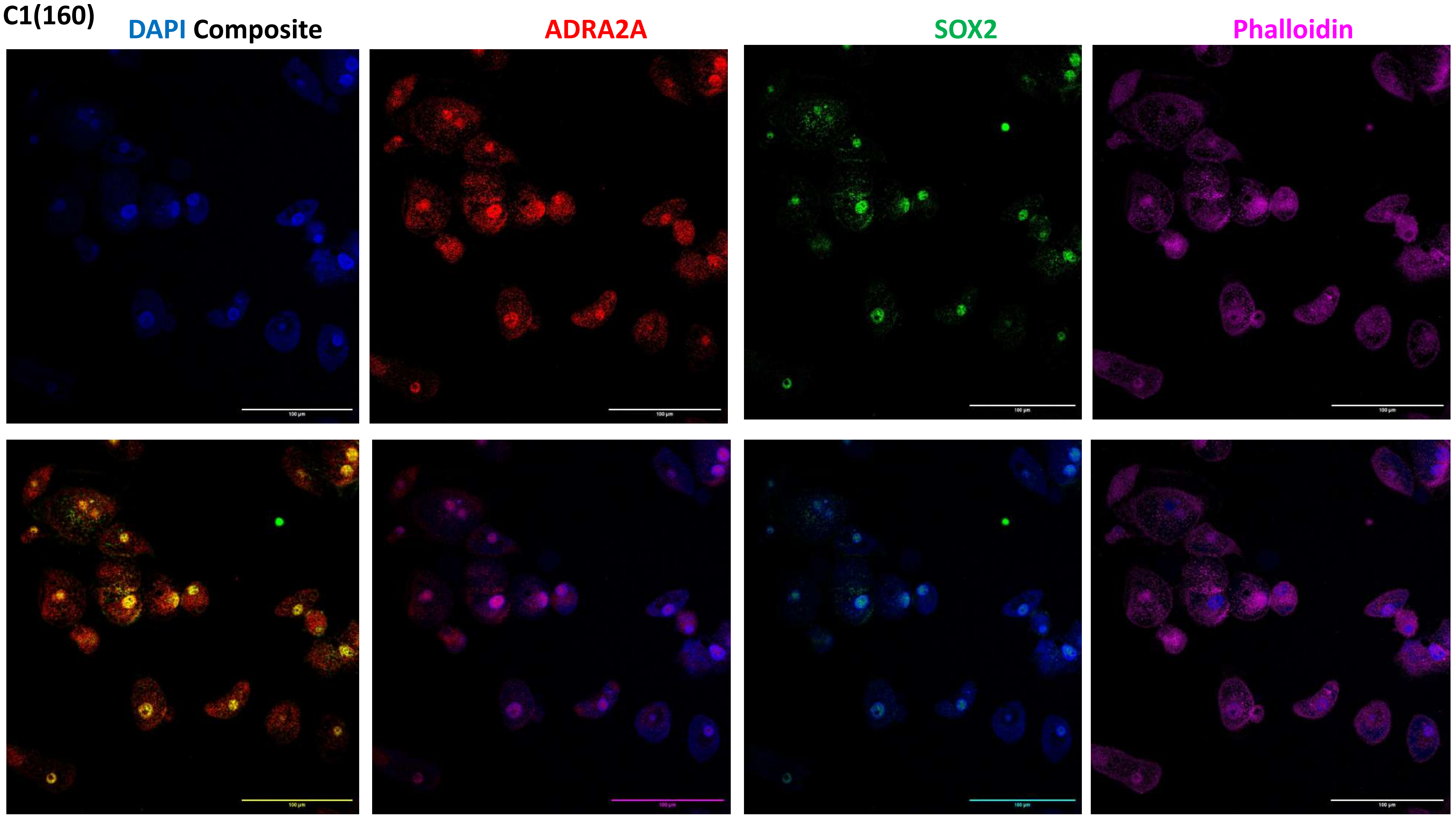
C1(160)
DAPI Composite
ADRA2A
SOX2
Phalloidin
100 µm

C1(160)

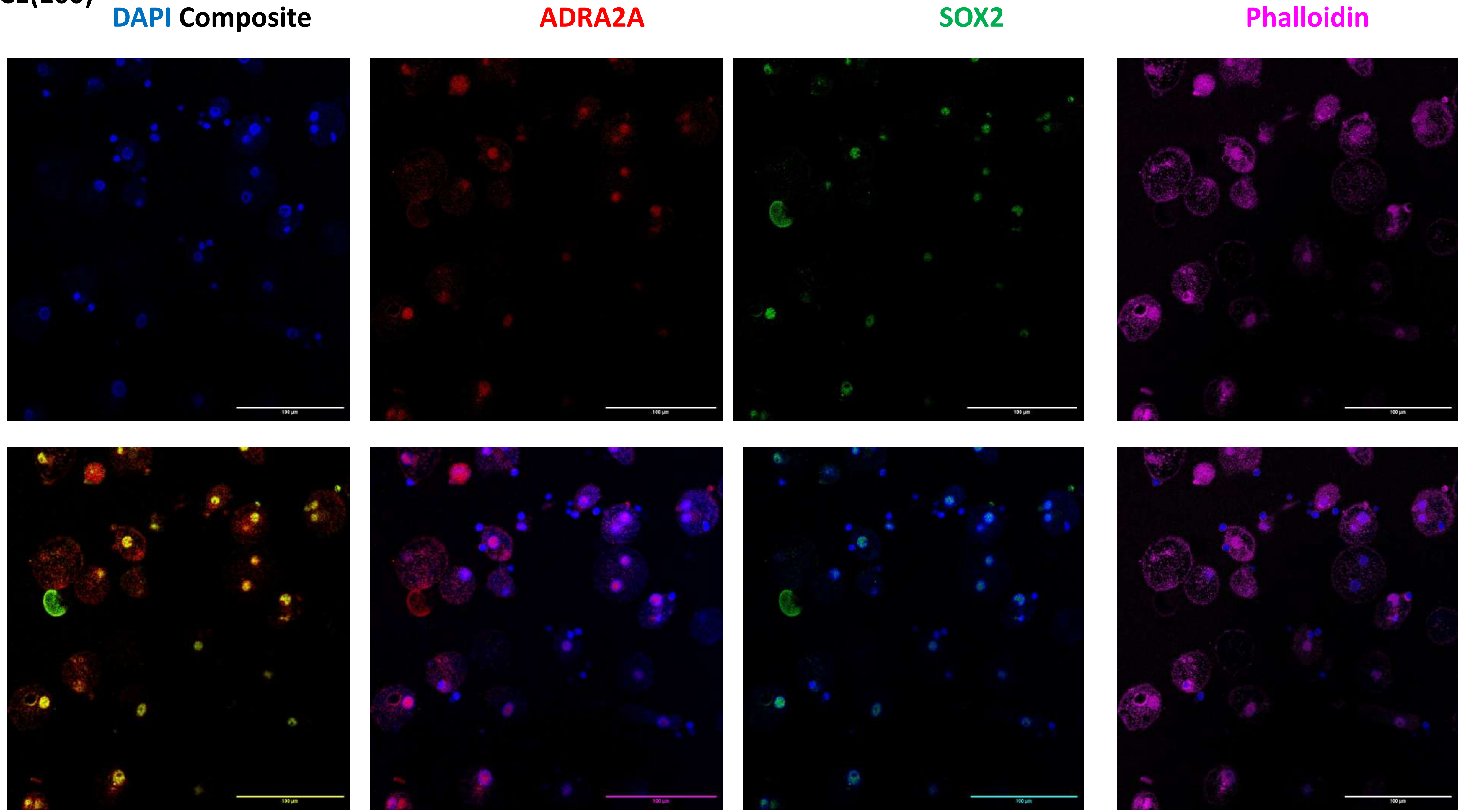

C5

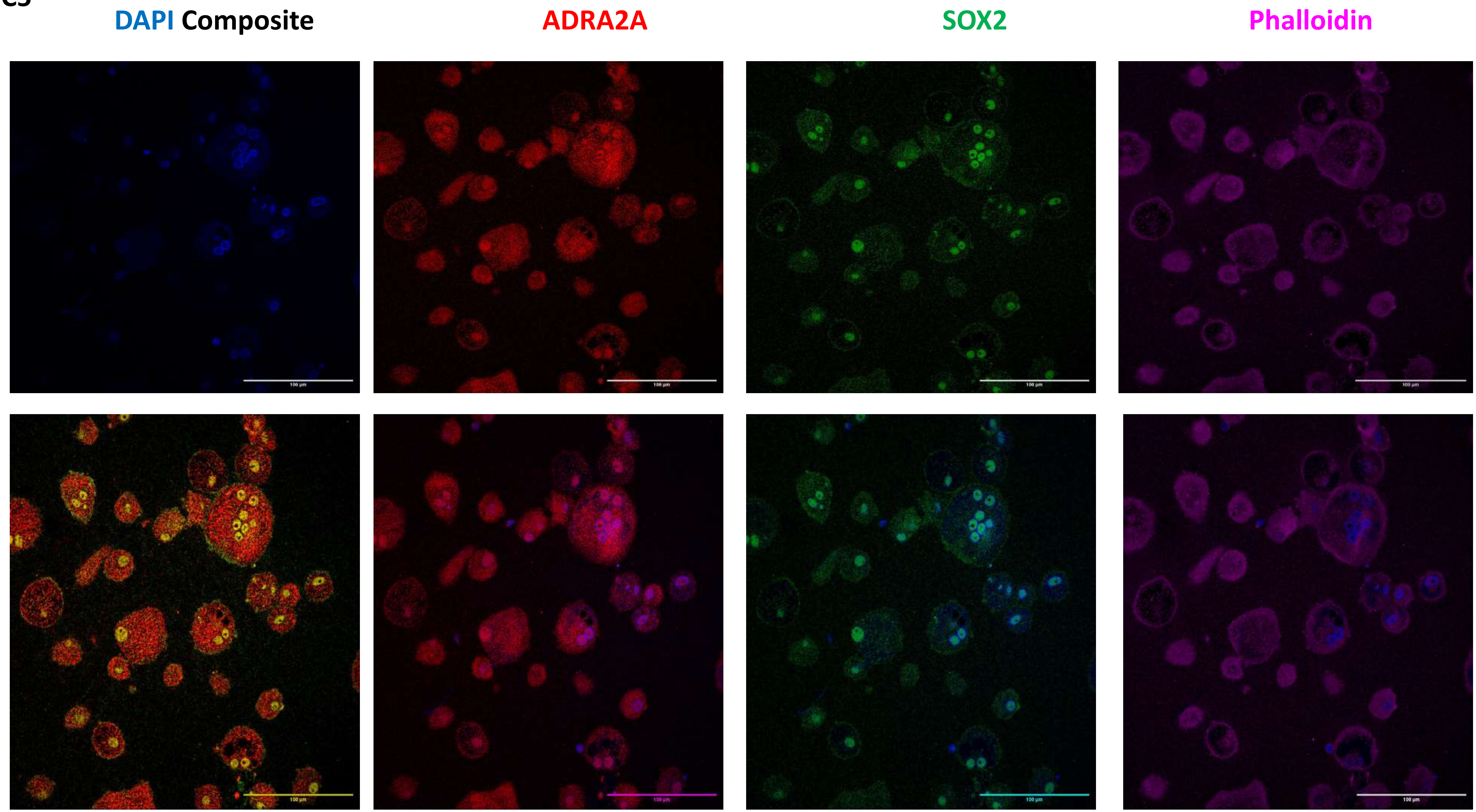

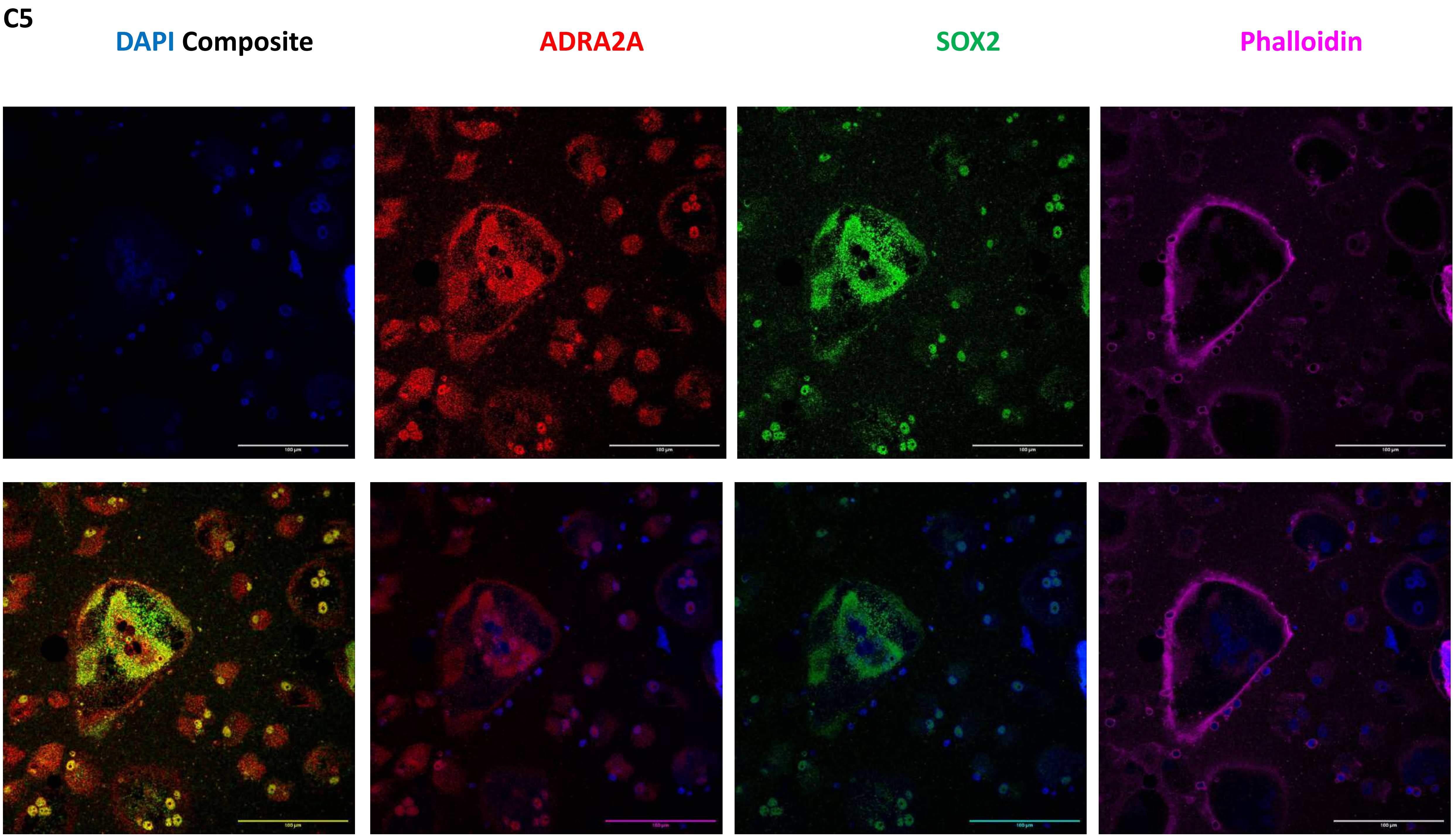
C5
DAPI Composite
ADRA2A
SOX2
Phalloidin

C5

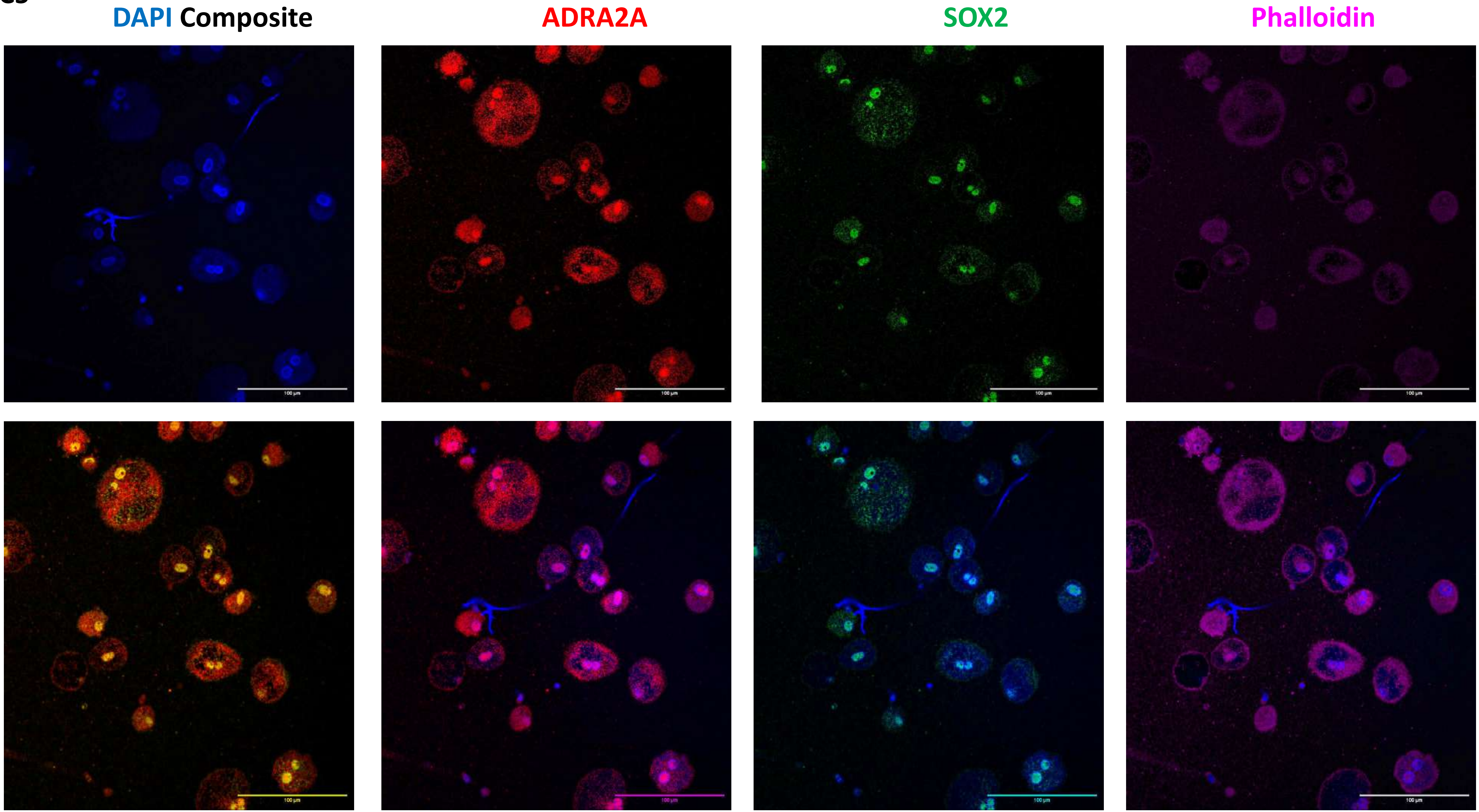

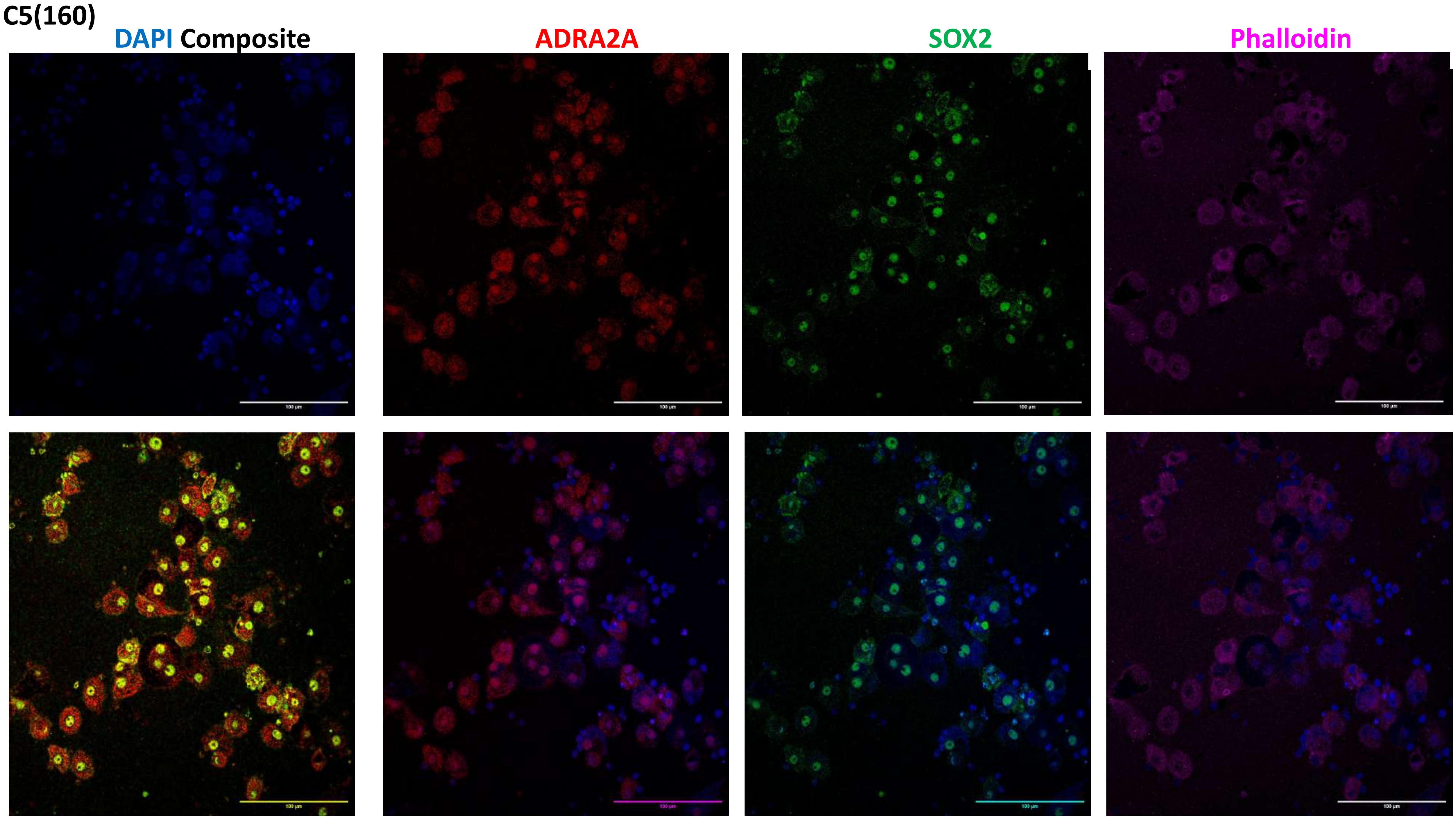
C5(160)
DAPI Composite
ADRA2A
SOX2
Phalloidin

C5(160)

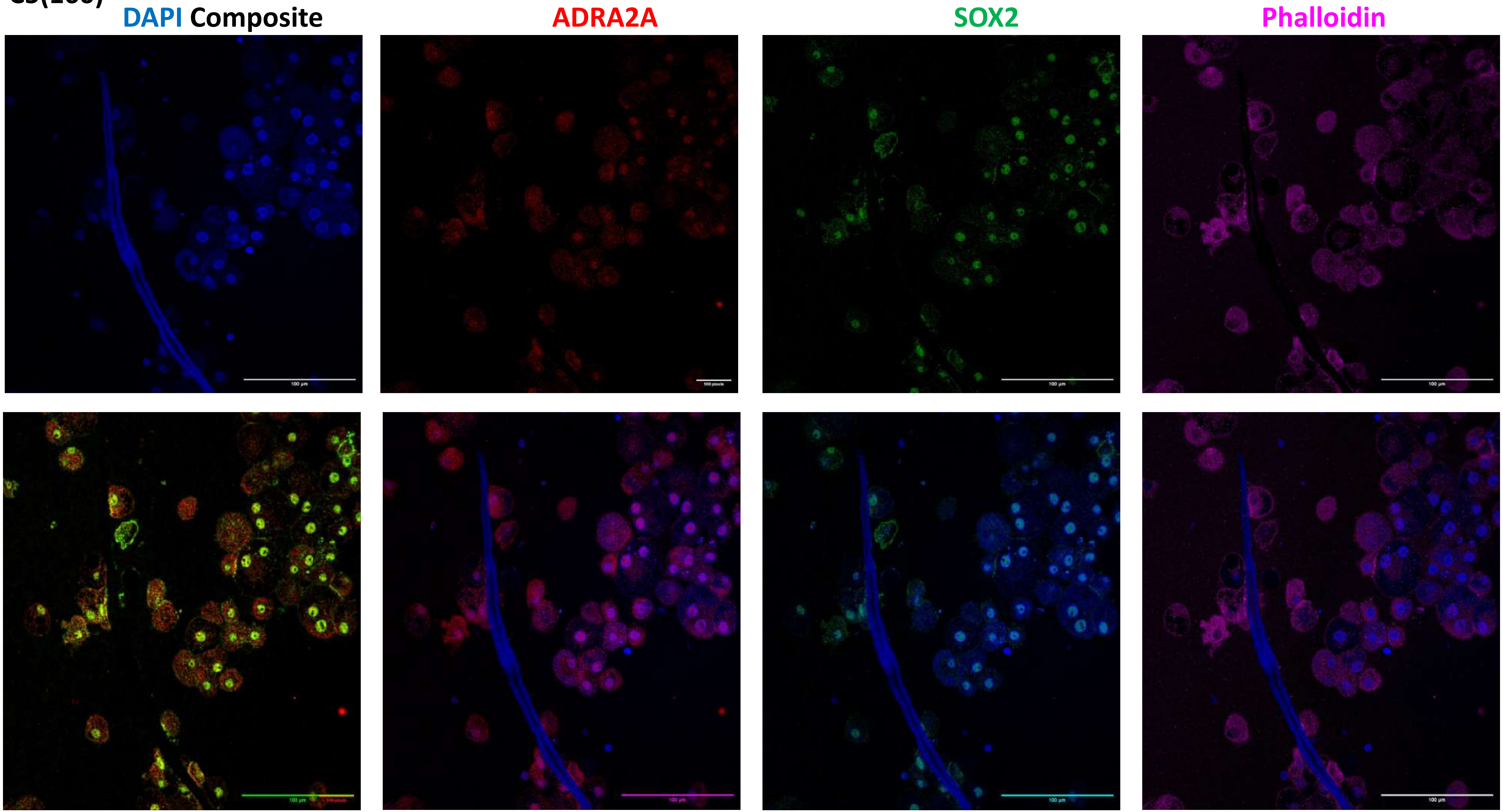

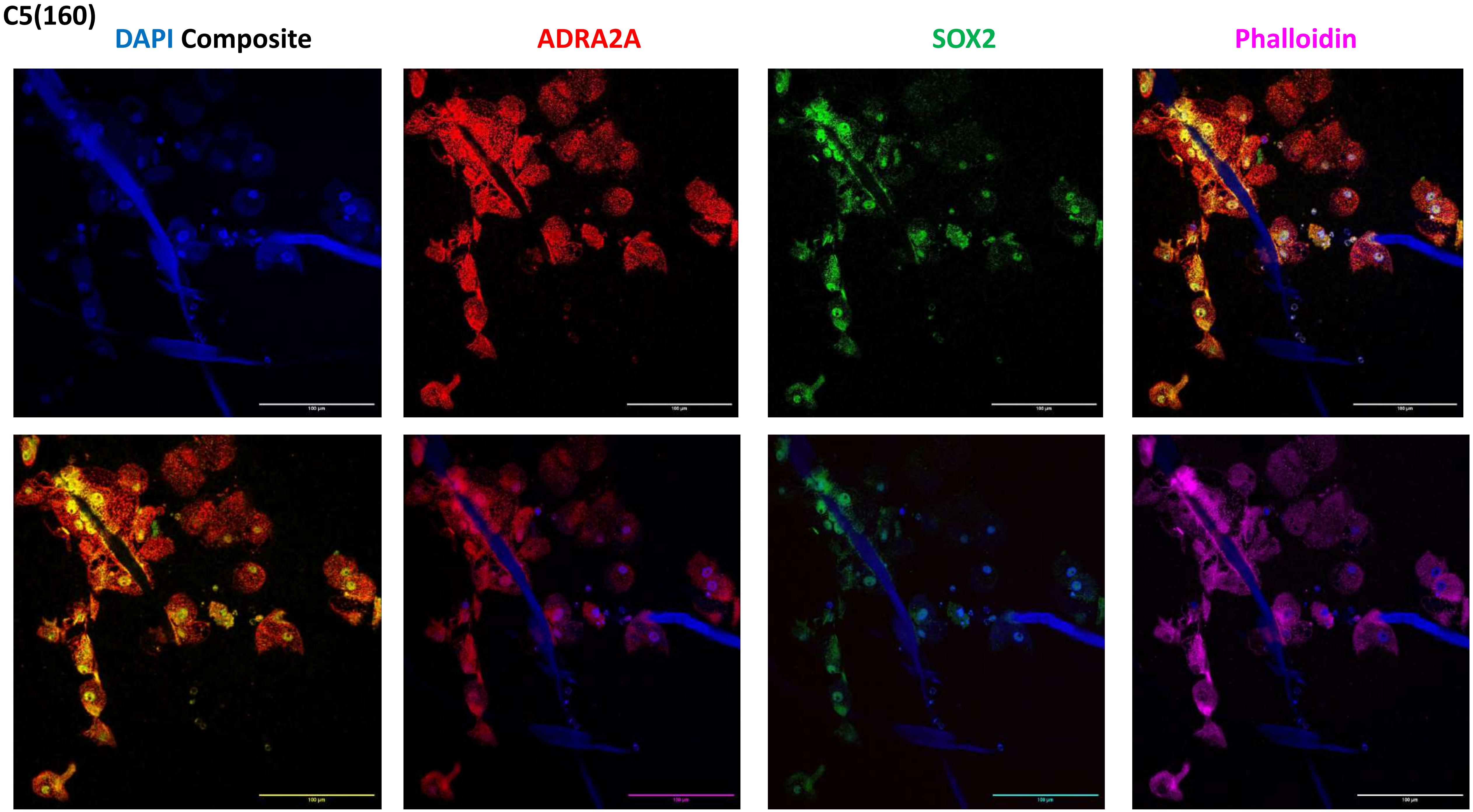
C5(160)
DAPI Composite
ADRA2A
SOX2
Phalloidin

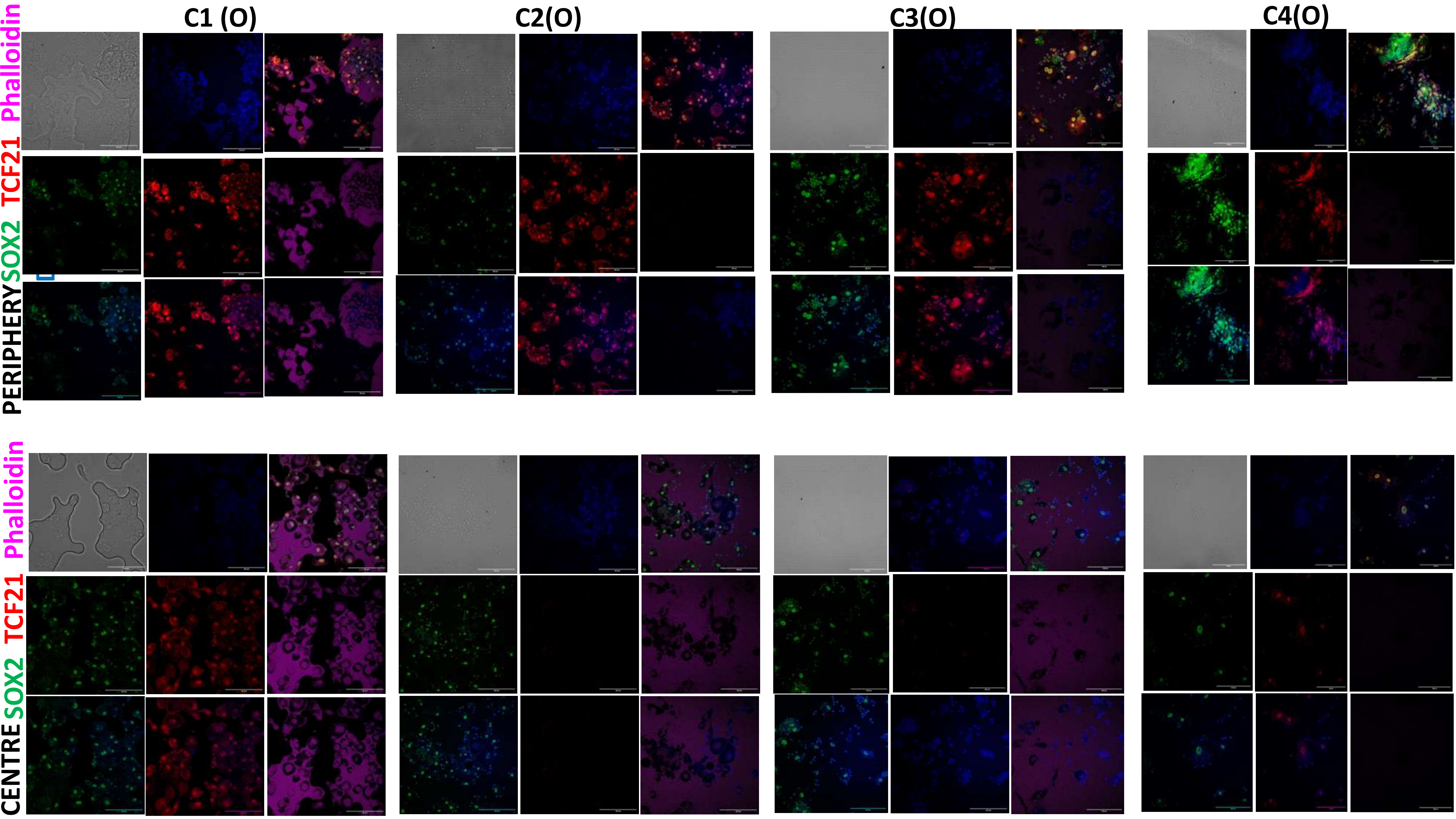
C1 (O)
C2(O)
C3(O)
C4(O)
PERIPHERY SOX2 TCF21 Phalloidin
CENTRE SOX2 TCF21 Phalloidin

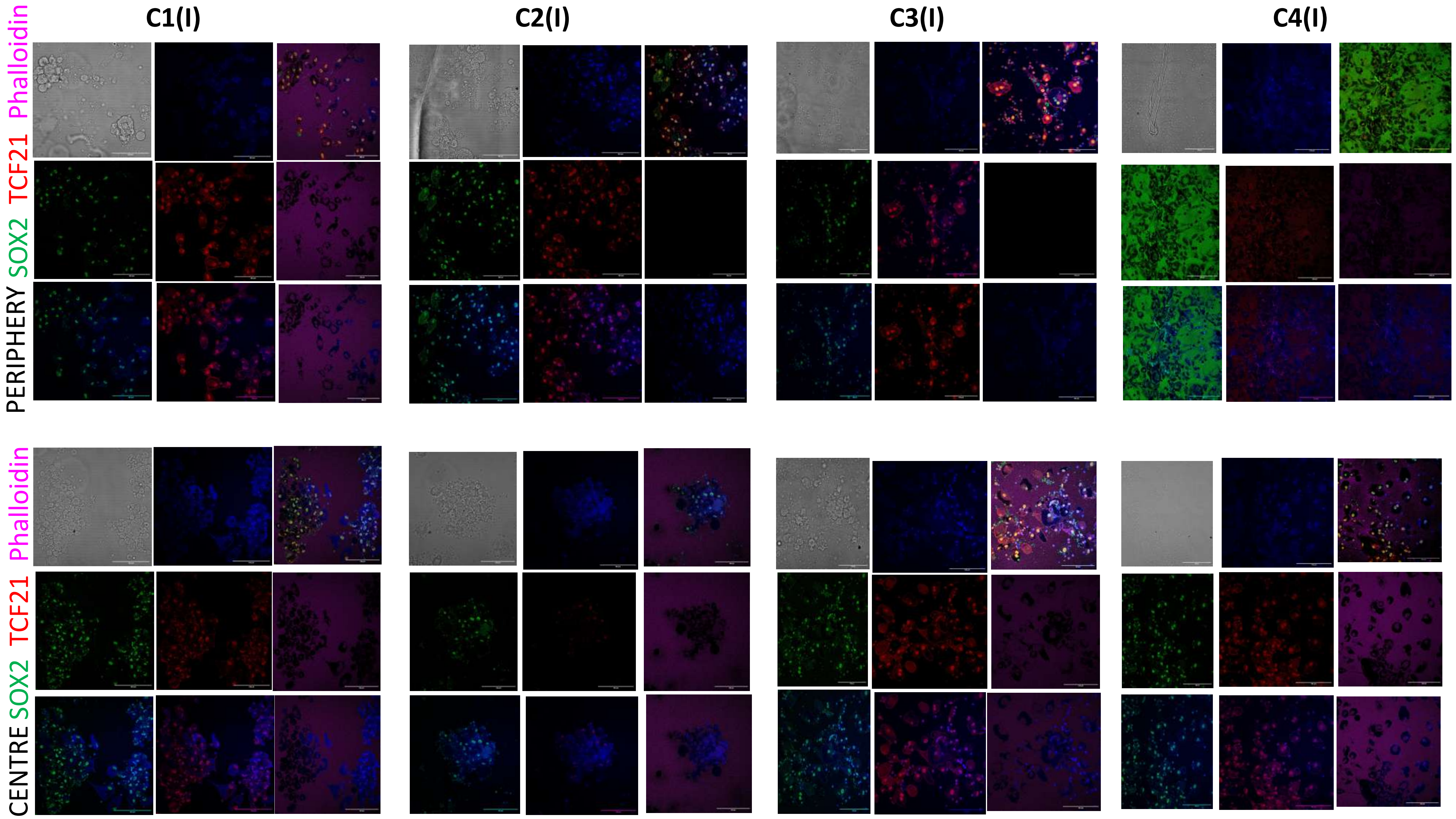
C1(I)
C2(I)
C3(I)
C4(I)
Phalloidin
TCF21
SOX2
PERIPHERY
Phalloidin
TCF21
SOX2
CENTRE

### Rotem for Complement- Coagulation

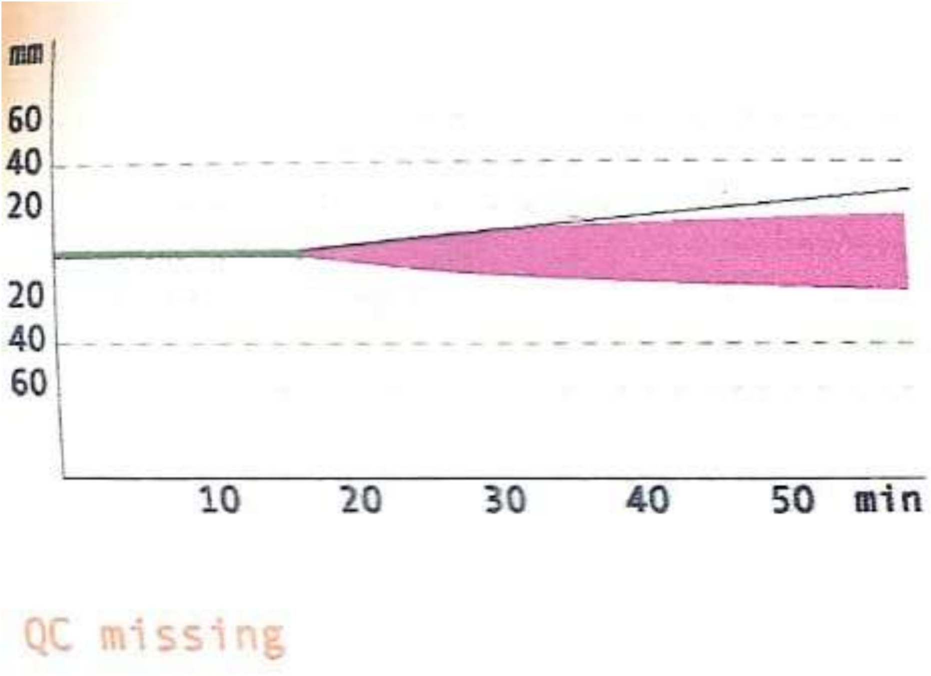


1 EXTEM [default]

1:1 2:1

RT: 01:00:00 ST: 2023-07-05T08:15:41

| | | | | | | | | |
|---|---|---|---|---|---|---|---|---|
| CT | : | | 1107 | s | [ 38 | - | 79] | ▲ |
| CFT | : | | | s | [ 34 | - | 159] | |
| α | : | | 9 | ° | [ 63 | - | 83] | ▼ |
| A10 | : | | 8 | mm | [ 43 | - | 65] | ▼ |
| A20 | : | | 12 | mm | [ 50 | - | 71] | ▼ |
| MCF | : | * | 16 | mm | [ 50 | - | 72] | |
| ML | : | * | 0 | % | [ 0 | - | 15] | |

**C1 Day 2**

### Rotem for Complement- Coagulation

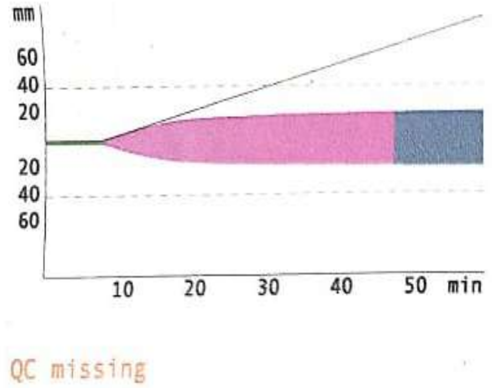


3 EXTEM [default] EP, 1
1:EP 2:1
RT: 01:00:00 ST: 2023-09-20T06:43:52

| | | | | | | | |
|---|---|---|---|---|---|---|---|
| CT | : | | 484 | s | [ 38 - | 79] | ▲ |
| CFT | : | | 2391 | s | [ 34 - | 159] | ▲ |
| α | : | | 24 | ° | [ 63 - | 83] | ▼ |
| A10 | : | | 15 | mm | [ 43 - | 65] | ▼ |
| A20 | : | | 18 | mm | [ 50 - | 71] | ▼ |
| MCF | : | * | 21 | mm | [ 50 - | 72] | |
| ML | : | * | 0 | % | [ 0 - | 15] | |

**C1 Day 3**

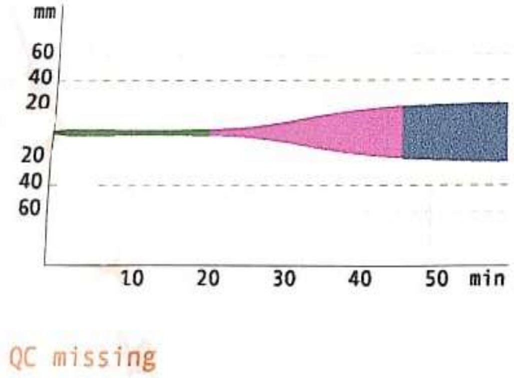


4 INTEM [default] EP, 1
1:EP 2:1
RT: 01:00:00 ST: 2023-09-20T06:45:01

| | | | | | | | |
|---|---|---|---|---|---|---|---|
| CT | : | | 849 | s | [ 100 - | 240] | ▲ |
| CFT | : | | 1923 | s | [ 30 - | 110] | ▲ |
| α | : | * | 8 | ° | [ 70 - | 83] | |
| A10 | : | | 3 | mm | [ 44 - | 66] | ▼ |
| A20 | : | | 12 | mm | [ 50 - | 71] | ▼ |
| MCF | : | * | 23 | mm | [ 50 - | 72] | |
| ML | : | * | 0 | % | [ 0 - | 15] | |

**C1 Day 3**

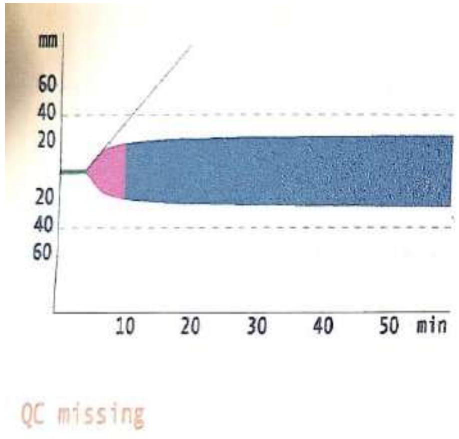


1 EXTEM [default] EP+ 2, 2
1:EP + 2 2:2
RT: 01:00:01 ST: 2023-09-20T06:52:39

| | | | | | | | |
|---|---|---|---|---|---|---|---|
| CT | : | | 249 | s | [ 38 - | 79] | ▲ |
| CFT | : | | 361 | s | [ 34 - | 159] | ▲ |
| α | : | | 55 | ° | [ 63 - | 83] | ▼ |
| A10 | : | | 22 | mm | [ 43 - | 65] | ▼ |
| A20 | : | | 24 | mm | [ 50 - | 71] | ▼ |
| MCF | : | | 25 | mm | [ 50 - | 72] | ▼ |
| ML | : | * | 0 | % | [ 0 - | 15] | |

**C2 Day 3**

Figure: The Rotem test reveals a notable upregulation of the complement cascade in (EDHA) P on Day 3. Moreover, this upregulation is even more pronounced in EDHA P+2 at Day 3.

### Rotem for Complement- Coagulation

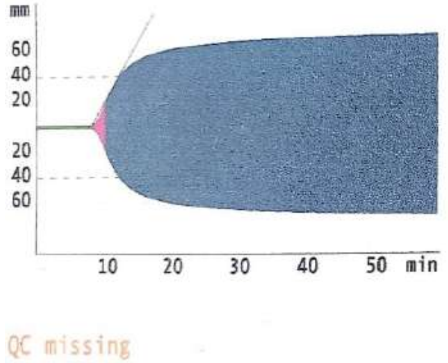


1 INTEM [default] 4, 4
1:4 2:

RT: 01:00:00 ST: 2023-09-27T08:42:56

| | | | | | |
|---|---|---|---|---|---|
| CT | : | 510 | s | [ 100 - 240] | ▲ |
| CFT | : | 121 | s | [ 30 - 110] | ▲ |
| α | : | 68 | ° | [ 70 - 83] | ▼ |
| A10 | : | 59 | mm | [ 44 - 66] | |
| A20 | : | 66 | mm | [ 50 - 71] | |
| MCF | : * | 71 | mm | [ 50 - 72] | |
| ML | : * | 0 | % | [ 0 - 15] | |

**C1 Day 3**

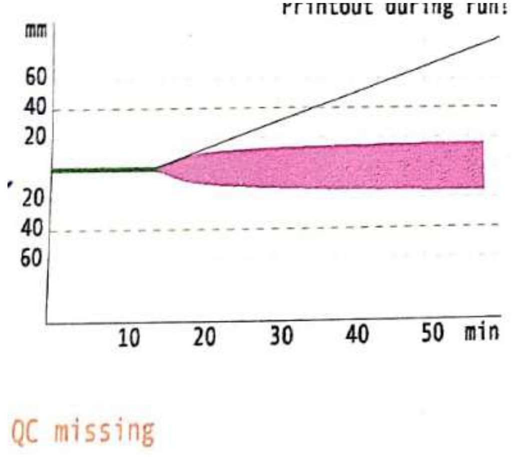


3 INTEM [default] 6, 6
1:6 2:6

RT: 00:58:35 ST: 2023-09-27T08:44:51

| | | | | | |
|---|---|---|---|---|---|
| CT | : | 864 | s | [ 100 - 240] | ▲ |
| CFT | : | | s | [ 30 - 110] | |
| α | : | 26 | ° | [ 70 - 83] | ▼ |
| A10 | : | 12 | mm | [ 44 - 66] | ▼ |
| A20 | : | 14 | mm | [ 50 - 71] | ▼ |
| MCF | : * | 16 | mm | [ 50 - 72] | |
| ML | : * | 0 | % | [ 0 - 15] | |

**C1 Day2**

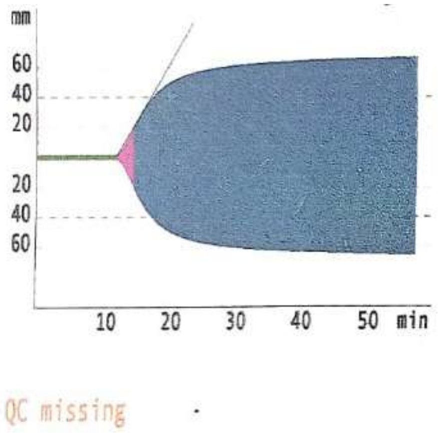


4 INTEM [default] 3, 3
1:3 2:3

RT: 00:58:21 ST: 2023-09-27T07:37:00

| | | | | | |
|---|---|---|---|---|---|
| CT | : | 732 | s | [ 100 - 240] | ▲ |
| CFT | : | 149 | s | [ 30 - 110] | ▲ |
| α | : | 63 | ° | [ 70 - 83] | ▼ |
| A10 | : | 54 | mm | [ 44 - 66] | |
| A20 | : | 62 | mm | [ 50 - 71] | |
| MCF | : * | 66 | mm | [ 50 - 72] | |
| ML | : * | 0 | % | [ 0 - 15] | |

**C2 Day 3**

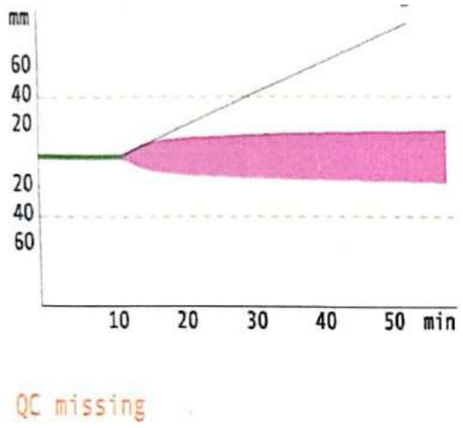


2 INTEM [default] 5, 5
1:5 2:5

RT: 00:59:34 ST: 2023-09-27T08:43:52

| | | | | | |
|---|---|---|---|---|---|
| CT | : | 732 | s | [ 100 - 240] | ▲ |
| CFT | : | | s | [ 30 - 110] | |
| α | : | 29 | ° | [ 70 - 83] | ▼ |
| A10 | : | 13 | mm | [ 44 - 66] | ▼ |
| A20 | : | 15 | mm | [ 50 - 71] | ▼ |
| MCF | : * | 18 | mm | [ 50 - 72] | |
| ML | : * | 0 | % | [ 0 - 15] | |

**C2 Day 2**

Figure: The Rotem test reveals a significant upregulation of the complement cascade in (EDHA) P on Day 3. Notably, this upregulation is more pronounced in EDHA P+2 at Day 3, demonstrating a marked increase compared to both EDHA P and P+2 at Day 2.

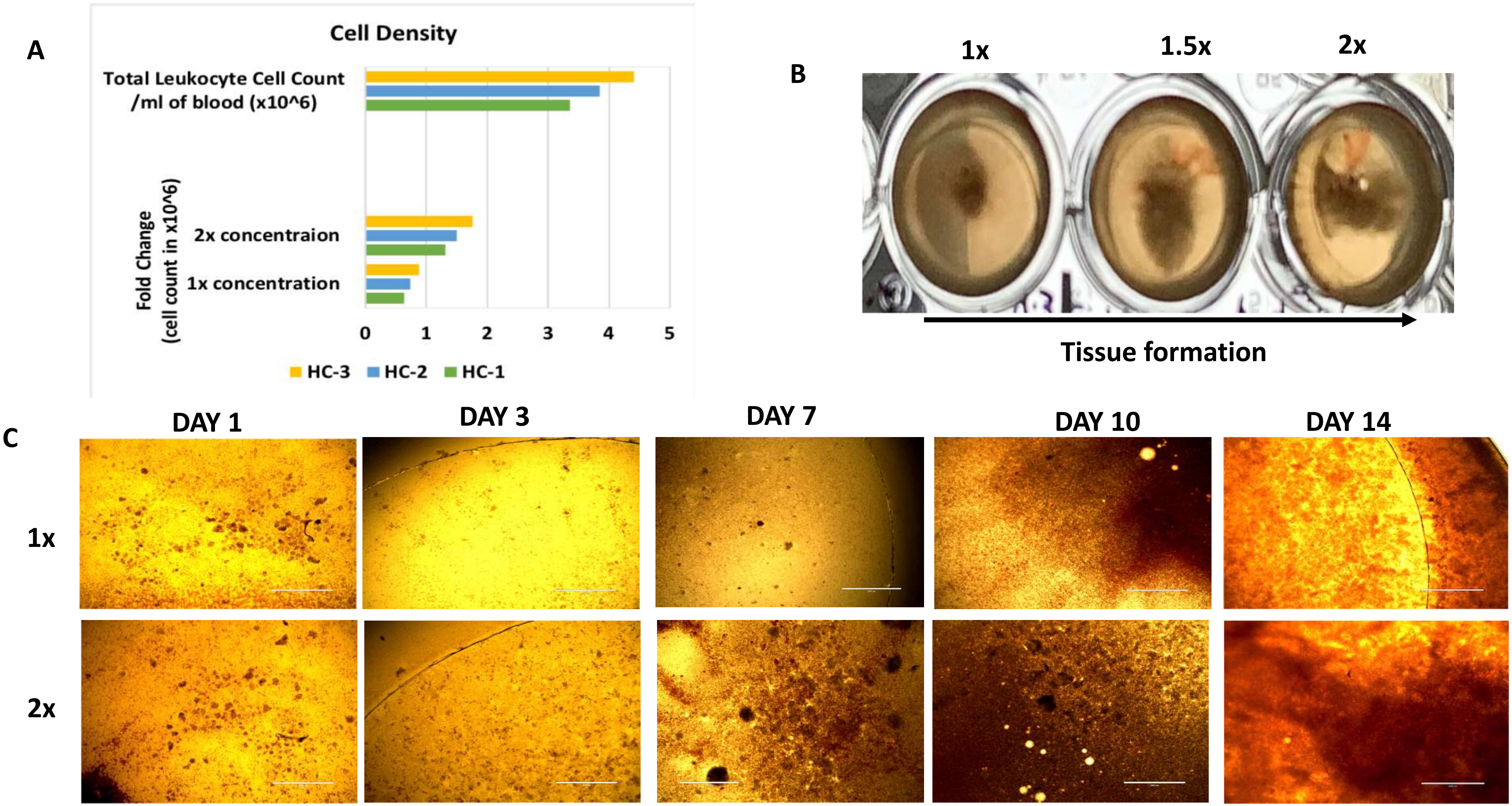
A
Cell Density
Total Leukocyte Cell Count
/ml of blood (x10^6)
Fold Change
(cell count in x10^6)
2x concentraion
1x concentration
0
1
2
3
4
5
HC-3
HC-2
HC-1
B
1x
1.5x
2x
Tissue formation
C
DAY 1
DAY 3
DAY 7
DAY 10
DAY 14
1x
2x

HC-1

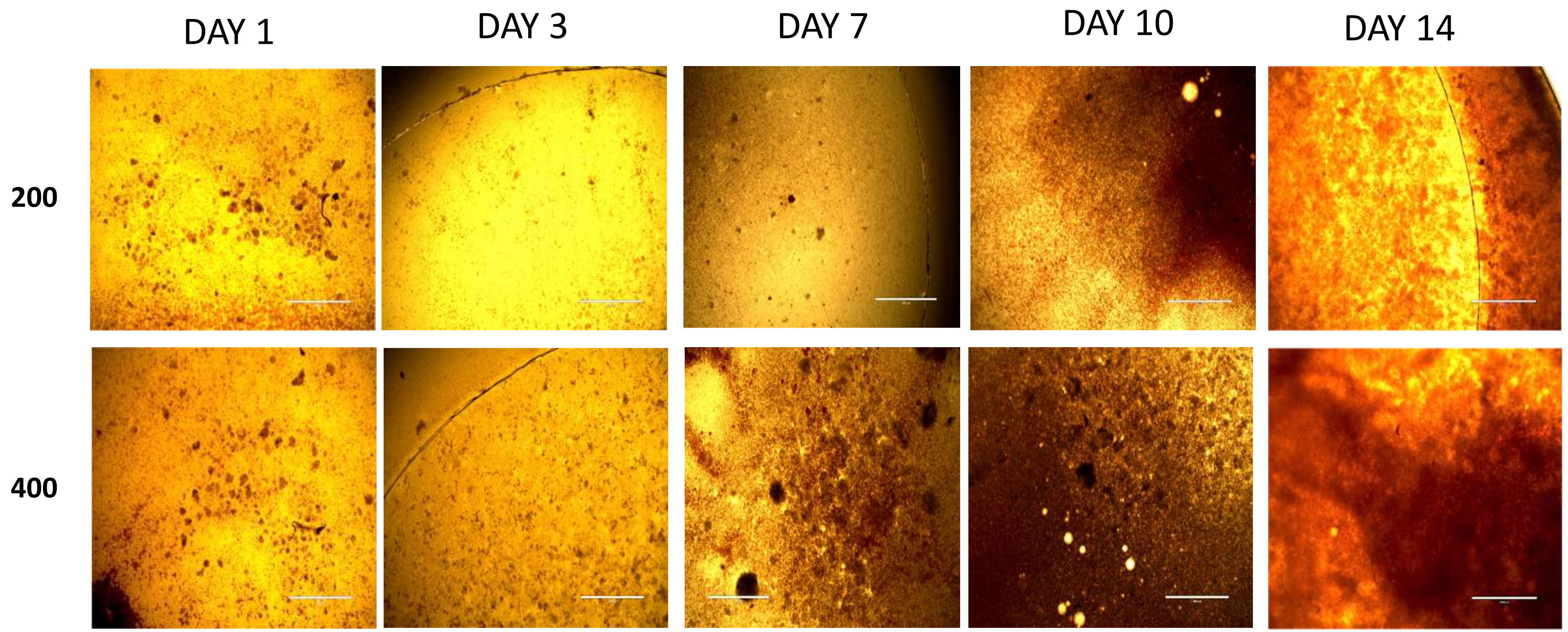

HC-2

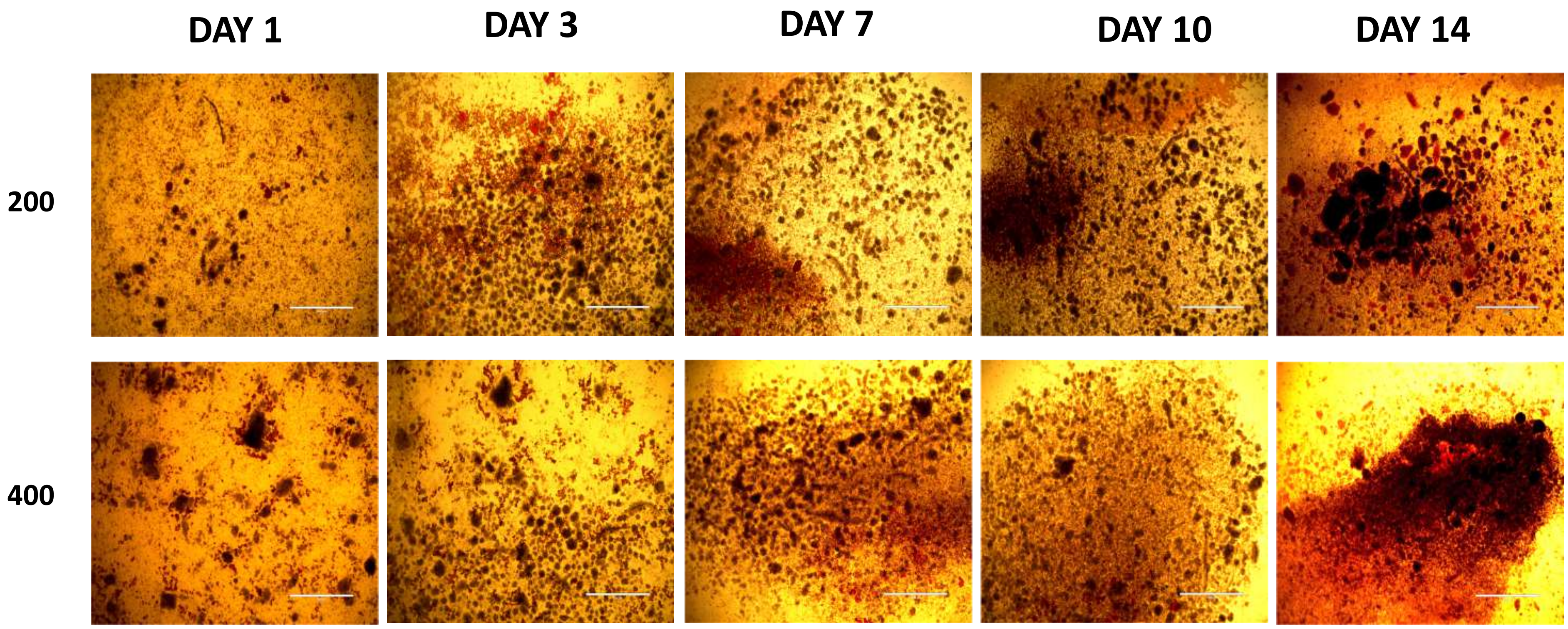

HC-3

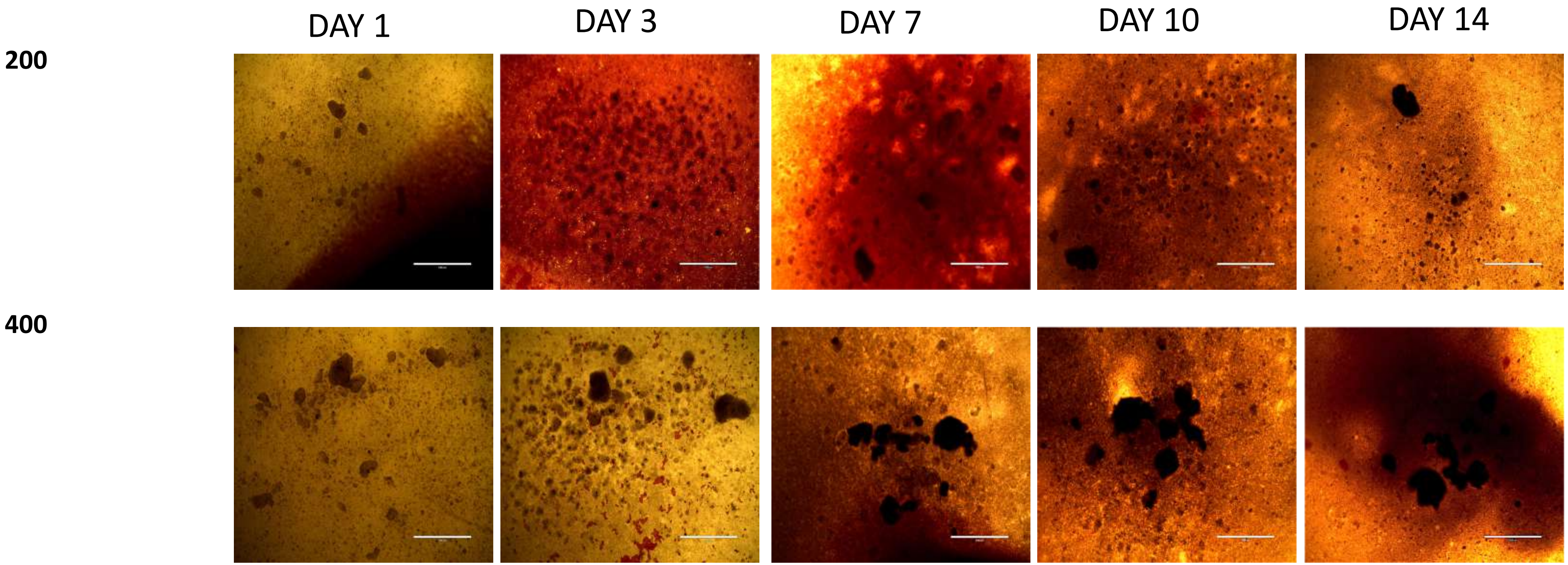
